# Comparative genomics analysis of three conserved plasmid families in the Western Hemisphere soft tick-borne relapsing fever borreliae provides insight into variation in genome structure and antigenic variation systems

**DOI:** 10.1101/2023.03.06.531354

**Authors:** Alexander R. Kneubehl, Job E. Lopez

**Affiliations:** Department of Pediatrics, Baylor College of Medicine, Houston, TX, USA; Department of Molecular Virology and Microbiology, Baylor College of Medicine, Houston, TX, USA

## Abstract

*Borrelia* spirochetes, causative agents of Lyme disease and relapsing fever (RF), have a uniquely complex genome consisting of a linear chromosome and circular and linear plasmids. The plasmids harbor genes important for the vector-host life cycle of these tick-borne bacteria. The role of Lyme disease causing *Borrelia* plasmids is more refined compared to RF spirochetes because of limited plasmid-resolved genomes for RF spirochetes. We recently addressed this limitation and found that three linear plasmid families (F6, F27, and F28) were syntenic across all species. Given this conservation, we further investigated the three plasmid families. The F6 family, also known as the megaplasmid, contained regions of repetitive DNA. The F27 was the smallest, encoding genes with unknown function. The F28 family encoded the expression locus for antigenic variation in all species except *Borrelia hermsii* and *Borrelia anserina.* Taken together, this work provides a foundation for future investigations to identify essential plasmid-localized genes that drive the vector-host life cycle of RF *Borrelia*.

**IMPORTANCE:** *Borrelia* spp. spirochetes are arthropod-borne bacteria found globally and infect humans and other vertebrates. RF borreliae are understudied and misdiagnosed pathogens because of the vague clinical presentation of disease and the elusive feeding behavior of argasid ticks. Consequently, genomics resources for RF spirochetes have been limited. Analyses of *Borrelia* plasmids have been challenging because they are often highly fragmented and unassembled. By utilizing Oxford Nanopore Technologies, we recently generated plasmid-resolved genomes for seven *Borrelia* spp. found in the Western Hemisphere. This current study is a more in-depth investigation into the linear plasmids that were conserved and syntenic across species. This analysis determined differences in genome structure and, importantly, in antigenic variation systems between species. This work is an important step in identifying crucial plasmid-borne genetic elements essential for the life cycle of RF spirochetes.

## INTRODUCTION

The plasmid content of the *Borreliaceae* is the most unique and complex among bacteria (1–5). No other bacterial organism harbors a repertoire of linear and circular plasmids, which are necessary for the completion of the microbes infectious cycle through the tick vector and vertebrate host (6–18). Extensive research has been conducted on the function of plasmids in *Borreliella* (*Borrelia*) *burgdorferi* (3-6, 19-22). This has proven important in the delineation of inter- and intra-species plasmid relationships and the identification of essential genes. Comparatively less work has been performed in tick-borne relapsing fever (RF) spirochetes because plasmid-resolved genomes for these microbes have been lacking.

We previously reported a comparative genomic analysis of seven species of Western Hemisphere soft tick-borne RF (WHsTBRF) spirochetes (1). A phylogenetic analysis of the PF32 plasmid partitioning loci identified 30 different plasmid families (F1 – F30). Of these, three (F6, F27, F28) were conserved and largely syntenic across the clade. The F6 plasmid family, also known as the megaplasmid, was the largest linear plasmid (110-194kb). The F27 plasmid was the smallest (10-12kb) encoding 12-14 genes of unknown function. The F28 plasmid family was related to the *B. burgdorferi* cp26 plasmid, which is essential is *B. burgdorferi*.

In this current study, we performed a comparative analysis of the F6, F27, and F28 plasmid families across the WHsTBRF spirochete clade. The species we evaluated were *Borrelia hermsii, Borrelia turicatae, Borrelia parkeri, Borrelia anserina, Borrelia coriaceae, Borrelia puertoricensis,* and *Borrelia venezuelensis.* This investigation revealed extensive repetitive gene content differences across the F6 megaplasmid family. Moreover, the F27 plasmid family was determined to be larger than previously reported due to an ancestral inverted duplication event. Interestingly, the F28 plasmid family contained the expression site for antigenic variation for all species except *B. hermsii* and *B. anserina*. This finding led to further investigation of the antigenic variation systems across the WHsTBRF spirochete clade. Collectively, this work is foundational for studies that will investigate the role of plasmid families in RF spirochete vector colonization, host use, vector specificity, and pathogenesis.

## RESULTS

### F6 (Megaplasmid) Plasmid Family Analysis

As the largest, non-chromosomal linear replicon found in RF spirochete genomes (23), the overall length of the F6 plasmid family varied. The largest megaplasmid was found in *B. hermsii* YOR at 194 kb and the smallest was in *B. anserina* BA2 at 88 kb. An analysis of the relatedness of WHsTBRF spirochetes’ megaplasmids by Mauve alignment indicated variable sequence conservation (**Fig. 1**). We identified three regions that were similar to those reported by Miller et al. (23). Starting from the 5’ end of the megaplasmid, region 1 was variable in length between all the species and contained *bbk32*/fibronectin-binding protein-type genes and multiple copies of complement regulator-acquiring surface protein genes (CRASPs). Region 2 contained *ribonucleoside-diphosphate reductase 2 subunit beta* (*nrdF*), *ribonucleoside-diphosphate reductase 2 subunit alpha* (*nrdE*), a *ribonucleotide reductase Class Ib* (*nrdI*), and plasmid partitioning genes (1). A gene encoding the *Borrelia immunogenic protein A* (*bipA*), an important diagnostic antigen (24–26), was found in region 2 of all isolates except for *B. anserina* BA2, which did not encode a homologue. Intact phage-related genes were also found in region 2 (e.g. *PBSX family phage terminase large subunit*, *multi-copy lipoprotein* [*mlp*] family, and *blyA* family holin genes). Region 3 was variable in length and gene content between all species and contained repetitive sequences. Overall, the three regions of conserved gene content varied in length across the WHsTBRF spirochete megaplasmids.

**Fig. 1.**
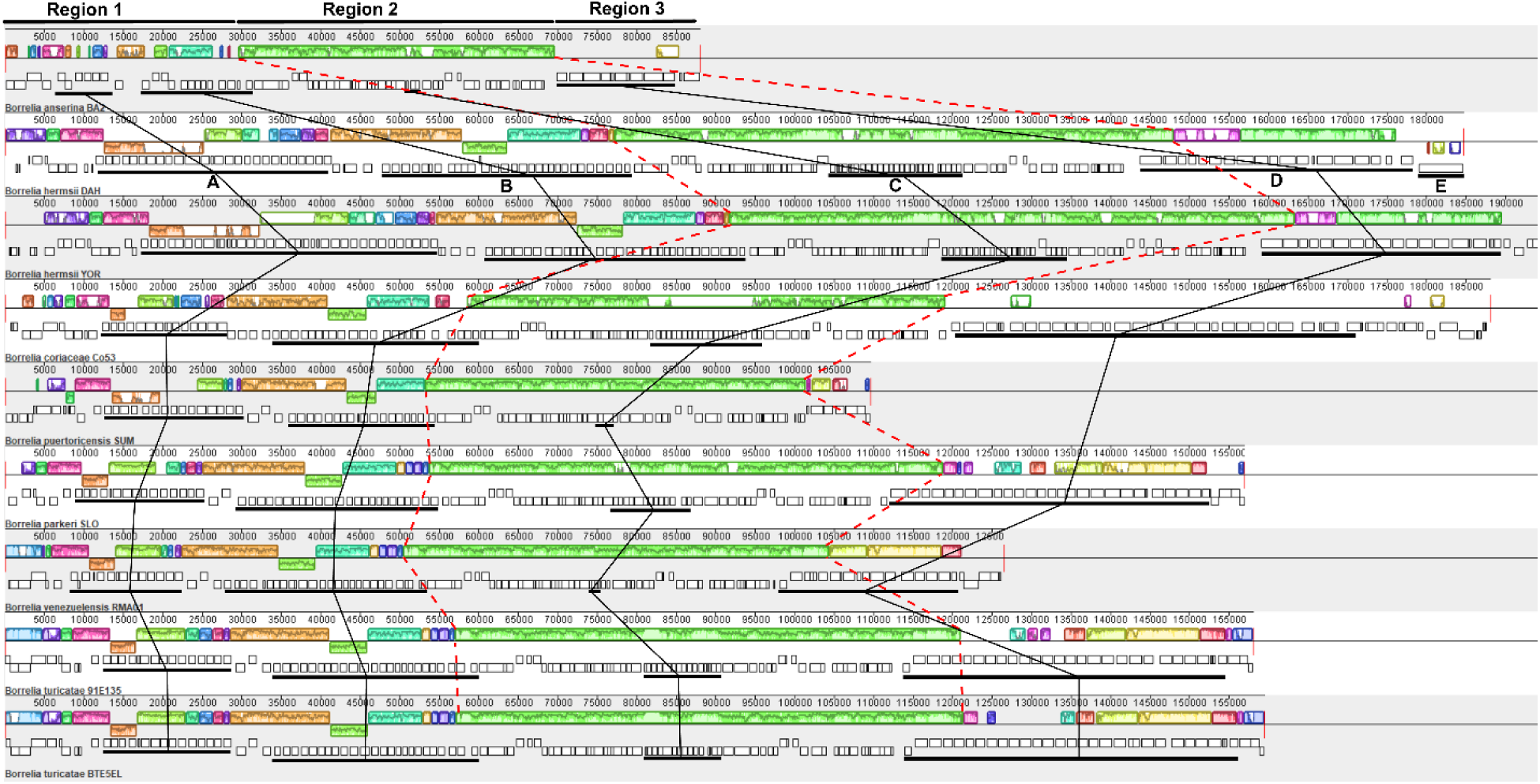
Mauve alignment of WHsTBRF megaplasmids (F6 Family). Three regions of interest are highlighted, which correspond to areas of variability and conservation. Red dashed lines were used to show where region 2 starts and stops for each isolate. The repetitive regions previously defined in *B. hermsii* are indicated on the *B. hermsii* DAH sequence and black lines indicate where those regions are in each genome (23). Colored boxes are shown indicating areas of collinearity between isolates where similarly colored boxes correspond to each other across taxa (histograms within these boxes correspond to the level of nucleotide similarity, the higher the bar the higher the similarity). Colored boxes below the mid-line correspond to inversions compared to the reference sequence (*B. anserina* BA2). Genes are shown for each isolate, below the collinearity boxes, with genes above the mid-line in the positive-sense and those below in the negative-sense.

The variation observed in the megaplasmids was primarily due to repetitive nucleotide sequences. This was reported in *B. hermsii* where five areas of genes with a high degree of repetitive nucleotide sequence were identified (23). We applied the designations of repetitive nucleotide blocks A-E across the WHsTBRF clade (**Fig. 1**).

The designated genes in blocks A-E are indicated in **Supplemental File 1** with their InterProScan classifications. Repetitive blocks A and B were found in what we designated as region 1, block C is in region 2, and blocks D and E are in region 3 (annotated in **Fig. 1**). Self-dot plot analysis revealed that the presence of these repetitive blocks was variable across the WHsTBRF clade (**Fig. 2**).

**Fig. 2.**
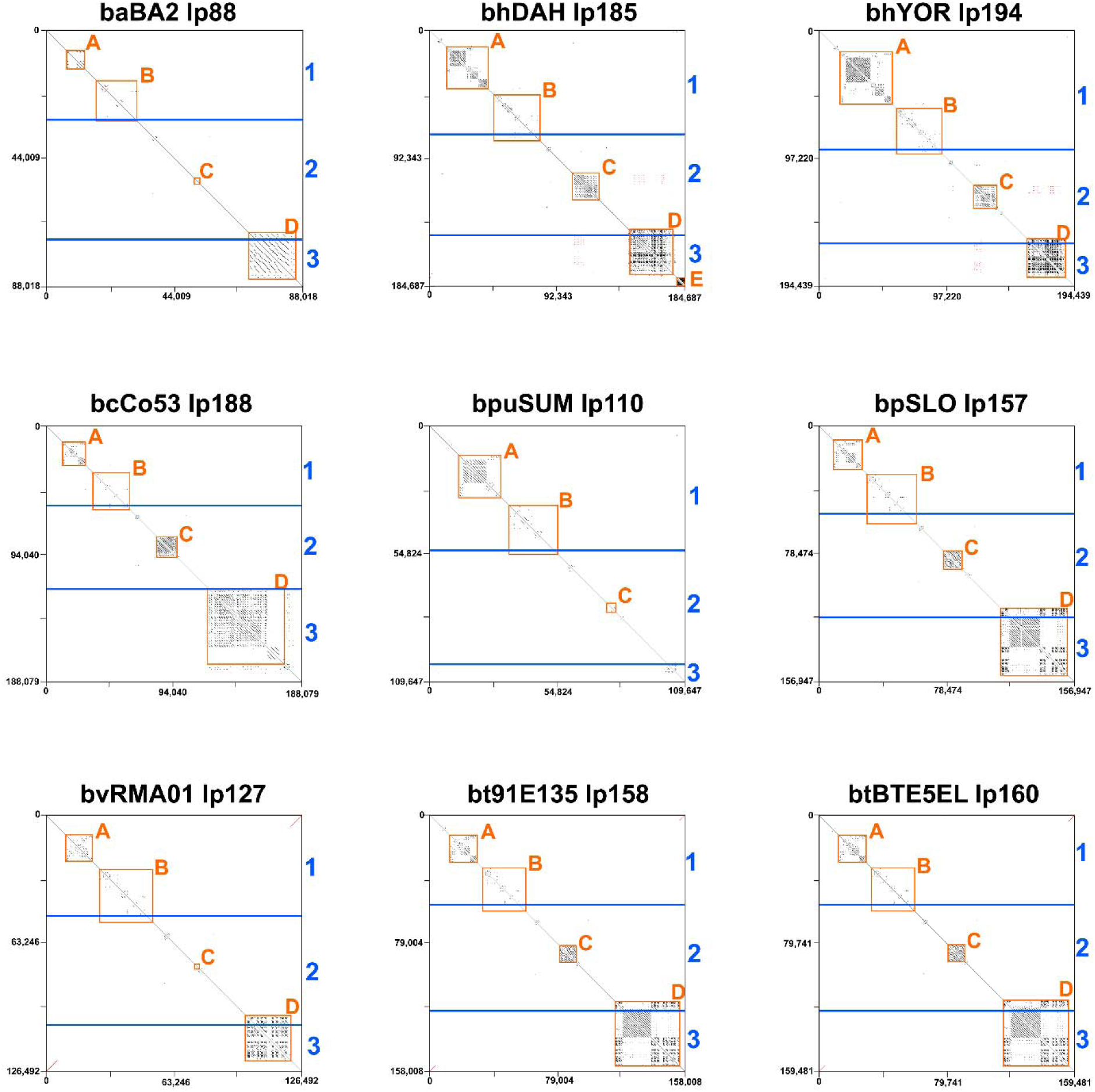
Self-dot plot of the megaplasmids. Self-dot plots were generated to observe differences in the repetitive gene content across the clade. Black lines from to left to bottom right indicate areas of direct repeats. Red lines from bottom left to top right indicate areas of inverted repeats. The blue lines and numbers indicate the locations of the three conserved regions in the megaplasmids. The orange boxes and numbers indicate the locations of the five repetitive gene blocks.

Block A was variable in length across species due to the number of CRASP- related genes. The size of this block ranged from ∼6kb in *B. anserina* BA2 to ∼37kb in *B. hermsii* YOR and was the largest in *B. hermsii* DAH and YOR and *B. puertoricensis* SUM (**Fig. 2**). The genes located here were identified by InterProScan as containing a Bbcrasp-1 domain (PF05714). On average there were 16 genes containing a Bbcrasp-1 domain. *B. anserina* BA2 had the least with six genes, while *B. hermsii* YOR had the most with 30 (**Supplemental File 1**). Interestingly, the number of genes was not associated with the number of nucleotide repeats. For example, block A of *B. coriaceae* Co53 and *B. puertoricensis* SUM had similar numbers of BbCRASP-1 domain containing genes. However, block A of *B. coriaceae* Co53 contained fewer repetitive sequences compared to *B. puertoricensis* SUM (**Fig. 2**).

Block B was evident across all species and isolates and ranged between ∼14kb in *B. anserina* BA2 to ∼33kb in *B. hermsii* YOR (**Fig. 2**). The genes found in block B were typically classified by InterProScan or GenBank’s Prokaryotic Genome Annotation Pipeline (PGAP) as a hypothetical protein. However, InterProScan classified one BbCRASP-1 domain containing gene in *B. puertoricensis* SUM, two in *B. venezuelensis* RMA01, three in *B. parkeri* SLO, and one each in *B. turicatae* 91E135 and BTE5EL (**Supplemental File 1**).

Repetitive nucleotide sequences in Block C were apparent in all species except *B. anserina* BA2, *B. puertoricensis* SUM, and *B. venezuelensis* RMA01 (**Fig. 2**). The size of block C ranged from ∼1.6kb in *B. anserina* BA2 to ∼16.5kb in *B. hermsii* DAH. The majority of genes found in this repetitive block were annotated by InterProScan as containing a domain related to *bbe16* also designated the *borrelial persistence in ticks protein A* (*bptA*) (14). This gene is a single copy, essential gene in *B. burgdorferi* (14), and we determined that two to 14 copies of *bptA* were present in all WHsTBRF spirochete isolates (**Supplemental File 2**). Phylogenetic analysis of the nucleotide sequence of these putative *bptA* genes, including two genes from *B. burgdorferi* BB31 that contained a *bptA* domain (GCF_000008685.2_ASM868v2), identified two clades (**Fig. 2 and Fig. S1)**. One clade contained both *B. burgdorferi bptA* and a known related protein in *B. burgdorferi, bbj47* (27). Each WHsTBRF spirochete assembly had one gene that clustered with *bbj47*. The other clade contained all other WHsTBRF genes that contained a *bptA* domain. These genes were highly divergent from the *B. burgdorferi bptA* (*bbe16*), with apparent duplication events occurring within each species (**Fig. S1**). Interestingly, *bptA*-like genes from *B. hermsii* isolates clustered to the exclusion of all other species’ *bptA*-like genes. The species that were missing repetitive nucleotide sequences in block C also had the fewest number of *bptA*-like genes (**Fig. S1**). Moreover, some of these genes were substantially truncated (>50% of sequence missing) and are likely pseudogenes. We did not considered these genes in our phylogenetic analysis (bhDAH_001245, bhDAH_001246, bhDAH_001248, bhDAH_001249, and bvRMA01_000992).

Block D was the most variable block in both in sequence similarity and length. This block ranged in size from non-existent in *B. puertoricensis* SUM to ∼53kb in *B. coriaceae* Co53. The amount of repetitive nucleotide sequence in block D of *B. coriaceae* Co53 was less compared to other species (e.g. *B. turicatae* and *B. parkeri*) (**Fig. 2**). In both *B. hermsii* isolates and in *B. coriaceae* Co53 the genes in this block were annotated as hypothetical proteins and not classified by InterProScan (**Supplemental File 1**). *Borrelia anserina* BA2 had only one classified gene in block D, a phage fiber gene. *Borrelia parkeri* SLO had three genes classified, DUF1617, a *collagen-like* gene, and a *trimeric UDP-N-acetylglucosamine acetyltransferase* (*lpxA*)- like enzyme family. *Borrelia venezuelensis* RMA01 had a single gene that was classified, which contained a conserved domain in the phosphatidylinositol phosphate kinase (PIPK) catalytic family. Lastly, both *B. turicatae* isolates contained only two genes classified as DUF1617.

Repetitive nucleotide block E was only present in *B. hermsii* DAH with a single gene (**Fig. 2**). *bhDAH_001311* is a large hypothetical gene (5,513bp) coding for an ∼210kda protein. BLASTn and BLASTp analysis of this gene failed to identify homologs outside of *Borreliaceae* (28). Within the WHsTBRF spirochetes, BLASTp found similar proteins (∼40% sequence similarity, 99% query coverage, e-value= 0.0) in *B. puertoricensis* SUM (bpuSUM_001813, linear plasmid 46 ‘lp46’), *B. parkeri* SLO (bpSLO_001255, lp28), and in *B. venezuelensis* RMA01 (bvRMA01_001103, lp25). Interestingly, these plasmids are not in the same plasmid family (1). Further, BLASTn nor BLASTp did not detect genes or proteins similar to *bhDAH_001311* in *B. hermsii* YOR or any other *B. hermsii* genomic group II sequences deposited in GenBank.

### F27 Plasmid Family Analysis

The F27 plasmid family was the shortest linear plasmid family found in all WHsTBRF species (1). The size ranged between ∼10kb to ∼12kb. Previously, this family of plasmids was reported to be either a circular or a linear 5 – 6kb plasmid (29–31). However, using long-read sequencing data and manual inspection, we determined that the F27 plasmid family is linear with evidence of an inverted duplication event (**Fig. S2A-C**). Our rationale that the F27 plasmid family was linear was three-fold. First, contigs that were assembled for this plasmid family often had at least one complete and one incomplete telomeric sequence, which would be uncommon for a circular or linear plasmid (**Fig. S2A-C**). Second, we observed that the longest Oxford Nanopore Technology (ONT) reads of the F27 plasmids were approximately twice the size of the final plasmid sequence. These lengths suggested linearity because the telomeres of linear plasmids are covalently linked and both the positive and negative sense strands would be sequenced. This would cause long inverted telomeric repeats (**Fig. S2A-C**) (32, 33). Third, dot plot analysis of the longest reads showed patterns consistent with a plasmid that was fully sequenced around the telomeres. Interestingly, we noticed a “wavey” appearance in the dot plot of the contigs and reads in this plasmid family, which indicated a physical DNA molecule translocation issues during sequencing (**Fig. S2A, B, and D**). Indeed, inverted duplication events can cause increased translocation rates in ONT sequencing (34). This occurs through the formation of secondary structures of the translocated DNA, which causes decreased sampling rates and lower basecall accuracy. The sequence at the inverted duplication junction that disrupted the primordial telomere was noncoding and variable in length and sequence identity across species. The disruptive sequence was not found anywhere else on the F27 plasmid or in any other replicon in that plasmid’s genome.

Alignment of the F27 plasmid family showed that the plasmids was conserved across species (**Fig. 3**). Interestingly, the plasmid partitioning genes in this plasmid family were in a unique configuration compared to other plasmid families and lacked PF50 genes (1). Aside from the plasmid partitioning genes, there were five to seven other predicted genes on the F27 plasmids that were annotated as hypothetical proteins. While the genes were conserved across species, the putative role of this plasmid and its genes remained elusive.

**Fig. 3.**
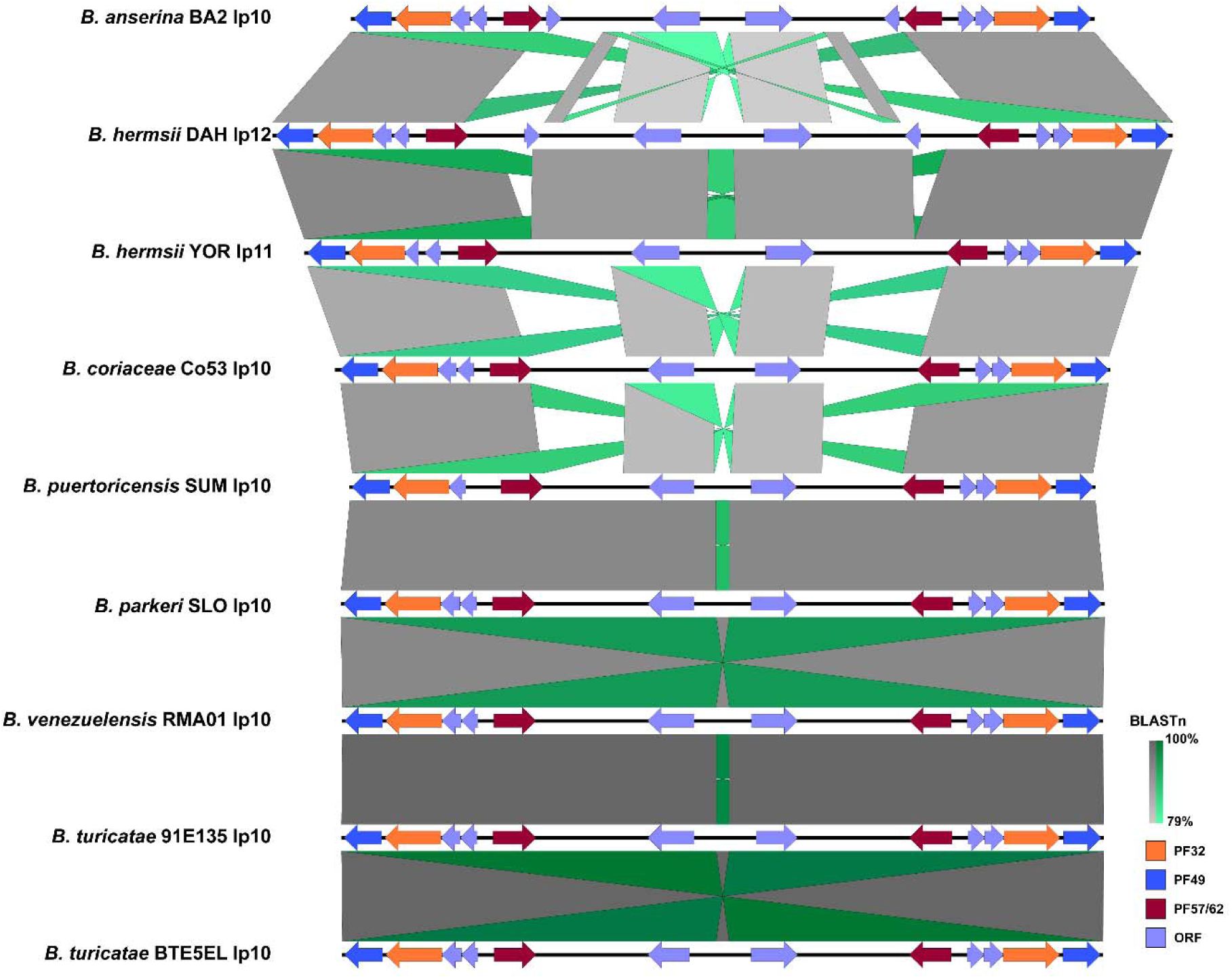
Alignment and visualization of the F27 plasmid family. The minimum BLASTn alignment was set to 250nt. BLASTn results scale from 79-100% going from light to dark grey. Nucleotide inversions scale similarly but from light to dark green. Open reading frames (ORFs) and plasmid partitioning loci are indicated by specific colors.

### F28 Plasmid Family Analysis and Comparing Antigenic Variation Systems in WHsTBRF Spirochetes

Lyme disease and RF causing spirochetes employ antigenic variation systems to evade host immunity, and this has been extensively studied for RF spirochetes in *B. hermsii* (35–37). The protein family driving antigenic variation for RF spirochetes is designated the variable major proteins (Vmps) (38). Antigenic variation is achieved through the recombination of silent, archived genes into a single expression site (39, 40). Since there is only one *vmp* expression locus in a single bacterium, we refer to both the expressed and silent archived *vmp*s as ‘alleles’ based on the nomenclature put forth by Rich et al. (41). Our classification of *vmp*s includes predicted pseudogenes as annotated by PGAP. The predicted protein products of pseudogenes were on average ∼40-50% shorter than the length of predicted Vmp alleles. Furthermore, Vmps are classified by molecular weight, with the variable small proteins (Vsps) at ∼22 Kda and variable large proteins (Vlps) ∼37 Kda (42–44). The antigenic variation expression site in *B. hermsii* uses a σ70-like promotor (45), and is characterized by upstream genetic elements that include three stem loop structures and a transcription-enhancing tract of ∼12 to 16 thymines (40, 46, 47). In our analysis, the *B. hermsii* promoter sequence and the thymine tract were found exclusively on the *B. hermsii* F20 plasmids, which were lp26 and lp23 plasmids for DAH and YOR isolates, respectively (**Fig. S3**). We also identified the upstream and downstream homology sequences (UHS and DHS respectively) on these two plasmids (**Fig. S3**). The UHS and DHS were previously identified as the two areas of sequence homology that allow directed recombination to occur in *B. hermsii* (40). However, analysis of the other WHsTBRF spirochete F20 plasmids indicated this plasmid family was not syntenic and we were unable to identify a similar *vmp* expression site in other species (**Fig. S4**).

Since *B. turicatae* utilizes an *B. burgdorferi ospC-*like promoter to express *vmp*s (48, 49), we analyzed the genomes for the presence of this promotor to identify the expression sites. The promoter sequence was identified on the F28 plasmids (**Fig. 4A**). Interestingly, for *B. hermsii* the promoter on these plasmids regulates the expression of the variable tick protein (*vtp*) gene (further discussed below). Sequence alignments determined that the -35 and -10 sites and the ribosomal binding site were largely conserved, with the -10 site possessing nucleotide variability (**Fig. 4A**). The -10 site was conserved in *B. puertoricensis* SUM, *B. parkeri* SLO*, B. venezuelensis* RMA01, and *B. turicatae* 91E135 and BTE5EL. However, these species also had the pseudo -35 site immediately upstream of the -10 site, which was hypothesized to cause RNA polymerase holoenzyme mis-alignment thereby potentially requiring a trans-activating protein (45). As previously described for *B. turicatae* (48, 50, 51), the promoter was downstream of an oligopeptide permease-like protein gene for all genomes except *B. anserina* BA2, which was missing this gene.

**Fig. 4.**
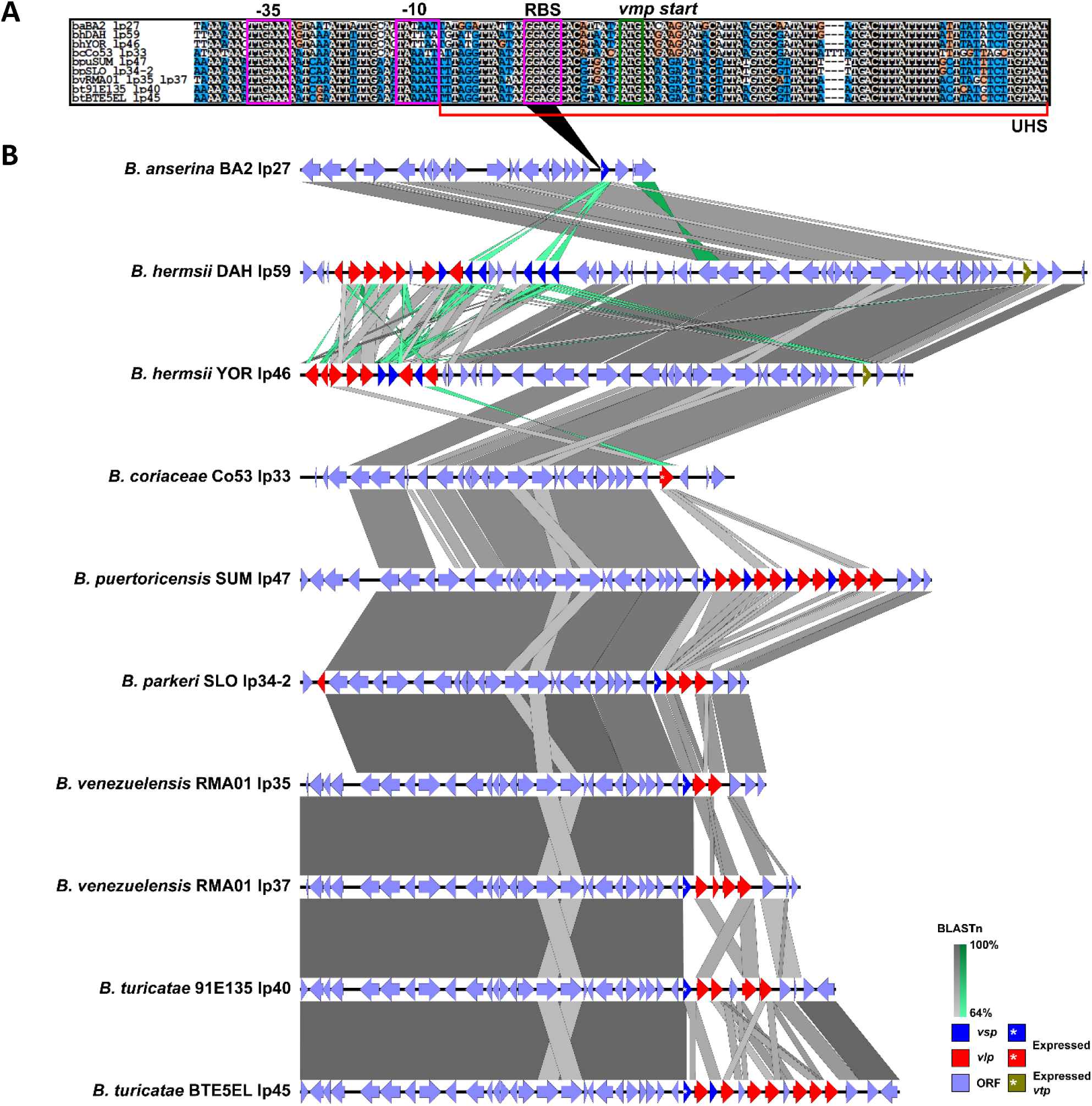
Alignment and visualization of F28 plasmid family and *vmp* promoter. The *vmp* promoter alignment is seen in the top sequence (**A**). Promoter features are annotated and boxed in purple (RBS= ribosome binding site). The ATG start codon is boxed in green. The upstream homology sequence (UHS) is indicated with a red bracket. For the alignment, a white text in a black square means the sequences are the same, black text in a white square indicates sequences are different, black text in an orange box indicates a transition substitution, and white text in blue squares indicates that that nucleotide matches the consensus for that position. The plasmids were aligned using EasyFig (**B**). This analysis demonstrated sequence conservation and synteny across species. The minimum BLASTn alignment was set to 250nt. BLASTn results scale from 64-100% going from light to dark grey whereas inversions scale the same but from light to dark green. Open reading frames (ORFs) and the *vmp* alleles are shown in specific colors. The *vmp* in the expression site is indicated by an asterisk within the indicated ORF.

We also attempted to identify sequence analogous to the UHS and DHS of the *B. hermsii* F28 plasmids. We detected an UHS across the species (**Fig. 4A**). However, we did not detect DHS sequence. The sequence downstream of the expression site, including the *vmp* loaded there, was nearly identical to other plasmids for all genomes except *B. coriaceae* Co53 (**Table 1**). The plasmids that had sequence similar to the expression plasmid did not fall into any particular plasmid family (**Table 1**), nor did they contain a complete promoter upstream of the *vmp* found in the F28 *vmp* expression site. In *B. coriaceae* Co53, we did not find sequence in the region where the DHS would be that had sequence identity >90% to other parts of its genome. We were also not able to find the archived gene of the *vlp* in the *B. coriaceae* Co53’s putative *vmp* expression site.

**Table 1.** Plasmids with Similar Sequence Downstream of the F28 Expression Site

| Genome | F28 Plasmid | Plasmid with Similar Sequence | Blastn % Sequence Coverage | Blastn % Sequence Identity | Plasmid Family |
| --- | --- | --- | --- | --- | --- |
| *B. coriaceae* Co53 | lp33 | none | N/A | N/A | N/A |
| *B. puertoricensis* SUM | lp47 | lp39 | 96 | 99.97 | F16 |
|  |  | lp46 | 100 | 99.96 | F14 |
|  |  | lp57 | 98 | 99.37 | F18 |
|  |  | lp65 | 100 | 99.98 | F1 |
| *B. parkeri* SLO | lp34-2 | lp34-1 | 99 | 99.97 | F7 |
|  |  | lp60 | 99 | 99.96 | F29 |
| *B. venezuelensis* RMA01 | lp35 | lp30 | 100 | 100 | F12 |
|  |  | lp31 | 100 | 97.13 | F10 |
|  | lp37 | lp25 | 97 | 99.82 | F14 |
|  |  | lp30 | 99 | 99.97 | F12 |
| *B. turicatae* 91E135 | lp40 | lp38 | 100 | 99.96 | F19 |
| *B. turicatae* BTE5EL | lp45 | lp28 | 99 | 99.94 | F14 |

N/A=not available

There were additional structural differences in the *vmp* expression locus between species of RF spirochete. Similar to what was reported for *B. turicatae* (48, 50, 51), the putative *vmp* expression site located on the F28 plasmids was internal with *vmp* alleles and non-*vmp* genes downstream (**Fig. 4B**). This is contrary to the *B. hermsii vmp* expression site on the F20 plasmids, which is terminal on the telomere and only one *vmp* is present and downstream genes are absent (**Fig. S3**) (40).

Intriguingly, analysis of the F28 family showed that *B. venezuelensis* RMA01 had two nearly identical plasmids in the F28 family, lp35 and lp37. Both *B. venezuelensis* RMA01 F28 plasmids contained the o*spC-*like expression site. However, lp37 had a pseudogenized *vsp*, caused by a point mutation resulting in a frameshift in this gene, whereas lp35 had an intact *vsp* gene. The *vmp* alleles located downstream of the expression site for lp35 and lp37 are different in number and sequence, and both plasmids differ at the 3’ ends. The 3’ end of lp35 matches the 5’ telomeres of lp30 and lp31. The 3’ end of lp37 matches the 3’ end of lp30 and 5’ telomere of lp25. These data suggest a recombination events occurred that replaced the entirety of the 3’ end rather than just *vmp* loci downstream of the expression site.

We confirmed the presence of the two 3’ end configurations for *B. venezuelensis* RMA01 F28 plasmids. Mapping of long reads that were initially used for the assembly of the *B. venezuelensis* RMA01 genome identified reads specific to both 3’ end configurations (**Fig. S5**). Adaptive sampling using the ONT MinION platform was also performed to selectively sequence the lp35 and lp37 plasmids of *B. venezuelensis* RMA01 and the related lp40 plasmid of *B. turicatae* 91E135. Adaptive sampling data supported the two different 3’ end configurations of *B. venezuelensis* RMA01’s lp35 (**Fig. S6)** and lp37 (**Fig. S7**) plasmids, while only one 3’ end configuration was present in *B. turicatae* 91E135’s lp40 plasmid (**Fig. S8**). Taken together, these data confirmed that *B. venezuelensis* RMA01 contained two nearly identical plasmids with the *vmp* expression locus but differed in their 3’ ends.

We further investigated the antigenic variation systems of the WHsTBRF spirochetes by determining the number of *vmp* alleles for each isolate. The number varied among species but *vlp* alleles outnumbered *vsp* alleles in every genome (**Fig. S9**). A phylogenetic analysis of *vmp* alleles determined relatedness within *vsp* and *vlp* groups. We did not observe phylogenetic structure in *vsp* alleles, which agreed with what Kuleshov, et al. reported for *B. miyamotoi vsp* alleles (**Fig. S10A**) (29). Phylogenetic analysis of the *vlp* alleles separated them into the four previously described subfamilies (alpha, beta, gamma, delta) (52). The exception was *B. anserina* BA2, which did not have *vlp* alleles (**Fig. S10B**). We analyzed each isolate individually to determine the subfamily designation for each *vlp* allele (**Fig. 5, Supplemental File 4**). The proportions of each *vlp* subfamily indicated that the beta subfamily *vlp* alleles remained relatively constant while the alpha, gamma, and delta subfamilies varied between isolates (**Fig. S11**).

**Fig. 5.**
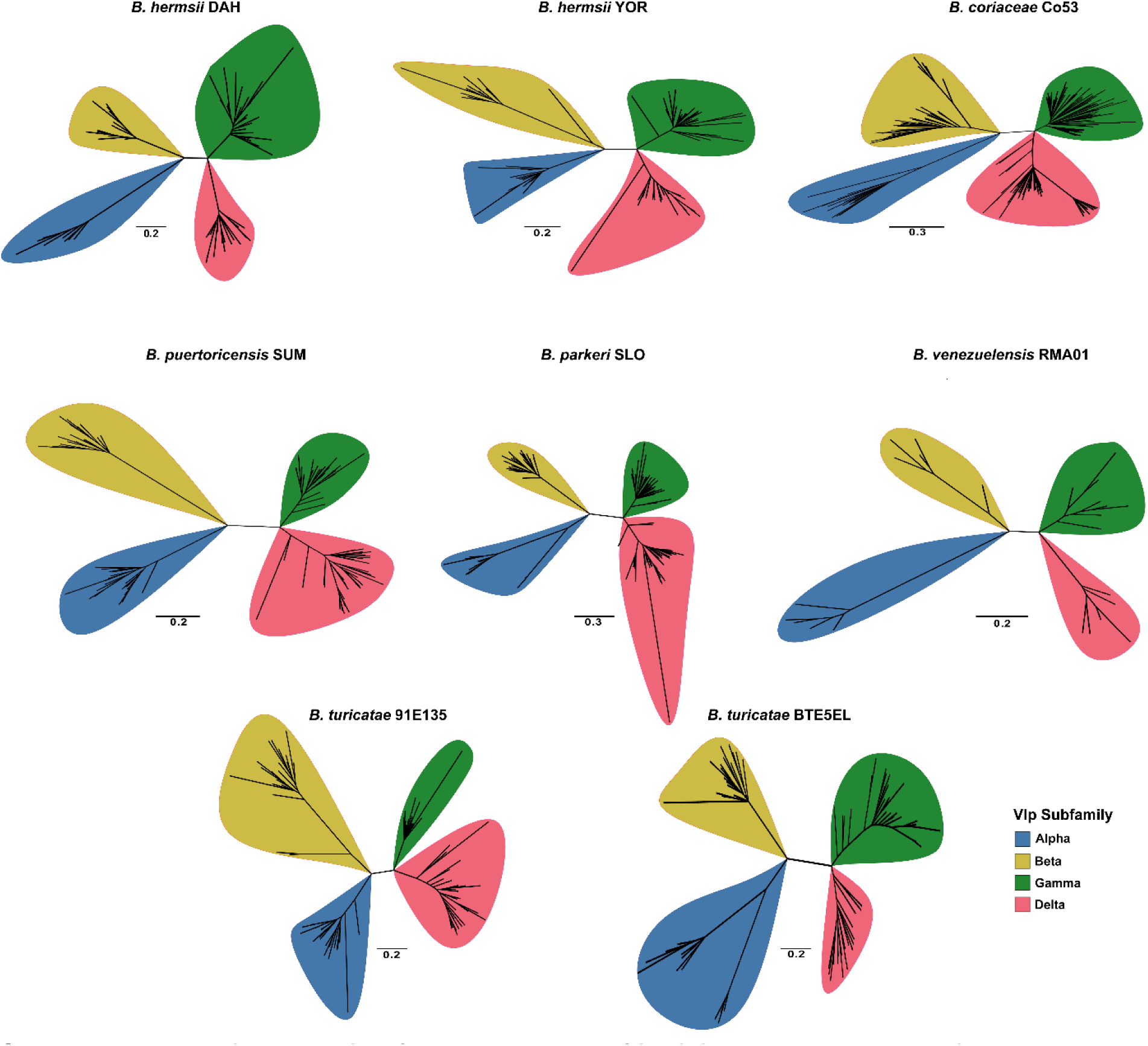
Phylogenetic analysis of the *vlp* alleles of individual WHsTBRF spirochete isolates. Each isolate’s *vlp* allele nucleotide sequences were used to infer a maximum likelihood tree with 1,000 bootstraps. Branches with less than 50% support were collapsed. The scale bar indicates substitutions per site. Note, *B. anserina* BA2 did not have any *vlp* alleles so it is not shown here.

### Assessment of the Variable Tick Protein (Vtp) Expression Locus Between Species

The F28 plasmids in *B. hermsii* contained the variable tick protein (Vtp) expression site (53, 54), and we performed an analysis to identify orthologs in other species of WHsTBRF spirochetes. The *vtp* is a *vsp* with a unique signal peptide sequence (53), exists in only one copy in the *B. hermsii* genome (45), and is controlled by a different promoter compared to the antigenic variation system (55). The *vtp* is expressed during spirochete colonization of the salivary glands (54), during early mammalian infection (56), and is crucial for infecting the host (17). We assessed our *vsp* phylogeny to see if we could detect a difference in the phylogenetic relationship between *vtp* genes (bhDAH_001490 and bhYOR_001299) and other *vsp* alleles (**Fig. S10A**). There was no discernable difference. This was not surprising since the only major difference between Vtps and Vsps is the signal peptide sequence (42, 53). Consequently, we evaluated the presence of Vtp in the WHsTBRF spirochetes in our dataset through cluster analysis of signal peptides (first ∼20 amino acids) of Vsps. Only the *B. hermsii* Vtp, *B. burgdorferi* OspC, and *B. anserina* BA2 Vsp signal peptides clustered together indicating that there were no other Vtp-like signal peptides that had at least a >80% amino acid sequence identity. The Vsp from *B. anserina* BA2 had 90% similarity to the OspC signal peptide. This similarity between the OspC and the *B. anserina* Vsp signal peptides was consistent to findings reported by Schwan and co-workers (15).

## DISCUSSION

We investigated the F6, F27, and F28 plasmid families because they were conserved in all species of WHsTBRF spirochetes with genomes currently available. The F6 family (megaplasmid) is the largest linear plasmid reported for RF spirochetes, while the F27 plasmid family is the smallest plasmid (∼10 kb) and had no close relative in the LD spirochetes (1). We also characterized the F28 plasmid family, which encoded the *vtp* gene of *B. hermsii*; however, in the other species, these plasmids contained the expression locus for antigenic variation. Collectively, our findings identified distinctions in the repetitive gene content and antigenic variation systems across the species of WHsTBRF spirochetes.

In RF spirochetes, the F6 megaplasmid is hypothesized to be important for tick colonization and mammalian infection based on its gene content and transcriptional profile in tick vector and mammalian host (23, 57, 58). For example, a gene expression analysis of the *B. turicatae* megaplasmid indicated that ∼67% of the genes found on the megaplasmid were up-regulated when grown under conditions mimicking the tick environment (57). Further analysis validated the up-regulation of genes localized at the 3’ end of the megaplasmid in the tick compared to infected mouse blood. Additionally, the megaplasmid is related to the essential lp54 plasmid in LD causing spirochetes (4, 22). Lp54 encodes genes, such as *outer surface lipoproteins A* and *B*, *cspA*, and *decorin binding proteins A* and *B*, which play important roles in tick colonization and vertebrate infection (59–63).

Previous work with the megaplasmid identified three regions of conservation with five blocks of repetitive nucleotide content in *B. hermsii, B. turicatae,* and *Borrelia duttonii* (23). Our work supported these findings for the additional species of WHsTBRF spirochete evaluated. However, the length of the three conserved regions varied across the species, which was driven by the number of repetitive sequences in the five blocks. Two repetitive regions that stood out were blocks A and C. For example, block A of the megaplasmid contains homologs of *B. burgdorferi* CRASP genes. In *B. burgdorferi*. the proteins are important for complement resistance in the vertebrate host (64–67).

Interestingly, these genes were variable in number and repetitive sequence across the WHsTBRF spirochetes. *Ornithodoros* species are known to feed on a variety of vertebrate hosts (68–73), and the observed variability in CRASP homologs may contribute to different vertebrate host use and complement resistance traits.

Repetitive region C was variable across isolates and contained multiple copies of *bptA* homologs. BptA is a surface exposed lipoprotein that is essential for persistence of *B. burgdorferi* in *Ixodes scapularis* (14). Multiple copies of *bptA*-like genes were found in prior work on the megaplasmids of *B. hermsii* and *B. turicatae*, however, their functional role remains uninvestigated (23). Previous investigations using microarray analysis of the megaplasmid in *B. turicatae* reported over a 3-fold up-regulation of expression of these genes in tick-like *in vitro* growth conditions relative to spirochetes isolated from mammalian blood (57). Transcriptional profiles and the gene’s duplication and diversification in WHsTBRF spirochetes suggest that they may be important in vector colonization. Indeed, gene duplication and diversification has been linked to environmental adaption in other prokaryotes (74, 75). The megaplasmid in RF spirochetes warrants further investigation to determine its role in the infectious life cycle of these pathogens.

Our analysis also indicated that the F27 family is also conserved in species of WHsTBRF spirochetes. In prior relapsing fever spirochete genome assemblies, F27 plasmids were often reported as a 5-6kb circular plasmid (*B. turicatae* 91E135 GCA_000012085.2, *B. hermsii* HS1 GCA_001660005.1, and *B. anserina* Es GCA_001936255.1). However, a plasmid analysis of WHsTBRF spirochetes using pulsed-field gel electrophoresis indicated that every genome possessed a ∼10 to 12 kb linear plasmid and a phylogenetic analysis determined that this small plasmid clustered into the F27 family (1). Newer third generation sequencing technology and assembly strategies demonstrated that the F27 plasmid family was not only linear but has undergone an inverted duplication event. While the F27 plasmid is conserved across species, the overall function is elusive.

The F28 plasmid family contains housekeeping genes and the putative expression site for antigenic variation for all species of the WHsTBRF clade except for *B. hermsii* and *B. anserina* (6, 76–79). Interestingly, compared to the terminal *vmp* expression site on *B. hermsii* F20 plasmids, the *vmp* expression sites on the F28 plasmids were internal, as previously reported for *B. turicatae* (51). Downstream of the F28 plasmid expression site were multiple *vmp* alleles and other non-*vmp* genes. Dai et al. previously reported that multiple *vmp*s can be present downstream of the F20 plasmid family’s antigenic variation expression site in *B. hermsii,* but that further recombination events take place to remove these alleles leaving only one *vmp* (40). The *vmp* gene in the expression site is directly adjacent to the telomere of the F20 plasmid. Our work determined that, for non-*B. hermsii* species there were multiple *vmp* alleles downstream of the expression site and non-*vmp* genes. Additionally, there were long stretches of sequence downstream of the expression site are nearly identical to other plasmids storing the archived *vmp*. These findings were noted previously in *B. turicatae* (48, 50, 51), and our findings further highlight the potentially exceptional nature of antigenic variation system in *B. hermsii*.

Surprisingly, we found that *B. venezuelensis* RMA01 had two F28 family plasmids. These plasmids (lp35 and lp37) were identical except for the sequence from the *vmp* located in the expression site to the end of 3’ telomere. Indeed, Pennington et al. reported in separate *B. turicatae* clones similar findings of extensive sequence similarity downstream of the *vmp* in the expression site between the expression plasmid and plasmids harboring the archived *vmp* gene (51). *B. venezuelensis* RMA01’s lp35 and lp37 plasmids may exist in different populations within the polyclonal population sequenced and requires further investigation via sequencing of clonally derived isolates. Future efforts comparing the evolution of the antigenic variation systems across all the RF spirochetes could identify conserved themes in the biology of these pathogens.

The *B. hermsii* F28 plasmids contain the expression site of the *vtp* gene rather than the *vmp* expression site (45); however our attempt to identify a related *vtp* gene in the other isolates was largely unsuccessful. Our efforts to determine other putative *vtp* genes were based on the signal peptide of the protein, which is the only way to discriminate a Vsp from a Vtp (53). In agreement with previous reports (15), our findings indicated that *B. anserina* BA2’s Vsp’s signal peptide was highly similar to the *B. hermsii* Vtp signal peptide (90%). These data coupled with *B. anserina*’s clinical presentation in fowl and the Vsp’s expression in the host suggests that *B. anserina* BA2’s single *vmp* gene may have a more nuanced function compared to conventional Vsp and Vtp proteins (80–84).

Our analysis is currently limited by the number of isolates available and the functional annotation of many borrelial proteins. While the more heavily studied species (*B. hermsii*, *B. turicatae*, *B. parkeri*) have multiple isolates available, others have only one isolate (*B. puertoricensis* SUM and *B. venezuelensis* RMA01) or there are limited numbers of isolates (*B. anserina* and *B. coriaceae*). As more isolates are analyzed, a refined understanding of intraspecies plasmid conservation will be accomplished. Perhaps one of the biggest limitations is that functional annotation of borrelial proteins is poor. A considerable number of genes were annotated as “hypothetical proteins” or “domain of unknown function containing proteins”, which complicates comparative genomics analysis. Moreover, we built on prior work with *B. turicatae* and its promoter to identify the putative expression locus for the *vmps* in the remaining species evaluated. We did not check every *vmp* allele in this dataset (∼1,000 predicted *vmp* alleles) for a complete promoter and ribosome binding site, there could be novel expression sites from what we investigated. Transcriptional studies are needed to determine if the F28 plasmid houses the expression locus for antigenic variation in species other than *B. turicatae*. These shortcomings represent opportunities for future work to understand the diversity and genome biology of RF spirochetes and how in relation to human health.

Given the unique complexity of borrelial genomes and our limited understanding of the role of plasmids in RF spirochete biology, assessment of plasmid-resolved genomes is important. This current study sets the foundation to evaluate the role of conserved plasmids in pathogenesis and vector competence. Given the advances in TBRF spirochete genetics (17, 85–87), studies can now be performed to assess the role of these plasmids and the genes they contain in the life cycle of RF spirochetes.

## METHODS

### Bioinformatics

Example commands used with the respective software are found in **Supplemental File 3.**

### Sequence Analysis

#### Genomes

The genomes used in this work were generated by our lab previously (1). The GenBank accession for these genomes were as follows: *B. anserina* BA2 (GCA_023035575.1), *B. hermsii* DAH (GCA_023035675.1), *B. hermsii* YOR (GCA_023035795.1), *B. puertoricensis* SUM (GCA_023035875.1), *B. parkeri* SLO (GCA_023035815.1*), B. venezuelensis* RMA01 (GCA_023035835.1), *B. turicatae* 91E135 (GCA_023035855.1), *B. turicatae* BTE5EL (GCA_003568645.1).

#### Dot Plot Analysis

Self-dot plots of the F6 plasmid family were generated using the nucleotide sequences of each megaplasmid with the LAST aligner (88). The dot plots were annotated in Inkscape (89).

Dot plots of the F27 contigs and read for *B. hermsii* DAH were performed using FlexiDot (v1.06) (90) on the initial assembly and read data generated by Kneubehl et al. (1). The dot plots were annotated in Inkscape.

#### Mauve

ProgressiveMauve (v20150226 build 10) was used to visualize sequence similarities for the megaplasmids(91). ProgressiveMauve was run without assuming collinearity with default options. Visualization was done using the Mauve program and annotated in Inkscape.

#### bptA Phylogenetic Analysis

Using the InterProScan analysis from Kneubehl et al.(1) we identified in each isolate and in the *B. burgdorferi* B31 genome the hits for BptA using the Pfam designation PF17044 (InterProScan results for loci are located in **Supplemental File 2**). Nucleotide sequences were aligned using MAFFT (v7.486) using the --auto option (92). Using these alignments we inferred a maximum likelihood tree using IQ-TREE2 with the -m MFP and -B 1000 options (93–95). The tree was visualized using iTOL (v6) and annotated in Inkscape (96).

#### EasyFig

To visualize whole plasmid similarity for the F6, F20, F27, and F28 plasmid families we used EasyFig (v2.2.5) (97). Each plasmid’s GBK file was uploaded to EasyFig which was run using a minimum cutoff length of 250nt for the BLASTn results (98). Synteny was shown going from grey to darker grey, inverted synteny were shown going from lighter green to darker green, and individual gene features that were highlighted were done by appending the specific gene’s information in the GBK files of each plasmid to include “/colour=“ followed by an RGB code indicating a specific color.

#### Promoter Analysis

Vmp expression site promoters were investigated based on previous reports in *B. hermsii* (46) and *B. turicatae* (50). The identified promoters were aligned using MAFFT and visualized using pyBoxshade.py (99). Promoter features were based on previous reports in B. hermsii (40, 45–47) and B. turicatae (48, 50, 51). These were annotated on the alignment image with Inkscape.

#### vmp Analysis

*Vmp* alleles were assessed using the results of each genome’s InterProScan analysis from Kneubehl et al.(1) and PGAP’s annot.gff file (*vsp*=PF01441, *vlp*=PF00921) per Kuleshov et al. (29) (InterProScan results for loci are indicated in **Supplemental File 4**). This analysis was performed using our WHsTBRF spirochete dataset (1). The phylogenetic analysis for the total *vsp* and *vlp* loci tree inferences as well as individual isolate *vlp* loci trees were performed similar to the *bptA*. This same analysis was carried out for the individual isolate’s *vlp* complements as well. The trees were visualized using iTOL (v6) and annotated in Inkscape (96). The total *vsp* and *vlp* alleles for each genome were graphed using Graphpad Prism 8. The *vlp* subfamilies were determined using the previously typed *vlp*s in *B. hermsii* (*52*). We identified similar alleles in our *B. hermsii* DAH genome using BLASTn and these were used to type the other isolates’ *vlp* alleles (98). Vtp proteins were investigated by using CD-HIT (v4.8.1) clustering of the signal peptide (first 20 amino acids) of each Vsp protein and OspC of *B. burgdorferi* B31 using a sequence identity threshold of 80% (100).

#### Borrelia venezuelensis RMA01 F28 Plasmid 3’ Telomere Long-Read Mapping

To map long-reads specific to the F28 plasmids in *B. venezuelensis* RMA01 we filtered the reads originally used to generate the *B. venezuelensis* RMA01 genome (SRR15006050) using NanoFilt (v2.8.0) with the -q 10 and -l 15000 (101). Minimap2 (v2.24-r1122) was used to map the filtered reads to the *B. venezuelensis* RMA01 genome assembly (GCA_023035835.1) (102, 103). Samtools (v1.11) was used to extract primary reads mapping for lp35 and lp37 using the -b, -F 0x104, and -q 30 options (104). The resulting BAM file for each plasmid was visualized in the Integrated Genomics Viewer (IGV, v2.6.2) and annotated in Inkscape. In the IGV visualizations, indels <10bp were masked, supplementary reads were linked, and reads were sorted by read order.

### Adaptive Sampling Sequencing and Analysis

To confirm the presence of multiple 3’ ends in the F28 plasmids in *B. venezuelensis* RMA01, we performed adaptive sampling using the same genomic DNA that was sequenced to generate the RMA01 genome assembly. We used *B. turicatae* 91E135 as a negative control since the F28 plasmid’s 3’ end should only be present in a single configuration.

To increase the amount of available DNA ends for ligation for the ONT sequencing adapters and fragment the DNA for better adaptive sampling performance, we performed a restriction digest on the genomic DNA. A single BstXI site was present on the F28 plasmids within the PF50 plasmid partition gene, which is ∼3kb upstream of the *vmp* expression site. The genomic DNA was digested with BstXI (New England Biolabs) for one hour at 37°C with enzyme inactivation at 65°C per manufacturer’s protocol. The genomic DNA was cleaned up with a 1:1 (v/v) NEBNext Sample Purification Beads with two 80% ethanol washes. The digested DNA was eluted into 10uL of water and quantified using a Qubit 4 with the Qubit 1x dsDNA broad range assay kit.

The digested DNA was sequenced on ONT’s MinION Mk1B platform. 250ng of digested DNA was end-prepped and dA-tailed with the NEBNext Companion Module for ONT Ligation Sequencing. The prepared DNA was barcoded and adapter’s were ligated from the SQK-NBD114.24 kit following manufacturer’s instructions. 60ng of prepared DNA was loaded onto an R10.4.1 flow cell and sequenced using MinKNOW 22.10.7 with an RTX 3060 Ti enabled. Adaptive sampling was enabled to enrich for reads mapping to *B. venezuelensis* RMA01’s lp35 (CP073229.1) and lp37 (CP073230.1) and *B. turicatae* 91E135’s lp40 (CP073188.1) using a FASTA file containing the sequences for these plasmids. The FAST5 data were basecalled with Guppy (v6.4.2) using the super accurate model, a Q-score filter of 10, --detect_mid_strand_adapter, and -- calib_detect options. The basecalled data were demultiplexed using the same version of Guppy.

Basecalled data were filtered and mapped to their respective genome assemblies to visualize read support for F28 plasmid 3’ ends. The data were filtered using NanoFilt (v2.8.0) -q 10 and -l 15000. The filtered data were mapped to their respective genome assemblies (either *B. venezuelensis* RMA01 GCA_023035835.1 or *B. turicatae* 91E135 GCA_023035855.1) using minimap2. Primary reads were extracted using Samtools (v1.11) with the -b, -F 0x104, and -q 30 options. The resulting BAM file for each plasmid was visualized IGV and annotated in Inkscape. In the IGV visualizations, indels <10bp were masked, supplementary reads were linked, and reads were sorted by read order.

## Data Availability

Sequencing data generated by this study have been deposited to NCBI’s Sequence Read Archive (SRA) and are available through the BioProject PRJNA918510. The adaptive sampling FASTQ file for *B. turicatae* 91E135’s lp40 plasmid was accessioned as SRR22993278. The adaptive sampling FASTQ file for *B. venezuelensis* RMA01’s lp35 and lp37 plasmids was accessioned as SRR22993258.

## Acknowledgements and Funding Statement

We would like to thank Sebastián Muñoz Leal and Marcelo B. Labruna for providing us the *B. venezuelensis* RMA01 isolate used for adaptive sequencing. This work was funded by NIH grants AI137412 (JEL) and AI123652 (JEL). The funders had no role in study design, data collection and interpretation, or the decision to submit the work for publication. ARK conceptualized the idea, performed the formal analysis and investigation, and wrote and edited the manuscript. JEL conceptualized the idea, acquired the funding and supervision, and wrote and edited the manuscript.

## Notes

### Competing Interest Statement

The authors have declared no competing interest.

## References

1. Kneubehl AR, Krishnavajhala A, Leal SM, Replogle AJ, Kingry LC, Bermúdez SE, Labruna MB, Lopez JE. 2022. Comparative genomics of the Western Hemisphere soft tick-borne relapsing fever borreliae highlights extensive plasmid diversity. BMC Genom 23:1–22.

2. Becker NS, Rollins RE, Nosenko K, Paulus A, Martin S, Krebs S, Takano A, Sato K, Kovalev SY, Kawabata H, Fingerle V, Margos G. 2020. High conservation combined with high plasticity: genomics and evolution of *Borrelia bavariensis*. BMC Genom 21:702.

3. Casjens SR, Gilcrease EB, Vujadinovic M, Mongodin EF, Luft BJ, Schutzer SE, Fraser CM, Qiu WG. 2017. Plasmid diversity and phylogenetic consistency in the Lyme disease agent *Borrelia burgdorferi*. BMC Genom 18:165.

4. Casjens SR, Mongodin EF, Qiu WG, Luft BJ, Schutzer SE, Gilcrease EB, Huang WM, Vujadinovic M, Aron JK, Vargas LC, Freeman S, Radune D, Weidman JF, Dimitrov GI, Khouri HM, Sosa JE, Halpin RA, Dunn JJ, Fraser CM. 2012. Genome stability of Lyme disease spirochetes: comparative genomics of *Borrelia burgdorferi* plasmids. PLoS One 7:e33280.

5. Casjens SR, Di L, Akther S, Mongodin EF, Luft BJ, Schutzer SE, Fraser CM, Qiu WG. 2018. Primordial origin and diversification of plasmids in Lyme disease agent bacteria. BMC Genom 19:218.

6. Byram R, Stewart PE, Rosa P. 2004. The essential nature of the ubiquitous 26- kilobase circular replicon of *Borrelia burgdorferi*. J Bacteriol 186:3561–3569.

7. Phelan JP, Kern A, Ramsey ME, Lundt ME, Sharma B, Lin T, Gao L, Norris SJ, Hyde JA, Skare JT, Hu LT. 2019. Genome-wide screen identifies novel genes required for *Borrelia burgdorferi* survival in its *Ixodes* tick vector. PLoS Pathog 15:e1007644.

8. Labandeira-Rey M, Seshu J, Skare JT. 2003. The absence of linear plasmid 25 or 28-1 of *Borrelia burgdorferi* dramatically alters the kinetics of experimental infection via distinct mechanisms. Infect Immun 71:4608–4613.

9. Labandeira-Rey M, Skare JT. 2001. Decreased infectivity in *Borrelia burgdorferi* strain B31 is associated with loss of linear plasmid 25 or 28-1. Infect Immun 69:446–455.

10. Purser JE, Lawrenz MB, Caimano MJ, Howell JK, Radolf JD, Norris SJ. 2003. A plasmid-encoded nicotinamidase (PncA) is essential for infectivity of *Borrelia burgdorferi* in a mammalian host. Mol Microbiol 48:753–764.

11. Purser JE, Norris SJ. 2000. Correlation between plasmid content and infectivity in *Borrelia burgdorferi*. Proc Natl Acad Sci USA 97:13865–13870.

12. Simpson WJ, Garon CF, Schwan TG. 1990. Analysis of supercoiled circular plasmids in infectious and non-infectious *Borrelia burgdorferi*. Microb Pathogen 8:109–118.

13. Xu Y, Kodner C, Coleman L, Johnson RC. 1996. Correlation of plasmids with infectivity of *Borrelia burgdorferi* sensu stricto type strain B31. Infect Immun 64:3870–3876.

14. Revel AT, Blevins JS, Almazán C, Neil L, Kocan KM, de la Fuente J, Hagman KE, Norgard MV. 2005. *bptA* (bbe16) is essential for the persistence of the Lyme disease spirochete, Borrelia burgdorferi, in its natural tick vector. Proc Natl Acad Sci U S A 102:6972–7.

15. Schwan TG, Raffel SJ, Battisti JM. 2020. Transgenic functional complementation with a transmission -associated protein restores spirochete infectivity by tick bite. Ticks Tick Borne Dis 11:101377.

16. Jewett MW, Lawrence K, Bestor AC, Tilly K, Grimm D, Shaw P, VanRaden M, Gherardini F, Rosa PA. 2007. The critical role of the linear plasmid lp36 in the infectious cycle of Borrelia burgdorferi. Mol Microbiol 64:1358–74.

17. Raffel SJ, Battisti JM, Fischer RJ, Schwan TG. 2014. Inactivation of genes for antigenic variation in the relapsing fever spirochete *Borrelia hermsii* reduces infectivity in mice and transmission by ticks. PLoS Pathog 10:e1004056.

18. Grimm D, Tilly K, Bueschel DM, Fisher MA, Policastro PF, Gherardini FC, Schwan TG, Rosa PA. 2005. Defining plasmids required by *Borrelia burgdorferi f*or colonization of tick vector *Ixodes scapularis* (Acari: Ixodidae). J Med Entomol 42:676–684.

19. Stewart PE, Rosa PA. 2008. Transposon mutagenesis of the lyme disease agent *Borrelia burgdorferi*. Methods Mol Biol 431:85–95.

20. Grimm D, Eggers CH, Caimano MJ, Tilly K, Stewart PE, Elias AF, Radolf JD, Rosa PA. 2004. Experimental assessment of the roles of linear plasmids lp25 and lp28-1 of *Borrelia burgdorferi* throughout the infectious cycle. Infect Immun 72:5938–46.

21. Casjens S, Palmer N, van Vugt R, Huang WM, Stevenson B, Rosa P, Lathigra R, Sutton G, Peterson J, Dodson RJ, Haft D, Hickey E, Gwinn M, White O, Fraser CM. 2000. A bacterial genome in flux: the twelve linear and nine circular extrachromosomal DNAs in an infectious isolate of the Lyme disease spirochete *Borrelia burgdorferi*. Mol Microbiol 35:490–516.

22. Stewart PE, Byram R, Grimm D, Tilly K, Rosa PA. 2005. The plasmids of *Borrelia burgdorferi*: essential genetic elements of a pathogen. Plasmid 53:1–13.

23. Miller SC, Porcella SF, Raffel SJ, Schwan TG, Barbour AG. 2013. Large linear plasmids of *Borrelia* species that cause relapsing fever. J Bacteriol 195:3629–3639.

24. Lopez JE, Schrumpf ME, Nagarajan V, Raffel SJ, McCoy BN, Schwan TG. 2010. A novel surface antigen of relapsing fever spirochetes can discriminate between relapsing fever and Lyme borreliosis. Clin Vaccine Immunol 17:564–571.

25. Lopez JE, Wilder HK, Boyle W, Drumheller LB, Thornton JA, Willeford B, Morgan TW, Varela-Stokes A. 2013. Sequence analysis and serological responses against *Borrelia turicatae* BipA, a putative species-specific antigen. PLoS Negl Trop Dis 7:e2454.

26. Curtis MW, Krishnavajhala A, Kneubehl AR, Embers ME, Gettings JR, Yabsley MJ, Lopez JE. 2022. Characterization of immunological responses to Borrelia Immunogenic Protein A (BipA), a species-specific antigen for North American tick-borne relapsing fever. Microbiol Spectr:e01722–21.

27. Dowdell AS, Murphy MD, Azodi C, Swanson SK, Florens L, Chen S, Zückert WR. 2017. Comprehensive spatial analysis of the *Borrelia burgdorferi* lipoproteome reveals a compartmentalization bias toward the bacterial surface. J Bacteriol 199.

28. Johnson M, Zaretskaya I, Raytselis Y, Merezhuk Y, McGinnis S, Madden TL. 2008. NCBI BLAST: a better web interface. Nucleic Acids Res 36:W5–9.

29. Kuleshov KV, Margos G, Fingerle V, Koetsveld J, Goptar IA, Markelov ML, Kolyasnikova NM, Sarksyan DS, Kirdyashkina NP, Shipulin GA, Hovius JW, Platonov AE. 2020. Whole genome sequencing of *Borrelia miyamotoi* isolate Izh- 4: reference for a complex bacterial genome. BMC Genom 21:16.

30. Barbour AG. 2016. Chromosome and plasmids of the tick-borne relapsing fever agent *Borrelia hermsii*. Genome Announc 4.

31. Kingry LC, Replogle A, Batra D, Rowe LA, Sexton C, Dolan M, Connally N, Petersen JM, Schriefer ME. 2017. Toward a complete North American *Borrelia miyamotoi* genome. Genome Announc 5.

32. Chaconas G, Kobryn K. 2010. Structure, function, and evolution of linear replicons in *Borrelia*. Annu Rev Microbiol 64:185–202.

33. Barbour AG, Garon CF. 1987. Linear plasmids of the bacterium *Borrelia burgdorferi* have covalently closed ends. Science 237:409–411.

34. Spealman P, Burrell J, Gresham D. 2020. Inverted duplicate DNA sequences increase translocation rates through sequencing nanopores resulting in reduced base calling accuracy. Nucleic Acids Res 48:4940–4945.

35. Barbour A. 1989. Antigenic variation in relapsing fever *Borrelia* species: genetic aspects, p 783–789. *In* Berg DE, Howe MM (ed), Mobile DNA. American Society for Microbiology, Washington, DC.

36. Zhang JR, Hardham JM, Barbour AG, Norris SJ. 1997. Antigenic variation in Lyme disease borreliae by promiscuous recombination of VMP-like sequence cassettes. Cell 89:275–85.

37. Barbour AG. 2003. Antigenic variation in Borrelia: relapsing fever and Lyme borreliosis, p 319–356. *In* Craig A, Scherf A (ed), Antigenic Variation. Academic Press, London.

38. Barbour AG, Tessier SL, Stoenner HG. 1982. Variable major proteins of *Borrelia hermsii*. J Exp Med 156:1312–1324.

39. Barbour AG, Burman N, Carter CJ, Kitten T, Bergström S. 1991. Variable antigen genes of the relapsing fever agent *Borrelia hermsii* are activated by promoter addition. Mol Microbiol 5:489–493.

40. Dai Q, Restrepo BI, Porcella SF, Raffel SJ, Schwan TG, Barbour AG. 2006. Antigenic variation by *Borrelia hermsii* occurs through recombination between extragenic repetitive elements on linear plasmids. Mol Microbiol 60:1329–1343.

41. Rich SM, Sawyer SA, Barbour AG. 2001. Antigen polymorphism in *Borrelia hermsii*, a clonal pathogenic bacterium. Proc Nat Acad Sci USA 98:15038–15043.

42. Carter CJ, Bergström S, Norris SJ, Barbour AG. 1994. A family of surface-exposed proteins of 20 kilodaltons in the genus *Borrelia*. Infect Immun 62:2792–2799.

43. Margolis N, Hogan D, Cieplak W, Jr., Schwan TG, Rosa PA. 1994. Homology between *Borrelia burgdorferi* OspC and members of the family of *Borrelia hermsii* variable major proteins. Gene 143:105–10.

44. Norris SJ. 2006. Antigenic variation with a twist--the *Borrelia* story. Mol Microbiol 60:1319–22.

45. Barbour AG, Carter CJ, Sohaskey CD. 2000. Surface protein variation by expression site switching in the relapsing fever agent *Borrelia hermsii*. Infect Immun 68:7114–7121.

46. Barbour AG, Carter CJ, Burman N, Freitag CS, Garon CF, Bergström S. 1991. Tandem insertion sequence-like elements define the expression site for variable antigen genes of *Borrelia hermsii*. Infect Immun 59:390–397.

47. Sohaskey CD, Zückert WR, Barbour AG. 1999. The extended promoters for two outer membrane lipoprotein genes of *Borrelia* spp. uniquely include a T-rich region. Mol Microbiol 33:41–51.

48. Pennington PM, Cadavid D, Barbour AG. 1999. Characterization of VspB of *Borrelia turicatae*, a major outer membrane protein expressed in blood and tissues of mice. Infect Immun 67:4637–4645.

49. Kitten T, Barrera AV, Barbour AG. 1993. Intragenic recombination and a chimeric outer membrane protein in the relapsing fever agent *Borrelia hermsii*. J Bacteriol 175:2516–2522.

50. Cadavid D, Pennington PM, Kerentseva TA, Bergström S, Barbour AG. 1997. Immunologic and genetic analyses of VmpA of a neurotropic strain of *Borrelia turicatae*. Infect Immun 65:3352–3360.

51. Pennington PM, Cadavid D, Bunikis J, Norris SJ, Barbour AG. 1999. Extensive interplasmidic duplications change the virulence phenotype of the relapsing fever agent *Borrelia turicatae*. Mol Microbiol 34:1120–1132.

52. Hinnebusch BJ, Barbour AG, Restrepo BI, Schwan TG. 1998. Population structure of the relapsing fever spirochete *Borrelia hermsii* as indicated by polymorphism of two multigene families that encode immunogenic outer surface lipoproteins. Infect Immun 66:432–440.

53. Porcella SF, Raffel SJ, Anderson Jr. DE, Gilk SD, Bono JL, Schrumpf ME, Schwan TG. 2005. Variable tick protein in two genomic groups of the relapsing fever spirochete *Borrelia hermsii* in western North America. Infect Immun 73:6647–6658.

54. Schwan TG, Hinnebusch BJ. 1998. Bloodstream- versus tick-associated variants of a relapsing fever bacterium. Science 280:1938–40.

55. Barbour AG, Burman N, Carter CJ, Kitten T, Bergström S. 1991. Variable antigen genes of the relapsing fever agent Borrelia hermsii are activated by promoter addition. Mol Microbiol 5:489–93.

56. Marcsisin RA, Lewis ER, Barbour AG. 2016. Expression of the tick-associated Vtp protein of *Borrelia hermsii* in a murine model of relapsing fever. PLoS One 11:e0149889.

57. Wilder HK, Raffel SJ, Barbour AG, Porcella SF, Sturdevant DE, Vaisvil B, Kapatral V, Schmitt DP, Schwan TG, Lopez JE. 2016. Transcriptional profiling the 150 kb linear megaplasmid of *Borrelia turicatae* suggests a role in vector colonization and initiating mammalian infection. PLoS One 11:e0147707.

58. Neelakanta G, Sultana H, Sonenshine DE, Marconi RT. 2017. An in vitro blood-feeding method revealed differential *Borrelia turicatae* (Spirochaetales: Spirochaetaceae) gene expression after spirochete acquisition and colonization in the soft tick *Ornithodoros turicata* (Acari: Argasidae). J Med Entomol 54:441–449.

59. Yang XF, Pal U, Alani SM, Fikrig E, Norgard MV. 2004. Essential role for OspA/B in the life cycle of the Lyme disease spirochete. J Exp Med 199:641–8.

60. Fischer JR, Parveen N, Magoun L, Leong JM. 2003. Decorin-binding proteins A and B confer distinct mammalian cell type-specific attachment by *Borrelia burgdorferi*, the Lyme disease spirochete. Proc Natl Acad Sci USA 100:7307–7312.

61. Shi Y, Xu Q, McShan K, Liang FT. 2008. Both decorin-binding proteins A and B are critical for the overall virulence of *Borrelia burgdorferi*. Infect Immun 76:1239–1246.

62. Brooks CS, Vuppala SR, Jett AM, Alitalo A, Meri S, Akins DR. 2005. Complement regulator-acquiring surface protein 1 imparts resistance to human serum in *Borrelia burgdorferi*. J Immunol 175:3299–308.

63. Kraiczy P, Hellwage J, Skerka C, Becker H, Kirschfink M, Simon MM, Brade V, Zipfel PF, Wallich R. 2004. Complement resistance of *Borrelia burgdorferi* correlates with the expression of BbCRASP-1, a novel linear plasmid-encoded surface protein that interacts with human factor H and FHL-1 and is unrelated to Erp proteins. J Biol Chem 279:2421–9.

64. Kraiczy P, Stevenson B. 2013. Complement regulator-acquiring surface proteins of *Borrelia burgdorferi:* Structure, function and regulation of gene expression. Ticks Tick Borne Dis 4:26–34.

65. Hammerschmidt C, Hallstrom T, Skerka C, Wallich R, Stevenson B, Zipfel PF, Kraiczy P. 2012. Contribution of the infection-associated complement regulator- acquiring surface protein 4 (ErpC) to complement resistance of *Borrelia burgdorferi*. Clin Dev Immunol 2012:349657.

66. Bykowski T, Woodman ME, Cooley AE, Brissette CA, Wallich R, Brade V, Kraiczy P, Stevenson B. 2008. *Borrelia burgdorferi* complement regulator- acquiring surface proteins (BbCRASPs): Expression patterns during the mammal-tick infection cycle. Int J Med Microbiol 298 Suppl 1:249–56.

67. Rossmann E, Kraiczy P, Herzberger P, Skerka C, Kirschfink M, Simon MM, Zipfel PF, Wallich R. 2007. Dual binding specificity of a *Borrelia hermsii*- associated complement regulator-acquiring surface protein for factor H and plasminogen discloses a putative virulence factor of relapsing fever spirochetes. J Immunol 178:7292–7301.

68. Balashov YS. 1972. Bloodsucking ticks (Ixodoidea)- vectors of diseases of man and animals. Misc Publ Entomol Soc Am 8:161–376.

69. Cooley RA, Kohls GM. 1944. The Agarasidae of North America, Central America, and Cuba, Monography No. 1 ed. The University Press, Notre Dame, Indiana.

70. Johnson TL, Fischer RJ, Raffel SJ, Schwan TG. 2016. Host associations and genomic diversity of *Borrelia hermsii* in an endemic focus of tick-borne relapsing fever in western North America. Parasites Vectors 9:575.

71. Nieto NC, Teglas MB. 2014. Relapsing fever group *Borrelia* in Southern California rodents. J Med Entomol 51:1029–1034.

72. Nieto NC, Teglas MB, Stewart KM, Wasley T, Wolff PL. 2012. Detection of relapsing fever spirochetes (*Borrelia hermsii* and *Borrelia coriaceae*) in free- ranging mule deer (*Odocoileus hemionus*) from Nevada, United States. Vector Borne Zoonotic Dis 12:99–105.

73. Donaldson TG, Perez de Leon AA, Li AI, Castro-Arellano I, Wozniak E, Boyle WK, Hargrove R, Wilder HK, Kim HJ, Teel PD, Lopez JE. 2016. Assessment of the geographic distribution of *Ornithodoros turicata* (Argasidae): climate variation and host diversity. PLoS Negl Trop Dis 10:e0004383.

74. Kondrashov FA. 2012. Gene duplication as a mechanism of genomic adaptation to a changing environment. Proc Royal Soc B 279:5048–5057.

75. Bratlie MS, Johansen J, Sherman BT, Lempicki RA, Drabløs F. 2010. Gene duplications in prokaryotes can be associated with environmental adaptation. BMC Genom 11:1–17.

76. Jain S, Sutchu S, Rosa PA, Byram R, Jewett MW. 2012. *Borrelia burgdorferi* harbors a transport system essential for purine salvage and mammalian infection. Infect Immun 80:3086–93.

77. Jewett MW, Lawrence KA, Bestor A, Byram R, Gherardini F, Rosa PA. 2009. GuaA and GuaB are essential for *Borrelia burgdorferi* survival in the tick-mouse infection cycle. J Bacteriol 191:6231–41.

78. Tilly K, Krum JG, Bestor A, Jewett MW, Grimm D, Bueschel D, Byram R, Dorward D, Vanraden MJ, Stewart P, Rosa P. 2006. *Borrelia burgdorferi* OspC protein required exclusively in a crucial early stage of mammalian infection. Infect Immun 74:3554–3564.

79. Stewart PE, Wang X, Bueschel DM, Clifton DR, Grimm D, Tilly K, Carroll JA, Weis JJ, Rosa PA. 2006. Delineating the requirement for the *Borrelia burgdorferi* virulence factor OspC in the mammalian host. Infect Immun 74:3547–53.

80. Sambri V, Marangoni A, Olmo A, Storni E, Montagnani M, Fabbi M, Cevenini R. 1999. Specific antibodies reactive with the 22-kilodalton major outer surface protein of *Borrelia anserina* Ni-NL protect chicks from infection. Infect Immun 67:2633–7.

81. DaMassa AJ, Adler HE. 1979. Avian spirochetosis: natural transmission by *Argas* (*Persicargas*) *sanchezi* (Ixodoidea: Argasidae) and existence of different serologic and immunologic types of *Borrelia anserina* in the United States. Am J Vet Res 40:154–7.

82. Levine JF, Dykstra MJ, Nicholson WL, Walker RL, Massey G, Barnes HJ. 1990. Attenuation of *Borrelia anserina* by serial passage in liquid medium. Res Vet Sci 48:64–69.

83. Lisbôa RS, Teixeira RC, Rangel CP, Santos HA, Massard CL, Fonseca AH. 2009. Avian spirochetosis in chickens following experimental transmission of *Borrelia anserina* by *Argas* (*Persicargas*) *miniatus*. Avian Dis 53:166–8.

84. McNeil E, Hinshaw WR, Kissling RE. 1949. A study of *Borrelia anserina* infection (spirochetosis) in turkeys. J Bacteriol 57:191–206.

85. Battisti JM, Raffel SJ, Schwan TG. 2008. A system for site-specific genetic manipulation of the relapsing fever spirochete *Borrelia hermsii*. Methods Mol Biol 431:69–84.

86. Krishnavajhala A, Wilder HK, Boyle WK, Damania A, Thornton JA, Perez de Leon AA, Teel PD, Lopez JE. 2017. Imaging of *Borrelia turicatae* producing the green fluorescent protein reveals persistent colonization of the *Ornithodoros turicata* midgut and salivary glands from nymphal acquisition through transmission. Appl Environ Microbiol 83.

87. Lopez JE, Wilder HK, Hargrove R, Brooks CP, Peterson KE, Beare PA, Sturdevant DE, Nagarajan V, Raffel SJ, Schwan TG. 2013. Development of genetic system to inactivate a *Borrelia turicatae* surface protein selectively produced within the salivary glands of the arthropod vector. PLoS Negl Trop Dis 7:e2514.

88. Frith MC, Kawaguchi R. 2015. Split-alignment of genomes finds orthologies more accurately. Genome Biol 16:1–17.

89. Anonymous. 2022. Inkscape. https://inkscape.org. Accessed

90. Seibt KM, Schmidt T, Heitkam T. 2018. FlexiDot: highly customizable, ambiguity-aware dotplots for visual sequence analyses. Bioinformatics 34:3575–3577.

91. Darling AE, Mau B, Perna NT. 2010. progressiveMauve: multiple genome alignment with gene gain, loss and rearrangement. PLoS One 5:e11147.

92. Katoh K, Standley DM. 2013. MAFFT multiple sequence alignment software version 7: improvements in performance and usability. Mol Biol Evol 30:772–80.

93. Minh BQ, Schmidt HA, Chernomor O, Schrempf D, Woodhams MD, von Haeseler A, Lanfear R. 2020. IQ-TREE 2: New models and efficient methods for phylogenetic inference in the genomic era. Mol Biol Evol 37:1530–1534.

94. Hoang DT, Chernomor O, von Haeseler A, Minh BQ, Vinh LS. 2018. UFBoot2: Improving the ultrafast bootstrap approximation. Mol Biol Evol 35:518–522.

95. Kalyaanamoorthy S, Minh BQ, Wong TKF, von Haeseler A, Jermiin LS. 2017. ModelFinder: fast model selection for accurate phylogenetic estimates. Nat Methods 14:587–589.

96. Letunic I, Bork P. 2021. Interactive Tree Of Life (iTOL) v5: an online tool for phylogenetic tree display and annotation. Nucleic Acids Res 49:W293–W296.

97. Sullivan MJ, Petty NK, Beatson SA. 2011. Easyfig: a genome comparison visualizer. Bioinformatics 27:1009–10.

98. Camacho C, Coulouris G, Avagyan V, Ma N, Papadopoulos J, Bealer K, Madden TL. 2009. BLAST+: architecture and applications. BMC Bioinform 10:421.

99. Anonymous. pyBoxshade. https://github.com/mdbaron42/pyBoxshade. Accessed November 1.

100. Fu L, Niu B, Zhu Z, Wu S, Li W. 2012. CD-HIT: accelerated for clustering the next-generation sequencing data. Bioinformatics 28:3150–2.

101. De Coster W, D’Hert S, Schultz DT, Cruts M, Van Broeckhoven C. 2018. NanoPack: visualizing and processing long-read sequencing data. Bioinformatics 34:2666–2669.

102. Jain C, Rhie A, Zhang H, Chu C, Walenz BP, Koren S, Phillippy AM. 2020. Weighted minimizer sampling improves long read mapping. Bioinformatics 36:i111–i118.

103. Li H. 2018. Minimap2: pairwise alignment for nucleotide sequences. Bioinformatics 34:3094–3100.

104. Li H, Handsaker B, Wysoker A, Fennell T, Ruan J, Homer N, Marth G, Abecasis G, Durbin R. 2009. The sequence alignment/map format and SAMtools. Bioinformatics 25:2078–9.

