## Supplemental Figures for "Comparative genomics analysis of three conserved plasmid families in the Western Hemisphere soft tick-borne relapsing fever borreliae provides insight into variation in genome structure and antigenic variation systems"

### Figure S1

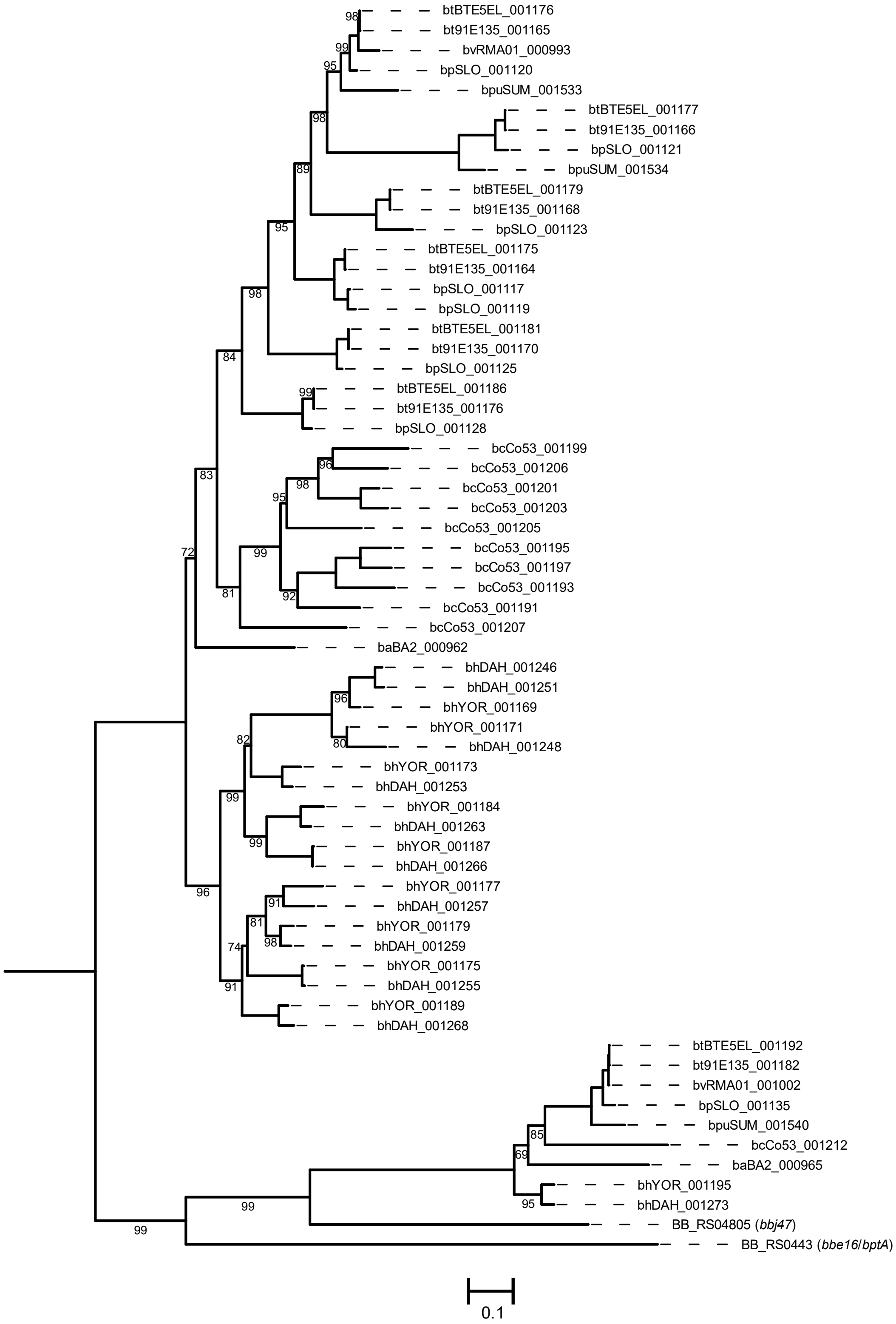

**Figure S1**. Phylogenetic analysis of genes related to *bptA*. A maximum likelihood tree was inferred using the nucleotide sequences of genes similar to *bptA* in *Borreliella* (*Borrelia*) *burgdorferi*. A midpoint rooted tree is shown. . Only branch supports less than 100% are shown. Branch supports indicated are the percentage of 1,000 ultrafast bootstrap replicates. The scale bar represents substitutions per site.

Figure S2

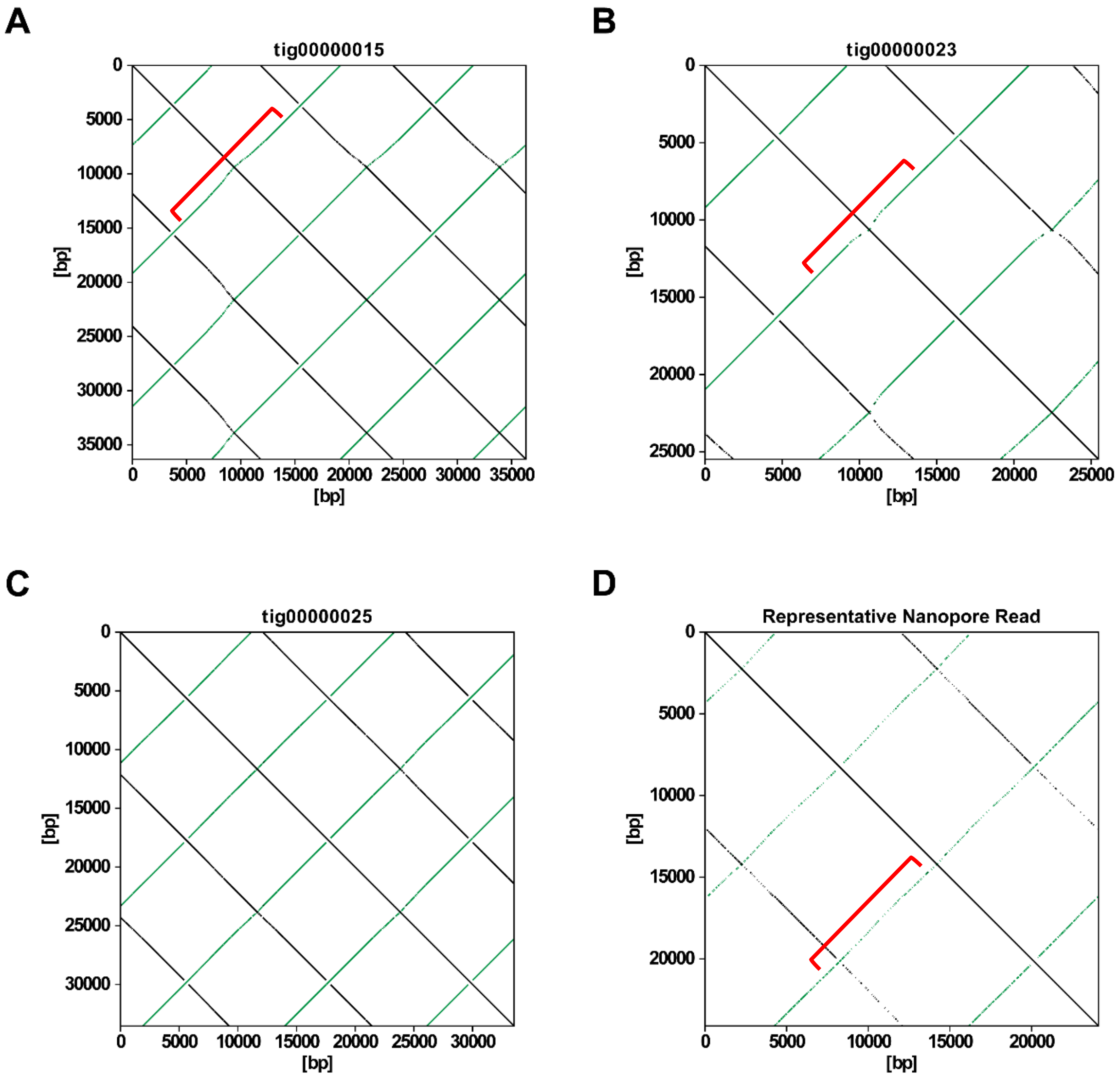


**Figure S2.** F27 plasmid family topology and size analysis. Self-dot plots are shown from the initial *B. hermsii* DAH assembly (**A**-**D**). Three different concatemeric contigs were generated in the initial assembly that depicted complete and incomplete inverted duplications (**A**-**C**). Dot plot analysis of a representative ONT read (~24kb) is shown in **D**. Dot plots were generated using FlexiDot. Black from top left to bottom right indicate direct sequence homology and green from bottom left to top right indicate inverted sequence homology. "Wavey" regions indicating areas of translocation issues through the nanopore due to inverted duplications are indicated by red brackets in the single read in **D** and are magnified in the contigs in **A** and **B**.

### Figure S3

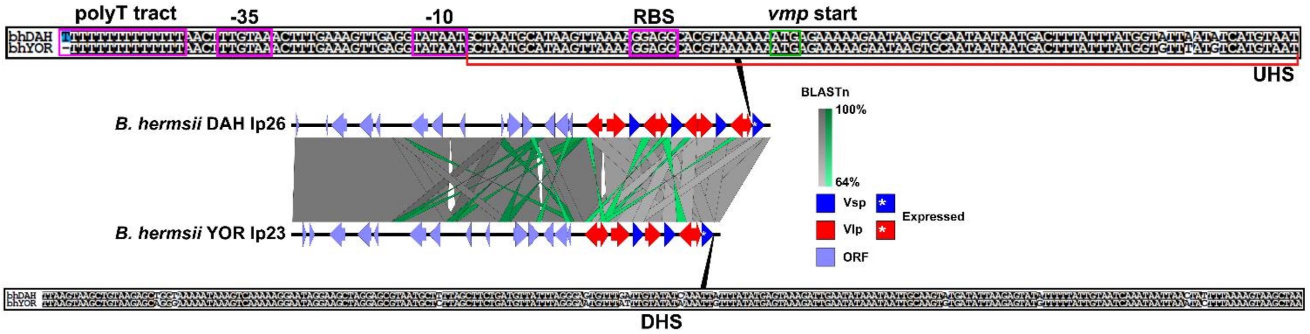


**Figure S3.** *Borrelia hermsii* antigenic variation Vmp expression plasmid alignment and visualization. The F20 plasmids of both *B. hermsii* isolates were aligned using EasyFig. The minimum BLASTn alignment was set to 250nt. BLASTn results scale from 64-100% going from light grey to dark grey whereas inversions scale the same but from light green to dark green. The *vmp* alleles and other open reading frames (ORFs) are shown in specific colors. The *vmp* in the expression site is indicated by an asterisk within the indicated ORF. The vmp promoter alignment is seen in the top sequence. Promoter features are annotated and boxed in purple (RBS=ribosome binding site). The ATG start codon is boxed in green. The upstream homology sequence (UHS) is indicated with a red bracket. The alignment of the downstream homology sequence (DHS) is seen below. For the alignments: a white text in a black square means the sequences are the same, black text in a white square indicates sequences are different, and white texts in blue squares indicates that that nucleotide matches the consensus for that position.

**Figure S4**

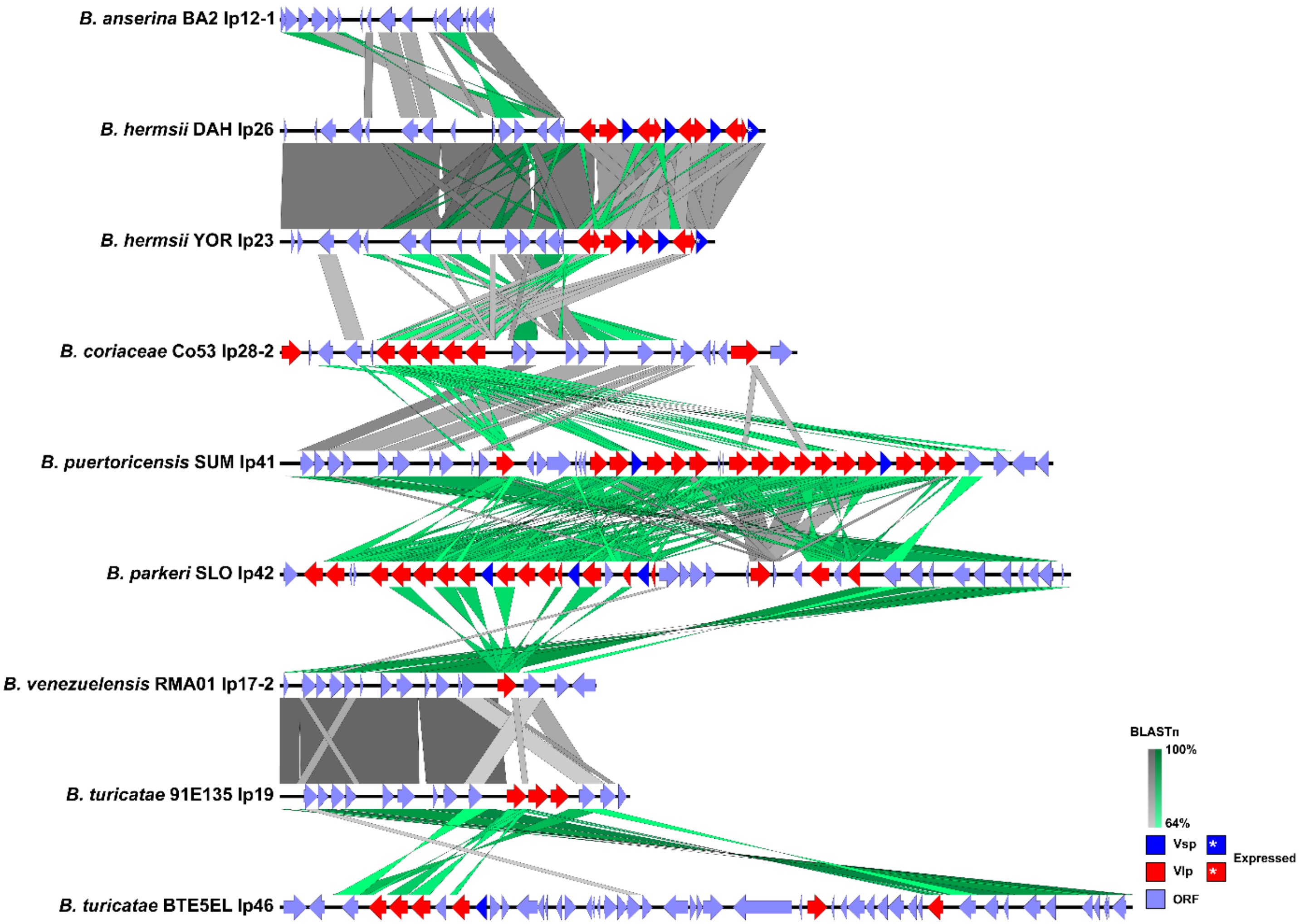


**Figure S4.** Alignment and visualization of the F20 plasmid family. All F20 plasmids were aligned using EasyFig demonstrating the lack of synteny across all species. The minimum BLASTn alignment was set to 250nt. BLASTn results scale from 64-100% going from light grey to dark grey whereas inversions scale the same but from light green to dark green. The *vmp* alleles and other open reading frames (ORFs) are shown in specific colors. The *vmp* in the expression site is indicated by an asterisk within the indicated ORF.

Figure S5

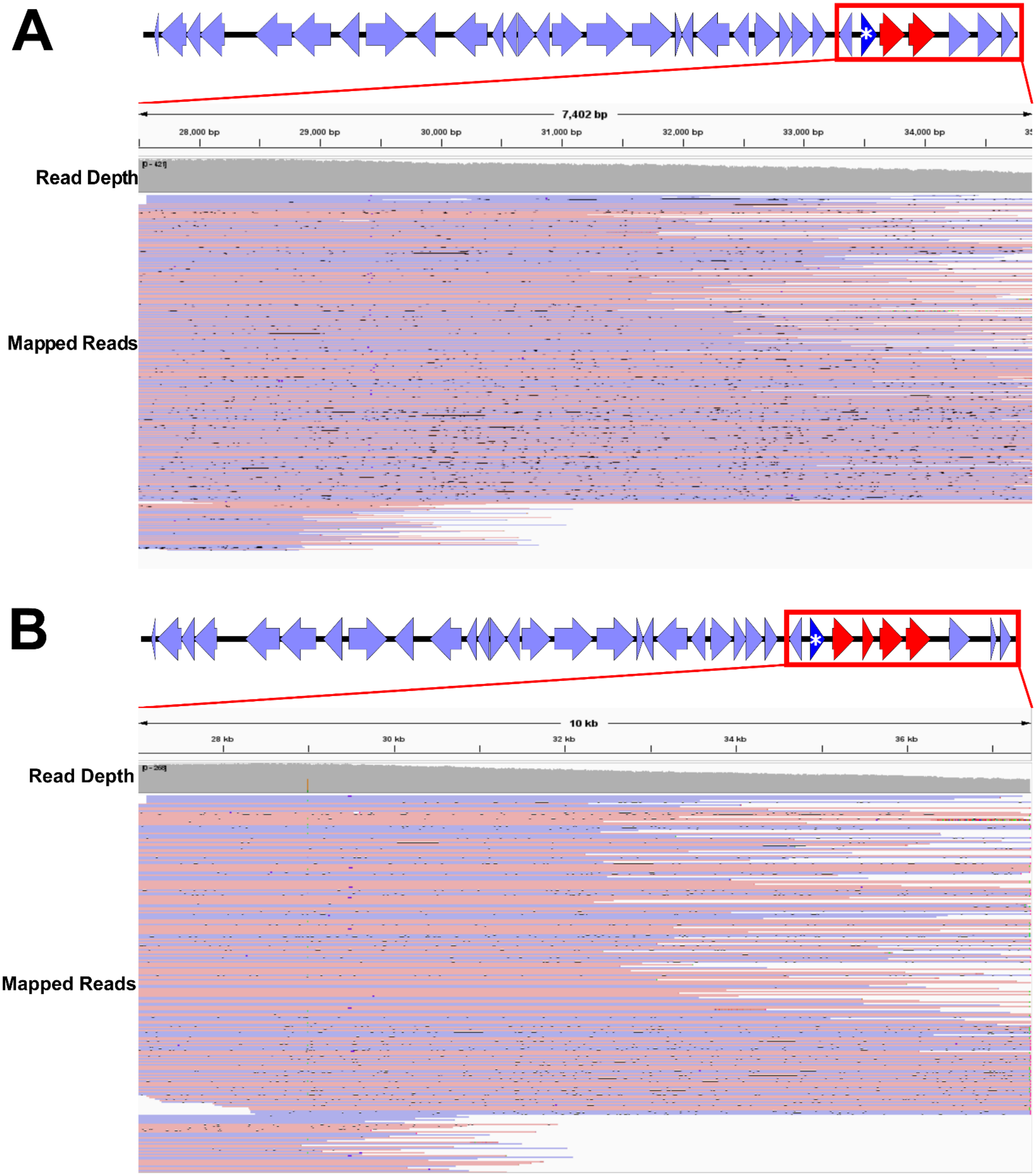
A
7,402 bp
28,000 bp
29,000 bp
30,000 bp
31,000 bp
32,000 bp
33,000 bp
34,000 bp
Read Depth
Mapped Reads
B
10 kb
28 kb
30 kb
32 kb
34 kb
36 kb
Read Depth
Mapped Reads

**Figure S5**. *B. venezuelensis* RMA01 F28 plasmid long-read mapping. Long-read mapping was performed on the lp35 (**A**) and lp37 (**B**) plasmids (F28 plasmids) of *B. venezuelensis* RMA01. Shown are the graphical representations of *B. venezuelensis* RMA01's F28 plasmids and the BAM file visualization in the Integrated Genomics Viewer of the highlighted 3' region starting from the conserved, upstream *oligopeptide permease-like protein* gene. In the graphical representation of the plasmids, the lavender arrows indicate ORFs and their directions, blue triangles indicate *vsp* alleles, and red indicates *vlp* alleles. The blue triangles with the asterisk indicate the *vmp* expression site. Supplementary alignments have been linked with the forward alignment shown in red with the reverse alignment overlayed in blue. Where reads conflict in sequence with the reference, the conflicting base in the read is colored green for adenine, blue for cytosine, red for thymine, and yellow for guanine. Gaps or deletions are shown in the read as a black line. Insertions are indicated with a purple line in the read; however, indels <10bp have been masked.

**Figure S6**

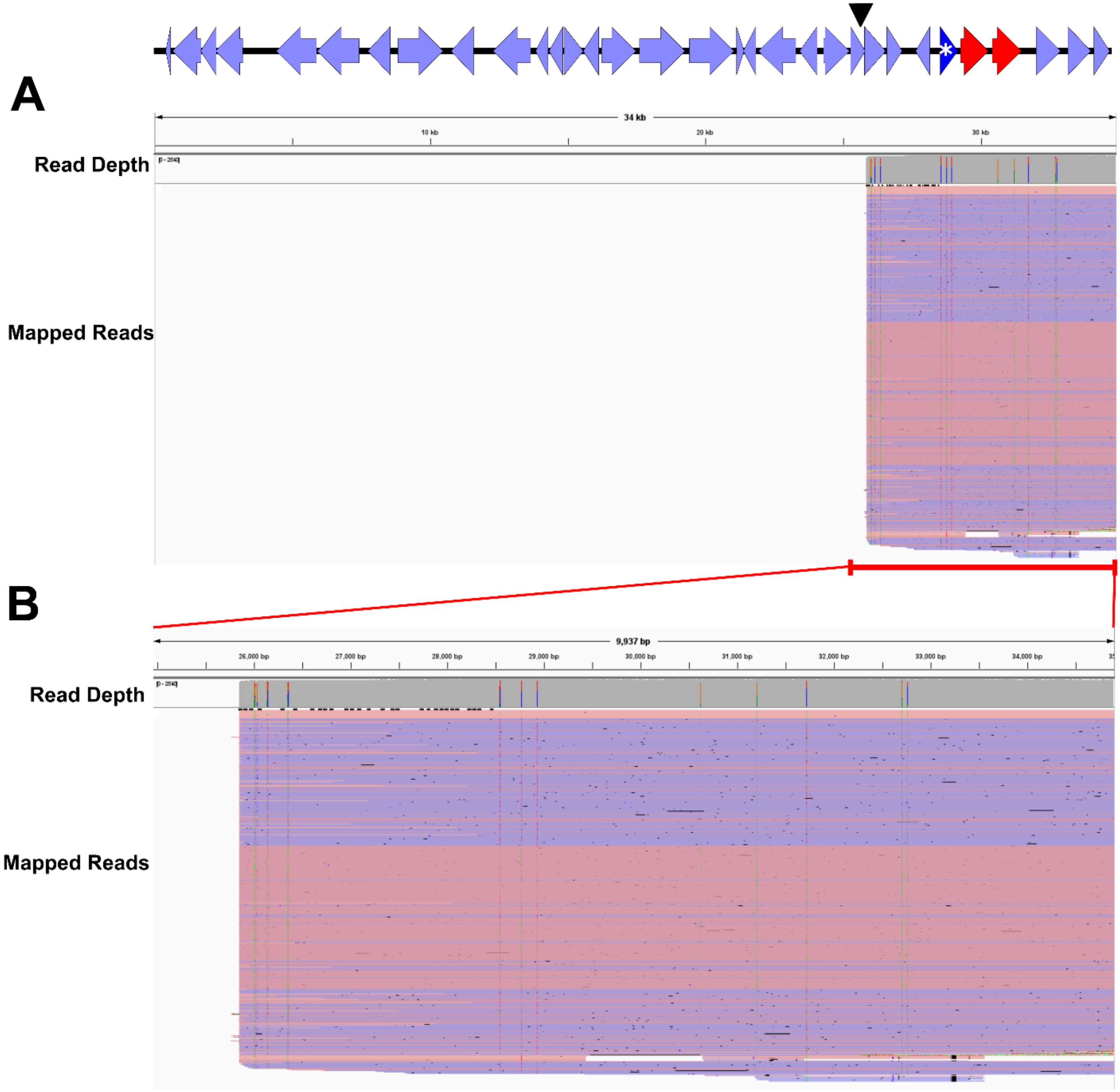

A
34 kb
10 kb
20 kb
30 kb
Read Depth
Mapped Reads
B
9,937 bp
26,000 bp
27,000 bp
28,000 bp
29,000 bp
30,000 bp
31,000 bp
32,000 bp
33,000 bp
34,000 bp
Read Depth
Mapped Reads

**Figure S6.** Mapped adaptive sampling reads for *B. venezuelensis* RMA01 lp35**.** Shown are the graphical representations of *B. venezuelensis* RMA01's lp35 plasmid and the BAM file visualization in the Integrated Genomics Viewer. In the graphical representation of the plasmids, the lavender arrows indicate ORFs and their directions, blue triangles indicate *vsp* alleles, and red indicates *vlp* alleles. The blue triangles with the asterisk indicate the *vmp* expression site. The black triangle represents the location of the BstXI restriction site on this plasmid. Only primary mapped reads are shown. Supplementary alignments have been linked with the forward alignment shown in red with the reverse alignment overlayed in blue. Where reads conflict in sequence with the reference, the conflicting base in the read is colored green for adenine, blue for cytosine, red for thymine, and yellow for guanine. Gaps or deletions are shown in the read as a black line. Insertions are indicated with a purple line in the read; however, indels <10bp have been masked. Reads mapping across the length of the plasmid are shown in **A.** Panel **B** is a enlarged view of the reads mapping to the 3' end.

Figure S7

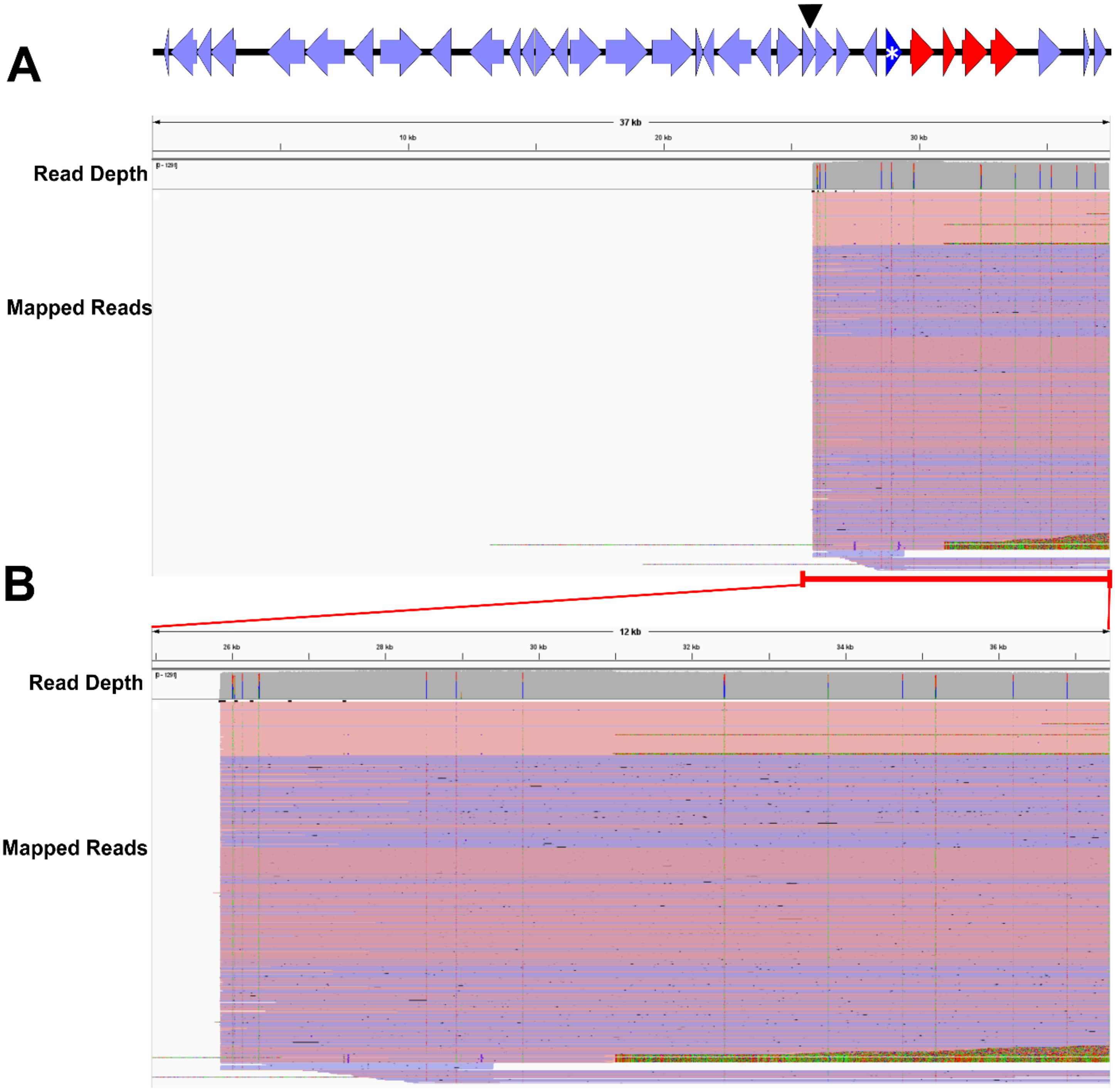
A
37 kb
10 kb
20 kb
30 kb
Read Depth
Mapped Reads
B
12 kb
26 kb
28 kb
30 kb
32 kb
34 kb
36 kb
Read Depth
Mapped Reads

**Figure S7.** Mapped adaptive sampling reads for *B. venezuelensis* RMA01 lp37**.** Shown are the graphical representations of *B. venezuelensis* RMA01's lp37 plasmid and the BAM file visualization in the Integrated Genomics Viewer. In the graphical representation of the plasmids, the lavender arrows indicate ORFs and their directions, blue triangles indicate *vsp* alleles, and red indicates *vlp* alleles. The blue triangles with the asterisk indicate the *vmp* expression site. The black triangle represents the location of the BstXI restriction site on this plasmid. Only primary mapped reads are shown. Supplementary alignments have been linked with the forward alignment shown in red with the reverse alignment overlayed in blue. Where reads conflict in sequence with the reference, the conflicting base in the read is colored green for adenine, blue for cytosine, red for thymine, and yellow for guanine. Gaps or deletions are shown in the read as a black line. Insertions are indicated with a purple line in the read; however, indels <10bp have been masked. Reads mapping across the length of the plasmid are shown in **A.** Panel **B** is a enlarged view of the reads mapping to the 3' end.

**Figure S8**

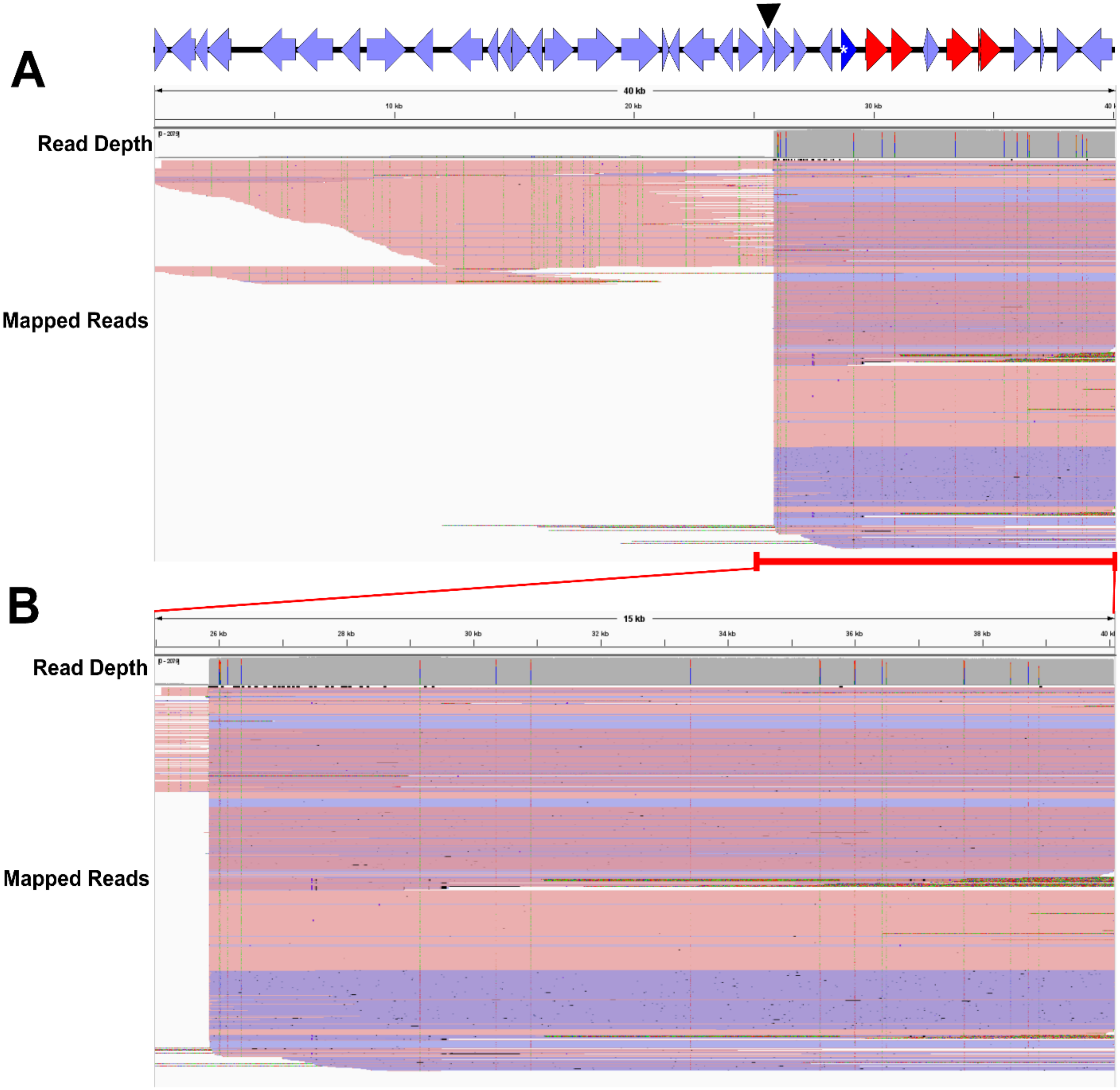
A
40 kb
10 kb
20 kb
30 kb
Read Depth
Mapped Reads
B
15 kb
26 kb
28 kb
30 kb
32 kb
34 kb
36 kb
38 kb
Read Depth
Mapped Reads

**Figure S8.** Mapped adaptive sampling reads for *B. turicatae* 91E135 lp40**.** Shown are the graphical representations of *B. turicatae* 91E135 lp40 plasmid and the BAM file visualization in the Integrated Genomics Viewer. In the graphical representation of the plasmids, the lavender arrows indicate ORFs and their directions, blue triangles indicate *vsp* alleles, and red indicates *vlp* alleles. The blue triangles with the asterisk indicate the *vmp* expression site. The black triangle represents the location of the BstXI restriction site on this plasmid. Only primary mapped reads are shown. Supplementary alignments have been linked with the forward alignment shown in red with the reverse alignment overlayed in blue. Where reads conflict in sequence with the reference, the conflicting base in the read is colored green for adenine, blue for cytosine, red for thymine, and yellow for guanine. Gaps or deletions are shown in the read as a black line. Insertions are indicated with a purple line in the read; however, indels <10bp have been masked. Reads mapping across the length of the plasmid are shown in **A.** Panel **B** is a enlarged view of the reads mapping to the 3' end.

**Figure S9**

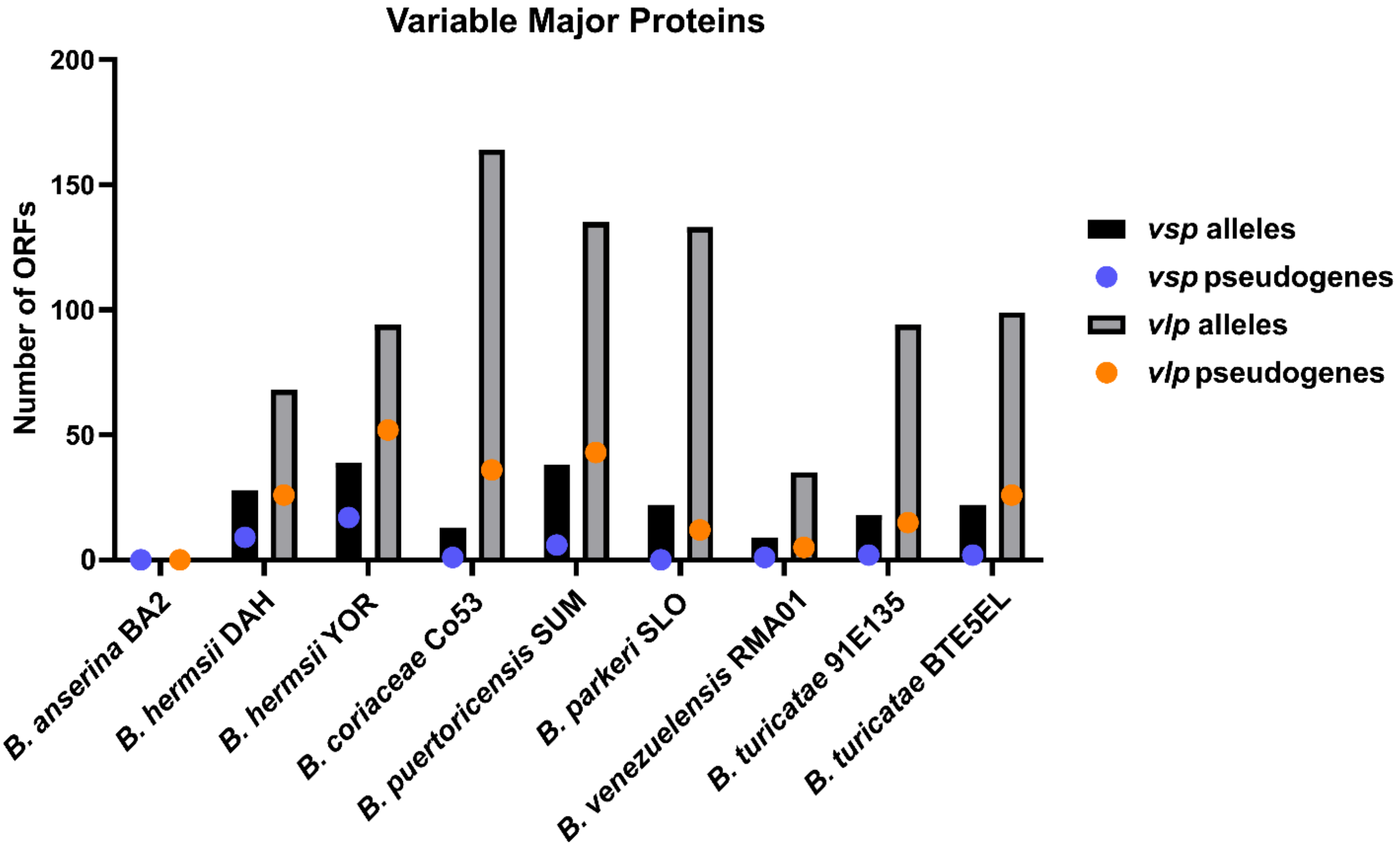


**Figure S9.** *vmp* allele composition. Total numbers of *vmp* alleles (including silent, archived genes as well as predicted pseudogenes) were determined from the InterProScan data for each isolate and are shown as bars. Pseudogenes were determined by PGAP annotation. The number of pseudogenes, as part of the whole *vmp* allele complement, are shown as circles. Note: *B. anserina* BA2 had only one vsp allele and no *vmp* pseudogenes.

**Figure S10**

A

B

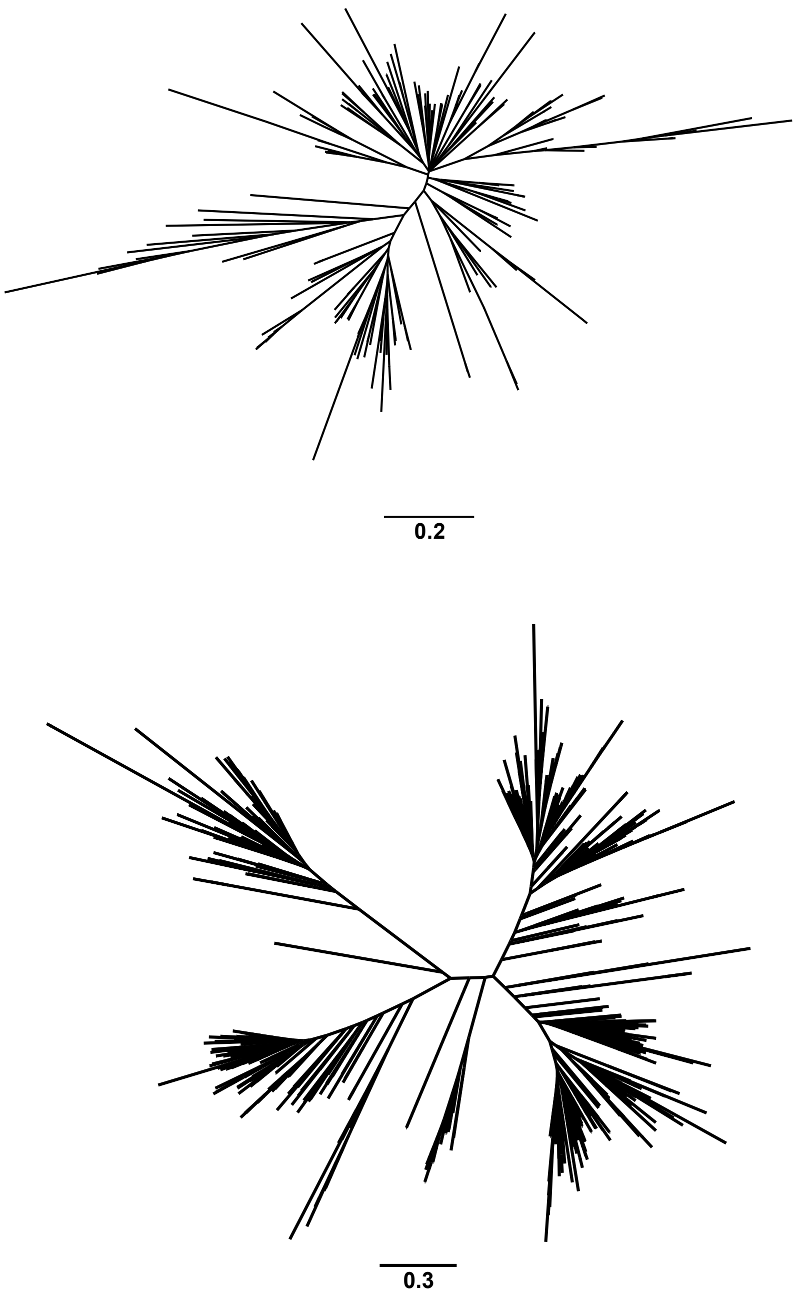


**Figure S10.** Combined phylogenetic analysis of *vmp* alleles. A maximum-likelihood tree was inferred with 1,000 ultrafast bootstrap replicates for all *vsp* (**A**) and *vlp* (**B**) alleles and pseudogenes. Branches with less than 50% support were collapsed. Unrooted trees were visualized in FigTree. The scale bar represents substitutions per site.

**Figure S11**

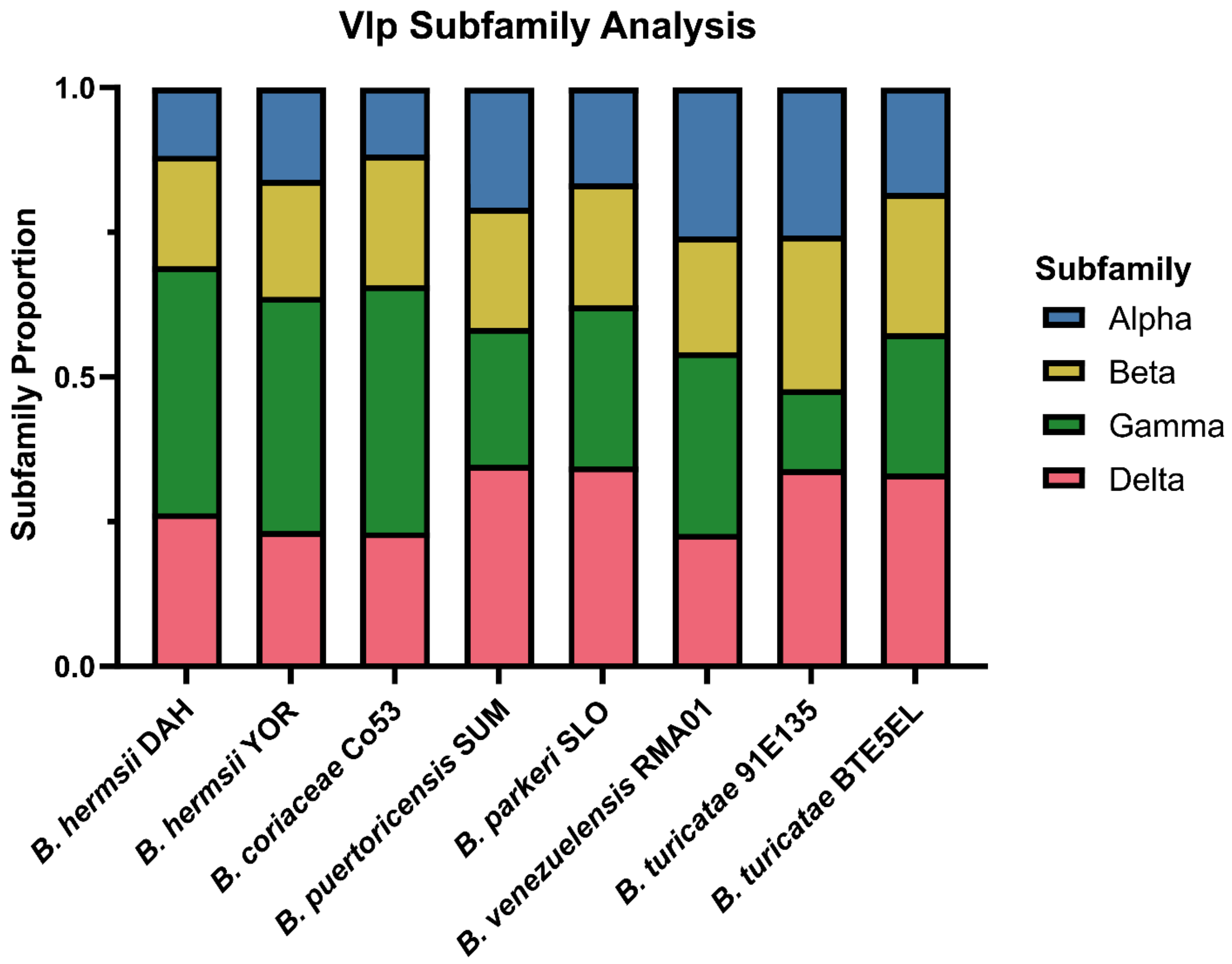


**Figure S11**. *vlp* subfamily proportions. The proportion that each *vlp* subfamily contributed to the whole *vlp* complement of a given genome was determined using the phylogenetic analysis from **Figure 5**.
