## Supplemental File 1 for "Comparative genomics analysis of three conserved plasmid families in the Western Hemisphere soft tick-borne relapsing fever borreliae provides insight into variation in genome structure and antigenic variation systems"

This file contains a sheet for each of the 5 repetitive blocks. The sheets contain the gene loci found in each of the 5 repetitive blocks for all the isolates analyzed.

| Block A | locus | PFAM/SSF Designation by InterProScan | Description |
|---|---|---|---|
| *B. anserina* BA2 | baBA2 000918 | PF05714 | Borrelia Bbcrasp-1 domain containing protein |
| | baBA2 000919 | PF05714 | Borrelia Bbcrasp-1 domain containing protein |
| | baBA2 000920 | PF05714 | Borrelia Bbcrasp-1 domain containing protein |
| | baBA2 000921 | PF05714 | Borrelia Bbcrasp-1 domain containing protein |
| | baBA2 000922 | PF05714 | Borrelia Bbcrasp-1 domain containing protein |
| | baBA2 000923 | PF05714 | Borrelia Bbcrasp-1 domain containing protein |
| *B. hermsii* DAH | bhDAH 001156 | PF05714 | Borrelia Bbcrasp-1 domain containing protein |
| | bhDAH 001157 | PF05714 | Borrelia Bbcrasp-1 domain containing protein |
| | bhDAH 001158 | PF05714 | Borrelia Bbcrasp-1 domain containing protein |
| | bhDAH 001159 | PF05714 | Borrelia Bbcrasp-1 domain containing protein |
| | bhDAH 001160 | PF05714 | Borrelia Bbcrasp-1 domain containing protein |
| | bhDAH 001161 | PF05714 | Borrelia Bbcrasp-1 domain containing protein |
| | bhDAH 001162 | PF05714 | Borrelia Bbcrasp-1 domain containing protein |
| | bhDAH 001163 | PF05714 | Borrelia Bbcrasp-1 domain containing protein |
| | bhDAH 001164 | PF05714 | Borrelia Bbcrasp-1 domain containing protein |
| | bhDAH 001165 | PF05714 | Borrelia Bbcrasp-1 domain containing protein |
| | bhDAH 001166 | PF05714 | Borrelia Bbcrasp-1 domain containing protein |
| | bhDAH 001167 | PF05714 | Borrelia Bbcrasp-1 domain containing protein |
| | bhDAH 001168 | not classified by interproscan | |
| | bhDAH 001169 | PF05714 | Borrelia Bbcrasp-1 domain containing protein |
| | bhDAH 001170 | PF05714 | Borrelia Bbcrasp-1 domain containing protein |
| | bhDAH 001171 | PF05714 | Borrelia Bbcrasp-1 domain containing protein |
| | bhDAH 001172 | PF05714 | Borrelia Bbcrasp-1 domain containing protein |
| | bhDAH 001173 | PF05714 | Borrelia Bbcrasp-1 domain containing protein |
| | bhDAH 001174 | PF05714 | Borrelia Bbcrasp-1 domain containing protein |
| | bhDAH 001175 | PF05714 | Borrelia Bbcrasp-1 domain containing protein |
| | bhDAH 001176 | PF05714 | Borrelia Bbcrasp-1 domain containing protein |
| | bhDAH 001177 | PF05714 | Borrelia Bbcrasp-1 domain containing protein |
| | bhDAH 001178 | PF05714 | Borrelia Bbcrasp-1 domain containing protein |
| | bhDAH 001179 | PF05714 | Borrelia Bbcrasp-1 domain containing protein |
| | bhDAH 001180 | PF05714 | Borrelia Bbcrasp-1 domain containing protein |
| | bhDAH 001181 | PF05714 | Borrelia Bbcrasp-1 domain containing protein |
| *B. hermsii* YOR | bhYOR 001068 | PF05714 | Borrelia Bbcrasp-1 domain containing protein |
| | bhYOR 001069 | PF05714 | Borrelia Bbcrasp-1 domain containing protein |
| | bhYOR 001070 | PF05714 | Borrelia Bbcrasp-1 domain containing protein |
| | bhYOR 001071 | PF05714 | Borrelia Bbcrasp-1 domain containing protein |
| | bhYOR 001072 | PF05714 | Borrelia Bbcrasp-1 domain containing protein |
| | bhYOR 001073 | PF05714 | Borrelia Bbcrasp-1 domain containing protein |
| | bhYOR 001074 | PF05714 | Borrelia Bbcrasp-1 domain containing protein |
| | bhYOR 001075 | PF05714 | Borrelia Bbcrasp-1 domain containing protein |
| | bhYOR 001076 | not classified by interproscan | |
| | bhYOR 001077 | not classified by interproscan | |
| | bhYOR 001078 | PF05714 | Borrelia Bbcrasp-1 domain containing protein |
| | bhYOR 001079 | not classified by interproscan | |
| | bhYOR 001080 | not classified by interproscan | |
| | bhYOR 001081 | PF05714 | Borrelia Bbcrasp-1 domain containing protein |
| | bhYOR 001082 | PF05714 | Borrelia Bbcrasp-1 domain containing protein |
| | bhYOR 001083 | PF05714 | Borrelia Bbcrasp-1 domain containing protein |
| | bhYOR 001084 | PF05714 | Borrelia Bbcrasp-1 domain containing protein |
| | bhYOR 001085 | PF05714 | Borrelia Bbcrasp-1 domain containing protein |
| | bhYOR 001086 | PF05714 | Borrelia Bbcrasp-1 domain containing protein |
| | bhYOR 001087 | SSF51126 | pectin lyase-like fold |
| | bhYOR 001088 | PF05714 | Borrelia Bbcrasp-1 domain containing protein |
| | bhYOR 001089 | not classified by interproscan | |
| | bhYOR 001090 | not classified by interproscan | |
| | bhYOR 001091 | PF05714 | Borrelia Bbcrasp-1 domain containing protein |
| | bhYOR 001092 | not classified by interproscan | |
| | bhYOR 001093 | PF05714 | Borrelia Bbcrasp-1 domain containing protein |
| | bhYOR 001094 | PF05714 | Borrelia Bbcrasp-1 domain containing protein |
| | bhYOR 001095 | PF05714 | Borrelia Bbcrasp-1 domain containing protein |
| | bhYOR 001096 | PF05714 | Borrelia Bbcrasp-1 domain containing protein |
| | bhYOR 001097 | PF05714 | Borrelia Bbcrasp-1 domain containing protein |
| | bhYOR 001098 | PF05714 | Borrelia Bbcrasp-1 domain containing protein |
| | bhYOR 001099 | PF05714 | Borrelia Bbcrasp-1 domain containing protein |
| | bhYOR 001100 | PF05714 | Borrelia Bbcrasp-1 domain containing protein |
| | bhYOR 001101 | PF05714 | Borrelia Bbcrasp-1 domain containing protein |
| | bhYOR 001102 | PF05714 | Borrelia Bbcrasp-1 domain containing protein |
| | bhYOR 001103 | PF05714 | Borrelia Bbcrasp-1 domain containing protein |
| | bhYOR 001104 | PF05714 | Borrelia Bbcrasp-1 domain containing protein |
| | bhYOR 001105 | PF05714 | Borrelia Bbcrasp-1 domain containing protein |
| *B. coriaceae* Co53 | bcCo53 001122 | PF05714 | Borrelia Bbcrasp-1 domain containing protein |
| | bcCo53 001123 | PF05714 | Borrelia Bbcrasp-1 domain containing protein |
| | bcCo53 001124 | PF05714 | Borrelia Bbcrasp-1 domain containing protein |
| | bcCo53 001125 | PF05714 | Borrelia Bbcrasp-1 domain containing protein |
| | bcCo53 001126 | PF05714 | Borrelia Bbcrasp-1 domain containing protein |
| | bcCo53 001127 | PF05714 | Borrelia Bbcrasp-1 domain containing protein |
| | bcCo53 001128 | PF05714 | Borrelia Bbcrasp-1 domain containing protein |
| | bcCo53 001129 | not classified by interproscan | |
| | bcCo53 001130 | PF05714 | Borrelia Bbcrasp-1 domain containing protein |
| | bcCo53 001131 | PF05714 | Borrelia Bbcrasp-1 domain containing protein |
| | bcCo53 001132 | PF05714 | Borrelia Bbcrasp-1 domain containing protein |
| | bcCo53 001133 | PF05714 | Borrelia Bbcrasp-1 domain containing protein |
| | bcCo53 001134 | PF05714 | Borrelia Bbcrasp-1 domain containing protein |
| | bcCo53 001135 | PF05714 | Borrelia Bbcrasp-1 domain containing protein |
| | bcCo53 001136 | PF05714 | Borrelia Bbcrasp-1 domain containing protein |
| *B. puertoricensis* SUM | bpuSUM 001474 | PF05714 | Borrelia Bbcrasp-1 domain containing protein |
| | bpuSUM 001475 | PF05714 | Borrelia Bbcrasp-1 domain containing protein |
| | bpuSUM 001476 | PF05714 | Borrelia Bbcrasp-1 domain containing protein |
| | bpuSUM 001477 | PF05714 | Borrelia Bbcrasp-1 domain containing protein |
| | bpuSUM 001478 | PF05714 | Borrelia Bbcrasp-1 domain containing protein |
| | bpuSUM 001479 | PF05714 | Borrelia Bbcrasp-1 domain containing protein |
| | bpuSUM 001480 | PF05714 | Borrelia Bbcrasp-1 domain containing protein |
| | bpuSUM 001481 | PF05714 | Borrelia Bbcrasp-1 domain containing protein |
| | bpuSUM 001482 | PF05714 | Borrelia Bbcrasp-1 domain containing protein |
| | bpuSUM 001483 | PF05714 | Borrelia Bbcrasp-1 domain containing protein |
| | bpuSUM 001484 | PF05714 | Borrelia Bbcrasp-1 domain containing protein |
| | bpuSUM 001485 | PF05714 | Borrelia Bbcrasp-1 domain containing protein |
| | bpuSUM 001486 | PF05714 | Borrelia Bbcrasp-1 domain containing protein |
| | bpuSUM 001487 | PF05714 | Borrelia Bbcrasp-1 domain containing protein |
| | bpuSUM 001488 | PF05714 | Borrelia Bbcrasp-1 domain containing protein |
| *B. parkeri* SLO | bpSLO 001051 | PF05714 | Borrelia Bbcrasp-1 domain containing protein |
| | bpSLO 001052 | PF05714 | Borrelia Bbcrasp-1 domain containing protein |
| | bpSLO 001053 | PF05714 | Borrelia Bbcrasp-1 domain containing protein |
| | bpSLO 001054 | PF05714 | Borrelia Bbcrasp-1 domain containing protein |
| | bpSLO 001055 | not classified by interproscan | |
| | bpSLO 001056 | PF05714 | Borrelia Bbcrasp-1 domain containing protein |
| | bpSLO 001057 | PF05714 | Borrelia Bbcrasp-1 domain containing protein |
| | bpSLO 001058 | PF05714 | Borrelia Bbcrasp-1 domain containing protein |
| | bpSLO 001059 | PF05714 | Borrelia Bbcrasp-1 domain containing protein |
| | bpSLO 001060 | PF05714 | Borrelia Bbcrasp-1 domain containing protein |
| | bpSLO 001061 | PF05714 | Borrelia Bbcrasp-1 domain containing protein |
| | bpSLO 001062 | PF05714 | Borrelia Bbcrasp-1 domain containing protein |
| | bpSLO 001063 | PF05714 | Borrelia Bbcrasp-1 domain containing protein |
| | bpSLO 001064 | PF05714 | Borrelia Bbcrasp-1 domain containing protein |
| | bpSLO 001065 | PF05714 | Borrelia Bbcrasp-1 domain containing protein |
| *B. venezuelensis* RMA01 | bvRMA01 000926 | PF05714 | Borrelia Bbcrasp-1 domain containing protein |
| | bvRMA01 000927 | PF05714 | Borrelia Bbcrasp-1 domain containing protein |
| | bvRMA01 000928 | PF05714 | Borrelia Bbcrasp-1 domain containing protein |
| | bvRMA01 000929 | PF05714 | Borrelia Bbcrasp-1 domain containing protein |
| | bvRMA01 000930 | not classified by interproscan | |
| | bvRMA01 000931 | PF05714 | Borrelia Bbcrasp-1 domain containing protein |
| | bvRMA01 000932 | PF05714 | Borrelia Bbcrasp-1 domain containing protein |
| | bvRMA01 000933 | PF05714 | Borrelia Bbcrasp-1 domain containing protein |
| | bvRMA01 000934 | PF05714 | Borrelia Bbcrasp-1 domain containing protein |
| | bvRMA01 000935 | PF05714 | Borrelia Bbcrasp-1 domain containing protein |
| | bvRMA01 000936 | PF05714 | Borrelia Bbcrasp-1 domain containing protein |
| | bvRMA01 000937 | PF05714 | Borrelia Bbcrasp-1 domain containing protein |
| *B. turicatae* 91E135 | bt91E135 001096 | PF05714 | Borrelia Bbcrasp-1 domain containing protein |
| | bt91E135 001097 | PF05714 | Borrelia Bbcrasp-1 domain containing protein |
| | bt91E135 001098 | PF05714 | Borrelia Bbcrasp-1 domain containing protein |
| | bt91E135 001099 | PF05714 | Borrelia Bbcrasp-1 domain containing protein |
| | bt91E135 001100 | not classified by interproscan | |
| | bt91E135 001101 | PF05714 | Borrelia Bbcrasp-1 domain containing protein |
| | bt91E135 001102 | PF05714 | Borrelia Bbcrasp-1 domain containing protein |
| | bt91E135 001103 | PF05714 | Borrelia Bbcrasp-1 domain containing protein |
| | bt91E135 001104 | PF05714 | Borrelia Bbcrasp-1 domain containing protein |
| | bt91E135 001105 | PF05714 | Borrelia Bbcrasp-1 domain containing protein |
| | bt91E135 001106 | PF05714 | Borrelia Bbcrasp-1 domain containing protein |
| | bt91E135 001107 | PF05714 | Borrelia Bbcrasp-1 domain containing protein |
| | bt91E135 001108 | PF05714 | Borrelia Bbcrasp-1 domain containing protein |
| | bt91E135 001109 | PF05714 | Borrelia Bbcrasp-1 domain containing protein |
| | bt91E135 001110 | PF05714 | Borrelia Bbcrasp-1 domain containing protein |
| *B. turicatae* BTE5EL | btBTE5EL 001107 | PF05714 | Borrelia Bbcrasp-1 domain containing protein |
| | btBTE5EL 001108 | PF05714 | Borrelia Bbcrasp-1 domain containing protein |
| | btBTE5EL 001109 | PF05714 | Borrelia Bbcrasp-1 domain containing protein |
| | btBTE5EL 001110 | PF05714 | Borrelia Bbcrasp-1 domain containing protein |
| | btBTE5EL 001111 | not classified by interproscan | |
| | btBTE5EL 001112 | PF05714 | Borrelia Bbcrasp-1 domain containing protein |
| | btBTE5EL 001113 | PF05714 | Borrelia Bbcrasp-1 domain containing protein |
| | btBTE5EL 001114 | PF05714 | Borrelia Bbcrasp-1 domain containing protein |
| | btBTE5EL 001115 | PF05714 | Borrelia Bbcrasp-1 domain containing protein |
| | btBTE5EL 001116 | PF05714 | Borrelia Bbcrasp-1 domain containing protein |
| | btBTE5EL 001117 | PF05714 | Borrelia Bbcrasp-1 domain containing protein |
| | btBTE5EL 001118 | PF05714 | Borrelia Bbcrasp-1 domain containing protein |
| | btBTE5EL 001119 | PF05714 | Borrelia Bbcrasp-1 domain containing protein |
| | btBTE5EL 001120 | PF05714 | Borrelia Bbcrasp-1 domain containing protein |
| | btBTE5EL 001121 | PF05714 | Borrelia Bbcrasp-1 domain containing protein |

| Block B | locus | PFAM/SSF Designation by InterProScan | Description |
|---|---|---|---|
| *B. anserina* BA2 | baBA2_000925 | not classified by interproscan | |
| | baBA2_000926 | not classified by interproscan | |
| | baBA2_000927 | not classified by interproscan | |
| | baBA2_000928 | not classified by interproscan | |
| | baBA2_000929 | not classified by interproscan | |
| | baBA2_000930 | not classified by interproscan | |
| | baBA2_000931 | not classified by interproscan | |
| | baBA2_000932 | not classified by interproscan | |
| | baBA2_000933 | not classified by interproscan | |
| | baBA2_000934 | not classified by interproscan | |
| | baBA2_000935 | not classified by interproscan | |
| | baBA2_000936 | not classified by interproscan | |
| | baBA2_000937 | not classified by interproscan | |
| *B. hermsii* DAH | bhDAH_001186 | PF12815 | Spt5 C-terminal nonapeptide repeat binding Spt4 |
| | bhDAH_001187 | not classified by interproscan | |
| | bhDAH_001188 | cd03796 | GT4_PIG-A-like |
| | bhDAH_001189 | not classified by interproscan | |
| | bhDAH_001190 | not classified by interproscan | |
| | bhDAH_001191 | not classified by interproscan | |
| | bhDAH_001192 | not classified by interproscan | |
| | bhDAH_001193 | not classified by interproscan | |
| | bhDAH_001194 | not classified by interproscan | |
| | bhDAH_001195 | not classified by interproscan | |
| | bhDAH_001196 | not classified by interproscan | |
| | bhDAH_001197 | not classified by interproscan | |
| | bhDAH_001198 | not classified by interproscan | |
| | bhDAH_001199 | not classified by interproscan | |
| | bhDAH_001200 | not classified by interproscan | |
| | bhDAH_001201 | not classified by interproscan | |
| | bhDAH_001202 | not classified by interproscan | |
| | bhDAH_001203 | not classified by interproscan | |
| | bhDAH_001204 | not classified by interproscan | |
| | bhDAH_001205 | not classified by interproscan | |
| | bhDAH_001206 | PIN_VapC-like | cd18707 |
| | bhDAH_001207 | not classified by interproscan | |
| | bhDAH_001208 | not classified by interproscan | |
| | bhDAH_001209 | not classified by interproscan | |
| | bhDAH_001210 | not classified by interproscan | |
| | bhDAH_001211 | not classified by interproscan | |
| | bhDAH_001212 | not classified by interproscan | |
| | bhDAH_001213 | not classified by interproscan | |
| | bhDAH_001214 | not classified by interproscan | |
| | bhDAH_001215 | not classified by interproscan | |
| *B. hermsii* YOR | bhYOR_001110 | not classified by interproscan | |
| | bhYOR_001111 | not classified by interproscan | |
| | bhYOR_001112 | not classified by interproscan | |
| | bhYOR_001113 | cd03796 | GT4_PIG-A-like |
| | bhYOR_001114 | not classified by interproscan | |
| | bhYOR_001115 | not classified by interproscan | |
| | bhYOR_001116 | not classified by interproscan | |
| | bhYOR_001117 | not classified by interproscan | |
| | bhYOR_001118 | not classified by interproscan | |
| | bhYOR_001119 | not classified by interproscan | |
| | bhYOR_001120 | not classified by interproscan | |
| | bhYOR_001121 | not classified by interproscan | |
| | bhYOR_001122 | not classified by interproscan | |
| | bhYOR_001123 | not classified by interproscan | |
| | bhYOR_001124 | not classified by interproscan | |
| | bhYOR_001125 | not classified by interproscan | |
| | bhYOR_001126 | not classified by interproscan | |
| | bhYOR_001127 | not classified by interproscan | |
| | bhYOR_001128 | not classified by interproscan | |
| | bhYOR_001129 | not classified by interproscan | |
| | bhYOR_001130 | not classified by interproscan | |
| | bhYOR_001131 | not classified by interproscan | |
| | bhYOR_001132 | not classified by interproscan | |
| | bhYOR_001133 | not classified by interproscan | |
| | bhYOR_001134 | not classified by interproscan | |
| | bhYOR_001135 | not classified by interproscan | |
| | bhYOR_001136 | not classified by interproscan | |
| | bhYOR_001137 | not classified by interproscan | |
| | bhYOR_001138 | not classified by interproscan | |
| | bhYOR_001139 | not classified by interproscan | |
| *B. coriaceae* Co53 | bcCo53_001141 | not classified by interproscan | |
| | bcCo53_001142 | not classified by interproscan | |
| | bcCo53_001143 | not classified by interproscan | |
| | bcCo53_001144 | not classified by interproscan | |
| | bcCo53_001145 | not classified by interproscan | |
| | bcCo53_001146 | not classified by interproscan | |
| | bcCo53_001147 | not classified by interproscan | |
| | bcCo53_001148 | not classified by interproscan | |
| | bcCo53_001149 | not classified by interproscan | |
| | bcCo53_001150 | not classified by interproscan | |
| | bcCo53_001151 | not classified by interproscan | |
| | bcCo53_001152 | not classified by interproscan | |
| | bcCo53_001153 | not classified by interproscan | |
| | bcCo53_001154 | not classified by interproscan | |
| | bcCo53_001155 | not classified by interproscan | |
| | bcCo53_001156 | not classified by interproscan | |
| | bcCo53_001157 | not classified by interproscan | |
| | bcCo53_001158 | not classified by interproscan | |
| | bcCo53_001159 | not classified by interproscan | |
| | bcCo53_001160 | not classified by interproscan | |
| | bcCo53_001161 | not classified by interproscan | |
| | bcCo53_001162 | not classified by interproscan | |
| | bcCo53_001163 | not classified by interproscan | |
| *B. puertoricensis* SUM | bpuSUM_001492 | not classified by interproscan | |
| | bpuSUM_001493 | not classified by interproscan | |
| | bpuSUM_001494 | PF05714 | Borrelia Bbcrasp-1 domain containing protein |
| | bpuSUM_001495 | PF01391 | Collagen triple helix repeat (20 copies) |
| | bpuSUM_001496 | PF01391 | Collagen triple helix repeat (20 copies) |
| | bpuSUM_001497 | not classified by interproscan | |
| | bpuSUM_001498 | not classified by interproscan | |
| | bpuSUM_001499 | not classified by interproscan | |
| | bpuSUM_001500 | not classified by interproscan | |
| | bpuSUM_001501 | not classified by interproscan | |
| | bpuSUM_001502 | not classified by interproscan | |
| | bpuSUM_001503 | not classified by interproscan | |
| | bpuSUM_001504 | not classified by interproscan | |
| | bpuSUM_001505 | not classified by interproscan | |
| | bpuSUM_001506 | not classified by interproscan | |
| | bpuSUM_001507 | not classified by interproscan | |
| | bpuSUM_001508 | not classified by interproscan | |
| | bpuSUM_001509 | not classified by interproscan | |
| *B. parkeri* SLO | bpSLO_001069 | not classified by interproscan | |
| | bpSLO_001070 | not classified by interproscan | |
| | bpSLO_001071 | not classified by interproscan | |
| | bpSLO_001072 | PF05714 | Borrelia Bbcrasp-1 domain containing protein |
| | bpSLO_001073 | PF01391 | Collagen triple helix repeat (20 copies) |
| | bpSLO_001074 | not classified by interproscan | |
| | bpSLO_001075 | not classified by interproscan | |
| | bpSLO_001076 | not classified by interproscan | |
| | bpSLO_001077 | not classified by interproscan | |
| | bpSLO_001078 | not classified by interproscan | |
| | bpSLO_001079 | not classified by interproscan | |
| | bpSLO_001080 | not classified by interproscan | |
| | bpSLO_001081 | not classified by interproscan | |
| | bpSLO_001082 | not classified by interproscan | |
| | bpSLO_001083 | not classified by interproscan | |
| | bpSLO_001084 | PF05714 | Borrelia Bbcrasp-1 domain containing protein |
| | bpSLO_001085 | not classified by interproscan | |
| | bpSLO_001086 | not classified by interproscan | |
| | bpSLO_001087 | not classified by interproscan | |
| | bpSLO_001088 | PF05714 | Borrelia Bbcrasp-1 domain containing protein |
| | bpSLO_001089 | not classified by interproscan | |
| | bpSLO_001090 | not classified by interproscan | |
| | bpSLO_001091 | not classified by interproscan | |
| | bpSLO_001092 | not classified by interproscan | |
| *B. venezuelensis* RMA01 | bvRMA01_000941 | not classified by interproscan | |
| | bvRMA01_000942 | not classified by interproscan | |
| | bvRMA01_000943 | not classified by interproscan | |
| | bvRMA01_000944 | PF05714 | Borrelia Bbcrasp-1 domain containing protein |
| | bvRMA01_000945 | PF01391 | Collagen triple helix repeat (20 copies) |
| | bvRMA01_000946 | not classified by interproscan | |
| | bvRMA01_000947 | not classified by interproscan | |
| | bvRMA01_000948 | not classified by interproscan | |
| | bvRMA01_000949 | not classified by interproscan | |
| | bvRMA01_000950 | not classified by interproscan | |
| | bvRMA01_000951 | not classified by interproscan | |
| | bvRMA01_000952 | not classified by interproscan | |
| | bvRMA01_000953 | not classified by interproscan | |
| | bvRMA01_000954 | not classified by interproscan | |
| | bvRMA01_000955 | not classified by interproscan | |
| | bvRMA01_000956 | not classified by interproscan | |
| | bvRMA01_000957 | not classified by interproscan | |
| | bvRMA01_000958 | not classified by interproscan | |
| | bvRMA01_000959 | not classified by interproscan | |
| | bvRMA01_000960 | not classified by interproscan | |
| | bvRMA01_000961 | PF05714 | Borrelia Bbcrasp-1 domain containing protein |
| | bvRMA01_000962 | not classified by interproscan | |
| | bvRMA01_000963 | not classified by interproscan | |
| | bvRMA01_000964 | not classified by interproscan | |
| | bvRMA01_000965 | not classified by interproscan | |
| | bvRMA01_000966 | not classified by interproscan | |
| | bvRMA01_000967 | not classified by interproscan | |
| *B. turicatae* 91E135 | bt91E135_001114 | not classified by interproscan | |
| | bt91E135_001115 | not classified by interproscan | |
| | bt91E135_001116 | not classified by interproscan | |
| | bt91E135_001117 | not classified by interproscan | |
| | bt91E135_001118 | PF01391 | Collagen triple helix repeat (20 copies) |
| | bt91E135_001119 | not classified by interproscan | |
| | bt91E135_001120 | not classified by interproscan | |
| | bt91E135_001121 | not classified by interproscan | |
| | bt91E135_001122 | not classified by interproscan | |
| | bt91E135_001123 | not classified by interproscan | |
| | bt91E135_001124 | not classified by interproscan | |
| | bt91E135_001125 | not classified by interproscan | |
| | bt91E135_001126 | not classified by interproscan | |
| | bt91E135_001127 | not classified by interproscan | |
| | bt91E135_001128 | not classified by interproscan | |
| | bt91E135_001129 | not classified by interproscan | |
| | bt91E135_001130 | not classified by interproscan | |
| | bt91E135_001131 | not classified by interproscan | |
| | bt91E135_001132 | not classified by interproscan | |
| | bt91E135_001133 | PF05714 | Borrelia Bbcrasp-1 domain containing protein |
| | bt91E135_001134 | not classified by interproscan | |
| | bt91E135_001135 | not classified by interproscan | |
| | bt91E135_001136 | not classified by interproscan | |
| | bt91E135_001137 | not classified by interproscan | |
| *B. turicatae* BTE5EL | btBTE5EL_001125 | not classified by interproscan | |
| | btBTE5EL_001126 | not classified by interproscan | |
| | btBTE5EL_001127 | not classified by interproscan | |
| | btBTE5EL_001128 | not classified by interproscan | |
| | btBTE5EL_001129 | PF01391 | Collagen triple helix repeat (20 copies) |
| | btBTE5EL_001130 | not classified by interproscan | |
| | btBTE5EL_001131 | not classified by interproscan | |
| | btBTE5EL_001132 | not classified by interproscan | |
| | btBTE5EL_001133 | not classified by interproscan | |
| | btBTE5EL_001134 | not classified by interproscan | |
| | btBTE5EL_001135 | not classified by interproscan | |
| | btBTE5EL_001136 | not classified by interproscan | |
| | btBTE5EL_001137 | not classified by interproscan | |
| | btBTE5EL_001138 | not classified by interproscan | |
| | btBTE5EL_001139 | not classified by interproscan | |
| | btBTE5EL_001140 | not classified by interproscan | |
| | btBTE5EL_001141 | not classified by interproscan | |
| | btBTE5EL_001142 | not classified by interproscan | |
| | btBTE5EL_001143 | not classified by interproscan | |
| | btBTE5EL_001144 | PF05714 | Borrelia Bbcrasp-1 domain containing protein |
| | btBTE5EL_001145 | not classified by interproscan | |
| | btBTE5EL_001146 | not classified by interproscan | |
| | btBTE5EL_001147 | not classified by interproscan | |
| | btBTE5EL_001148 | not classified by interproscan | |

| **Block C** | **locus** | **PFAM/SSF Designation by InterProScan** | **Description** |
|---|---|---|---|
| ***B. anserina* BA2** | baBA2_000961 | not classified by interproscan | |
| | baBA2_000962 | PF17044 | Borrelial persistence in ticks protein A |
| | baBA2_000963 | not classified by interproscan | |
| ***B. hermsii* DAH** | bhDAH_001244 | not classified by interproscan | |
| | bhDAH_001245 | PF17044 | Borrelial persistence in ticks protein A |
| | bhDAH_001246 | PF17044 | Borrelial persistence in ticks protein A |
| | bhDAH_001247 | not classified by interproscan | not classified by interproscan |
| | bhDAH_001248 | PF17044 | Borrelial persistence in ticks protein A |
| | bhDAH_001249 | PF17044 | Borrelial persistence in ticks protein A |
| | bhDAH_001250 | not classified by interproscan | |
| | bhDAH_001251 | PF17044 | Borrelial persistence in ticks protein A |
| | bhDAH_001252 | not classified by interproscan | |
| | bhDAH_001253 | PF17044 | Borrelial persistence in ticks protein A |
| | bhDAH_001254 | not classified by interproscan | |
| | bhDAH_001255 | PF17044 | Borrelial persistence in ticks protein A |
| | bhDAH_001256 | not classified by interproscan | |
| | bhDAH_001257 | PF17044 | Borrelial persistence in ticks protein A |
| | bhDAH_001258 | not classified by interproscan | |
| | bhDAH_001259 | PF17044 | Borrelial persistence in ticks protein A |
| | bhDAH_001260 | not classified by interproscan | |
| | bhDAH_001261 | not classified by interproscan | |
| | bhDAH_001262 | not classified by interproscan | |
| | bhDAH_001263 | PF17044 | Borrelial persistence in ticks protein A |
| | bhDAH_001264 | not classified by interproscan | |
| | bhDAH_001265 | PF02524 | KID repeat |
| | bhDAH_001266 | PF17044 | Borrelial persistence in ticks protein A |
| | bhDAH_001267 | not classified by interproscan | |
| | bhDAH_001268 | PF17044 | Borrelial persistence in ticks protein A |
| | bhDAH_001269 | not classified by interproscan | |
| ***B. hermsii* YOR** | bhYOR_001168 | not classified by interproscan | |
| | bhYOR_001169 | PF17044 | Borrelial persistence in ticks protein A |
| | bhYOR_001170 | not classified by interproscan | |
| | bhYOR_001171 | PF17044 | Borrelial persistence in ticks protein A |
| | bhYOR_001172 | not classified by interproscan | |
| | bhYOR_001173 | PF17044 | Borrelial persistence in ticks protein A |
| | bhYOR_001174 | not classified by interproscan | |
| | bhYOR_001175 | PF17044 | Borrelial persistence in ticks protein A |
| | bhYOR_001176 | not classified by interproscan | |
| | bhYOR_001177 | PF17044 | Borrelial persistence in ticks protein A |
| | bhYOR_001178 | not classified by interproscan | |
| | bhYOR_001179 | PF17044 | Borrelial persistence in ticks protein A |
| | bhYOR_001180 | not classified by interproscan | |
| | bhYOR_001181 | not classified by interproscan | |
| | bhYOR_001182 | not classified by interproscan | |
| | bhYOR_001183 | not classified by interproscan | |
| | bhYOR_001184 | PF17044 | Borrelial persistence in ticks protein A |
| | bhYOR_001185 | not classified by interproscan | |
| | bhYOR_001186 | SSF58113 | Apolipoprotein A-I |
| | bhYOR_001187 | PF17044 | Borrelial persistence in ticks protein A |
| | bhYOR_001188 | not classified by interproscan | |
| | bhYOR_001189 | PF17044 | Borrelial persistence in ticks protein A |
| | bhYOR_001190 | not classified by interproscan | |
| ***B. coriaceae* Co53** | bcCo53_001189 | not classified by interproscan | |
| | bcCo53_001190 | not classified by interproscan | |
| | bcCo53_001191 | PF17044 | Borrelial persistence in ticks protein A |
| | bcCo53_001192 | not classified by interproscan | |
| | bcCo53_001193 | PF17044 | Borrelial persistence in ticks protein A |
| | bcCo53_001194 | not classified by interproscan | |
| | bcCo53_001195 | PF17044 | Borrelial persistence in ticks protein A |
| | bcCo53_001196 | not classified by interproscan | |
| | bcCo53_001197 | PF17044 | Borrelial persistence in ticks protein A |
| | bcCo53_001198 | not classified by interproscan | |
| | bcCo53_001199 | PF17044 | Borrelial persistence in ticks protein A |
| | bcCo53_001200 | not classified by interproscan | |
| | bcCo53_001201 | PF17044 | Borrelial persistence in ticks protein A |
| | bcCo53_001202 | not classified by interproscan | |
| | bcCo53_001203 | PF17044 | Borrelial persistence in ticks protein A |
| | bcCo53_001204 | not classified by interproscan | |
| | bcCo53_001205 | PF17044 | Borrelial persistence in ticks protein A |
| | bcCo53_001206 | PF17044 | Borrelial persistence in ticks protein A |
| | bcCo53_001207 | PF17044 | Borrelial persistence in ticks protein A |
| | bcCo53_001208 | not classified by interproscan | |
| ***B. puertoricensis* SUM** | bpuSUM_001533 | PF17044 | Borrelial persistence in ticks protein A |
| | bpuSUM_001534 | PF17044 | Borrelial persistence in ticks protein A |
| | bpuSUM_001535 | not classified by interproscan | |
| ***B. parkeri* SLO** | bpSLO_001117 | PF17044 | Borrelial persistence in ticks protein A |
| | bpSLO_001118 | not classified by interproscan | |
| | bpSLO_001119 | PF17044 | Borrelial persistence in ticks protein A |
| | bpSLO_001120 | PF17044 | Borrelial persistence in ticks protein A |
| | bpSLO_001121 | PF17044 | Borrelial persistence in ticks protein A |
| | bpSLO_001122 | not classified by interproscan | |
| | bpSLO_001123 | PF17044 | Borrelial persistence in ticks protein A |
| | bpSLO_001124 | PF17044 | Borrelial persistence in ticks protein A |
| | bpSLO_001125 | not classified by interproscan | |
| | bpSLO_001126 | not classified by interproscan | |
| | bpSLO_001127 | not classified by interproscan | |
| | bpSLO_001128 | PF17044 | Borrelial persistence in ticks protein A |
| | bpSLO_001129 | not classified by interproscan | |
| ***B. venezuelensis* RMA01** | bvRMA01_000992 | PF17044 | Borrelial persistence in ticks protein A |
| | bvRMA01_000993 | PF17044 | Borrelial persistence in ticks protein A |
| | bvRMA01_000994 | not classified by interproscan | |
| | bvRMA01_000995 | not classified by interproscan | |
| ***B. turicatae* 91E135** | bt91E135_001162 | not classified by interproscan | |
| | bt91E135_001163 | not classified by interproscan | |
| | bt91E135_001164 | PF17044 | Borrelial persistence in ticks protein A |
| | bt91E135_001165 | PF17044 | Borrelial persistence in ticks protein A |
| | bt91E135_001166 | PF17044 | Borrelial persistence in ticks protein A |
| | bt91E135_001167 | not classified by interproscan | |
| | bt91E135_001168 | PF17044 | Borrelial persistence in ticks protein A |
| | bt91E135_001169 | not classified by interproscan | |
| | bt91E135_001170 | PF17044 | Borrelial persistence in ticks protein A |
| | bt91E135_001171 | not classified by interproscan | |
| | bt91E135_001172 | not classified by interproscan | |
| | bt91E135_001173 | not classified by interproscan | |
| | bt91E135_001174 | not classified by interproscan | |
| | bt91E135_001175 | not classified by interproscan | |
| | bt91E135_001176 | PF17044 | Borrelial persistence in ticks protein A |
| | bt91E135_001177 | not classified by interproscan | |
| ***B. turicatae* BTE5EL** | btBTE5EL_001173 | not classified by interproscan | |
| | btBTE5EL_001174 | not classified by interproscan | |
| | btBTE5EL_001175 | PF17044 | Borrelial persistence in ticks protein A |
| | btBTE5EL_001176 | PF17044 | Borrelial persistence in ticks protein A |
| | btBTE5EL_001177 | PF17044 | Borrelial persistence in ticks protein A |
| | btBTE5EL_001178 | not classified by interproscan | |
| | btBTE5EL_001179 | PF17044 | Borrelial persistence in ticks protein A |
| | btBTE5EL_001180 | not classified by interproscan | |
| | btBTE5EL_001181 | PF17044 | Borrelial persistence in ticks protein A |
| | btBTE5EL_001182 | not classified by interproscan | |
| | btBTE5EL_001183 | not classified by interproscan | |
| | btBTE5EL_001184 | not classified by interproscan | |
| | btBTE5EL_001185 | not classified by interproscan | |
| | btBTE5EL_001186 | PF17044 | Borrelial persistence in ticks protein A |
| | btBTE5EL_001187 | not classified by interproscan | |

| Block D | locus | PFAM/SSF Designation by InterProScan | Description |
|---|---|---|---|
| ***B. anserina*** **BA2** | baBA2_000981 | not classified by interproscan | |
| | baBA2_000982 | not classified by interproscan | |
| | baBA2_000983 | not classified by interproscan | |
| | baBA2_000984 | not classified by interproscan | |
| | baBA2_000985 | not classified by interproscan | |
| | baBA2_000986 | not classified by interproscan | |
| | baBA2_000987 | not classified by interproscan | |
| | baBA2_000988 | not classified by interproscan | |
| | baBA2_000989 | not classified by interproscan | |
| | baBA2_000990 | SSF69349 | Phage fibre proteins |
| ***B. hermsii*** **DAH** | bhDAH_001291 | not classified by interproscan | |
| | bhDAH_001292 | not classified by interproscan | |
| | bhDAH_001293 | not classified by interproscan | |
| | bhDAH_001294 | not classified by interproscan | |
| | bhDAH_001295 | not classified by interproscan | |
| | bhDAH_001296 | not classified by interproscan | |
| | bhDAH_001297 | not classified by interproscan | |
| | bhDAH_001298 | not classified by interproscan | |
| | bhDAH_001299 | not classified by interproscan | |
| | bhDAH_001300 | not classified by interproscan | |
| | bhDAH_001301 | not classified by interproscan | |
| | bhDAH_001302 | not classified by interproscan | |
| | bhDAH_001303 | not classified by interproscan | |
| | bhDAH_001304 | not classified by interproscan | |
| | bhDAH_001305 | not classified by interproscan | |
| | bhDAH_001306 | not classified by interproscan | |
| | bhDAH_001307 | not classified by interproscan | |
| | bhDAH_001308 | not classified by interproscan | |
| | bhDAH_001309 | not classified by interproscan | |
| ***B. hermsii*** **YOR** | bhYOR_001214 | not classified by interproscan | |
| | bhYOR_001215 | not classified by interproscan | |
| | bhYOR_001216 | not classified by interproscan | |
| | bhYOR_001217 | not classified by interproscan | |
| | bhYOR_001218 | not classified by interproscan | |
| | bhYOR_001219 | not classified by interproscan | |
| | bhYOR_001220 | not classified by interproscan | |
| | bhYOR_001221 | not classified by interproscan | |
| | bhYOR_001222 | not classified by interproscan | |
| | bhYOR_001223 | not classified by interproscan | |
| | bhYOR_001224 | not classified by interproscan | |
| | bhYOR_001225 | not classified by interproscan | |
| | bhYOR_001226 | not classified by interproscan | |
| | bhYOR_001227 | not classified by interproscan | |
| | bhYOR_001228 | not classified by interproscan | |
| | bhYOR_001229 | not classified by interproscan | |
| | bhYOR_001230 | not classified by interproscan | |
| | bhYOR_001231 | not classified by interproscan | |
| | bhYOR_001232 | not classified by interproscan | |
| | bcCo53_001231 | not classified by interproscan | |
| ***B. coriaceae*** **Co53** | bcCo53_001232 | not classified by interproscan | |
| | bcCo53_001233 | not classified by interproscan | |
| | bcCo53_001234 | not classified by interproscan | |
| | bcCo53_001235 | not classified by interproscan | |
| | bcCo53_001236 | not classified by interproscan | |
| | bcCo53_001237 | not classified by interproscan | |
| | bcCo53_001238 | not classified by interproscan | |
| | bcCo53_001239 | not classified by interproscan | |
| | bcCo53_001240 | not classified by interproscan | |
| | bcCo53_001241 | not classified by interproscan | |
| | bcCo53_001242 | not classified by interproscan | |
| | bcCo53_001243 | not classified by interproscan | |
| | bcCo53_001244 | not classified by interproscan | |
| | bcCo53_001245 | not classified by interproscan | |
| | bcCo53_001246 | not classified by interproscan | |
| | bcCo53_001247 | not classified by interproscan | |
| | bcCo53_001248 | not classified by interproscan | |
| | bcCo53_001249 | not classified by interproscan | |
| | bcCo53_001250 | not classified by interproscan | |
| | bcCo53_001251 | not classified by interproscan | |
| | bcCo53_001252 | not classified by interproscan | |
| | bcCo53_001253 | not classified by interproscan | |
| | bcCo53_001254 | not classified by interproscan | |
| | bcCo53_001255 | not classified by interproscan | |
| | bcCo53_001256 | not classified by interproscan | |
| | bcCo53_001257 | not classified by interproscan | |
| | bcCo53_001258 | not classified by interproscan | |
| | bcCo53_001259 | not classified by interproscan | |
| | bcCo53_001260 | not classified by interproscan | |
| | bcCo53_001261 | not classified by interproscan | |
| ***B. puertoricensis*** **SUM** | none | | |
| ***B. parkeri*** **SLO** | bpSLO_001155 | not classified by interproscan | |
| | bpSLO_001156 | not classified by interproscan | |
| | bpSLO_001157 | not classified by interproscan | |
| | bpSLO_001158 | not classified by interproscan | |
| | bpSLO_001159 | PF07761 | Protein of unknown function (DUF1617) |
| | bpSLO_001160 | not classified by interproscan | |
| | bpSLO_001161 | not classified by interproscan | |
| | bpSLO_001162 | PF01391 | Collagen triple helix repeat (20 copies) |
| | bpSLO_001163 | not classified by interproscan | |
| | bpSLO_001164 | not classified by interproscan | |
| | bpSLO_001165 | not classified by interproscan | |
| | bpSLO_001166 | not classified by interproscan | |
| | bpSLO_001167 | not classified by interproscan | |
| | bpSLO_001168 | not classified by interproscan | |
| | bpSLO_001169 | not classified by interproscan | |
| | bpSLO_001170 | not classified by interproscan | |
| | bpSLO_001171 | not classified by interproscan | |
| | bpSLO_001172 | not classified by interproscan | |
| | bpSLO_001173 | SSF51161 | Trimeric LpxA-like enzymes |
| | bpSLO_001174 | not classified by interproscan | |
| | bpSLO_001175 | not classified by interproscan | |
| | bpSLO_001176 | not classified by interproscan | |
| | bpSLO_001177 | not classified by interproscan | |
| | bpSLO_001178 | not classified by interproscan | |
| | bpSLO_001179 | not classified by interproscan | |
| | bpSLO_001180 | not classified by interproscan | |
| | bpSLO_001181 | not classified by interproscan | |
| ***B. venezuelensis*** **RMA01** | bvRMA01_001022 | not classified by interproscan | |
| | bvRMA01_001023 | not classified by interproscan | |
| | bvRMA01_001024 | not classified by interproscan | |
| | bvRMA01_001025 | not classified by interproscan | |
| | bvRMA01_001026 | not classified by interproscan | |
| | bvRMA01_001027 | not classified by interproscan | |
| | bvRMA01_001028 | not classified by interproscan | |
| | bvRMA01_001029 | not classified by interproscan | |
| | bvRMA01_001030 | not classified by interproscan | |
| | bvRMA01_001031 | not classified by interproscan | |
| | bvRMA01_001032 | not classified by interproscan | |
| | bvRMA01_001033 | not classified by interproscan | |
| | bvRMA01_001034 | not classified by interproscan | |
| | bvRMA01_001035 | not classified by interproscan | |
| | bvRMA01_001036 | not classified by interproscan | |
| | bvRMA01_001037 | cd00139 | PIPKc |
| | bvRMA01_001038 | not classified by interproscan | |
| | bvRMA01_001039 | not classified by interproscan | |
| | bvRMA01_001040 | not classified by interproscan | |
| ***B. turicatae*** **91E135** | bt91E135_001201 | not classified by interproscan | |
| | bt91E135_001202 | not classified by interproscan | |
| | bt91E135_001203 | not classified by interproscan | |
| | bt91E135_001204 | not classified by interproscan | |
| | bt91E135_001205 | not classified by interproscan | |
| | bt91E135_001206 | PF07761 | Protein of unknown function (DUF1617) |
| | bt91E135_001207 | not classified by interproscan | |
| | bt91E135_001208 | not classified by interproscan | |
| | bt91E135_001209 | not classified by interproscan | |
| | bt91E135_001210 | not classified by interproscan | |
| | bt91E135_001211 | not classified by interproscan | |
| | bt91E135_001212 | not classified by interproscan | |
| | bt91E135_001213 | PF07761 | Protein of unknown function (DUF1617) |
| | bt91E135_001214 | not classified by interproscan | |
| | bt91E135_001215 | not classified by interproscan | |
| | bt91E135_001216 | not classified by interproscan | |
| | bt91E135_001217 | not classified by interproscan | |
| | bt91E135_001218 | not classified by interproscan | |
| | bt91E135_001219 | not classified by interproscan | |
| | bt91E135_001220 | not classified by interproscan | |
| | bt91E135_001221 | not classified by interproscan | |
| | bt91E135_001222 | not classified by interproscan | |
| | bt91E135_001223 | not classified by interproscan | |
| | bt91E135_001224 | not classified by interproscan | |
| | bt91E135_001225 | not classified by interproscan | |
| | bt91E135_001226 | not classified by interproscan | |
| | bt91E135_001227 | not classified by interproscan | |
| | bt91E135_001228 | not classified by interproscan | |
| ***B. turicatae*** **BTE5EL** | btBTE5EL_001211 | not classified by interproscan | |
| | btBTE5EL_001212 | not classified by interproscan | |
| | btBTE5EL_001213 | not classified by interproscan | |
| | btBTE5EL_001214 | not classified by interproscan | |
| | btBTE5EL_001215 | not classified by interproscan | |
| | btBTE5EL_001216 | PF07761 | Protein of unknown function (DUF1617) |
| | btBTE5EL_001217 | not classified by interproscan | |
| | btBTE5EL_001218 | not classified by interproscan | |
| | btBTE5EL_001219 | not classified by interproscan | |
| | btBTE5EL_001220 | not classified by interproscan | |
| | btBTE5EL_001221 | not classified by interproscan | |
| | btBTE5EL_001222 | not classified by interproscan | |
| | btBTE5EL_001223 | PF07761 | Protein of unknown function (DUF1617) |
| | btBTE5EL_001224 | not classified by interproscan | |
| | btBTE5EL_001225 | not classified by interproscan | |
| | btBTE5EL_001226 | not classified by interproscan | |
| | btBTE5EL_001227 | not classified by interproscan | |
| | btBTE5EL_001228 | not classified by interproscan | |
| | btBTE5EL_001229 | not classified by interproscan | |
| | btBTE5EL_001230 | not classified by interproscan | |
| | btBTE5EL_001231 | not classified by interproscan | |
| | btBTE5EL_001232 | not classified by interproscan | |
| | btBTE5EL_001233 | not classified by interproscan | |
| | btBTE5EL_001234 | not classified by interproscan | |
| | btBTE5EL_001235 | not classified by interproscan | |
| | btBTE5EL_001236 | not classified by interproscan | |
| | btBTE5EL_001237 | not classified by interproscan | |
| | btBTE5EL_001238 | not classified by interproscan | |
| | btBTE5EL_001239 | not classified by interproscan | |
| | btBTE5EL_001240 | not classified by interproscan | |

| Block E | locus | PFAM/SSF Designation by InterProScan | Description |
| --- | --- | --- | --- |
| ***B. anserina*** **BA2** | none | | |
| ***B. hermsii*** **DAH** | bhDAH_001311 | not classified by interproscan | |
| ***B. hermsii*** **YOR** | none | | |
| ***B. coriaceae*** **Co53** | none | | |
| ***B. puertoricensis*** **SUM** | none | | |
| ***B. parkeri*** **SLO** | none | | |
| ***B. venezuelensis*** **RMA01** | none | | |
| ***B. turicatae*** **91E135** | none | | |
| ***B. turicatae*** **BTE5EL** | none | | |
