## Supplemental File 2 for "Comparative genomics analysis of three conserved plasmid families in the Western Hemisphere soft tick-borne relapsing fever borreliae provides insight into variation in genome structure and antigenic variation systems"

**Genes Classified as "Borrelial persistence in ticks protein A" by InterProScan**

| ***Borrelia anserina* BA2** | **Gene or Pseudogene** | **Database** | **Start Location** | **Stop Location** | **Score** | **Signature Description** |
|---|---|---|---|---|---|---|
| gene-baBA2_000962_orf21554 | gene | Pfam | 24 | 198 | 8.30E-28 | Target=null 24 198;ID=match$1465_24_198;signature_desc=Borrelial persistence in ticks protein A;Name=PF17044;status=T;Dbxref="InterPro:IPR031471" |
| gene-baBA2_000965_orf21610 | gene | Pfam | 50 | 215 | 1.80E-29 | Target=null 50 215;ID=match$387_50_215;signature_desc=Borrelial persistence in ticks protein A;Name=PF17044;status=T;Dbxref="InterPro:IPR031471" |
| ***Borrelia hermsii* DAH** | | | | | | |
| gene-bhDAH_001245_orf25941 | gene | Pfam | 6 | 66 | 1.10E-11 | Target=null 6 66;ID=match$56_6_66;signature_desc=Borrelial persistence in ticks protein A;Name=PF17044;status=T;Dbxref="InterPro:IPR031471" |
| gene-bhDAH_001246_orf25949 | gene | Pfam | 36 | 106 | 6.30E-08 | Target=null 36 106;ID=match$906_36_106;signature_desc=Borrelial persistence in ticks protein A;Name=PF17044;status=T;Dbxref="InterPro:IPR031471" |
| gene-bhDAH_001248_orf25970 | gene | Pfam | 6 | 66 | 2.60E-12 | Target=null 6 66;ID=match$1225_6_66;signature_desc=Borrelial persistence in ticks protein A;Name=PF17044;status=T;Dbxref="InterPro:IPR031471" |
| gene-bhDAH_001249_orf25977 | gene | Pfam | 36 | 106 | 3.10E-08 | Target=null 36 106;ID=match$60_36_106;signature_desc=Borrelial persistence in ticks protein A;Name=PF17044;status=T;Dbxref="InterPro:IPR031471" |
| gene-bhDAH_001251_orf26003 | gene | Pfam | 35 | 194 | 5.70E-27 | Target=null 35 194;ID=match$757_35_194;signature_desc=Borrelial persistence in ticks protein A;Name=PF17044;status=T;Dbxref="InterPro:IPR031471" |
| gene-bhDAH_001253_orf26034 | gene | Pfam | 32 | 196 | 2.60E-29 | Target=null 32 196;ID=match$1378_32_196;signature_desc=Borrelial persistence in ticks protein A;Name=PF17044;status=T;Dbxref="InterPro:IPR031471" |
| gene-bhDAH_001255_orf26065 | gene | Pfam | 36 | 205 | 1.20E-31 | Target=null 36 205;ID=match$2471_36_205;signature_desc=Borrelial persistence in ticks protein A;Name=PF17044;status=T;Dbxref="InterPro:IPR031471" |
| gene-bhDAH_001257_orf26094 | gene | Pfam | 33 | 199 | 1.50E-30 | Target=null 33 199;ID=match$762_33_199;signature_desc=Borrelial persistence in ticks protein A;Name=PF17044;status=T;Dbxref="InterPro:IPR031471" |
| gene-bhDAH_001259_orf26124 | gene | Pfam | 33 | 199 | 1.60E-31 | Target=null 33 199;ID=match$2174_33_199;signature_desc=Borrelial persistence in ticks protein A;Name=PF17044;status=T;Dbxref="InterPro:IPR031471" |
| gene-bhDAH_001263_orf26173 | gene | Pfam | 32 | 195 | 7.90E-31 | Target=null 32 195;ID=match$78_32_195;signature_desc=Borrelial persistence in ticks protein A;Name=PF17044;status=T;Dbxref="InterPro:IPR031471" |
| gene-bhDAH_001266_orf26205 | gene | Pfam | 29 | 199 | 1.20E-31 | Target=null 29 199;ID=match$452_29_199;signature_desc=Borrelial persistence in ticks protein A;Name=PF17044;status=T;Dbxref="InterPro:IPR031471" |
| gene-bhDAH_001268_orf26230 | gene | Pfam | 29 | 199 | 2.40E-33 | Target=null 29 199;ID=match$67_29_199;signature_desc=Borrelial persistence in ticks protein A;Name=PF17044;status=T;Dbxref="InterPro:IPR031471" |
| gene-bhDAH_001273_orf26340 | gene | Pfam | 37 | 213 | 1.50E-34 | Target=null 37 213;ID=match$2091_37_213;signature_desc=Borrelial persistence in ticks protein A;Name=PF17044;status=T;Dbxref="InterPro:IPR031471" |
| ***Borrelia hermsii* YOR** | | | | | | |
| gene-bhYOR_001169_orf24828 | gene | Pfam | 40 | 197 | 5.00E-25 | Target=null 40 197;ID=match$552_40_197;signature_desc=Borrelial persistence in ticks protein A;Name=PF17044;status=T;Dbxref="InterPro:IPR031471" |
| gene-bhYOR_001171_orf24860 | gene | Pfam | 38 | 197 | 7.80E-26 | Target=null 38 197;ID=match$1041_38_197;signature_desc=Borrelial persistence in ticks protein A;Name=PF17044;status=T;Dbxref="InterPro:IPR031471" |
| gene-bhYOR_001173_orf24894 | gene | Pfam | 34 | 199 | 5.70E-28 | Target=null 34 199;ID=match$114_34_199;signature_desc=Borrelial persistence in ticks protein A;Name=PF17044;status=T;Dbxref="InterPro:IPR031471" |
| gene-bhYOR_001175_orf24926 | gene | Pfam | 36 | 210 | 1.40E-30 | Target=null 36 210;ID=match$3282_36_210;signature_desc=Borrelial persistence in ticks protein A;Name=PF17044;status=T;Dbxref="InterPro:IPR031471" |
| gene-bhYOR_001177_orf24953 | gene | Pfam | 28 | 199 | 3.80E-30 | Target=null 28 199;ID=match$257_28_199;signature_desc=Borrelial persistence in ticks protein A;Name=PF17044;status=T;Dbxref="InterPro:IPR031471" |
| gene-bhYOR_001179_orf24984 | gene | Pfam | 29 | 199 | 1.70E-31 | Target=null 29 199;ID=match$3170_29_199;signature_desc=Borrelial persistence in ticks protein A;Name=PF17044;status=T;Dbxref="InterPro:IPR031471" |
| gene-bhYOR_001184_orf25033 | gene | Pfam | 32 | 194 | 2.40E-29 | Target=null 32 194;ID=match$943_32_194;signature_desc=Borrelial persistence in ticks protein A;Name=PF17044;status=T;Dbxref="InterPro:IPR031471" |
| gene-bhYOR_001187_orf25074 | gene | Pfam | 29 | 199 | 1.20E-31 | Target=null 29 199;ID=match$1559_29_199;signature_desc=Borrelial persistence in ticks protein A;Name=PF17044;status=T;Dbxref="InterPro:IPR031471" |
| gene-bhYOR_001189_orf25100 | gene | Pfam | 32 | 199 | 1.40E-32 | Target=null 32 199;ID=match$788_32_199;signature_desc=Borrelial persistence in ticks protein A;Name=PF17044;status=T;Dbxref="InterPro:IPR031471" |
| gene-bhYOR_001195_orf25241 | gene | Pfam | 37 | 212 | 9.50E-35 | Target=null 37 212;ID=match$1586_37_212;signature_desc=Borrelial persistence in ticks protein A;Name=PF17044;status=T;Dbxref="InterPro:IPR031471" |
| ***Borrelia coriaceae* Co53** | | | | | | |
| gene-bcCo53_001191_orf24686 | gene | Pfam | 34 | 203 | 5.10E-26 | Target=null 34 203;ID=match$1867_34_203;signature_desc=Borrelial persistence in ticks protein A;Name=PF17044;status=T;Dbxref="InterPro:IPR031471" |
| gene-bcCo53_001193_orf24710 | gene | Pfam | 27 | 191 | 1.70E-26 | Target=null 27 191;ID=match$484_27_191;signature_desc=Borrelial persistence in ticks protein A;Name=PF17044;status=T;Dbxref="InterPro:IPR031471" |
| gene-bcCo53_001195_orf24734 | gene | Pfam | 35 | 201 | 4.40E-23 | Target=null 35 201;ID=match$291_35_201;signature_desc=Borrelial persistence in ticks protein A;Name=PF17044;status=T;Dbxref="InterPro:IPR031471" |
| gene-bcCo53_001197_orf24760 | gene | Pfam | 35 | 197 | 1.00E-23 | Target=null 35 197;ID=match$125_35_197;signature_desc=Borrelial persistence in ticks protein A;Name=PF17044;status=T;Dbxref="InterPro:IPR031471" |
| gene-bcCo53_001199_orf24787 | gene | Pfam | 30 | 202 | 1.60E-15 | Target=null 30 202;ID=match$955_30_202;signature_desc=Borrelial persistence in ticks protein A;Name=PF17044;status=T;Dbxref="InterPro:IPR031471" |
| gene-bcCo53_001201_orf24816 | gene | Pfam | 35 | 201 | 1.00E-27 | Target=null 35 201;ID=match$1375_35_201;signature_desc=Borrelial persistence in ticks protein A;Name=PF17044;status=T;Dbxref="InterPro:IPR031471" |
| gene-bcCo53_001203_orf24845 | gene | Pfam | 32 | 200 | 3.40E-27 | Target=null 32 200;ID=match$997_32_200;signature_desc=Borrelial persistence in ticks protein A;Name=PF17044;status=T;Dbxref="InterPro:IPR031471" |
| gene-bcCo53_001205_orf24873 | gene | Pfam | 31 | 200 | 3.70E-29 | Target=null 31 200;ID=match$2307_31_200;signature_desc=Borrelial persistence in ticks protein A;Name=PF17044;status=T;Dbxref="InterPro:IPR031471" |
| gene-bcCo53_001206_orf24888 | gene | Pfam | 31 | 198 | 8.70E-23 | Target=null 31 198;ID=match$2765_31_198;signature_desc=Borrelial persistence in ticks protein A;Name=PF17044;status=T;Dbxref="InterPro:IPR031471" |
| gene-bcCo53_001207_orf24903 | gene | Pfam | 31 | 205 | 1.30E-31 | Target=null 31 205;ID=match$1891_31_205;signature_desc=Borrelial persistence in ticks protein A;Name=PF17044;status=T;Dbxref="InterPro:IPR031471" |
| gene-bcCo53_001212_orf25026 | gene | Pfam | 17 | 184 | 1.10E-31 | Target=null 17 184;ID=match$142_17_184;signature_desc=Borrelial persistence in ticks protein A;Name=PF17044;status=T;Dbxref="InterPro:IPR031471" |
| ***Borrelia puertoricensis* SUM** | | | | | | |
| gene-bpuSUM_001533_orf29709 | gene | Pfam | 38 | 211 | 3.10E-21 | Target=null 38 211;ID=match$295_38_211;signature_desc=Borrelial persistence in ticks protein A;Name=PF17044;status=T;Dbxref="InterPro:IPR031471" |
| gene-bpuSUM_001534_orf29731 | gene | Pfam | 224 | 388 | 3.80E-27 | Target=null 224 388;ID=match$3369_224_388;signature_desc=Borrelial persistence in ticks protein A;Name=PF17044;status=T;Dbxref="InterPro:IPR031471" |
| gene-bpuSUM_001540_orf29870 | gene | Pfam | 50 | 212 | 3.10E-31 | Target=null 50 212;ID=match$2318_50_212;signature_desc=Borrelial persistence in ticks protein A;Name=PF17044;status=T;Dbxref="InterPro:IPR031471" |
| ***Borrelia parkeri* SLO** | | | | | | |
| gene-bpSLO_001117_orf23803 | gene | Pfam | 35 | 202 | 8.30E-24 | Target=null 35 202;ID=match$766_35_202;signature_desc=Borrelial persistence in ticks protein A;Name=PF17044;status=T;Dbxref="InterPro:IPR031471" |
| gene-bpSLO_001119_orf23829 | gene | Pfam | 35 | 202 | 9.90E-25 | Target=null 35 202;ID=match$1925_35_202;signature_desc=Borrelial persistence in ticks protein A;Name=PF17044;status=T;Dbxref="InterPro:IPR031471" |
| gene-bpSLO_001120_orf23842 | gene | Pfam | 35 | 203 | 4.30E-30 | Target=null 35 203;ID=match$549_35_203;signature_desc=Borrelial persistence in ticks protein A;Name=PF17044;status=T;Dbxref="InterPro:IPR031471" |
| gene-bpSLO_001121_orf23862 | gene | Pfam | 205 | 370 | 5.70E-26 | Target=null 205 370;ID=match$1131_205_370;signature_desc=Borrelial persistence in ticks protein A;Name=PF17044;status=T;Dbxref="InterPro:IPR031471" |
| gene-bpSLO_001123_orf23898 | gene | Pfam | 35 | 207 | 2.00E-21 | Target=null 35 207;ID=match$233_35_207;signature_desc=Borrelial persistence in ticks protein A;Name=PF17044;status=T;Dbxref="InterPro:IPR031471" |
| gene-bpSLO_001125_orf23927 | gene | Pfam | 36 | 211 | 1.60E-32 | Target=null 36 211;ID=match$2635_36_211;signature_desc=Borrelial persistence in ticks protein A;Name=PF17044;status=T;Dbxref="InterPro:IPR031471" |
| gene-bpSLO_001128_orf23967 | gene | Pfam | 12 | 185 | 1.00E-27 | Target=null 12 185;ID=match$3146_12_185;signature_desc=Borrelial persistence in ticks protein A;Name=PF17044;status=T;Dbxref="InterPro:IPR031471" |
| gene-bpSLO_001135_orf24124 | gene | Pfam | 53 | 212 | 5.00E-29 | Target=null 53 212;ID=match$594_53_212;signature_desc=Borrelial persistence in ticks protein A;Name=PF17044;status=T;Dbxref="InterPro:IPR031471" |
| ***Borrelia venezuelensis* RMA01** | | | | | | |
| gene-bvRMA01_000992_orf22101 | gene | Pfam | 2 | 66 | 4.40E-10 | Target=null 2 66;ID=match$2634_2_66;signature_desc=Borrelial persistence in ticks protein A;Name=PF17044;status=T;Dbxref="InterPro:IPR031471" |
| gene-bvRMA01_000993_orf22108 | gene | Pfam | 34 | 115 | 3.20E-07 | Target=null 34 115;ID=match$2640_34_115;signature_desc=Borrelial persistence in ticks protein A;Name=PF17044;status=T;Dbxref="InterPro:IPR031471" |
| gene-bvRMA01_001002_orf22225 | gene | Pfam | 46 | 212 | 2.00E-29 | Target=null 46 212;ID=match$1139_46_212;signature_desc=Borrelial persistence in ticks protein A;Name=PF17044;status=T;Dbxref="InterPro:IPR031471" |
| ***Borrelia turicatae* 91E135** | | | | | | |
| gene-bt91E135_001164_orf24394 | gene | Pfam | 36 | 202 | 4.50E-22 | Target=null 36 202;ID=match$1617_36_202;signature_desc=Borrelial persistence in ticks protein A;Name=PF17044;status=T;Dbxref="InterPro:IPR031471" |
| gene-bt91E135_001165_orf24409 | gene | Pfam | 34 | 203 | 1.40E-28 | Target=null 34 203;ID=match$464_34_203;signature_desc=Borrelial persistence in ticks protein A;Name=PF17044;status=T;Dbxref="InterPro:IPR031471" |
| gene-bt91E135_001166_orf24430 | gene | Pfam | 206 | 370 | 1.70E-23 | Target=null 206 370;ID=match$196_206_370;signature_desc=Borrelial persistence in ticks protein A;Name=PF17044;status=T;Dbxref="InterPro:IPR031471" |
| gene-bt91E135_001168_orf24466 | gene | Pfam | 36 | 207 | 8.30E-24 | Target=null 36 207;ID=match$2567_36_207;signature_desc=Borrelial persistence in ticks protein A;Name=PF17044;status=T;Dbxref="InterPro:IPR031471" |
| gene-bt91E135_001170_orf24501 | gene | Pfam | 39 | 215 | 1.40E-30 | Target=null 39 215;ID=match$2452_39_215;signature_desc=Borrelial persistence in ticks protein A;Name=PF17044;status=T;Dbxref="InterPro:IPR031471" |
| gene-bt91E135_001176_orf24565 | gene | Pfam | 31 | 201 | 4.50E-27 | Target=null 31 201;ID=match$717_31_201;signature_desc=Borrelial persistence in ticks protein A;Name=PF17044;status=T;Dbxref="InterPro:IPR031471" |
| gene-bt91E135_001182_orf24694 | gene | Pfam | 46 | 212 | 1.80E-29 | Target=null 46 212;ID=match$1459_46_212;signature_desc=Borrelial persistence in ticks protein A;Name=PF17044;status=T;Dbxref="InterPro:IPR031471" |
| ***Borrelia turicatae* BTE5EL** | | | | | | |
| gene-btBTE5EL_001175_orf24641 | gene | Pfam | 36 | 202 | 4.50E-22 | Target=null 36 202;ID=match$1663_36_202;signature_desc=Borrelial persistence in ticks protein A;Name=PF17044;status=T;Dbxref="InterPro:IPR031471" |
| gene-btBTE5EL_001176_orf24656 | gene | Pfam | 34 | 203 | 1.40E-28 | Target=null 34 203;ID=match$486_34_203;signature_desc=Borrelial persistence in ticks protein A;Name=PF17044;status=T;Dbxref="InterPro:IPR031471" |
| gene-btBTE5EL_001177_orf24677 | gene | Pfam | 206 | 370 | 1.40E-23 | Target=null 206 370;ID=match$1547_206_370;signature_desc=Borrelial persistence in ticks protein A;Name=PF17044;status=T;Dbxref="InterPro:IPR031471" |
| gene-btBTE5EL_001179_orf24713 | gene | Pfam | 36 | 207 | 8.30E-24 | Target=null 36 207;ID=match$2692_36_207;signature_desc=Borrelial persistence in ticks protein A;Name=PF17044;status=T;Dbxref="InterPro:IPR031471" |
| gene-btBTE5EL_001181_orf24748 | gene | Pfam | 39 | 215 | 1.40E-30 | Target=null 39 215;ID=match$2530_39_215;signature_desc=Borrelial persistence in ticks protein A;Name=PF17044;status=T;Dbxref="InterPro:IPR031471" |
| gene-btBTE5EL_001186_orf24802 | gene | Pfam | 31 | 201 | 4.50E-27 | Target=null 31 201;ID=match$749_31_201;signature_desc=Borrelial persistence in ticks protein A;Name=PF17044;status=T;Dbxref="InterPro:IPR031471" |
| gene-btBTE5EL_001192_orf24933 | gene | Pfam | 46 | 212 | 1.80E-29 | Target=null 46 212;ID=match$1479_46_212;signature_desc=Borrelial persistence in ticks protein A;Name=PF17044;status=T;Dbxref="InterPro:IPR031471" |

likely a pseudogene due to gene truncation

*bbj47*- like gene
