## Supplemental File 3 for "Comparative genomics analysis of three conserved plasmid families in the Western Hemisphere soft tick-borne relapsing fever borreliae provides insight into variation in genome structure and antigenic variation systems"

```
##example commands

##Last aligner (v1418), self-dot plot of megaplasmid sequences. A fasta file for each megaplasmid was
used for self-dot plot analysis.
#make a database using lastdb, a prefix for the sample, and the megaplasmid sequence to be analyzed.
lastdb database_prefix megaplasmid.fasta
#align the same megaplasmid sequence against itself with only significant alignments reported in the
megaplasmid.maf file
lastal -E0.05 database megaplasmid.fasta > megaplasmid.maf
#generate dotplot with forward alignments as black and reverse alignments as red. Outputs the dot plot
as a png file.
last-dotplot -c black -r red megaplasmid.maf megaplasmid.png

##Flexidot (v1.06) dot plot analysis of the F27 plasmid.
python flexidot_v1.06.py -i /path/to/sequences/ -p 0 -D y -f 1 -k 20 -r y -x n -m 3 -P 15

##Maximum likelihood phylogenies.
#Nucleotide sequences of vmp loci and bptA genes identified by InterProScan were aligned with MAFFT
(v7.475)
mafft --auto /path/to/sequences/ > sequence.aln
#ML phylogeny using IQTREE2
iqtree2 -s /path/to/sequence.aln -m MFP -B 1000  -T AUTO
#collapse resulting tree at branches with less than 50% ultra-fast bootstraps for vlp analysis
iqtree2 -t /path/to/tree/output/ -minsupnew 50

##cluster analysis of vsp signal peptides
#CD-HIT (v4.8.1) was used to cluster the first 20 amino acids of the vsp genes including the first 20
amino acids of OspC from B. burgdorferi B31.
cd-hit -c 0.8 -i /path/to/sequences -o /path/to/output

##Long read mapping for B.venezuelensis RMA01 lp35 and lp37 analysis
#first filter reads out <q10 and <15kb with NanoFilt (v2.8.0)
NanoFilt -q 10 -l 15000 reads.fq > filt_reads.fq
#map reads to B. venezuelensis RMA01 genome (GCA_023035835.1) using minimap2 (v2.24-r1122)
minimap2 -ax map-ont B.venezuelensis_genome.fa filt_reads.fq > mapped_reads.sam
#convert mapped reads to bam file, selecting for primary mapped reads with a mapping quality better
than q30 that mapped to lp35 or lp37. The fasta headers in the B. venezuelensis RMA01 genome that was
mapped with minimap2 were >lp35 and >lp37 for the respective plasmids.
samtools view -b -F 0x104 -q20 lp35 > primary_mapped_reads_lp35.bam
samtools view -b -F 0x104 -q20 lp37 > primary_mapped_reads_lp37.bam
#sort the bam files
samtools sort primary_mapped_reads_lp35.bam > sorted_primary_mapped_reads_lp35.bam
samtools sort primary_mapped_reads_lp37.bam > sorted_primary_mapped_reads_lp37.bam
#index sorted bam files for visualization in IGV
samtools index sorted_primary_mapped_reads_lp35.bam
samtools index sorted_primary_mapped_reads_lp37.bam

##Adaptive sampling analysis for B. venezuelensis RMA01 lp35 and lp37
#basecall FAST5 files with Guppy (v6.4.2)
guppy_basecaller --input_path /path/to/fast5_files --save_path basecalled_data --config
dna_r10.4.1_e8.2_400bps_sup.cfg -x 'auto' -r --detect_mid_strand_adapter --calib_detect --min_qscore 10
#demultiplex fastq files using Guppy (v6.4.2)
guppy_barcoder -i /path/to/basecalled_data/pass -s demultiplexed_data --barcode_kits "SQK-NBD114.24" -x
'auto' --detect_mid_strand_barcoders
#Mapping and analysis of the adaptive sampling was performed the same as described above for the
original read data for the B. venezuelensis RMA01 genome, using the same versions of software. These
data were visualized in IGV as well.
```
