## Supplemental File 4 for "Comparative genomics analysis of three conserved plasmid families in the Western Hemisphere soft tick-borne relapsing fever borreliae provides insight into variation in genome structure and antigenic variation systems"

This file contains the loci classified as either a vsp or vlp for each of the isolates investigated. Vlp genes are further classified by subfamily.

| | | | | | | | | | | |
|---|---|---|---|---|---|---|---|---|---|---|
| Vsp | replicon | gene/pseudo | | | | | | | | |
| gene-baBA2_0009 | lp27 | gene | Pfam | protein_matc | 37 | 205 | 4.3E-58 | + | . | Target=null 37 205;ID=match$1484_37_205;signature_desc=Lipoprotein;Name=PF01441;status=T;Dbxref="InterPro:IPR001800" |

Vlp
none

| Vsp | replicon | gene/pseudo | | | | | | | | |
|---|---|---|---|---|---|---|---|---|---|---|
| gene-bhDAH_001040 ( | cp90 | gene | Pfam | protein_matc | 38 | 205 | 6E-59 | + | . | Target=null 38 205;ID=match$1421_38_205;signature_desc=Lipoprotein;Name=PF01441;status=T;Dbxref="InterPro:IPR001800" |
| gene-bhDAH_001041 ( | cp90 | pseudo | Pfam | protein_matc | 21 | 61 | 7.3E-13 | + | . | Target=null 21 61;ID=match$1591_21_61;signature_desc=Lipoprotein;Name=PF01441;status=T;Dbxref="InterPro:IPR001800" |
| gene-bhDAH_001053 ( | cp90 | pseudo | Pfam | protein_matc | 257 | 398 | 4.7E-32 | + | . | Target=null 257 398;ID=match$493_257_398;signature_desc=Lipoprotein;Name=PF01441;status=T;Dbxref="InterPro:IPR001800" |
| gene-bhDAH_001055 ( | cp90 | gene | Pfam | protein_matc | 2 | 65 | 9E-10 | + | . | Target=null 2 65;ID=match$1289_2_65;signature_desc=Lipoprotein;Name=PF01441;status=T;Dbxref="InterPro:IPR001800" |
| gene-bhDAH_001058 ( | cp90 | gene | Pfam | protein_matc | 47 | 209 | 5.8E-38 | + | . | Target=null 47 209;ID=match$2796_47_209;signature_desc=Lipoprotein;Name=PF01441;status=T;Dbxref="InterPro:IPR001800" |
| gene-bhDAH_001089 ( | cp90 | pseudo | Pfam | protein_matc | 40 | 114 | 6.7E-27 | + | . | Target=null 40 114;ID=match$103_40_114;signature_desc=Lipoprotein;Name=PF01441;status=T;Dbxref="InterPro:IPR001800" |
| gene-bhDAH_001089 ( | cp90 | pseudo | Pfam | protein_matc | 2 | 35 | 1.8E-08 | + | . | Target=null 2 35;ID=match$934_2_35;signature_desc=Lipoprotein;Name=PF01441;status=T;Dbxref="InterPro:IPR001800" |
| gene-bhDAH_001093 ( | cp90 | pseudo | Pfam | protein_matc | 2 | 167 | 2.3E-52 | + | . | Target=null 2 167;ID=match$1056_2_167;signature_desc=Lipoprotein;Name=PF01441;status=T;Dbxref="InterPro:IPR001800" |
| gene-bhDAH_001101 ( | cp90 | gene | Pfam | protein_matc | 1 | 169 | 2.7E-67 | + | . | Target=null 1 169;ID=match$2690_1_169;signature_desc=Lipoprotein;Name=PF01441;status=T;Dbxref="InterPro:IPR001800" |
| gene-bhDAH_001121 ( | cp90 | pseudo | Pfam | protein_matc | 38 | 165 | 8.3E-36 | + | . | Target=null 38 165;ID=match$528_38_165;signature_desc=Lipoprotein;Name=PF01441;status=T;Dbxref="InterPro:IPR001800" |
| gene-bhDAH_001121 ( | cp90 | pseudo | Pfam | protein_matc | 3 | 50 | 1.1E-10 | + | . | Target=null 3 50;ID=match$2604_3_50;signature_desc=Lipoprotein;Name=PF01441;status=T;Dbxref="InterPro:IPR001800" |
| gene-bhDAH_001328 ( | lp26 | gene | Pfam | protein_matc | 29 | 197 | 2.1E-66 | + | . | Target=null 29 197;ID=match$2243_29_197;signature_desc=Lipoprotein;Name=PF01441;status=T;Dbxref="InterPro:IPR001800" |
| gene-bhDAH_001331 ( | lp26 | pseudo | Pfam | protein_matc | 38 | 79 | 4E-16 | + | . | Target=null 38 79;ID=match$1921_38_79;signature_desc=Lipoprotein;Name=PF01441;status=T;Dbxref="InterPro:IPR001800" |
| gene-bhDAH_001331 ( | lp26 | pseudo | Pfam | protein_matc | 2 | 126 | 8.2E-41 | + | . | Target=null 2 126;ID=match$1455_2_126;signature_desc=Lipoprotein;Name=PF01441;status=T;Dbxref="InterPro:IPR001800" |
| gene-bhDAH_001334 ( | lp26 | pseudo | Pfam | protein_matc | 29 | 70 | 2.2E-17 | + | . | Target=null 29 70;ID=match$1948_29_70;signature_desc=Lipoprotein;Name=PF01441;status=T;Dbxref="InterPro:IPR001800" |
| gene-bhDAH_001334 ( | lp26 | pseudo | Pfam | protein_matc | 2 | 127 | 1.1E-40 | + | . | Target=null 2 127;ID=match$1050_2_127;signature_desc=Lipoprotein;Name=PF01441;status=T;Dbxref="InterPro:IPR001800" |
| gene-bhDAH_001337 ( | lp26 | gene | Pfam | protein_matc | 38 | 207 | 2.7E-66 | + | . | Target=null 38 207;ID=match$204_38_207;signature_desc=Lipoprotein;Name=PF01441;status=T;Dbxref="InterPro:IPR001800" |
| gene-bhDAH_001354 ( | lp27 | gene | Pfam | protein_matc | 29 | 200 | 8.7E-45 | + | . | Target=null 29 200;ID=match$1681_29_200;signature_desc=Lipoprotein;Name=PF01441;status=T;Dbxref="InterPro:IPR001800" |
| gene-bhDAH_001405 ( | lp31 | gene | Pfam | protein_matc | 22 | 192 | 1.3E-51 | + | . | Target=null 22 192;ID=match$956_22_192;signature_desc=Lipoprotein;Name=PF01441;status=T;Dbxref="InterPro:IPR001800" |
| gene-bhDAH_001407 ( | lp31 | gene | Pfam | protein_matc | 1 | 116 | 8.1E-35 | + | . | Target=null 1 116;ID=match$929_1_116;signature_desc=Lipoprotein;Name=PF01441;status=T;Dbxref="InterPro:IPR001800" |
| gene-bhDAH_001408 ( | lp31 | gene | Pfam | protein_matc | 38 | 207 | 1.9E-58 | + | . | Target=null 38 207;ID=match$19_38_207;signature_desc=Lipoprotein;Name=PF01441;status=T;Dbxref="InterPro:IPR001800" |
| gene-bhDAH_001419 ( | lp31 | gene | Pfam | protein_matc | 38 | 210 | 2.6E-58 | + | . | Target=null 38 210;ID=match$2482_38_210;signature_desc=Lipoprotein;Name=PF01441;status=T;Dbxref="InterPro:IPR001800" |
| gene-bhDAH_001420 ( | lp31 | gene | Pfam | protein_matc | 29 | 197 | 1.1E-62 | + | . | Target=null 29 197;ID=match$2024_29_197;signature_desc=Lipoprotein;Name=PF01441;status=T;Dbxref="InterPro:IPR001800" |
| gene-bhDAH_001422 ( | lp31 | pseudo | Pfam | protein_matc | 43 | 214 | 1.2E-53 | + | . | Target=null 43 214;ID=match$511_43_214;signature_desc=Lipoprotein;Name=PF01441;status=T;Dbxref="InterPro:IPR001800" |
| gene-bhDAH_001423 ( | lp31 | pseudo | Pfam | protein_matc | 7 | 172 | 1E-40 | + | . | Target=null 7 172;ID=match$1227_7_172;signature_desc=Lipoprotein;Name=PF01441;status=T;Dbxref="InterPro:IPR001800" |
| gene-bhDAH_001447 ( | lp59 | gene | Pfam | protein_matc | 29 | 199 | 2.1E-60 | + | . | Target=null 29 199;ID=match$1337_29_199;signature_desc=Lipoprotein;Name=PF01441;status=T;Dbxref="InterPro:IPR001800" |
| gene-bhDAH_001449 ( | lp59 | gene | Pfam | protein_matc | 25 | 188 | 4.6E-49 | + | . | Target=null 25 188;ID=match$562_25_188;signature_desc=Lipoprotein;Name=PF01441;status=T;Dbxref="InterPro:IPR001800" |
| gene-bhDAH_001450 ( | lp59 | gene | Pfam | protein_matc | 38 | 207 | 1.6E-52 | + | . | Target=null 38 207;ID=match$92_38_207;signature_desc=Lipoprotein;Name=PF01441;status=T;Dbxref="InterPro:IPR001800" |
| gene-bhDAH_001454 ( | lp59 | gene | Pfam | protein_matc | 38 | 209 | 1.3E-44 | + | . | Target=null 38 209;ID=match$1000_38_209;signature_desc=Lipoprotein;Name=PF01441;status=T;Dbxref="InterPro:IPR001800" |
| gene-bhDAH_001455 ( | lp59 | gene | Pfam | protein_matc | 38 | 206 | 6.3E-60 | + | . | Target=null 38 206;ID=match$1364_38_206;signature_desc=Lipoprotein;Name=PF01441;status=T;Dbxref="InterPro:IPR001800" |
| gene-bhDAH_001456 ( | lp59 | gene | Pfam | protein_matc | 38 | 206 | 3.2E-66 | + | . | Target=null 38 206;ID=match$2899_38_206;signature_desc=Lipoprotein;Name=PF01441;status=T;Dbxref="InterPro:IPR001800" |
| gene-bhDAH_001490 ( | lp59 | gene | Pfam | protein_matc | 38 | 201 | 6E-53 | + | . | Target=null 38 201;ID=match$1958_38_201;signature_desc=Lipoprotein;Name=PF01441;status=T;Dbxref="InterPro:IPR001800" |

| Vlp | | | Subfamily | | | | | | | | |
|---|---|---|---|---|---|---|---|---|---|---|---|
| gene-bhDAH_001038 ( | cp90 | gene | gamma | Pfam | protein_matc | 2 | 69 | 3.2E-10 | + | . | Target=null 2 69;ID=match$1517_2_69;signature_desc=Borrelia lipoprotein;Name=PF00921;status=T;Dbxref="InterPro:IPR000680" |
| gene-bhDAH_001039 ( | cp90 | gene | gamma | Pfam | protein_matc | 4 | 97 | 1.5E-14 | + | . | Target=null 4 97;ID=match$2620_4_97;signature_desc=Borrelia lipoprotein;Name=PF00921;status=T;Dbxref="InterPro:IPR000680" |
| gene-bhDAH_001048 ( | cp90 | gene | gamma | Pfam | protein_matc | 30 | 311 | 2E-83 | + | . | Target=null 30 311;ID=match$2791_30_311;signature_desc=Borrelia lipoprotein;Name=PF00921;status=T;Dbxref="InterPro:IPR000680" |
| gene-bhDAH_001049 ( | cp90 | gene | delta | Pfam | protein_matc | 43 | 322 | 1.6E-84 | + | . | Target=null 43 322;ID=match$2080_43_322;signature_desc=Borrelia lipoprotein;Name=PF00921;status=T;Dbxref="InterPro:IPR000680" |
| gene-bhDAH_001050 ( | cp90 | gene | gamma | Pfam | protein_matc | 43 | 321 | 6E-93 | + | . | Target=null 43 321;ID=match$2084_43_321;signature_desc=Borrelia lipoprotein;Name=PF00921;status=T;Dbxref="InterPro:IPR000680" |
| gene-bhDAH_001051 ( | cp90 | pseudo | beta | Pfam | protein_matc | 10 | 265 | 4E-42 | + | . | Target=null 10 265;ID=match$1812_10_265;signature_desc=Borrelia lipoprotein;Name=PF00921;status=T;Dbxref="InterPro:IPR000680" |
| gene-bhDAH_001052 ( | cp90 | pseudo | gamma | Pfam | protein_matc | 7 | 53 | 2.8E-15 | + | . | Target=null 7 53;ID=match$363_7_53;signature_desc=Borrelia lipoprotein;Name=PF00921;status=T;Dbxref="InterPro:IPR000680" |
| gene-bhDAH_001052 ( | cp90 | pseudo | | Pfam | protein_matc | 1 | 240 | 6.9E-60 | + | . | Target=null 1 240;ID=match$265_1_240;signature_desc=Borrelia lipoprotein;Name=PF00921;status=T;Dbxref="InterPro:IPR000680" |
| gene-bhDAH_001053 ( | cp90 | pseudo | delta | Pfam | protein_matc | 43 | 88 | 4.2E-16 | + | . | Target=null 43 88;ID=match$438_43_88;signature_desc=Borrelia lipoprotein;Name=PF00921;status=T;Dbxref="InterPro:IPR000680" |
| gene-bhDAH_001053 ( | cp90 | pseudo | | Pfam | protein_matc | 2 | 249 | 4.5E-56 | + | . | Target=null 2 249;ID=match$492_2_249;signature_desc=Borrelia lipoprotein;Name=PF00921;status=T;Dbxref="InterPro:IPR000680" |
| gene-bhDAH_001054 ( | cp90 | gene | gamma | Pfam | protein_matc | 11 | 271 | 4.8E-75 | + | . | Target=null 11 271;ID=match$1767_11_271;signature_desc=Borrelia lipoprotein;Name=PF00921;status=T;Dbxref="InterPro:IPR000680" |
| gene-bhDAH_001055 ( | cp90 | gene | delta | Pfam | protein_matc | 25 | 271 | 7E-56 | + | . | Target=null 25 271;ID=match$1291_25_271;signature_desc=Borrelia lipoprotein;Name=PF00921;status=T;Dbxref="InterPro:IPR000680" |
| gene-bhDAH_001056 ( | cp90 | gene | gamma | Pfam | protein_matc | 34 | 312 | 5.4E-93 | + | . | Target=null 34 312;ID=match$2655_34_312;signature_desc=Borrelia lipoprotein;Name=PF00921;status=T;Dbxref="InterPro:IPR000680" |
| gene-bhDAH_001057 ( | cp90 | pseudo | gamma | Pfam | protein_matc | 25 | 317 | 6.8E-94 | + | . | Target=null 25 317;ID=match$2987_25_317;signature_desc=Borrelia lipoprotein;Name=PF00921;status=T;Dbxref="InterPro:IPR000680" |
| gene-bhDAH_001059 ( | cp90 | pseudo | gamma | Pfam | protein_matc | 25 | 317 | 3.6E-91 | + | . | Target=null 25 317;ID=match$2272_25_317;signature_desc=Borrelia lipoprotein;Name=PF00921;status=T;Dbxref="InterPro:IPR000680" |
| gene-bhDAH_001060 ( | cp90 | gene | beta | Pfam | protein_matc | 47 | 336 | 2.3E-51 | + | . | Target=null 47 336;ID=match$764_47_336;signature_desc=Borrelia lipoprotein;Name=PF00921;status=T;Dbxref="InterPro:IPR000680" |
| gene-bhDAH_001069 ( | cp90 | gene | delta | Pfam | protein_matc | 43 | 335 | 1.6E-86 | + | . | Target=null 43 335;ID=match$197_43_335;signature_desc=Borrelia lipoprotein;Name=PF00921;status=T;Dbxref="InterPro:IPR000680" |
| gene-bhDAH_001070 ( | cp90 | pseudo | gamma | Pfam | protein_matc | 13 | 45 | 1.1E-11 | + | . | Target=null 13 45;ID=match$378_13_45;signature_desc=Borrelia lipoprotein;Name=PF00921;status=T;Dbxref="InterPro:IPR000680" |
| gene-bhDAH_001092 ( | cp90 | gene | alpha | Pfam | protein_matc | 59 | 348 | 1.4E-67 | + | . | Target=null 59 348;ID=match$1719_59_348;signature_desc=Borrelia lipoprotein;Name=PF00921;status=T;Dbxref="InterPro:IPR000680" |
| gene-bhDAH_001094 ( | cp90 | gene | beta | Pfam | protein_matc | 55 | 345 | 1.9E-42 | + | . | Target=null 55 345;ID=match$408_55_345;signature_desc=Borrelia lipoprotein;Name=PF00921;status=T;Dbxref="InterPro:IPR000680" |
| gene-bhDAH_001095 ( | cp90 | gene | delta | Pfam | protein_matc | 43 | 326 | 5.6E-83 | + | . | Target=null 43 326;ID=match$1735_43_326;signature_desc=Borrelia lipoprotein;Name=PF00921;status=T;Dbxref="InterPro:IPR000680" |
| gene-bhDAH_001096 ( | cp90 | gene | alpha | Pfam | protein_matc | 27 | 327 | 7.7E-64 | + | . | Target=null 27 327;ID=match$1180_27_327;signature_desc=Borrelia lipoprotein;Name=PF00921;status=T;Dbxref="InterPro:IPR000680" |
| gene-bhDAH_001097 ( | cp90 | gene | beta | Pfam | protein_matc | 20 | 310 | 6.2E-49 | + | . | Target=null 20 310;ID=match$1029_20_310;signature_desc=Borrelia lipoprotein;Name=PF00921;status=T;Dbxref="InterPro:IPR000680" |
| gene-bhDAH_001099 ( | cp90 | gene | gamma | Pfam | protein_matc | 43 | 207 | 5.3E-44 | + | . | Target=null 43 207;ID=match$2900_43_207;signature_desc=Borrelia lipoprotein;Name=PF00921;status=T;Dbxref="InterPro:IPR000680" |
| gene-bhDAH_001099 ( | cp90 | gene | | Pfam | protein_matc | 200 | 275 | 4.1E-28 | + | . | Target=null 200 275;ID=match$2900_200_275;signature_desc=Borrelia lipoprotein;Name=PF00921;status=T;Dbxref="InterPro:IPR000680" |
| gene-bhDAH_001100 ( | cp90 | gene | delta | Pfam | protein_matc | 20 | 304 | 4.8E-87 | + | . | Target=null 20 304;ID=match$80_20_304;signature_desc=Borrelia lipoprotein;Name=PF00921;status=T;Dbxref="InterPro:IPR000680" |
| gene-bhDAH_001102 ( | cp90 | gene | gamma | Pfam | protein_matc | 176 | 251 | 3.6E-27 | + | . | Target=null 176 251;ID=match$230_176_251;signature_desc=Borrelia lipoprotein;Name=PF00921;status=T;Dbxref="InterPro:IPR000680" |
| gene-bhDAH_001102 ( | cp90 | gene | | Pfam | protein_matc | 19 | 183 | 4.1E-44 | + | . | Target=null 19 183;ID=match$230_19_183;signature_desc=Borrelia lipoprotein;Name=PF00921;status=T;Dbxref="InterPro:IPR000680" |
| gene-bhDAH_001119 ( | cp90 | pseudo | delta | Pfam | protein_matc | 34 | 127 | 3.2E-18 | + | . | Target=null 34 127;ID=match$2703_34_127;signature_desc=Borrelia lipoprotein;Name=PF00921;status=T;Dbxref="InterPro:IPR000680" |
| gene-bhDAH_001120 ( | cp90 | gene | gamma | Pfam | protein_matc | 19 | 306 | 4E-91 | + | . | Target=null 19 306;ID=match$706_19_306;signature_desc=Borrelia lipoprotein;Name=PF00921;status=T;Dbxref="InterPro:IPR000680" |
| gene-bhDAH_001122 ( | cp90 | gene | beta | Pfam | protein_matc | 20 | 312 | 3.6E-49 | + | . | Target=null 20 312;ID=match$2340_20_312;signature_desc=Borrelia lipoprotein;Name=PF00921;status=T;Dbxref="InterPro:IPR000680" |
| gene-bhDAH_001123 ( | cp90 | gene | alpha | Pfam | protein_matc | 59 | 349 | 2.4E-70 | + | . | Target=null 59 349;ID=match$2156_59_349;signature_desc=Borrelia lipoprotein;Name=PF00921;status=T;Dbxref="InterPro:IPR000680" |
| gene-bhDAH_001124 ( | cp90 | pseudo | delta | Pfam | protein_matc | 1 | 168 | 9.1E-33 | + | . | Target=null 1 168;ID=match$2128_1_168;signature_desc=Borrelia lipoprotein;Name=PF00921;status=T;Dbxref="InterPro:IPR000680" |
| gene-bhDAH_001124 ( | cp90 | pseudo | | Pfam | protein_matc | 8 | 39 | 3.1E-10 | + | . | Target=null 8 39;ID=match$1019_8_39;signature_desc=Borrelia lipoprotein;Name=PF00921;status=T;Dbxref="InterPro:IPR000680" |
| gene-bhDAH_001124 ( | cp90 | pseudo | | Pfam | protein_matc | 24 | 58 | 1.2E-08 | + | . | Target=null 24 58;ID=match$1994_24_58;signature_desc=Borrelia lipoprotein;Name=PF00921;status=T;Dbxref="InterPro:IPR000680" |
| gene-bhDAH_001125 ( | cp90 | gene | delta | Pfam | protein_matc | 53 | 75 | 0.000022 | + | . | Target=null 53 75;ID=match$2545_53_75;signature_desc=Borrelia lipoprotein;Name=PF00921;status=T;Dbxref="InterPro:IPR000680" |
| gene-bhDAH_001128 ( | cp90 | pseudo | gamma | Pfam | protein_matc | 13 | 45 | 1.1E-11 | + | . | Target=null 13 45;ID=match$376_13_45;signature_desc=Borrelia lipoprotein;Name=PF00921;status=T;Dbxref="InterPro:IPR000680" |
| gene-bhDAH_001129 ( | cp90 | gene | delta | Pfam | protein_matc | 43 | 326 | 2.9E-84 | + | . | Target=null 43 326;ID=match$614_43_326;signature_desc=Borrelia lipoprotein;Name=PF00921;status=T;Dbxref="InterPro:IPR000680" |
| gene-bhDAH_001130 ( | cp90 | gene | gamma | Pfam | protein_matc | 5 | 38 | 5.7E-08 | + | . | Target=null 5 38;ID=match$2909_5_38;signature_desc=Borrelia lipoprotein;Name=PF00921;status=T;Dbxref="InterPro:IPR000680" |
| gene-bhDAH_001326 ( | lp26 | gene | gamma | Pfam | protein_matc | 19 | 296 | 8.1E-80 | + | . | Target=null 19 296;ID=match$1350_19_296;signature_desc=Borrelia lipoprotein;Name=PF00921;status=T;Dbxref="InterPro:IPR000680" |
| gene-bhDAH_001327 ( | lp26 | gene | delta | Pfam | protein_matc | 43 | 332 | 6.5E-83 | + | . | Target=null 43 332;ID=match$1139_43_332;signature_desc=Borrelia lipoprotein;Name=PF00921;status=T;Dbxref="InterPro:IPR000680" |
| gene-bhDAH_001329 ( | lp26 | gene | gamma | Pfam | protein_matc | 29 | 313 | 4.1E-90 | + | . | Target=null 29 313;ID=match$2163_29_313;signature_desc=Borrelia lipoprotein;Name=PF00921;status=T;Dbxref="InterPro:IPR000680" |
| gene-bhDAH_001330 ( | lp26 | pseudo | gamma | Pfam | protein_matc | 2 | 118 | 7.2E-28 | + | . | Target=null 2 118;ID=match$748_2_118;signature_desc=Borrelia lipoprotein;Name=PF00921;status=T;Dbxref="InterPro:IPR000680" |
| gene-bhDAH_001332 ( | lp26 | gene | gamma | Pfam | protein_matc | 9 | 235 | 4.2E-59 | + | . | Target=null 9 235;ID=match$2061_9_235;signature_desc=Borrelia lipoprotein;Name=PF00921;status=T;Dbxref="InterPro:IPR000680" |
| gene-bhDAH_001333 ( | lp26 | pseudo | delta | Pfam | protein_matc | 1 | 88 | 3.9E-15 | + | . | Target=null 1 88;ID=match$1688_1_88;signature_desc=Borrelia lipoprotein;Name=PF00921;status=T;Dbxref="InterPro:IPR000680" |
| gene-bhDAH_001333 ( | lp26 | pseudo | | Pfam | protein_matc | 4 | 84 | 4.2E-14 | + | . | Target=null 4 84;ID=match$152_4_84;signature_desc=Borrelia lipoprotein;Name=PF00921;status=T;Dbxref="InterPro:IPR000680" |
| gene-bhDAH_001333 ( | lp26 | pseudo | | Pfam | protein_matc | 3 | 81 | 4.6E-28 | + | . | Target=null 3 81;ID=match$2106_3_81;signature_desc=Borrelia lipoprotein;Name=PF00921;status=T;Dbxref="InterPro:IPR000680" |
| gene-bhDAH_001335 ( | lp26 | gene | gamma | Pfam | protein_matc | 29 | 304 | 1.1E-81 | + | . | Target=null 29 304;ID=match$2816_29_304;signature_desc=Borrelia lipoprotein;Name=PF00921;status=T;Dbxref="InterPro:IPR000680" |
| gene-bhDAH_001336 ( | lp26 | pseudo | gamma | Pfam | protein_matc | 2 | 66 | 1.4E-09 | + | . | Target=null 2 66;ID=match$1839_2_66;signature_desc=Borrelia lipoprotein;Name=PF00921;status=T;Dbxref="InterPro:IPR000680" |
| gene-bhDAH_001346 ( | lp27 | gene | delta | Pfam | protein_matc | 34 | 314 | 7.4E-87 | + | . | Target=null 34 314;ID=match$2881_34_314;signature_desc=Borrelia lipoprotein;Name=PF00921;status=T;Dbxref="InterPro:IPR000680" |
| gene-bhDAH_001347 ( | lp27 | pseudo | beta | Pfam | protein_matc | 20 | 74 | 2.7E-08 | + | . | Target=null 20 74;ID=match$981_20_74;signature_desc=Borrelia lipoprotein;Name=PF00921;status=T;Dbxref="InterPro:IPR000680" |
| gene-bhDAH_001347 ( | lp27 | pseudo | | Pfam | protein_matc | 7 | 221 | 1.1E-29 | + | . | Target=null 7 221;ID=match$573_7_221;signature_desc=Borrelia lipoprotein;Name=PF00921;status=T;Dbxref="InterPro:IPR000680" |
| gene-bhDAH_001348 ( | lp27 | pseudo | gamma | Pfam | protein_matc | 4 | 148 | 1.1E-42 | + | . | Target=null 4 148;ID=match$1068_4_148;signature_desc=Borrelia lipoprotein;Name=PF00921;status=T;Dbxref="InterPro:IPR000680" |
| gene-bhDAH_001355 ( | lp27 | pseudo | delta | Pfam | protein_matc | 9 | 196 | 8.2E-42 | + | . | Target=null 9 196;ID=match$570_9_196;signature_desc=Borrelia lipoprotein;Name=PF00921;status=T;Dbxref="InterPro:IPR000680" |
| gene-bhDAH_001355 ( | lp27 | pseudo | | Pfam | protein_matc | 33 | 108 | 1.6E-26 | + | . | Target=null 33 108;ID=match$630_33_108;signature_desc=Borrelia lipoprotein;Name=PF00921;status=T;Dbxref="InterPro:IPR000680" |
| gene-bhDAH_001356 ( | lp27 | pseudo | alpha | Pfam | protein_matc | 7 | 74 | 3.3E-15 | + | . | Target=null 7 74;ID=match$963_7_74;signature_desc=Borrelia lipoprotein;Name=PF00921;status=T;Dbxref="InterPro:IPR000680" |
| gene-bhDAH_001356 ( | lp27 | pseudo | | Pfam | protein_matc | 12 | 224 | 1.4E-52 | + | . | Target=null 12 224;ID=match$2821_12_224;signature_desc=Borrelia lipoprotein;Name=PF00921;status=T;Dbxref="InterPro:IPR000680" |
| gene-bhDAH_001357 ( | lp27 | pseudo | beta | Pfam | protein_matc | 20 | 77 | 4.4E-10 | + | . | Target=null 20 77;ID=match$542_20_77;signature_desc=Borrelia lipoprotein;Name=PF00921;status=T;Dbxref="InterPro:IPR000680" |
| gene-bhDAH_001357 ( | lp27 | pseudo | | Pfam | protein_matc | 7 | 241 | 4.2E-34 | + | . | Target=null 7 241;ID=match$268_7_241;signature_desc=Borrelia lipoprotein;Name=PF00921;status=T;Dbxref="InterPro:IPR000680" |
| gene-bhDAH_001358 ( | lp27 | gene | alpha | Pfam | protein_matc | 60 | 351 | 1.9E-71 | + | . | Target=null 60 351;ID=match$2736_60_351;signature_desc=Borrelia lipoprotein;Name=PF00921;status=T;Dbxref="InterPro:IPR000680" |
| gene-bhDAH_001359 ( | lp27 | gene | beta | Pfam | protein_matc | 20 | 308 | 2.1E-42 | + | . | Target=null 20 308;ID=match$1105_20_308;signature_desc=Borrelia lipoprotein;Name=PF00921;status=T;Dbxref="InterPro:IPR000680" |
| gene-bhDAH_001360 ( | lp27 | pseudo | delta | Pfam | protein_matc | 14 | 290 | 8.4E-85 | + | . | Target=null 14 290;ID=match$429_14_290;signature_desc=Borrelia lipoprotein;Name=PF00921;status=T;Dbxref="InterPro:IPR000680" |
| gene-bhDAH_001361 ( | lp27 | pseudo | gamma | Pfam | protein_matc | 12 | 89 | 4.3E-28 | + | . | Target=null 12 89;ID=match$1200_12_89;signature_desc=Borrelia lipoprotein;Name=PF00921;status=T;Dbxref="InterPro:IPR000680" |
| gene-bhDAH_001361 ( | lp27 | pseudo | | Pfam | protein_matc | 9 | 70 | 2.8E-09 | + | . | Target=null 9 70;ID=match$1569_9_70;signature_desc=Borrelia lipoprotein;Name=PF00921;status=T;Dbxref="InterPro:IPR000680" |
| gene-bhDAH_001361 ( | lp27 | pseudo | | Pfam | protein_matc | 6 | 131 | 2.4E-28 | + | . | Target=null 6 131;ID=match$2607_6_131;signature_desc=Borrelia lipoprotein;Name=PF00921;status=T;Dbxref="InterPro:IPR000680" |
| gene-bhDAH_001363 ( | lp27 | gene | delta | Pfam | protein_matc | 13 | 98 | 6.6E-14 | + | . | Target=null 13 98;ID=match$2798_13_98;signature_desc=Borrelia lipoprotein;Name=PF00921;status=T;Dbxref="InterPro:IPR000680" |
| gene-bhDAH_001404 ( | lp31 | gene | gamma | Pfam | protein_matc | 13 | 65 | 2E-10 | + | . | Target=null 13 65;ID=match$1363_13_65;signature_desc=Borrelia lipoprotein;Name=PF00921;status=T;Dbxref="InterPro:IPR000680" |
| gene-bhDAH_001406 ( | lp31 | gene | alpha | Pfam | protein_matc | 57 | 354 | 5.3E-70 | + | . | Target=null 57 354;ID=match$1805_57_354;signature_desc=Borrelia lipoprotein;Name=PF00921;status=T;Dbxref="InterPro:IPR000680" |
| gene-bhDAH_001417 ( | lp31 | gene | delta | Pfam | protein_matc | 43 | 324 | 2E-88 | + | . | Target=null 43 324;ID=match$2580_43_324;signature_desc=Borrelia lipoprotein;Name=PF00921;status=T;Dbxref="InterPro:IPR000680" |
| gene-bhDAH_001418 ( | lp31 | gene | beta | Pfam | protein_matc | 47 | 334 | 4.6E-48 | + | . | Target=null 47 334;ID=match$2044_47_334;signature_desc=Borrelia lipoprotein;Name=PF00921;status=T;Dbxref="InterPro:IPR000680" |
| gene-bhDAH_001421 ( | lp31 | pseudo | gamma | Pfam | protein_matc | 1 | 45 | 5.7E-15 | + | . | Target=null 1 45;ID=match$3046_1_45;signature_desc=Borrelia lipoprotein;Name=PF00921;status=T;Dbxref="InterPro:IPR000680" |
| gene-bhDAH_001421 ( | lp31 | pseudo | | Pfam | protein_matc | 1 | 239 | 5E-60 | + | . | Target=null 1 239;ID=match$959_1_239;signature_desc=Borrelia lipoprotein;Name=PF00921;status=T;Dbxref="InterPro:IPR000680" |
| gene-bhDAH_001429 ( | lp31 | pseudo | gamma | Pfam | protein_matc | 2 | 70 | 2.8E-13 | + | . | Target=null 2 70;ID=match$233_2_70;signature_desc=Borrelia lipoprotein;Name=PF00921;status=T;Dbxref="InterPro:IPR000680" |
| gene-bhDAH_001429 ( | lp31 | pseudo | | Pfam | protein_matc | 2 | 82 | 1.2E-14 | + | . | Target=null 2 82;ID=match$574_2_82;signature_desc=Borrelia lipoprotein;Name=PF00921;status=T;Dbxref="InterPro:IPR000680" |
| gene-bhDAH_001431 ( | lp31 | gene | delta | Pfam | protein_matc | 1 | 46 | 2E-15 | + | . | Target=null 1 46;ID=match$766_1_46;signature_desc=Borrelia lipoprotein;Name=PF00921;status=T;Dbxref="InterPro:IPR000680" |
| gene-bhDAH_001432 ( | lp31 | pseudo | alpha | Pfam | protein_matc | 2 | 64 | 1.9E-09 | + | . | Target=null 2 64;ID=match$1482_2_64;signature_desc=Borrelia lipoprotein;Name=PF00921;status=T;Dbxref="InterPro:IPR000680" |
| gene-bhDAH_001433 ( | lp31 | gene | beta | Pfam | protein_matc | 6 | 46 | 3.2E-11 | + | . | Target=null 6 46;ID=match$2360_6_46;signature_desc=Borrelia lipoprotein;Name=PF00921;status=T;Dbxref="InterPro:IPR000680" |
| gene-bhDAH_001440 ( | lp59 | pseudo | beta | Pfam | protein_matc | 5 | 34 | 0.000013 | + | . | Target=null 5 34;ID=match$1250_5_34;signature_desc=Borrelia lipoprotein;Name=PF00921;status=T;Dbxref="InterPro:IPR000680" |
| gene-bhDAH_001441 ( | lp59 | pseudo | gamma | Pfam | protein_matc | 43 | 65 | 0.0000019 | + | . | Target=null 43 65;ID=match$1922_43_65;signature_desc=Borrelia lipoprotein;Name=PF00921;status=T;Dbxref="InterPro:IPR000680" |
| gene-bhDAH_001441 ( | lp59 | pseudo | | Pfam | protein_matc | 1 | 245 | 9.4E-67 | + | . | Target=null 1 245;ID=match$386_1_245;signature_desc=Borrelia lipoprotein;Name=PF00921;status=T;Dbxref="InterPro:IPR000680" |
| gene-bhDAH_001442 ( | lp59 | gene | beta | Pfam | protein_matc | 47 | 331 | 5.9E-45 | + | . | Target=null 47 331;ID=match$2922_47_331;signature_desc=Borrelia lipoprotein;Name=PF00921;status=T;Dbxref="InterPro:IPR000680" |
| gene-bhDAH_001443 ( | lp59 | gene | beta | Pfam | protein_matc | 47 | 340 | 2.2E-48 | + | . | Target=null 47 340;ID=match$961_47_340;signature_desc=Borrelia lipoprotein;Name=PF00921;status=T;Dbxref="InterPro:IPR000680" |
| gene-bhDAH_001444 ( | lp59 | pseudo | gamma | Pfam | protein_matc | 1 | 53 | 7.8E-08 | + | . | Target=null 1 53;ID=match$2458_1_53;signature_desc=Borrelia lipoprotein;Name=PF00921;status=T;Dbxref="InterPro:IPR000680" |
| gene-bhDAH_001444 ( | lp59 | pseudo | | Pfam | protein_matc | 4 | 89 | 1.9E-26 | + | . | Target=null 4 89;ID=match$1759_4_89;signature_desc=Borrelia lipoprotein;Name=PF00921;status=T;Dbxref="InterPro:IPR000680" |
| gene-bhDAH_001446 ( | lp59 | pseudo | alpha | Pfam | protein_matc | 57 | 300 | 3E-47 | + | . | Target=null 57 300;ID=match$849_57_300;signature_desc=Borrelia lipoprotein;Name=PF00921;status=T;Dbxref="InterPro:IPR000680" |
| gene-bhDAH_001446 ( | lp59 | pseudo | | Pfam | protein_matc | 2 | 50 | 9.2E-10 | + | . | Target=null 2 50;ID=match$2120_2_50;signature_desc=Borrelia lipoprotein;Name=PF00921;status=T;Dbxref="InterPro:IPR000680" |
| gene-bhDAH_001448 ( | lp59 | gene | gamma | Pfam | protein_matc | 30 | 322 | 9.8E-92 | + | . | Target=null 30 322;ID=match$2602_30_322;signature_desc=Borrelia lipoprotein;Name=PF00921;status=T;Dbxref="InterPro:IPR000680" |

Duplicates between Vsp and Vlp hits. Did not use for Vsp analysis.

| Vsp | | replicon | gene/pseudo | | | | | | | | |
|---|---|---|---|---|---|---|---|---|---|---|---|
| gene-bhYOR 001254 | orf2 | lp23 | gene | Pfam | protein matc | 25 | 193 | 1.4E-51 | + | . | Target=null 25 193;ID=match$608_25_193;signature_desc=Lipoprotein;Name=PF01441;status=T;Dbxref="InterPro:IPR001800" |
| gene-bhYOR 001256 | orf2 | lp23 | pseudo | Pfam | protein matc | 3 | 165 | 4.9E-55 | + | . | Target=null 3 165;ID=match$2592_3_165;signature_desc=Lipoprotein;Name=PF01441;status=T;Dbxref="InterPro:IPR001800" |
| gene-bhYOR 001259 | orf2 | lp23 | gene | Pfam | protein matc | 38 | 205 | 1E-62 | + | . | Target=null 38 205;ID=match$1182_38_205;signature_desc=Lipoprotein;Name=PF01441;status=T;Dbxref="InterPro:IPR001800" |
| gene-bhYOR 001265 | orf2 | lp46 | pseudo | Pfam | protein matc | 2 | 43 | 1.8E-17 | + | . | Target=null 2 43;ID=match$411_2_43;signature_desc=Lipoprotein;Name=PF01441;status=T;Dbxref="InterPro:IPR001800" |
| gene-bhYOR 001265 | orf2 | lp46 | pseudo | Pfam | protein matc | 13 | 144 | 6.9E-33 | + | . | Target=null 13 144;ID=match$407_13_144;signature_desc=Lipoprotein;Name=PF01441;status=T;Dbxref="InterPro:IPR001800" |
| gene-bhYOR 001266 | orf2 | lp46 | gene | Pfam | protein matc | 38 | 207 | 5.1E-64 | + | . | Target=null 38 207;ID=match$2931_38_207;signature_desc=Lipoprotein;Name=PF01441;status=T;Dbxref="InterPro:IPR001800" |
| gene-bhYOR 001268 | orf2 | lp46 | pseudo | Pfam | protein matc | 25 | 188 | 3.1E-58 | + | . | Target=null 25 188;ID=match$2823_25_188;signature_desc=Lipoprotein;Name=PF01441;status=T;Dbxref="InterPro:IPR001800" |
| gene-bhYOR 001299 | orf2 | lp46 | gene | Pfam | protein matc | 38 | 206 | 2.4E-61 | + | . | Target=null 38 206;ID=match$639_38_206;signature_desc=Lipoprotein;Name=PF01441;status=T;Dbxref="InterPro:IPR001800" |
| gene-bhYOR 001308 | orf2 | lp73 | gene | Pfam | protein matc | 38 | 205 | 4.2E-63 | + | . | Target=null 38 205;ID=match$1719_38_205;signature_desc=Lipoprotein;Name=PF01441;status=T;Dbxref="InterPro:IPR001800" |
| gene-bhYOR 001311 | orf2 | lp73 | gene | Pfam | protein matc | 1 | 83 | 7.9E-23 | + | . | Target=null 1 83;ID=match$654_1_83;signature_desc=Lipoprotein;Name=PF01441;status=T;Dbxref="InterPro:IPR001800" |
| gene-bhYOR 001312 | orf2 | lp73 | pseudo | Pfam | protein matc | 7 | 172 | 1.4E-40 | + | . | Target=null 7 172;ID=match$2640_7_172;signature_desc=Lipoprotein;Name=PF01441;status=T;Dbxref="InterPro:IPR001800" |
| gene-bhYOR 001313 | orf2 | lp73 | pseudo | Pfam | protein matc | 43 | 214 | 1.2E-53 | + | . | Target=null 43 214;ID=match$544_43_214;signature_desc=Lipoprotein;Name=PF01441;status=T;Dbxref="InterPro:IPR001800" |
| gene-bhYOR 001315 | orf2 | lp73 | gene | Pfam | protein matc | 29 | 195 | 1.1E-64 | + | . | Target=null 29 195;ID=match$356_29_195;signature_desc=Lipoprotein;Name=PF01441;status=T;Dbxref="InterPro:IPR001800" |
| gene-bhYOR 001316 | orf2 | lp73 | gene | Pfam | protein matc | 38 | 205 | 2.3E-59 | + | . | Target=null 38 205;ID=match$1395_38_205;signature_desc=Lipoprotein;Name=PF01441;status=T;Dbxref="InterPro:IPR001800" |
| gene-bhYOR 001354 | orf2 | lp73 | gene | Pfam | protein matc | 1 | 116 | 8.4E-37 | + | . | Target=null 1 116;ID=match$817_1_116;signature_desc=Lipoprotein;Name=PF01441;status=T;Dbxref="InterPro:IPR001800" |
| gene-bhYOR 001361 | orf2 | lp73 | gene | Pfam | protein matc | 38 | 211 | 1.1E-43 | + | . | Target=null 38 211;ID=match$697_38_211;signature_desc=Lipoprotein;Name=PF01441;status=T;Dbxref="InterPro:IPR001800" |
| gene-bhYOR 001363 | orf2 | lp73 | pseudo | Pfam | protein matc | 3 | 171 | 7.2E-48 | + | . | Target=null 3 171;ID=match$2664_3_171;signature_desc=Lipoprotein;Name=PF01441;status=T;Dbxref="InterPro:IPR001800" |
| gene-bhYOR 001374 | orf2 | lp73 | pseudo | Pfam | protein matc | 2 | 139 | 3.8E-51 | + | . | Target=null 2 139;ID=match$1956_2_139;signature_desc=Lipoprotein;Name=PF01441;status=T;Dbxref="InterPro:IPR001800" |
| gene-bhYOR 001376 | orf2 | lp73 | pseudo | Pfam | protein matc | 2 | 137 | 3.1E-49 | + | . | Target=null 2 137;ID=match$1698_2_137;signature_desc=Lipoprotein;Name=PF01441;status=T;Dbxref="InterPro:IPR001800" |
| gene-bhYOR 001397 | orf2 | lp89 | gene | Pfam | protein matc | 29 | 198 | 6.9E-61 | + | . | Target=null 29 198;ID=match$1970_29_198;signature_desc=Lipoprotein;Name=PF01441;status=T;Dbxref="InterPro:IPR001800" |
| gene-bhYOR 001399 | orf2 | lp89 | gene | Pfam | protein matc | 38 | 205 | 3.1E-62 | + | . | Target=null 38 205;ID=match$3043_38_205;signature_desc=Lipoprotein;Name=PF01441;status=T;Dbxref="InterPro:IPR001800" |
| gene-bhYOR 001403 | orf2 | lp89 | gene | Pfam | protein matc | 25 | 195 | 3.8E-45 | + | . | Target=null 25 195;ID=match$1596_25_195;signature_desc=Lipoprotein;Name=PF01441;status=T;Dbxref="InterPro:IPR001800" |
| gene-bhYOR 001407 | orf2 | lp89 | pseudo | Pfam | protein matc | 38 | 71 | 6.2E-11 | + | . | Target=null 38 71;ID=match$288_38_71;signature_desc=Lipoprotein;Name=PF01441;status=T;Dbxref="InterPro:IPR001800" |
| gene-bhYOR 001407 | orf2 | lp89 | pseudo | Pfam | protein matc | 8 | 99 | 5.9E-17 | + | . | Target=null 8 99;ID=match$1168_8_99;signature_desc=Lipoprotein;Name=PF01441;status=T;Dbxref="InterPro:IPR001800" |
| gene-bhYOR 001409 | orf2 | lp89 | pseudo | Pfam | protein matc | 38 | 82 | 1.4E-17 | + | . | Target=null 38 82;ID=match$1056_38_82;signature_desc=Lipoprotein;Name=PF01441;status=T;Dbxref="InterPro:IPR001800" |
| gene-bhYOR 001409 | orf2 | lp89 | pseudo | Pfam | protein matc | 1 | 81 | 1.7E-24 | + | . | Target=null 1 81;ID=match$2048_1_81;signature_desc=Lipoprotein;Name=PF01441;status=T;Dbxref="InterPro:IPR001800" |
| gene-bhYOR 001409 | orf2 | lp89 | pseudo | Pfam | protein matc | 18 | 65 | 1.1E-12 | + | . | Target=null 18 65;ID=match$449_18_65;signature_desc=Lipoprotein;Name=PF01441;status=T;Dbxref="InterPro:IPR001800" |
| gene-bhYOR 001418 | orf2 | lp89 | pseudo | Pfam | protein matc | 3 | 74 | 3.4E-28 | + | . | Target=null 3 74;ID=match$40_3_74;signature_desc=Lipoprotein;Name=PF01441;status=T;Dbxref="InterPro:IPR001800" |
| gene-bhYOR 001418 | orf2 | lp89 | pseudo | Pfam | protein matc | 7 | 44 | 9.4E-07 | + | . | Target=null 7 44;ID=match$43_7_44;signature_desc=Lipoprotein;Name=PF01441;status=T;Dbxref="InterPro:IPR001800" |
| gene-bhYOR 001445 | orf2 | lp89 | gene | Pfam | protein matc | 38 | 205 | 6.4E-68 | + | . | Target=null 38 205;ID=match$1670_38_205;signature_desc=Lipoprotein;Name=PF01441;status=T;Dbxref="InterPro:IPR001800" |
| gene-bhYOR 001447 | orf2 | lp89 | pseudo | Pfam | protein matc | 2 | 34 | 7.1E-11 | + | . | Target=null 2 34;ID=match$1104_2_34;signature_desc=Lipoprotein;Name=PF01441;status=T;Dbxref="InterPro:IPR001800" |
| gene-bhYOR 001449 | orf2 | lp89 | gene | Pfam | protein matc | 38 | 205 | 7.1E-61 | + | . | Target=null 38 205;ID=match$3003_38_205;signature_desc=Lipoprotein;Name=PF01441;status=T;Dbxref="InterPro:IPR001800" |
| gene-bhYOR 001451 | orf2 | lp89 | gene | Pfam | protein matc | 38 | 199 | 2.5E-40 | + | . | Target=null 38 199;ID=match$3187_38_199;signature_desc=Lipoprotein;Name=PF01441;status=T;Dbxref="InterPro:IPR001800" |
| gene-bhYOR 001454 | orf2 | lp89 | pseudo | Pfam | protein matc | 38 | 138 | 6.5E-36 | + | . | Target=null 38 138;ID=match$2869_38_138;signature_desc=Lipoprotein;Name=PF01441;status=T;Dbxref="InterPro:IPR001800" |
| gene-bhYOR 001454 | orf2 | lp89 | pseudo | Pfam | protein matc | 6 | 77 | 5.1E-18 | + | . | Target=null 6 77;ID=match$3226_6_77;signature_desc=Lipoprotein;Name=PF01441;status=T;Dbxref="InterPro:IPR001800" |
| gene-bhYOR 001469 | orf2 | lp89 | gene | Pfam | protein matc | 38 | 207 | 2.2E-61 | + | . | Target=null 38 207;ID=match$695_38_207;signature_desc=Lipoprotein;Name=PF01441;status=T;Dbxref="InterPro:IPR001800" |
| gene-bhYOR 001476 | orf2 | lp89 | pseudo | Pfam | protein matc | 29 | 169 | 1.1E-52 | + | . | Target=null 29 169;ID=match$2781_29_169;signature_desc=Lipoprotein;Name=PF01441;status=T;Dbxref="InterPro:IPR001800" |
| gene-bhYOR 001478 | orf2 | lp89 | pseudo | Pfam | protein matc | 16 | 43 | 8E-08 | + | . | Target=null 16 43;ID=match$220_16_43;signature_desc=Lipoprotein;Name=PF01441;status=T;Dbxref="InterPro:IPR001800" |
| gene-bhYOR 001479 | orf2 | lp89 | gene | Pfam | protein matc | 1 | 115 | 4.5E-24 | + | . | Target=null 1 115;ID=match$2336_1_115;signature_desc=Lipoprotein;Name=PF01441;status=T;Dbxref="InterPro:IPR001800" |
| gene-bhYOR 001481 | orf2 | lp89 | gene | Pfam | protein matc | 29 | 200 | 4E-58 | + | . | Target=null 29 200;ID=match$2694_29_200;signature_desc=Lipoprotein;Name=PF01441;status=T;Dbxref="InterPro:IPR001800" |
| gene-bhYOR 001488 | orf2 | lp93 | gene | Pfam | protein matc | 38 | 206 | 1.4E-59 | + | . | Target=null 38 206;ID=match$286_38_206;signature_desc=Lipoprotein;Name=PF01441;status=T;Dbxref="InterPro:IPR001800" |
| gene-bhYOR 001499 | orf2 | lp93 | pseudo | Pfam | protein matc | 2 | 43 | 1.8E-17 | + | . | Target=null 2 43;ID=match$413_2_43;signature_desc=Lipoprotein;Name=PF01441;status=T;Dbxref="InterPro:IPR001800" |
| gene-bhYOR 001499 | orf2 | lp93 | pseudo | Pfam | protein matc | 13 | 144 | 6.9E-33 | + | . | Target=null 13 144;ID=match$409_13_144;signature_desc=Lipoprotein;Name=PF01441;status=T;Dbxref="InterPro:IPR001800" |
| gene-bhYOR 001500 | orf2 | lp93 | gene | Pfam | protein matc | 38 | 207 | 5.1E-64 | + | . | Target=null 38 207;ID=match$2929_38_207;signature_desc=Lipoprotein;Name=PF01441;status=T;Dbxref="InterPro:IPR001800" |
| gene-bhYOR 001601 | orf3 | lp93 | gene | Pfam | protein matc | 39 | 207 | 4.5E-60 | + | . | Target=null 39 207;ID=match$701_39_207;signature_desc=Lipoprotein;Name=PF01441;status=T;Dbxref="InterPro:IPR001800" |
| gene-bhYOR 001605 | orf3 | lp93 | gene | Pfam | protein matc | 28 | 194 | 3.5E-63 | + | . | Target=null 28 194;ID=match$2360_28_194;signature_desc=Lipoprotein;Name=PF01441;status=T;Dbxref="InterPro:IPR001800" |

| Vlp | | | | Subfamily | | | | | | | | |
|---|---|---|---|---|---|---|---|---|---|---|---|---|
| gene-bhYOR 001251 | orf2 | lp23 | pseudo | gamma | Pfam | protein matc | 19 | 126 | 4.5E-37 | + | . | Target=null 19 126;ID=match$737_19_126;signature_desc=Borrelia lipoprotein;Name=PF00921;status=T;Dbxref="InterPro:IPR000680" |
| gene-bhYOR 001251 | orf2 | lp23 | pseudo | | Pfam | protein matc | 9 | 38 | 0.000046 | + | . | Target=null 9 38;ID=match$279_9_38;signature_desc=Borrelia lipoprotein;Name=PF00921;status=T;Dbxref="InterPro:IPR000680" |
| gene-bhYOR 001251 | orf2 | lp23 | pseudo | | Pfam | protein matc | 9 | 76 | 6.9E-10 | + | . | Target=null 9 76;ID=match$2190_9_76;signature_desc=Borrelia lipoprotein;Name=PF00921;status=T;Dbxref="InterPro:IPR000680" |
| gene-bhYOR 001251 | orf2 | lp23 | pseudo | | Pfam | protein matc | 13 | 64 | 2E-19 | + | . | Target=null 13 64;ID=match$24_13_64;signature_desc=Borrelia lipoprotein;Name=PF00921;status=T;Dbxref="InterPro:IPR000680" |
| gene-bhYOR 001252 | orf2 | lp23 | pseudo | gamma | Pfam | protein matc | 1 | 118 | 6E-30 | + | . | Target=null 1 118;ID=match$1257_1_118;signature_desc=Borrelia lipoprotein;Name=PF00921;status=T;Dbxref="InterPro:IPR000680" |
| gene-bhYOR 001253 | orf2 | lp23 | pseudo | delta | Pfam | protein matc | 21 | 284 | 3.2E-80 | + | . | Target=null 21 284;ID=match$314_21_284;signature_desc=Borrelia lipoprotein;Name=PF00921;status=T;Dbxref="InterPro:IPR000680" |
| gene-bhYOR 001255 | orf2 | lp23 | gene | delta | Pfam | protein matc | 1 | 276 | 1.3E-81 | + | . | Target=null 1 276;ID=match$719_1_276;signature_desc=Borrelia lipoprotein;Name=PF00921;status=T;Dbxref="InterPro:IPR000680" |
| gene-bhYOR 001257 | orf2 | lp23 | pseudo | gamma | Pfam | protein matc | 19 | 126 | 4.5E-37 | + | . | Target=null 19 126;ID=match$735_19_126;signature_desc=Borrelia lipoprotein;Name=PF00921;status=T;Dbxref="InterPro:IPR000680" |
| gene-bhYOR 001257 | orf2 | lp23 | pseudo | | Pfam | protein matc | 9 | 38 | 0.000046 | + | . | Target=null 9 38;ID=match$280_9_38;signature_desc=Borrelia lipoprotein;Name=PF00921;status=T;Dbxref="InterPro:IPR000680" |
| gene-bhYOR 001257 | orf2 | lp23 | pseudo | | Pfam | protein matc | 9 | 76 | 6.9E-10 | + | . | Target=null 9 76;ID=match$2188_9_76;signature_desc=Borrelia lipoprotein;Name=PF00921;status=T;Dbxref="InterPro:IPR000680" |
| gene-bhYOR 001257 | orf2 | lp23 | pseudo | | Pfam | protein matc | 13 | 82 | 4.5E-30 | + | . | Target=null 13 82;ID=match$565_13_82;signature_desc=Borrelia lipoprotein;Name=PF00921;status=T;Dbxref="InterPro:IPR000680" |
| gene-bhYOR 001258 | orf2 | lp23 | pseudo | gamma | Pfam | protein matc | 1 | 61 | 5.2E-10 | + | . | Target=null 1 61;ID=match$2186_1_61;signature_desc=Borrelia lipoprotein;Name=PF00921;status=T;Dbxref="InterPro:IPR000680" |
| gene-bhYOR 001260 | orf2 | lp46 | gene | delta | Pfam | protein matc | 34 | 313 | 6E-85 | + | . | Target=null 34 313;ID=match$1068_34_313;signature_desc=Borrelia lipoprotein;Name=PF00921;status=T;Dbxref="InterPro:IPR000680" |
| gene-bhYOR 001261 | orf2 | lp46 | pseudo | beta | Pfam | protein matc | 10 | 79 | 1.1E-19 | + | . | Target=null 10 79;ID=match$644_10_79;signature_desc=Borrelia lipoprotein;Name=PF00921;status=T;Dbxref="InterPro:IPR000680" |
| gene-bhYOR 001262 | orf2 | lp46 | gene | gamma | Pfam | protein matc | 30 | 314 | 1.2E-85 | + | . | Target=null 30 314;ID=match$363_30_314;signature_desc=Borrelia lipoprotein;Name=PF00921;status=T;Dbxref="InterPro:IPR000680" |
| gene-bhYOR 001263 | orf2 | lp46 | gene | alpha | Pfam | protein matc | 7 | 302 | 8.7E-72 | + | . | Target=null 7 302;ID=match$2223_7_302;signature_desc=Borrelia lipoprotein;Name=PF00921;status=T;Dbxref="InterPro:IPR000680" |
| gene-bhYOR 001264 | orf2 | lp46 | pseudo | beta | Pfam | protein matc | 20 | 120 | 1E-13 | + | . | Target=null 20 120;ID=match$338_20_120;signature_desc=Borrelia lipoprotein;Name=PF00921;status=T;Dbxref="InterPro:IPR000680" |
| gene-bhYOR 001264 | orf2 | lp46 | pseudo | | Pfam | protein matc | 25 | 204 | 1.6E-30 | + | . | Target=null 25 204;ID=match$1955_25_204;signature_desc=Borrelia lipoprotein;Name=PF00921;status=T;Dbxref="InterPro:IPR000680" |
| gene-bhYOR 001267 | orf2 | lp46 | pseudo | delta | Pfam | protein matc | 43 | 320 | 1.6E-83 | + | . | Target=null 43 320;ID=match$727_43_320;signature_desc=Borrelia lipoprotein;Name=PF00921;status=T;Dbxref="InterPro:IPR000680" |
| gene-bhYOR 001269 | orf2 | lp46 | pseudo | gamma | Pfam | protein matc | 43 | 87 | 1.3E-11 | + | . | Target=null 43 87;ID=match$2898_43_87;signature_desc=Borrelia lipoprotein;Name=PF00921;status=T;Dbxref="InterPro:IPR000680" |
| gene-bhYOR 001269 | orf2 | lp46 | pseudo | | Pfam | protein matc | 3 | 237 | 1.2E-61 | + | . | Target=null 3 237;ID=match$2662_3_237;signature_desc=Borrelia lipoprotein;Name=PF00921;status=T;Dbxref="InterPro:IPR000680" |
| gene-bhYOR 001303 | orf2 | lp73 | gene | alpha | Pfam | protein matc | 58 | 343 | 5.5E-62 | + | . | Target=null 58 343;ID=match$1691_58_343;signature_desc=Borrelia lipoprotein;Name=PF00921;status=T;Dbxref="InterPro:IPR000680" |
| gene-bhYOR 001305 | orf2 | lp73 | pseudo | beta | Pfam | protein matc | 2 | 190 | 6.8E-15 | + | . | Target=null 2 190;ID=match$1696_2_190;signature_desc=Borrelia lipoprotein;Name=PF00921;status=T;Dbxref="InterPro:IPR000680" |
| gene-bhYOR 001306 | orf2 | lp73 | gene | alpha | Pfam | protein matc | 23 | 132 | 3.5E-28 | + | . | Target=null 23 132;ID=match$595_23_132;signature_desc=Borrelia lipoprotein;Name=PF00921;status=T;Dbxref="InterPro:IPR000680" |
| gene-bhYOR 001307 | orf2 | lp73 | pseudo | gamma | Pfam | protein matc | 3 | 71 | 4.4E-23 | + | . | Target=null 3 71;ID=match$435_3_71;signature_desc=Borrelia lipoprotein;Name=PF00921;status=T;Dbxref="InterPro:IPR000680" |
| gene-bhYOR 001314 | orf2 | lp73 | pseudo | gamma | Pfam | protein matc | 1 | 285 | 6.2E-85 | + | . | Target=null 1 285;ID=match$1311_1_285;signature_desc=Borrelia lipoprotein;Name=PF00921;status=T;Dbxref="InterPro:IPR000680" |
| gene-bhYOR 001317 | orf2 | lp73 | gene | gamma | Pfam | protein matc | 34 | 319 | 1.6E-92 | + | . | Target=null 34 319;ID=match$1314_34_319;signature_desc=Borrelia lipoprotein;Name=PF00921;status=T;Dbxref="InterPro:IPR000680" |
| gene-bhYOR 001318 | orf2 | lp73 | gene | alpha | Pfam | protein matc | 9 | 130 | 2.6E-23 | + | . | Target=null 9 130;ID=match$831_9_130;signature_desc=Borrelia lipoprotein;Name=PF00921;status=T;Dbxref="InterPro:IPR000680" |
| gene-bhYOR 001320 | orf2 | lp73 | gene | beta | Pfam | protein matc | 47 | 334 | 7.5E-49 | + | . | Target=null 47 334;ID=match$38_47_334;signature_desc=Borrelia lipoprotein;Name=PF00921;status=T;Dbxref="InterPro:IPR000680" |
| gene-bhYOR 001321 | orf2 | lp73 | gene | gamma | Pfam | protein matc | 28 | 63 | 2E-10 | + | . | Target=null 28 63;ID=match$132_28_63;signature_desc=Borrelia lipoprotein;Name=PF00921;status=T;Dbxref="InterPro:IPR000680" |
| gene-bhYOR 001328 | orf2 | lp73 | pseudo | gamma | Pfam | protein matc | 3 | 151 | 3.7E-36 | + | . | Target=null 3 151;ID=match$730_3_151;signature_desc=Borrelia lipoprotein;Name=PF00921;status=T;Dbxref="InterPro:IPR000680" |
| gene-bhYOR 001328 | orf2 | lp73 | pseudo | | Pfam | protein matc | 4 | 91 | 6E-15 | + | . | Target=null 4 91;ID=match$772_4_91;signature_desc=Borrelia lipoprotein;Name=PF00921;status=T;Dbxref="InterPro:IPR000680" |
| gene-bhYOR 001338 | orf2 | lp73 | pseudo | delta | Pfam | protein matc | 4 | 287 | 6.1E-89 | + | . | Target=null 4 287;ID=match$1873_4_287;signature_desc=Borrelia lipoprotein;Name=PF00921;status=T;Dbxref="InterPro:IPR000680" |
| gene-bhYOR 001339 | orf2 | lp73 | pseudo | gamma | Pfam | protein matc | 2 | 89 | 1.1E-27 | + | . | Target=null 2 89;ID=match$269_2_89;signature_desc=Borrelia lipoprotein;Name=PF00921;status=T;Dbxref="InterPro:IPR000680" |
| gene-bhYOR 001352 | orf2 | lp73 | pseudo | gamma | Pfam | protein matc | 14 | 60 | 6.2E-13 | + | . | Target=null 14 60;ID=match$2321_14_60;signature_desc=Borrelia lipoprotein;Name=PF00921;status=T;Dbxref="InterPro:IPR000680" |
| gene-bhYOR 001353 | orf2 | lp73 | pseudo | gamma | Pfam | protein matc | 6 | 38 | 6.4E-13 | + | . | Target=null 6 38;ID=match$3162_6_38;signature_desc=Borrelia lipoprotein;Name=PF00921;status=T;Dbxref="InterPro:IPR000680" |
| gene-bhYOR 001355 | orf2 | lp73 | gene | alpha | Pfam | protein matc | 59 | 353 | 3.9E-72 | + | . | Target=null 59 353;ID=match$164_59_353;signature_desc=Borrelia lipoprotein;Name=PF00921;status=T;Dbxref="InterPro:IPR000680" |
| gene-bhYOR 001356 | orf2 | lp73 | gene | gamma | Pfam | protein matc | 1 | 236 | 1.9E-65 | + | . | Target=null 1 236;ID=match$1928_1_236;signature_desc=Borrelia lipoprotein;Name=PF00921;status=T;Dbxref="InterPro:IPR000680" |
| gene-bhYOR 001362 | orf2 | lp73 | pseudo | gamma | Pfam | protein matc | 3 | 116 | 1.7E-16 | + | . | Target=null 3 116;ID=match$757_3_116;signature_desc=Borrelia lipoprotein;Name=PF00921;status=T;Dbxref="InterPro:IPR000680" |
| gene-bhYOR 001362 | orf2 | lp73 | pseudo | | Pfam | protein matc | 6 | 111 | 3.3E-35 | + | . | Target=null 6 111;ID=match$3169_6_111;signature_desc=Borrelia lipoprotein;Name=PF00921;status=T;Dbxref="InterPro:IPR000680" |
| gene-bhYOR 001371 | orf2 | lp73 | pseudo | delta | Pfam | protein matc | 10 | 107 | 1.4E-27 | + | . | Target=null 10 107;ID=match$2111_10_107;signature_desc=Borrelia lipoprotein;Name=PF00921;status=T;Dbxref="InterPro:IPR000680" |
| gene-bhYOR 001372 | orf2 | lp73 | gene | alpha | Pfam | protein matc | 4 | 78 | 7.8E-24 | + | . | Target=null 4 78;ID=match$2080_4_78;signature_desc=Borrelia lipoprotein;Name=PF00921;status=T;Dbxref="InterPro:IPR000680" |
| gene-bhYOR 001373 | orf2 | lp73 | gene | beta | Pfam | protein matc | 21 | 315 | 1.2E-45 | + | . | Target=null 21 315;ID=match$1138_21_315;signature_desc=Borrelia lipoprotein;Name=PF00921;status=T;Dbxref="InterPro:IPR000680" |
| gene-bhYOR 001375 | orf2 | lp73 | pseudo | gamma | Pfam | protein matc | 1 | 209 | 4.7E-49 | + | . | Target=null 1 209;ID=match$2100_1_209;signature_desc=Borrelia lipoprotein;Name=PF00921;status=T;Dbxref="InterPro:IPR000680" |
| gene-bhYOR 001375 | orf2 | lp73 | pseudo | | Pfam | protein matc | 16 | 57 | 5.2E-14 | + | . | Target=null 16 57;ID=match$1180_16_57;signature_desc=Borrelia lipoprotein;Name=PF00921;status=T;Dbxref="InterPro:IPR000680" |
| gene-bhYOR 001377 | orf2 | lp73 | pseudo | delta | Pfam | protein matc | 3 | 84 | 9.2E-13 | + | . | Target=null 3 84;ID=match$972_3_84;signature_desc=Borrelia lipoprotein;Name=PF00921;status=T;Dbxref="InterPro:IPR000680" |
| gene-bhYOR 001377 | orf2 | lp73 | pseudo | | Pfam | protein matc | 2 | 182 | 8.2E-54 | + | . | Target=null 2 182;ID=match$765_2_182;signature_desc=Borrelia lipoprotein;Name=PF00921;status=T;Dbxref="InterPro:IPR000680" |
| gene-bhYOR 001378 | orf2 | lp73 | pseudo | gamma | Pfam | protein matc | 15 | 48 | 2.8E-07 | + | . | Target=null 15 48;ID=match$2948_15_48;signature_desc=Borrelia lipoprotein;Name=PF00921;status=T;Dbxref="InterPro:IPR000680" |
| gene-bhYOR 001379 | orf2 | lp73 | pseudo | beta | Pfam | protein matc | 13 | 97 | 0.0000034 | + | . | Target=null 13 97;ID=match$1784_13_97;signature_desc=Borrelia lipoprotein;Name=PF00921;status=T;Dbxref="InterPro:IPR000680" |
| gene-bhYOR 001390 | orf2 | lp89 | gene | gamma | Pfam | protein matc | 43 | 319 | 1.5E-86 | + | . | Target=null 43 319;ID=match$35_43_319;signature_desc=Borrelia lipoprotein;Name=PF00921;status=T;Dbxref="InterPro:IPR000680" |
| gene-bhYOR 001391 | orf2 | lp89 | gene | beta | Pfam | protein matc | 47 | 331 | 5.7E-45 | + | . | Target=null 47 331;ID=match$1424_47_331;signature_desc=Borrelia lipoprotein;Name=PF00921;status=T;Dbxref="InterPro:IPR000680" |
| gene-bhYOR 001392 | orf2 | lp89 | gene | alpha | Pfam | protein matc | 61 | 353 | 2.7E-69 | + | . | Target=null 61 353;ID=match$1600_61_353;signature_desc=Borrelia lipoprotein;Name=PF00921;status=T;Dbxref="InterPro:IPR000680" |
| gene-bhYOR 001393 | orf2 | lp89 | gene | beta | Pfam | protein matc | 20 | 307 | 5.4E-49 | + | . | Target=null 20 307;ID=match$153_20_307;signature_desc=Borrelia lipoprotein;Name=PF00921;status=T;Dbxref="InterPro:IPR000680" |
| gene-bhYOR 001394 | orf2 | lp89 | pseudo | gamma | Pfam | protein matc | 1 | 57 | 2.2E-08 | + | . | Target=null 1 57;ID=match$60_1_57;signature_desc=Borrelia lipoprotein;Name=PF00921;status=T;Dbxref="InterPro:IPR000680" |
| gene-bhYOR 001394 | orf2 | lp89 | pseudo | | Pfam | protein matc | 31 | 65 | 0.000014 | + | . | Target=null 31 65;ID=match$3185_31_65;signature_desc=Borrelia lipoprotein;Name=PF00921;status=T;Dbxref="InterPro:IPR000680" |
| gene-bhYOR 001394 | orf2 | lp89 | pseudo | | Pfam | protein matc | 10 | 43 | 0.000012 | + | . | Target=null 10 43;ID=match$37_10_43;signature_desc=Borrelia lipoprotein;Name=PF00921;status=T;Dbxref="InterPro:IPR000680" |
| gene-bhYOR 001394 | orf2 | lp89 | pseudo | | Pfam | protein matc | 10 | 85 | 9.7E-26 | + | . | Target=null 10 85;ID=match$1881_10_85;signature_desc=Borrelia lipoprotein;Name=PF00921;status=T;Dbxref="InterPro:IPR000680" |
| gene-bhYOR 001396 | orf2 | lp89 | gene | alpha | Pfam | protein matc | 58 | 355 | 1E-68 | + | . | Target=null 58 355;ID=match$240_58_355;signature_desc=Borrelia lipoprotein;Name=PF00921;status=T;Dbxref="InterPro:IPR000680" |
| gene-bhYOR 001398 | orf2 | lp89 | pseudo | gamma | Pfam | protein matc | 43 | 91 | 2.3E-14 | + | . | Target=null 43 91;ID=match$465_43_91;signature_desc=Borrelia lipoprotein;Name=PF00921;status=T;Dbxref="InterPro:IPR000680" |
| gene-bhYOR 001398 | orf2 | lp89 | pseudo | | Pfam | protein matc | 7 | 237 | 1.9E-65 | + | . | Target=null 7 237;ID=match$2167_7_237;signature_desc=Borrelia lipoprotein;Name=PF00921;status=T;Dbxref="InterPro:IPR000680" |
| gene-bhYOR 001404 | orf2 | lp89 | pseudo | delta | Pfam | protein matc | 14 | 305 | 1.3E-85 | + | . | Target=null 14 305;ID=match$513_14_305;signature_desc=Borrelia lipoprotein;Name=PF00921;status=T;Dbxref="InterPro:IPR000680" |
| gene-bhYOR 001405 | orf2 | lp89 | pseudo | beta | Pfam | protein matc | 20 | 162 | 1.3E-17 | + | . | Target=null 20 162;ID=match$1364_20_162;signature_desc=Borrelia lipoprotein;Name=PF00921;status=T;Dbxref="InterPro:IPR000680" |
| gene-bhYOR 001405 | orf2 | lp89 | pseudo | | Pfam | protein matc | 78 | 121 | 0.0000041 | + | . | Target=null 78 121;ID=match$2482_78_121;signature_desc=Borrelia lipoprotein;Name=PF00921;status=T;Dbxref="InterPro:IPR000680" |
| gene-bhYOR 001406 | orf2 | lp89 | gene | alpha | Pfam | protein matc | 7 | 304 | 5.2E-71 | + | . | Target=null 7 304;ID=match$2031_7_304;signature_desc=Borrelia lipoprotein;Name=PF00921;status=T;Dbxref="InterPro:IPR000680" |
| gene-bhYOR 001408 | orf2 | lp89 | pseudo | delta | Pfam | protein matc | 42 | 315 | 1.7E-83 | + | . | Target=null 42 315;ID=match$1113_42_315;signature_desc=Borrelia lipoprotein;Name=PF00921;status=T;Dbxref="InterPro:IPR000680" |
| gene-bhYOR 001411 | orf2 | lp89 | gene | beta | Pfam | protein matc | 21 | 310 | 1.6E-45 | + | . | Target=null 21 310;ID=match$2795_21_310;signature_desc=Borrelia lipoprotein;Name=PF00921;status=T;Dbxref="InterPro:IPR000680" |
| gene-bhYOR 001412 | orf2 | lp89 | pseudo | alpha | Pfam | protein matc | 58 | 358 | 4E-63 | + | . | Target=null 58 358;ID=match$2836_58_358;signature_desc=Borrelia lipoprotein;Name=PF00921;status=T;Dbxref="InterPro:IPR000680" |
| gene-bhYOR 001413 | orf2 | lp89 | gene | delta | Pfam | protein matc | 43 | 326 | 1.6E-83 | + | . | Target=null 43 326;ID=match$2732_43_326;signature_desc=Borrelia lipoprotein;Name=PF00921;status=T;Dbxref="InterPro:IPR000680" |
| gene-bhYOR 001414 | orf2 | lp89 | gene | beta | Pfam | protein matc | 55 | 345 | 2E-41 | + | . | Target=null 55 345;ID=match$2050_55_345;signature_desc=Borrelia lipoprotein;Name=PF00921;status=T;Dbxref="InterPro:IPR000680" |
| gene-bhYOR 001415 | orf2 | lp89 | pseudo | delta | Pfam | protein matc | 26 | 301 | 5.6E-83 | + | . | Target=null 26 301;ID=match$2525_26_301;signature_desc=Borrelia lipoprotein;Name=PF00921;status=T;Dbxref="InterPro:IPR000680" |
| gene-bhYOR 001423 | orf2 | lp89 | pseudo | gamma | Pfam | protein matc | 42 | 73 | 1.1E-07 | + | . | Target=null 42 73;ID=match$3215_42_73;signature_desc=Borrelia lipoprotein;Name=PF00921;status=T;Dbxref="InterPro:IPR000680" |
| gene-bhYOR 001423 | orf2 | lp89 | pseudo | | Pfam | protein matc | 5 | 121 | 3.2E-24 | + | . | Target=null 5 121;ID=match$1778_5_121;signature_desc=Borrelia lipoprotein;Name=PF00921;status=T;Dbxref="InterPro:IPR000680" |
| gene-bhYOR 001424 | orf2 | lp89 | gene | beta | Pfam | protein matc | 20 | 307 | 2.4E-44 | + | . | Target=null 20 307;ID=match$2008_20_307;signature_desc=Borrelia lipoprotein;Name=PF00921;status=T;Dbxref="InterPro:IPR000680" |
| gene-bhYOR 001425 | orf2 | lp89 | gene | alpha | Pfam | protein matc | 60 | 355 | 1.2E-66 | + | . | Target=null 60 355;ID=match$2407_60_355;signature_desc=Borrelia lipoprotein;Name=PF00921;status=T;Dbxref="InterPro:IPR000680" |
| gene-bhYOR 001429 | orf2 | lp89 | pseudo | gamma | Pfam | protein matc | 4 | 35 | 2.3E-10 | + | . | Target=null 4 35;ID=match$663_4_35;signature_desc=Borrelia lipoprotein;Name=PF00921;status=T;Dbxref="InterPro:IPR000680" |
| gene-bhYOR 001430 | orf2 | lp89 | gene | delta | Pfam | protein matc | 43 | 323 | 5.3E-89 | + | . | Target=null 43 323;ID=match$1266_43_323;signature_desc=Borrelia lipoprotein;Name=PF00921;status=T;Dbxref="InterPro:IPR000680" |
| gene-bhYOR 001431 | orf2 | lp89 | pseudo | gamma | Pfam | protein matc | 5 | 31 | 0.000025 | + | . | Target=null 5 31;ID=match$1829_5_31;signature_desc=Borrelia lipoprotein;Name=PF00921;status=T;Dbxref="InterPro:IPR000680" |
| gene-bhYOR 001439 | orf2 | lp89 | pseudo | delta | Pfam | protein matc | 3 | 39 | 1.7E-12 | + | . | Target=null 3 39;ID=match$1587_3_39;signature_desc=Borrelia lipoprotein;Name=PF00921;status=T;Dbxref="InterPro:IPR000680" |
| gene-bhYOR 001440 | orf2 | lp89 | pseudo | beta | Pfam | protein matc | 1 | 48 | 8.7E-12 | + | . | Target=null 1 48;ID=match$108_1_48;signature_desc=Borrelia lipoprotein;Name=PF00921;status=T;Dbxref="InterPro:IPR000680" |
| gene-bhYOR 001441 | orf2 | lp89 | gene | delta | Pfam | protein matc | 11 | 55 | 3.2E-08 | + | . | Target=null 11 55;ID=match$452_11_55;signature_desc=Borrelia lipoprotein;Name=PF00921;status=T;Dbxref="InterPro:IPR000680" |
| gene-bhYOR 001443 | orf2 | lp89 | pseudo | gamma | Pfam | protein matc | 4 | 64 | 2.5E-11 | + | . | Target=null 4 64;ID=match$2522_4_64;signature_desc=Borrelia lipoprotein;Name=PF00921;status=T;Dbxref="InterPro:IPR000680" |
| gene-bhYOR 001444 | orf2 | lp89 | gene | gamma | Pfam | protein matc | 2 | 94 | 1.5E-17 | + | . | Target=null 2 94;ID=match$2681_2_94;signature_desc=Borrelia lipoprotein;Name=PF00921;status=T;Dbxref="InterPro:IPR000680" |
| gene-bhYOR 001450 | orf2 | lp89 | pseudo | gamma | Pfam | protein matc | 34 | 56 | 0.000011 | + | . | Target=null 34 56;ID=match$1903_34_56;signature_desc=Borrelia lipoprotein;Name=PF00921;status=T;Dbxref="InterPro:IPR000680" |
| gene-bhYOR 001450 | orf2 | lp89 | pseudo | | Pfam | protein matc | 1 | 235 | 2.9E-60 | + | . | Target=null 1 235;ID=match$1614_1_235;signature_desc=Borrelia lipoprotein;Name=PF00921;status=T;Dbxref="InterPro:IPR000680" |
| gene-bhYOR 001452 | orf2 | lp89 | pseudo | gamma | Pfam | protein matc | 3 | 53 | 3.9E-18 | + | . | Target=null 3 53;ID=match$1001_3_53;signature_desc=Borrelia lipoprotein;Name=PF00921;status=T;Dbxref="InterPro:IPR000680" |
| gene-bhYOR 001453 | orf2 | lp89 | gene | gamma | Pfam | protein matc | 8 | 60 | 0.000022 | + | . | Target=null 8 60;ID=match$1286_8_60;signature_desc=Borrelia lipoprotein;Name=PF00921;status=T;Dbxref="InterPro:IPR000680" |
| gene-bhYOR 001455 | orf2 | lp89 | gene | gamma | Pfam | protein matc | 28 | 63 | 2.3E-10 | + | . | Target=null 28 63;ID=match$2890_28_63;signature_desc=Borrelia lipoprotein;Name=PF00921;status=T;Dbxref="InterPro:IPR000680" |
| gene-bhYOR 001464 | orf2 | lp89 | pseudo | delta | Pfam | protein matc | 1 | 171 | 1.7E-45 | + | . | Target=null 1 171;ID=match$1813_1_171;signature_desc=Borrelia lipoprotein;Name=PF00921;status=T;Dbxref="InterPro:IPR000680" |
| gene-bhYOR 001465 | orf2 | lp89 | pseudo | gamma | Pfam | protein matc | 6 | 38 | 6.4E-13 | + | . | Target=null 6 38;ID=match$1693_6_38;signature_desc=Borrelia lipoprotein;Name=PF00921;status=T;Dbxref="InterPro:IPR000680" |
| gene-bhYOR 001470 | orf2 | lp89 | gene | gamma | Pfam | protein matc | 4 | 96 | 1.1E-20 | + | . | Target=null 4 96;ID=match$3035_4_96;signature_desc=Borrelia lipoprotein;Name=PF00921;status=T;Dbxref="InterPro:IPR000680" |
| gene-bhYOR 001471 | orf2 | lp89 | pseudo | gamma | Pfam | protein matc | 1 | 29 | 4.8E-10 | + | . | Target=null 1 29;ID=match$1673_1_29;signature_desc=Borrelia lipoprotein;Name=PF00921;status=T;Dbxref="InterPro:IPR000680" |
| gene-bhYOR 001472 | orf2 | lp89 | pseudo | delta | Pfam | protein matc | 4 | 164 | 1.3E-38 | + | . | Target=null 4 164;ID=match$708_4_164;signature_desc=Borrelia lipoprotein;Name=PF00921;status=T;Dbxref="InterPro:IPR000680" |
| gene-bhYOR 001472 | orf2 | lp89 | pseudo | | Pfam | protein matc | 17 | 56 | 5.6E-12 | + | . | Target=null 17 56;ID=match$2400_17_56;signature_desc=Borrelia lipoprotein;Name=PF00921;status=T;Dbxref="InterPro:IPR000680" |
| gene-bhYOR 001473 | orf2 | lp89 | gene | alpha | Pfam | protein matc | 61 | 350 | 9.4E-71 | + | . | Target=null 61 350;ID=match$1959_61_350;signature_desc=Borrelia lipoprotein;Name=PF00921;status=T;Dbxref="InterPro:IPR000680" |
| gene-bhYOR 001474 | orf2 | lp89 | gene | beta | Pfam | protein matc | 20 | 308 | 3.7E-45 | + | . | Target=null 20 308;ID=match$1553_20_308;signature_desc=Borrelia lipoprotein;Name=PF00921;status=T;Dbxref="InterPro:IPR000680" |
| gene-bhYOR 001475 | orf2 | lp89 | gene | delta | Pfam | protein matc | 20 | 302 | 7.2E-85 | + | . | Target=null 20 302;ID=match$2601_20_302;signature_desc=Borrelia lipoprotein;Name=PF00921;status=T;Dbxref="InterPro:IPR000680" |
| gene-bhYOR 001477 | orf2 | lp89 | gene | delta | Pfam | protein matc | 30 | 315 | 3.1E-86 | + | . | Target=null 30 315;ID=match$1223_30_315;signature_desc=Borrelia lipoprotein;Name=PF00921;status=T;Dbxref="InterPro:IPR000680" |
| gene-bhYOR 001480 | orf2 | lp89 | pseudo | gamma | Pfam | protein matc | 7 | 30 | 0.0000039 | + | . | Target=null 7 30;ID=match$237_7_30;signature_desc=Borrelia lipoprotein;Name=PF00921;status=T;Dbxref="InterPro:IPR000680" |
| gene-bhYOR 001480 | orf2 | lp89 | pseudo | | Pfam | protein matc | 3 | 270 | 4.5E-74 | + | . | Target=null 3 270;ID=match$3126_3_270;signature_desc=Borrelia lipoprotein;Name=PF00921;status=T;Dbxref="InterPro:IPR000680" |
| gene-bhYOR 001482 | orf2 | lp89 | gene | alpha | Pfam | protein matc | 57 | 348 | 5.1E-66 | + | . | Target=null 57 348;ID=match$2449_57_348;signature_desc=Borrelia lipoprotein;Name=PF00921;status=T;Dbxref="InterPro:IPR000680" |
| gene-bhYOR 001483 | orf2 | lp89 | pseudo | beta | Pfam | protein matc | 34 | 77 | 0.0000086 | + | . | Target=null 34 77;ID=match$2953_34_77;signature_desc=Borrelia lipoprotein;Name=PF00921;status=T;Dbxref="InterPro:IPR000680" |
| gene-bhYOR 001483 | orf2 | lp89 | pseudo | | Pfam | protein matc | 3 | 168 | 2.3E-33 | + | . | Target=null 3 168;ID=match$3268_3_168;signature_desc=Borrelia lipoprotein;Name=PF00921;status=T;Dbxref="InterPro:IPR000680" |
| gene-bhYOR 001484 | orf2 | lp89 | pseudo | gamma | Pfam | protein matc | 3 | 37 | 0.0000087 | + | . | Target=null 3 37;ID=match$3069_3_37;signature_desc=Borrelia lipoprotein;Name=PF00921;status=T;Dbxref="InterPro:IPR000680" |
| gene-bhYOR 001485 | orf2 | lp89 | gene | beta | Pfam | protein matc | 22 | 240 | 2.3E-37 | + | . | Target=null 22 240;ID=match$2956_22_240;signature_desc=Borrelia lipoprotein;Name=PF00921;status=T;Dbxref="InterPro:IPR000680" |
| gene-bhYOR 001487 | orf2 | lp93 | pseudo | delta | Pfam | protein matc | 14 | 298 | 2.4E-87 | + | . | Target=null 14 298;ID=match$786_14_298;signature_desc=Borrelia lipoprotein;Name=PF00921;status=T;Dbxref="InterPro:IPR000680" |
| gene-bhYOR 001489 | orf2 | lp93 | pseudo | gamma | Pfam | protein matc | 2 | 37 | 2.4E-11 | + | . | Target=null 2 37;ID=match$2411_2_37;signature_desc=Borrelia lipoprotein;Name=PF00921;status=T;Dbxref="InterPro:IPR000680" |
| gene-bhYOR 001494 | orf2 | lp93 | gene | delta | Pfam | protein matc | 34 | 313 | 5.8E-85 | + | . | Target=null 34 313;ID=match$589_34_313;signature_desc=Borrelia lipoprotein;Name=PF00921;status=T;Dbxref="InterPro:IPR000680" |
| gene-bhYOR 001495 | orf2 | lp93 | pseudo | beta | Pfam | protein matc | 10 | 79 | 1.1E-19 | + | . | Target=null 10 79;ID=match$646_10_79;signature_desc=Borrelia lipoprotein;Name=PF00921;status=T;Dbxref="InterPro:IPR000680" |
| gene-bhYOR 001496 | orf2 | lp93 | gene | gamma | Pfam | protein matc | 30 | 314 | 1.2E-85 | + | . | Target=null 30 314;ID=match$365_30_314;signature_desc=Borrelia lipoprotein;Name=PF00921;status=T;Dbxref="InterPro:IPR000680" |
| gene-bhYOR 001497 | orf2 | lp93 | gene | alpha | Pfam | protein matc | 7 | 302 | 8.7E-72 | + | . | Target=null 7 302;ID=match$2221_7_302;signature_desc=Borrelia lipoprotein;Name=PF00921;status=T;Dbxref="InterPro:IPR000680" |
| gene-bhYOR 001498 | orf2 | lp93 | pseudo | beta | Pfam | protein matc | 20 | 120 | 1E-13 | + | . | Target=null 20 120;ID=match$336_20_120;signature_desc=Borrelia lipoprotein;Name=PF00921;status=T;Dbxref="InterPro:IPR000680" |
| gene-bhYOR 001498 | orf2 | lp93 | pseudo | | Pfam | protein matc | 25 | 204 | 1.6E-30 | + | . | Target=null 25 204;ID=match$1953_25_204;signature_desc=Borrelia lipoprotein;Name=PF00921;status=T;Dbxref="InterPro:IPR000680" |
| gene-bhYOR 001501 | orf2 | lp93 | pseudo | delta | Pfam | protein matc | 14 | 286 | 2.2E-85 | + | . | Target=null 14 286;ID=match$531_14_286;signature_desc=Borrelia lipoprotein;Name=PF00921;status=T;Dbxref="InterPro:IPR000680" |
| gene-bhYOR 001602 | orf3 | lp93 | gene | gamma | Pfam | protein matc | 1 | 275 | 8.6E-79 | + | . | Target=null 1 275;ID=match$3110_1_275;signature_desc=Borrelia lipoprotein;Name=PF00921;status=T;Dbxref="InterPro:IPR000680" |
| gene-bhYOR 001603 | orf3 | lp93 | gene | alpha | Pfam | protein matc | 7 | 302 | 8.7E-72 | + | . | Target=null 7 302;ID=match$2219_7_302;signature_desc=Borrelia lipoprotein;Name=PF00921;status=T;Dbxref="InterPro:IPR000680" |
| gene-bhYOR 001604 | orf3 | lp93 | pseudo | beta | Pfam | protein matc | 20 | 120 | 1E-13 | + | . | Target=null 20 120;ID=match$334_20_120;signature_desc=Borrelia lipoprotein;Name=PF00921;status=T;Dbxref="InterPro:IPR000680" |
| gene-bhYOR 001604 | orf3 | lp93 | pseudo | | Pfam | protein matc | 25 | 204 | 1.6E-30 | + | . | Target=null 25 204;ID=match$1823_25_204;signature_desc=Borrelia lipoprotein;Name=PF00921;status=T;Dbxref="InterPro:IPR000680" |
| gene-bhYOR 001606 | orf3 | lp93 | pseudo | delta | Pfam | protein matc | 8 | 99 | 1.1E-27 | + | . | Target=null 8 99;ID=match$3190_8_99;signature_desc=Borrelia lipoprotein;Name=PF00921;status=T;Dbxref="InterPro:IPR000680" |
| gene-bhYOR 001606 | orf3 | lp93 | pseudo | | Pfam | protein matc | 28 | 84 | 1.9E-07 | + | . | Target=null 28 84;ID=match$3032_28_84;signature_desc=Borrelia lipoprotein;Name=PF00921;status=T;Dbxref="InterPro:IPR000680" |
| gene-bhYOR 001606 | orf3 | lp93 | pseudo | | Pfam | protein matc | 2 | 82 | 6.8E-22 | + | . | Target=null 2 82;ID=match$238_2_82;signature_desc=Borrelia lipoprotein;Name=PF00921;status=T;Dbxref="InterPro:IPR000680" |
| gene-bhYOR 001606 | orf3 | lp93 | pseudo | | Pfam | protein matc | 2 | 41 | 2E-12 | + | . | Target=null 2 41;ID=match$2410_2_41;signature_desc=Borrelia lipoprotein;Name=PF00921;status=T;Dbxref="InterPro:IPR000680" |

| Vsp | replicon | gene/pseudo | | | | | | | | |
|---|---|---|---|---|---|---|---|---|---|---|
| gene-bcCo53 001291 c | lp20 | gene | Pfam | protein m | 44 | 210 | 2.00E-59 | + | . | Target=null 44 210;ID=match$761_44_210;signature_desc=Lipoprotein;Name=PF01441;status=T;Dbxref="InterPro:IPR001800" |
| gene-bcCo53 001358 c | Lp24-2 | gene | Pfam | protein m | 50 | 220 | 4.40E-35 | + | . | Target=null 50 220;ID=match$2107_50_220;signature_desc=Lipoprotein;Name=PF01441;status=T;Dbxref="InterPro:IPR001800" |
| gene-bcCo53 001519 c | lp31 | gene | Pfam | protein m | 44 | 210 | 5.80E-39 | + | . | Target=null 44 210;ID=match$985_44_210;signature_desc=Lipoprotein;Name=PF01441;status=T;Dbxref="InterPro:IPR001800" |
| gene-bcCo53 001542 c | lp32 | gene | Pfam | protein m | 39 | 211 | 6.60E-48 | + | . | Target=null 39 211;ID=match$2664_39_211;signature_desc=Lipoprotein;Name=PF01441;status=T;Dbxref="InterPro:IPR001800" |
| gene-bcCo53 001548 c | lp32 | gene | Pfam | protein m | 47 | 209 | 8.00E-42 | + | . | Target=null 47 209;ID=match$219_47_209;signature_desc=Lipoprotein;Name=PF01441;status=T;Dbxref="InterPro:IPR001800" |
| gene-bcCo53 001553 c | lp32 | gene | Pfam | protein m | 48 | 215 | 2.80E-41 | + | . | Target=null 48 215;ID=match$3114_48_215;signature_desc=Lipoprotein;Name=PF01441;status=T;Dbxref="InterPro:IPR001800" |
| gene-bcCo53 001554 c | lp32 | gene | Pfam | protein m | 44 | 208 | 1.00E-51 | + | . | Target=null 44 208;ID=match$2497_44_208;signature_desc=Lipoprotein;Name=PF01441;status=T;Dbxref="InterPro:IPR001800" |
| gene-bcCo53 001556 c | lp32 | gene | Pfam | protein m | 39 | 210 | 1.70E-52 | + | . | Target=null 39 210;ID=match$2418_39_210;signature_desc=Lipoprotein;Name=PF01441;status=T;Dbxref="InterPro:IPR001800" |
| gene-bcCo53 001603 c | lp34 | gene | Pfam | protein m | 44 | 214 | 2.00E-43 | + | . | Target=null 44 214;ID=match$2923_44_214;signature_desc=Lipoprotein;Name=PF01441;status=T;Dbxref="InterPro:IPR001800" |
| gene-bcCo53 001639 c | lp38 | gene | Pfam | protein m | 44 | 214 | 4.70E-43 | + | . | Target=null 44 214;ID=match$1934_44_214;signature_desc=Lipoprotein;Name=PF01441;status=T;Dbxref="InterPro:IPR001800" |
| gene-bcCo53 001642 c | lp38 | pseudo | Pfam | protein m | 1 | 36 | 5.40E-07 | + | . | Target=null 1 36;ID=match$2054_1_36;signature_desc=Lipoprotein;Name=PF01441;status=T;Dbxref="InterPro:IPR001800" |
| gene-bcCo53 001644 c | lp38 | gene | Pfam | protein m | 47 | 216 | 1.50E-36 | + | . | Target=null 47 216;ID=match$599_47_216;signature_desc=Lipoprotein;Name=PF01441;status=T;Dbxref="InterPro:IPR001800" |
| gene-bcCo53 001690 c | lp39 | gene | Pfam | protein m | 5 | 95 | 2.20E-10 | + | . | Target=null 5 95;ID=match$1138_5_95;signature_desc=Lipoprotein;Name=PF01441;status=T;Dbxref="InterPro:IPR001800" |

| Vlp | | | Subfamily | | | | | | | | |
|---|---|---|---|---|---|---|---|---|---|---|---|
| gene-bcCo53 000002 c | chromosome | gene | beta | Pfam | protein m | 275 | 497 | 5.90E-27 | + | . | Target=null 275 497;ID=match$2487_275_497;signature_desc=Borrelia lipoprotein;Name=PF00921;status=T;Dbxref="InterPro:IPR000680" |
| gene-bcCo53 001279 c | lp20 | gene | beta | Pfam | protein m | 9 | 235 | 1.60E-19 | + | . | Target=null 9 235;ID=match$2104_9_235;signature_desc=Borrelia lipoprotein;Name=PF00921;status=T;Dbxref="InterPro:IPR000680" |
| gene-bcCo53 001285 c | lp20 | gene | delta | Pfam | protein m | 49 | 326 | 3.40E-73 | + | . | Target=null 49 326;ID=match$1559_49_326;signature_desc=Borrelia lipoprotein;Name=PF00921;status=T;Dbxref="InterPro:IPR000680" |
| gene-bcCo53 001286 c | lp20 | gene | gamma | Pfam | protein m | 49 | 325 | 2.60E-84 | + | . | Target=null 49 325;ID=match$2448_49_325;signature_desc=Borrelia lipoprotein;Name=PF00921;status=T;Dbxref="InterPro:IPR000680" |
| gene-bcCo53 001287 c | lp20 | gene | gamma | Pfam | protein m | 49 | 334 | 2.30E-79 | + | . | Target=null 49 334;ID=match$586_49_334;signature_desc=Borrelia lipoprotein;Name=PF00921;status=T;Dbxref="InterPro:IPR000680" |
| gene-bcCo53 001288 c | lp20 | gene | gamma | Pfam | protein m | 49 | 342 | 4.20E-85 | + | . | Target=null 49 342;ID=match$992_49_342;signature_desc=Borrelia lipoprotein;Name=PF00921;status=T;Dbxref="InterPro:IPR000680" |
| gene-bcCo53 001289 c | lp20 | gene | gamma | Pfam | protein m | 49 | 342 | 4.20E-85 | + | . | Target=null 49 342;ID=match$3118_49_342;signature_desc=Borrelia lipoprotein;Name=PF00921;status=T;Dbxref="InterPro:IPR000680" |
| gene-bcCo53 001290 c | lp20 | gene | delta | Pfam | protein m | 49 | 330 | 9.00E-88 | + | . | Target=null 49 330;ID=match$1799_49_330;signature_desc=Borrelia lipoprotein;Name=PF00921;status=T;Dbxref="InterPro:IPR000680" |
| gene-bcCo53 001292 c | lp20 | gene | delta | Pfam | protein m | 49 | 328 | 1.70E-80 | + | . | Target=null 49 328;ID=match$2955_49_328;signature_desc=Borrelia lipoprotein;Name=PF00921;status=T;Dbxref="InterPro:IPR000680" |
| gene-bcCo53 001293 c | lp20 | gene | gamma | Pfam | protein m | 109 | 180 | 5.30E-17 | + | . | Target=null 109 180;ID=match$1320_109_180;signature_desc=Borrelia lipoprotein;Name=PF00921;status=T;Dbxref="InterPro:IPR000680" |
| gene-bcCo53 001293 c | lp20 | | | Pfam | protein m | 1 | 100 | 1.30E-09 | + | . | Target=null 1 100;ID=match$1320_1_100;signature_desc=Borrelia lipoprotein;Name=PF00921;status=T;Dbxref="InterPro:IPR000680" |
| gene-bcCo53 001294 c | lp20 | gene | gamma | Pfam | protein m | 6 | 38 | 3.30E-11 | + | . | Target=null 6 38;ID=match$1893_6_38;signature_desc=Borrelia lipoprotein;Name=PF00921;status=T;Dbxref="InterPro:IPR000680" |
| gene-bcCo53 001306 c | lp23 | gene | gamma | Pfam | protein m | 49 | 328 | 2.70E-89 | + | . | Target=null 49 328;ID=match$3338_49_328;signature_desc=Borrelia lipoprotein;Name=PF00921;status=T;Dbxref="InterPro:IPR000680" |
| gene-bcCo53 001307 c | lp23 | gene | beta | Pfam | protein m | 1 | 164 | 2.10E-24 | + | . | Target=null 1 164;ID=match$201_1_164;signature_desc=Borrelia lipoprotein;Name=PF00921;status=T;Dbxref="InterPro:IPR000680" |
| gene-bcCo53 001308 c | lp23 | gene | beta | Pfam | protein m | 1 | 275 | 6.60E-37 | + | . | Target=null 1 275;ID=match$1299_1_275;signature_desc=Borrelia lipoprotein;Name=PF00921;status=T;Dbxref="InterPro:IPR000680" |
| gene-bcCo53 001309 c | lp23 | gene | gamma | Pfam | protein m | 51 | 338 | 2.70E-82 | + | . | Target=null 51 338;ID=match$1653_51_338;signature_desc=Borrelia lipoprotein;Name=PF00921;status=T;Dbxref="InterPro:IPR000680" |
| gene-bcCo53 001310 c | lp23 | gene | gamma | Pfam | protein m | 48 | 290 | 2.50E-52 | + | . | Target=null 48 290;ID=match$2462_48_290;signature_desc=Borrelia lipoprotein;Name=PF00921;status=T;Dbxref="InterPro:IPR000680" |
| gene-bcCo53 001310 c | lp23 | | | Pfam | protein m | 1 | 58 | 7.00E-17 | + | . | Target=null 1 58;ID=match$2696_1_58;signature_desc=Borrelia lipoprotein;Name=PF00921;status=T;Dbxref="InterPro:IPR000680" |
| gene-bcCo53 001311 c | lp23 | gene | alpha | Pfam | protein m | 42 | 338 | 1.60E-57 | + | . | Target=null 42 338;ID=match$2076_42_338;signature_desc=Borrelia lipoprotein;Name=PF00921;status=T;Dbxref="InterPro:IPR000680" |
| gene-bcCo53 001312 c | lp23 | gene | delta | Pfam | protein m | 49 | 333 | 6.20E-61 | + | . | Target=null 49 333;ID=match$2868_49_333;signature_desc=Borrelia lipoprotein;Name=PF00921;status=T;Dbxref="InterPro:IPR000680" |
| gene-bcCo53 001313 c | lp23 | pseudo | alpha | Pfam | protein m | 50 | 75 | 3.80E-06 | + | . | Target=null 50 75;ID=match$1968_50_75;signature_desc=Borrelia lipoprotein;Name=PF00921;status=T;Dbxref="InterPro:IPR000680" |
| gene-bcCo53 001313 c | lp23 | | | Pfam | protein m | 5 | 139 | 1.00E-14 | + | . | Target=null 5 139;ID=match$1544_5_139;signature_desc=Borrelia lipoprotein;Name=PF00921;status=T;Dbxref="InterPro:IPR000680" |
| gene-bcCo53 001322 c | Lp24-1 | gene | delta | Pfam | protein m | 44 | 74 | 2.30E-07 | + | . | Target=null 44 74;ID=match$2193_44_74;signature_desc=Borrelia lipoprotein;Name=PF00921;status=T;Dbxref="InterPro:IPR000680" |
| gene-bcCo53 001323 c | Lp24-1 | gene | beta | Pfam | protein m | 50 | 331 | 8.70E-48 | + | . | Target=null 50 331;ID=match$2461_50_331;signature_desc=Borrelia lipoprotein;Name=PF00921;status=T;Dbxref="InterPro:IPR000680" |
| gene-bcCo53 001324 c | Lp24-1 | gene | gamma | Pfam | protein m | 7 | 73 | 5.20E-20 | + | . | Target=null 7 73;ID=match$2242_7_73;signature_desc=Borrelia lipoprotein;Name=PF00921;status=T;Dbxref="InterPro:IPR000680" |
| gene-bcCo53 001325 c | Lp24-1 | gene | gamma | Pfam | protein m | 3 | 126 | 5.60E-25 | + | . | Target=null 3 126;ID=match$1573_3_126;signature_desc=Borrelia lipoprotein;Name=PF00921;status=T;Dbxref="InterPro:IPR000680" |
| gene-bcCo53 001326 c | Lp24-1 | pseudo | gamma | Pfam | protein m | 1 | 45 | 6.10E-09 | + | . | Target=null 1 45;ID=match$2308_1_45;signature_desc=Borrelia lipoprotein;Name=PF00921;status=T;Dbxref="InterPro:IPR000680" |
| gene-bcCo53 001328 c | Lp24-1 | gene | alpha | Pfam | protein m | 17 | 307 | 5.60E-66 | + | . | Target=null 17 307;ID=match$2595_17_307;signature_desc=Borrelia lipoprotein;Name=PF00921;status=T;Dbxref="InterPro:IPR000680" |
| gene-bcCo53 001329 c | Lp24-1 | gene | gamma | Pfam | protein m | 50 | 335 | 3.00E-75 | + | . | Target=null 50 335;ID=match$2574_50_335;signature_desc=Borrelia lipoprotein;Name=PF00921;status=T;Dbxref="InterPro:IPR000680" |
| gene-bcCo53 001330 c | Lp24-1 | pseudo | alpha | Pfam | protein m | 50 | 85 | 2.10E-08 | + | . | Target=null 50 85;ID=match$2337_50_85;signature_desc=Borrelia lipoprotein;Name=PF00921;status=T;Dbxref="InterPro:IPR000680" |
| gene-bcCo53 001330 c | Lp24-1 | | | Pfam | protein m | 3 | 173 | 4.00E-17 | + | . | Target=null 3 173;ID=match$216_3_173;signature_desc=Borrelia lipoprotein;Name=PF00921;status=T;Dbxref="InterPro:IPR000680" |
| gene-bcCo53 001330 c | Lp24-1 | | | Pfam | protein m | 8 | 84 | 6.50E-26 | + | . | Target=null 8 84;ID=match$1507_8_84;signature_desc=Borrelia lipoprotein;Name=PF00921;status=T;Dbxref="InterPro:IPR000680" |
| gene-bcCo53 001331 c | Lp24-1 | pseudo | beta | Pfam | protein m | 49 | 234 | 3.80E-21 | + | . | Target=null 49 234;ID=match$1950_49_234;signature_desc=Borrelia lipoprotein;Name=PF00921;status=T;Dbxref="InterPro:IPR000680" |
| gene-bcCo53 001331 c | Lp24-1 | | | Pfam | protein m | 2 | 72 | 1.30E-06 | + | . | Target=null 2 72;ID=match$1709_2_72;signature_desc=Borrelia lipoprotein;Name=PF00921;status=T;Dbxref="InterPro:IPR000680" |
| gene-bcCo53 001331 c | Lp24-1 | | | Pfam | protein m | 8 | 50 | 5.50E-12 | + | . | Target=null 8 50;ID=match$182_8_50;signature_desc=Borrelia lipoprotein;Name=PF00921;status=T;Dbxref="InterPro:IPR000680" |
| gene-bcCo53 001332 c | Lp24-1 | gene | beta | Pfam | protein m | 50 | 331 | 1.40E-40 | + | . | Target=null 50 331;ID=match$534_50_331;signature_desc=Borrelia lipoprotein;Name=PF00921;status=T;Dbxref="InterPro:IPR000680" |
| gene-bcCo53 001333 c | Lp24-1 | gene | gamma | Pfam | protein m | 50 | 329 | 5.20E-68 | + | . | Target=null 50 329;ID=match$995_50_329;signature_desc=Borrelia lipoprotein;Name=PF00921;status=T;Dbxref="InterPro:IPR000680" |
| gene-bcCo53 001334 c | Lp24-1 | gene | delta | Pfam | protein m | 49 | 330 | 2.10E-78 | + | . | Target=null 49 330;ID=match$1698_49_330;signature_desc=Borrelia lipoprotein;Name=PF00921;status=T;Dbxref="InterPro:IPR000680" |
| gene-bcCo53 001350 c | Lp24-2 | gene | delta | Pfam | protein m | 45 | 286 | 6.00E-62 | + | . | Target=null 45 286;ID=match$912_45_286;signature_desc=Borrelia lipoprotein;Name=PF00921;status=T;Dbxref="InterPro:IPR000680" |
| gene-bcCo53 001351 c | Lp24-2 | gene | gamma | Pfam | protein m | 18 | 48 | 1.60E-09 | + | . | Target=null 18 48;ID=match$2636_18_48;signature_desc=Borrelia lipoprotein;Name=PF00921;status=T;Dbxref="InterPro:IPR000680" |
| gene-bcCo53 001352 c | Lp24-2 | gene | gamma | Pfam | protein m | 6 | 70 | 2.70E-10 | + | . | Target=null 6 70;ID=match$28_6_70;signature_desc=Borrelia lipoprotein;Name=PF00921;status=T;Dbxref="InterPro:IPR000680" |
| gene-bcCo53 001354 c | Lp24-2 | gene | gamma | Pfam | protein m | 49 | 344 | 6.90E-78 | + | . | Target=null 49 344;ID=match$1916_49_344;signature_desc=Borrelia lipoprotein;Name=PF00921;status=T;Dbxref="InterPro:IPR000680" |
| gene-bcCo53 001355 c | Lp24-2 | pseudo | alpha | Pfam | protein m | 6 | 45 | 1.60E-09 | + | . | Target=null 6 45;ID=match$8_6_45;signature_desc=Borrelia lipoprotein;Name=PF00921;status=T;Dbxref="InterPro:IPR000680" |
| gene-bcCo53 001355 c | Lp24-2 | | | Pfam | protein m | 5 | 204 | 1.10E-38 | + | . | Target=null 5 204;ID=match$2990_5_204;signature_desc=Borrelia lipoprotein;Name=PF00921;status=T;Dbxref="InterPro:IPR000680" |
| gene-bcCo53 001356 c | Lp24-2 | gene | gamma | Pfam | protein m | 49 | 338 | 2.00E-77 | + | . | Target=null 49 338;ID=match$2286_49_338;signature_desc=Borrelia lipoprotein;Name=PF00921;status=T;Dbxref="InterPro:IPR000680" |
| gene-bcCo53 001357 c | Lp24-2 | gene | beta | Pfam | protein m | 50 | 334 | 8.80E-50 | + | . | Target=null 50 334;ID=match$1958_50_334;signature_desc=Borrelia lipoprotein;Name=PF00921;status=T;Dbxref="InterPro:IPR000680" |
| gene-bcCo53 001359 c | Lp24-2 | gene | beta | Pfam | protein m | 50 | 340 | 2.80E-51 | + | . | Target=null 50 340;ID=match$3268_50_340;signature_desc=Borrelia lipoprotein;Name=PF00921;status=T;Dbxref="InterPro:IPR000680" |
| gene-bcCo53 001360 c | Lp24-2 | gene | gamma | Pfam | protein m | 21 | 309 | 4.60E-86 | + | . | Target=null 21 309;ID=match$1610_21_309;signature_desc=Borrelia lipoprotein;Name=PF00921;status=T;Dbxref="InterPro:IPR000680" |
| gene-bcCo53 001361 c | Lp24-2 | gene | gamma | Pfam | protein m | 252 | 321 | 3.70E-27 | + | . | Target=null 252 321;ID=match$2443_252_321;signature_desc=Borrelia lipoprotein;Name=PF00921;status=T;Dbxref="InterPro:IPR000680" |
| gene-bcCo53 001361 c | Lp24-2 | | | Pfam | protein m | 52 | 249 | 2.50E-48 | + | . | Target=null 52 249;ID=match$2443_52_249;signature_desc=Borrelia lipoprotein;Name=PF00921;status=T;Dbxref="InterPro:IPR000680" |
| gene-bcCo53 001362 c | Lp24-2 | gene | alpha | Pfam | protein m | 49 | 345 | 7.20E-64 | + | . | Target=null 49 345;ID=match$2405_49_345;signature_desc=Borrelia lipoprotein;Name=PF00921;status=T;Dbxref="InterPro:IPR000680" |
| gene-bcCo53 001399 c | Lp28-2 | gene | beta | Pfam | protein m | 99 | 353 | 2.80E-20 | + | . | Target=null 99 353;ID=match$1425_99_353;signature_desc=Borrelia lipoprotein;Name=PF00921;status=T;Dbxref="InterPro:IPR000680" |
| gene-bcCo53 001404 c | Lp28-2 | gene | gamma | Pfam | protein m | 49 | 326 | 3.40E-73 | + | . | Target=null 49 326;ID=match$1561_49_326;signature_desc=Borrelia lipoprotein;Name=PF00921;status=T;Dbxref="InterPro:IPR000680" |
| gene-bcCo53 001405 c | Lp28-2 | gene | gamma | Pfam | protein m | 49 | 325 | 2.60E-84 | + | . | Target=null 49 325;ID=match$2446_49_325;signature_desc=Borrelia lipoprotein;Name=PF00921;status=T;Dbxref="InterPro:IPR000680" |
| gene-bcCo53 001406 c | Lp28-2 | gene | gamma | Pfam | protein m | 49 | 334 | 2.30E-79 | + | . | Target=null 49 334;ID=match$588_49_334;signature_desc=Borrelia lipoprotein;Name=PF00921;status=T;Dbxref="InterPro:IPR000680" |
| gene-bcCo53 001407 c | Lp28-2 | gene | gamma | Pfam | protein m | 49 | 342 | 4.20E-85 | + | . | Target=null 49 342;ID=match$994_49_342;signature_desc=Borrelia lipoprotein;Name=PF00921;status=T;Dbxref="InterPro:IPR000680" |
| gene-bcCo53 001408 c | Lp28-2 | gene | gamma | Pfam | protein m | 49 | 342 | 4.20E-85 | + | . | Target=null 49 342;ID=match$3116_49_342;signature_desc=Borrelia lipoprotein;Name=PF00921;status=T;Dbxref="InterPro:IPR000680" |
| gene-bcCo53 001421 c | Lp28-2 | gene | beta | Pfam | protein m | 275 | 497 | 5.90E-27 | + | . | Target=null 275 497;ID=match$2485_275_497;signature_desc=Borrelia lipoprotein;Name=PF00921;status=T;Dbxref="InterPro:IPR000680" |
| gene-bcCo53 001452 c | Lp29-2 | gene | beta | Pfam | protein m | 72 | 347 | 4.60E-27 | + | . | Target=null 72 347;ID=match$1545_72_347;signature_desc=Borrelia lipoprotein;Name=PF00921;status=T;Dbxref="InterPro:IPR000680" |
| gene-bcCo53 001459 c | Lp29-2 | gene | gamma | Pfam | protein m | 49 | 336 | 3.00E-78 | + | . | Target=null 49 336;ID=match$1557_49_336;signature_desc=Borrelia lipoprotein;Name=PF00921;status=T;Dbxref="InterPro:IPR000680" |
| gene-bcCo53 001460 c | Lp29-2 | gene | beta | Pfam | protein m | 38 | 323 | 2.10E-43 | + | . | Target=null 38 323;ID=match$678_38_323;signature_desc=Borrelia lipoprotein;Name=PF00921;status=T;Dbxref="InterPro:IPR000680" |
| gene-bcCo53 001461 c | Lp29-2 | gene | gamma | Pfam | protein m | 50 | 348 | 2.80E-74 | + | . | Target=null 50 348;ID=match$211_50_348;signature_desc=Borrelia lipoprotein;Name=PF00921;status=T;Dbxref="InterPro:IPR000680" |
| gene-bcCo53 001462 c | Lp29-2 | gene | delta | Pfam | protein m | 49 | 335 | 2.70E-83 | + | . | Target=null 49 335;ID=match$123_49_335;signature_desc=Borrelia lipoprotein;Name=PF00921;status=T;Dbxref="InterPro:IPR000680" |
| gene-bcCo53 001463 c | Lp29-2 | pseudo | beta | Pfam | protein m | 49 | 159 | 5.10E-13 | + | . | Target=null 49 159;ID=match$2228_49_159;signature_desc=Borrelia lipoprotein;Name=PF00921;status=T;Dbxref="InterPro:IPR000680" |
| gene-bcCo53 001463 c | Lp29-2 | | | Pfam | protein m | 23 | 200 | 8.50E-27 | + | . | Target=null 23 200;ID=match$1422_23_200;signature_desc=Borrelia lipoprotein;Name=PF00921;status=T;Dbxref="InterPro:IPR000680" |
| gene-bcCo53 001464 c | Lp29-2 | gene | delta | Pfam | protein m | 53 | 342 | 8.50E-76 | + | . | Target=null 53 342;ID=match$328_53_342;signature_desc=Borrelia lipoprotein;Name=PF00921;status=T;Dbxref="InterPro:IPR000680" |
| gene-bcCo53 001465 c | Lp29-2 | gene | gamma | Pfam | protein m | 49 | 338 | 2.70E-83 | + | . | Target=null 49 338;ID=match$2997_49_338;signature_desc=Borrelia lipoprotein;Name=PF00921;status=T;Dbxref="InterPro:IPR000680" |
| gene-bcCo53 001466 c | Lp29-2 | gene | delta | Pfam | protein m | 48 | 333 | 2.30E-59 | + | . | Target=null 48 333;ID=match$1621_48_333;signature_desc=Borrelia lipoprotein;Name=PF00921;status=T;Dbxref="InterPro:IPR000680" |
| gene-bcCo53 001467 c | Lp29-2 | gene | gamma | Pfam | protein m | 50 | 329 | 7.20E-77 | + | . | Target=null 50 329;ID=match$390_50_329;signature_desc=Borrelia lipoprotein;Name=PF00921;status=T;Dbxref="InterPro:IPR000680" |
| gene-bcCo53 001468 c | Lp29-2 | gene | delta | Pfam | protein m | 52 | 329 | 9.10E-83 | + | . | Target=null 52 329;ID=match$1029_52_329;signature_desc=Borrelia lipoprotein;Name=PF00921;status=T;Dbxref="InterPro:IPR000680" |
| gene-bcCo53 001469 c | Lp29-2 | gene | gamma | Pfam | protein m | 49 | 332 | 1.20E-66 | + | . | Target=null 49 332;ID=match$3105_49_332;signature_desc=Borrelia lipoprotein;Name=PF00921;status=T;Dbxref="InterPro:IPR000680" |
| gene-bcCo53 001470 c | Lp29-2 | gene | gamma | Pfam | protein m | 49 | 344 | 9.50E-82 | + | . | Target=null 49 344;ID=match$150_49_344;signature_desc=Borrelia lipoprotein;Name=PF00921;status=T;Dbxref="InterPro:IPR000680" |
| gene-bcCo53 001477 c | lp30 | gene | beta | Pfam | protein m | 75 | 332 | 1.40E-26 | + | . | Target=null 75 332;ID=match$1108_75_332;signature_desc=Borrelia lipoprotein;Name=PF00921;status=T;Dbxref="InterPro:IPR000680" |
| gene-bcCo53 001486 c | lp30 | pseudo | gamma | Pfam | protein m | 45 | 334 | 9.10E-80 | + | . | Target=null 45 334;ID=match$1809_45_334;signature_desc=Borrelia lipoprotein;Name=PF00921;status=T;Dbxref="InterPro:IPR000680" |
| gene-bcCo53 001487 c | lp30 | gene | gamma | Pfam | protein m | 49 | 334 | 3.70E-71 | + | . | Target=null 49 334;ID=match$2004_49_334;signature_desc=Borrelia lipoprotein;Name=PF00921;status=T;Dbxref="InterPro:IPR000680" |
| gene-bcCo53 001488 c | lp30 | gene | delta | Pfam | protein m | 53 | 341 | 4.30E-59 | + | . | Target=null 53 341;ID=match$2949_53_341;signature_desc=Borrelia lipoprotein;Name=PF00921;status=T;Dbxref="InterPro:IPR000680" |
| gene-bcCo53 001489 c | lp30 | gene | gamma | Pfam | protein m | 52 | 339 | 1.40E-71 | + | . | Target=null 52 339;ID=match$311_52_339;signature_desc=Borrelia lipoprotein;Name=PF00921;status=T;Dbxref="InterPro:IPR000680" |
| gene-bcCo53 001490 c | lp30 | gene | delta | Pfam | protein m | 50 | 335 | 8.30E-61 | + | . | Target=null 50 335;ID=match$399_50_335;signature_desc=Borrelia lipoprotein;Name=PF00921;status=T;Dbxref="InterPro:IPR000680" |
| gene-bcCo53 001491 c | lp30 | gene | gamma | Pfam | protein m | 44 | 335 | 9.10E-82 | + | . | Target=null 44 335;ID=match$2655_44_335;signature_desc=Borrelia lipoprotein;Name=PF00921;status=T;Dbxref="InterPro:IPR000680" |
| gene-bcCo53 001492 c | lp30 | pseudo | gamma | Pfam | protein m | 7 | 45 | 3.70E-13 | + | . | Target=null 7 45;ID=match$3205_7_45;signature_desc=Borrelia lipoprotein;Name=PF00921;status=T;Dbxref="InterPro:IPR000680" |
| gene-bcCo53 001493 c | lp30 | gene | delta | Pfam | protein m | 49 | 337 | 3.60E-58 | + | . | Target=null 49 337;ID=match$2162_49_337;signature_desc=Borrelia lipoprotein;Name=PF00921;status=T;Dbxref="InterPro:IPR000680" |
| gene-bcCo53 001494 c | lp30 | gene | delta | Pfam | protein m | 49 | 327 | 1.10E-82 | + | . | Target=null 49 327;ID=match$298_49_327;signature_desc=Borrelia lipoprotein;Name=PF00921;status=T;Dbxref="InterPro:IPR000680" |
| gene-bcCo53 001495 c | lp30 | gene | delta | Pfam | protein m | 49 | 336 | 6.30E-61 | + | . | Target=null 49 336;ID=match$1303_49_336;signature_desc=Borrelia lipoprotein;Name=PF00921;status=T;Dbxref="InterPro:IPR000680" |
| gene-bcCo53 001496 c | lp30 | gene | gamma | Pfam | protein m | 49 | 351 | 1.30E-70 | + | . | Target=null 49 351;ID=match$1228_49_351;signature_desc=Borrelia lipoprotein;Name=PF00921;status=T;Dbxref="InterPro:IPR000680" |
| gene-bcCo53 001497 c | lp30 | pseudo | gamma | Pfam | protein m | 49 | 93 | 1.00E-09 | + | . | Target=null 49 93;ID=match$1555_49_93;signature_desc=Borrelia lipoprotein;Name=PF00921;status=T;Dbxref="InterPro:IPR000680" |
| gene-bcCo53 001497 c | lp30 | | | Pfam | protein m | 1 | 257 | 1.50E-50 | + | . | Target=null 1 257;ID=match$425_1_257;signature_desc=Borrelia lipoprotein;Name=PF00921;status=T;Dbxref="InterPro:IPR000680" |
| gene-bcCo53 001508 c | lp31 | gene | alpha | Pfam | protein m | 5 | 59 | 8.30E-17 | + | . | Target=null 5 59;ID=match$419_5_59;signature_desc=Borrelia lipoprotein;Name=PF00921;status=T;Dbxref="InterPro:IPR000680" |
| gene-bcCo53 001509 c | lp31 | pseudo | alpha | Pfam | protein m | 3 | 89 | 1.80E-09 | + | . | Target=null 3 89;ID=match$1868_3_89;signature_desc=Borrelia lipoprotein;Name=PF00921;status=T;Dbxref="InterPro:IPR000680" |
| gene-bcCo53 001510 c | lp31 | gene | alpha | Pfam | protein m | 57 | 196 | 6.10E-19 | + | . | Target=null 57 196;ID=match$2795_57_196;signature_desc=Borrelia lipoprotein;Name=PF00921;status=T;Dbxref="InterPro:IPR000680" |
| gene-bcCo53 001511 c | lp31 | pseudo | beta | Pfam | protein m | 50 | 181 | 5.90E-15 | + | . | Target=null 50 181;ID=match$706_50_181;signature_desc=Borrelia lipoprotein;Name=PF00921;status=T;Dbxref="InterPro:IPR000680" |
| gene-bcCo53 001511 c | lp31 | | | Pfam | protein m | 26 | 153 | 1.30E-23 | + | . | Target=null 26 153;ID=match$3042_26_153;signature_desc=Borrelia lipoprotein;Name=PF00921;status=T;Dbxref="InterPro:IPR000680" |
| gene-bcCo53 001512 c | lp31 | gene | alpha | Pfam | protein m | 50 | 350 | 3.60E-61 | + | . | Target=null 50 350;ID=match$1140_50_350;signature_desc=Borrelia lipoprotein;Name=PF00921;status=T;Dbxref="InterPro:IPR000680" |
| gene-bcCo53 001513 c | lp31 | pseudo | beta | Pfam | protein m | 6 | 275 | 1.40E-36 | + | . | Target=null 6 275;ID=match$3_6_275;signature_desc=Borrelia lipoprotein;Name=PF00921;status=T;Dbxref="InterPro:IPR000680" |
| gene-bcCo53 001514 c | lp31 | gene | delta | Pfam | protein m | 49 | 329 | 1.40E-80 | + | . | Target=null 49 329;ID=match$160_49_329;signature_desc=Borrelia lipoprotein;Name=PF00921;status=T;Dbxref="InterPro:IPR000680" |
| gene-bcCo53 001515 c | lp31 | gene | beta | Pfam | protein m | 42 | 333 | 1.20E-45 | + | . | Target=null 42 333;ID=match$2094_42_333;signature_desc=Borrelia lipoprotein;Name=PF00921;status=T;Dbxref="InterPro:IPR000680" |
| gene-bcCo53 001516 c | lp31 | gene | delta | Pfam | protein m | 49 | 330 | 3.30E-78 | + | . | Target=null 49 330;ID=match$571_49_330;signature_desc=Borrelia lipoprotein;Name=PF00921;status=T;Dbxref="InterPro:IPR000680" |
| gene-bcCo53 001517 c | lp31 | gene | gamma | Pfam | protein m | 49 | 328 | 7.10E-85 | + | . | Target=null 49 328;ID=match$466_49_328;signature_desc=Borrelia lipoprotein;Name=PF00921;status=T;Dbxref="InterPro:IPR000680" |
| gene-bcCo53 001518 c | lp31 | gene | delta | Pfam | protein m | 49 | 327 | 4.20E-80 | + | . | Target=null 49 327;ID=match$749_49_327;signature_desc=Borrelia lipoprotein;Name=PF00921;status=T;Dbxref="InterPro:IPR000680" |
| gene-bcCo53 001520 c | lp31 | gene | gamma | Pfam | protein m | 31 | 328 | 1.70E-82 | + | . | Target=null 31 328;ID=match$246_31_328;signature_desc=Borrelia lipoprotein;Name=PF00921;status=T;Dbxref="InterPro:IPR000680" |
| gene-bcCo53 001521 c | lp31 | gene | delta | Pfam | protein m | 38 | 315 | 9.90E-72 | + | . | Target=null 38 315;ID=match$55_38_315;signature_desc=Borrelia lipoprotein;Name=PF00921;status=T;Dbxref="InterPro:IPR000680" |
| gene-bcCo53 001527 c | lp31 | gene | gamma | Pfam | protein m | 1 | 32 | 8.50E-08 | + | . | Target=null 1 32;ID=match$2678_1_32;signature_desc=Borrelia lipoprotein;Name=PF00921;status=T;Dbxref="InterPro:IPR000680" |
| gene-bcCo53 001543 c | lp32 | gene | gamma | Pfam | protein m | 49 | 340 | 1.00E-81 | + | . | Target=null 49 340;ID=match$473_49_340;signature_desc=Borrelia lipoprotein;Name=PF00921;status=T;Dbxref="InterPro:IPR000680" |
| gene-bcCo53 001544 c | lp32 | pseudo | beta | Pfam | protein m | 44 | 273 | 1.20E-19 | + | . | Target=null 44 273;ID=match$2631_44_273;signature_desc=Borrelia lipoprotein;Name=PF00921;status=T;Dbxref="InterPro:IPR000680" |
| gene-bcCo53 001544 c | lp32 | | | Pfam | protein m | 6 | 63 | 3.90E-17 | + | . | Target=null 6 63;ID=match$463_6_63;signature_desc=Borrelia lipoprotein;Name=PF00921;status=T;Dbxref="InterPro:IPR000680" |
| gene-bcCo53 001545 c | lp32 | gene | gamma | Pfam | protein m | 33 | 264 | 7.90E-51 | + | . | Target=null 33 264;ID=match$1421_33_264;signature_desc=Borrelia lipoprotein;Name=PF00921;status=T;Dbxref="InterPro:IPR000680" |
| gene-bcCo53 001546 c | lp32 | pseudo | alpha | Pfam | protein m | 1 | 50 | 2.80E-11 | + | . | Target=null 1 50;ID=match$1873_1_50;signature_desc=Borrelia lipoprotein;Name=PF00921;status=T;Dbxref="InterPro:IPR000680" |
| gene-bcCo53 001547 c | lp32 | pseudo | gamma | Pfam | protein m | 49 | 207 | 2.10E-41 | + | . | Target=null 49 207;ID=match$2239_49_207;signature_desc=Borrelia lipoprotein;Name=PF00921;status=T;Dbxref="InterPro:IPR000680" |
| gene-bcCo53 001547 c | lp32 | | | Pfam | protein m | 48 | 168 | 2.90E-37 | + | . | Target=null 48 168;ID=match$1992_48_168;signature_desc=Borrelia lipoprotein;Name=PF00921;status=T;Dbxref="InterPro:IPR000680" |
| gene-bcCo53 001549 c | lp32 | pseudo | delta | Pfam | protein m | 44 | 223 | 2.40E-43 | + | . | Target=null 44 223;ID=match$2530_44_223;signature_desc=Borrelia lipoprotein;Name=PF00921;status=T;Dbxref="InterPro:IPR000680" |
| gene-bcCo53 001549 c | lp32 | | | Pfam | protein m | 6 | 100 | 1.60E-30 | + | . | Target=null 6 100;ID=match$3234_6_100;signature_desc=Borrelia lipoprotein;Name=PF00921;status=T;Dbxref="InterPro:IPR000680" |
| gene-bcCo53 001550 c | lp32 | pseudo | beta | Pfam | protein m | 23 | 146 | 3.90E-20 | + | . | Target=null 23 146;ID=match$647_23_146;signature_desc=Borrelia lipoprotein;Name=PF00921;status=T;Dbxref="InterPro:IPR000680" |
| gene-bcCo53 001552 c | lp32 | pseudo | beta | Pfam | protein m | 10 | 300 | 8.60E-47 | + | . | Target=null 10 300;ID=match$3133_10_300;signature_desc=Borrelia lipoprotein;Name=PF00921;status=T;Dbxref="InterPro:IPR000680" |
| gene-bcCo53 001555 c | lp32 | pseudo | delta | Pfam | protein m | 32 | 309 | 3.00E-82 | + | . | Target=null 32 309;ID=match$602_32_309;signature_desc=Borrelia lipoprotein;Name=PF00921;status=T;Dbxref="InterPro:IPR000680" |
| gene-bcCo53 001557 c | lp32 | pseudo | gamma | Pfam | protein m | 1 | 241 | 6.60E-60 | + | . | Target=null 1 241;ID=match$307_1_241;signature_desc=Borrelia lipoprotein;Name=PF00921;status=T;Dbxref="InterPro:IPR000680" |
| gene-bcCo53 001557 c | lp32 | | | Pfam | protein m | 17 | 54 | 3.70E-13 | + | . | Target=null 17 54;ID=match$1440_17_54;signature_desc=Borrelia lipoprotein;Name=PF00921;status=T;Dbxref="InterPro:IPR000680" |
| gene-bcCo53 001558 c | lp32 | gene | beta | Pfam | protein m | 50 | 198 | 1.50E-19 | + | . | Target=null 50 198;ID=match$1551_50_198;signature_desc=Borrelia lipoprotein;Name=PF00921;status=T;Dbxref="InterPro:IPR000680" |
| gene-bcCo53 001559 c | lp32 | gene | beta | Pfam | protein m | 6 | 113 | 2.80E-10 | + | . | Target=null 6 113;ID=match$3010_6_113;signature_desc=Borrelia lipoprotein;Name=PF00921;status=T;Dbxref="InterPro:IPR000680" |
| gene-bcCo53 001562 c | lp32 | gene | beta | Pfam | protein m | 64 | 329 | 3.90E-23 | + | . | Target=null 64 329;ID=match$108_64_329;signature_desc=Borrelia lipoprotein;Name=PF00921;status=T;Dbxref="InterPro:IPR000680" |
| gene-bcCo53 001589 c | lp33 | gene | gamma | Pfam | protein m | 45 | 337 | 3.10E-88 | + | . | Target=null 45 337;ID=match$469_45_337;signature_desc=Borrelia lipoprotein;Name=PF00921;status=T;Dbxref="InterPro:IPR000680" |
| gene-bcCo53 001597 c | lp34 | gene | gamma | Pfam | protein m | 49 | 337 | 2.30E-67 | + | . | Target=null 49 337;ID=match$2224_49_337;signature_desc=Borrelia lipoprotein;Name=PF00921;status=T;Dbxref="InterPro:IPR000680" |
| gene-bcCo53 001598 c | lp34 | gene | gamma | Pfam | protein m | 49 | 351 | 1.30E-70 | + | . | Target=null 49 351;ID=match$1226_49_351;signature_desc=Borrelia lipoprotein;Name=PF00921;status=T;Dbxref="InterPro:IPR000680" |
| gene-bcCo53 001599 c | lp34 | gene | alpha | Pfam | protein m | 53 | 336 | 1.10E-43 | + | . | Target=null 53 336;ID=match$1243_53_336;signature_desc=Borrelia lipoprotein;Name=PF00921;status=T;Dbxref="InterPro:IPR000680" |
| gene-bcCo53 001600 c | lp34 | gene | gamma | Pfam | protein m | 49 | 329 | 3.10E-80 | + | . | Target=null 49 329;ID=match$1271_49_329;signature_desc=Borrelia lipoprotein;Name=PF00921;status=T;Dbxref="InterPro:IPR000680" |
| gene-bcCo53 001601 c | lp34 | gene | delta | Pfam | protein m | 50 | 334 | 4.30E-64 | + | . | Target=null 50 334;ID=match$1751_50_334;signature_desc=Borrelia lipoprotein;Name=PF00921;status=T;Dbxref="InterPro:IPR000680" |
| gene-bcCo53 001602 c | lp34 | pseudo | alpha | Pfam | protein m | 54 | 109 | 9.80E-07 | + | . | Target=null 54 109;ID=match$1319_54_109;signature_desc=Borrelia lipoprotein;Name=PF00921;status=T;Dbxref="InterPro:IPR000680" |
| gene-bcCo53 001602 c | lp34 | | | Pfam | protein m | 24 | 233 | 2.40E-28 | + | . | Target=null 24 233;ID=match$1949_24_233;signature_desc=Borrelia lipoprotein;Name=PF00921;status=T;Dbxref="InterPro:IPR000680" |
| gene-bcCo53 001604 c | lp34 | gene | gamma | Pfam | protein m | 49 | 329 | 2.30E-79 | + | . | Target=null 49 329;ID=match$1127_49_329;signature_desc=Borrelia lipoprotein;Name=PF00921;status=T;Dbxref="InterPro:IPR000680" |
| gene-bcCo53 001605 c | lp34 | gene | gamma | Pfam | protein m | 49 | 338 | 3.80E-83 | + | . | Target=null 49 338;ID=match$2719_49_338;signature_desc=Borrelia lipoprotein;Name=PF00921;status=T;Dbxref="InterPro:IPR000680" |
| gene-bcCo53 001606 c | lp34 | gene | gamma | Pfam | protein m | 49 | 344 | 2.20E-85 | + | . | Target=null 49 344;ID=match$223_49_344;signature_desc=Borrelia lipoprotein;Name=PF00921;status=T;Dbxref="InterPro:IPR000680" |
| gene-bcCo53 001607 c | lp34 | gene | gamma | Pfam | protein m | 49 | 341 | 4.90E-83 | + | . | Target=null 49 341;ID=match$3276_49_341;signature_desc=Borrelia lipoprotein;Name=PF00921;status=T;Dbxref="InterPro:IPR000680" |
| gene-bcCo53 001608 c | lp34 | gene | gamma | Pfam | protein m | 49 | 335 | 1.10E-81 | + | . | Target=null 49 335;ID=match$2712_49_335;signature_desc=Borrelia lipoprotein;Name=PF00921;status=T;Dbxref="InterPro:IPR000680" |
| gene-bcCo53 001609 c | lp34 | gene | beta | Pfam | protein m | 50 | 337 | 7.00E-45 | + | . | Target=null 50 337;ID=match$1247_50_337;signature_desc=Borrelia lipoprotein;Name=PF00921;status=T;Dbxref="InterPro:IPR000680" |
| gene-bcCo53 001610 c | lp34 | gene | gamma | Pfam | protein m | 1 | 56 | 3.20E-16 | + | . | Target=null 1 56;ID=match$2876_1_56;signature_desc=Borrelia lipoprotein;Name=PF00921;status=T;Dbxref="InterPro:IPR000680" |
| gene-bcCo53 001611 c | lp34 | gene | delta | Pfam | protein m | 50 | 335 | 1.90E-60 | + | . | Target=null 50 335;ID=match$2155_50_335;signature_desc=Borrelia lipoprotein;Name=PF00921;status=T;Dbxref="InterPro:IPR000680" |
| gene-bcCo53 001612 c | lp34 | gene | beta | Pfam | protein m | 51 | 339 | 1.40E-40 | + | . | Target=null 51 339;ID=match$2755_51_339;signature_desc=Borrelia lipoprotein;Name=PF00921;status=T;Dbxref="InterPro:IPR000680" |
| gene-bcCo53 001626 c | lp38 | pseudo | beta | Pfam | protein m | 33 | 173 | 6.60E-19 | + | . | Target=null 33 173;ID=match$427_33_173;signature_desc=Borrelia lipoprotein;Name=PF00921;status=T;Dbxref="InterPro:IPR000680" |
| gene-bcCo53 001626 c | lp38 | | | Pfam | protein m | 18 | 139 | 2.00E-23 | + | . | Target=null 18 139;ID=match$3145_18_139;signature_desc=Borrelia lipoprotein;Name=PF00921;status=T;Dbxref="InterPro:IPR000680" |
| gene-bcCo53 001627 c | lp38 | gene | delta | Pfam | protein m | 44 | 232 | 9.30E-41 | + | . | Target=null 44 232;ID=match$146_44_232;signature_desc=Borrelia lipoprotein;Name=PF00921;status=T;Dbxref="InterPro:IPR000680" |
| gene-bcCo53 001629 c | lp38 | gene | delta | Pfam | protein m | 49 | 326 | 4.00E-77 | + | . | Target=null 49 326;ID=match$1169_49_326;signature_desc=Borrelia lipoprotein;Name=PF00921;status=T;Dbxref="InterPro:IPR000680" |
| gene-bcCo53 001630 c | lp38 | gene | delta | Pfam | protein m | 49 | 329 | 1.20E-73 | + | . | Target=null 49 329;ID=match$2916_49_329;signature_desc=Borrelia lipoprotein;Name=PF00921;status=T;Dbxref="InterPro:IPR000680" |
| gene-bcCo53 001631 c | lp38 | pseudo | alpha | Pfam | protein m | 50 | 80 | 4.70E-06 | + | . | Target=null 50 80;ID=match$1566_50_80;signature_desc=Borrelia lipoprotein;Name=PF00921;status=T;Dbxref="InterPro:IPR000680" |
| gene-bcCo53 001631 c | lp38 | | | Pfam | protein m | 1 | 193 | 2.20E-22 | + | . | Target=null 1 193;ID=match$1924_1_193;signature_desc=Borrelia lipoprotein;Name=PF00921;status=T;Dbxref="InterPro:IPR000680" |
| gene-bcCo53 001631 c | lp38 | | | Pfam | protein m | 8 | 76 | 3.30E-19 | + | . | Target=null 8 76;ID=match$1964_8_76;signature_desc=Borrelia lipoprotein;Name=PF00921;status=T;Dbxref="InterPro:IPR000680" |
| gene-bcCo53 001632 c | lp38 | gene | gamma | Pfam | protein m | 49 | 329 | 3.80E-84 | + | . | Target=null 49 329;ID=match$878_49_329;signature_desc=Borrelia lipoprotein;Name=PF00921;status=T;Dbxref="InterPro:IPR000680" |
| gene-bcCo53 001633 c | lp38 | gene | alpha | Pfam | protein m | 50 | 346 | 1.70E-58 | + | . | Target=null 50 346;ID=match$675_50_346;signature_desc=Borrelia lipoprotein;Name=PF00921;status=T;Dbxref="InterPro:IPR000680" |
| gene-bcCo53 001634 c | lp38 | gene | delta | Pfam | protein m | 49 | 291 | 1.70E-45 | + | . | Target=null 49 291;ID=match$1825_49_291;signature_desc=Borrelia lipoprotein;Name=PF00921;status=T;Dbxref="InterPro:IPR000680" |
| gene-bcCo53 001635 c | lp38 | gene | delta | Pfam | protein m | 1 | 35 | 8.30E-08 | + | . | Target=null 1 35;ID=match$157_1_35;signature_desc=Borrelia lipoprotein;Name=PF00921;status=T;Dbxref="InterPro:IPR000680" |
| gene-bcCo53 001636 c | lp38 | gene | beta | Pfam | protein m | 50 | 338 | 7.40E-45 | + | . | Target=null 50 338;ID=match$2096_50_338;signature_desc=Borrelia lipoprotein;Name=PF00921;status=T;Dbxref="InterPro:IPR000680" |
| gene-bcCo53 001637 c | lp38 | gene | gamma | Pfam | protein m | 49 | 342 | 9.50E-80 | + | . | Target=null 49 342;ID=match$1883_49_342;signature_desc=Borrelia lipoprotein;Name=PF00921;status=T;Dbxref="InterPro:IPR000680" |
| gene-bcCo53 001638 c | lp38 | gene | gamma | Pfam | protein m | 49 | 333 | 2.20E-83 | + | . | Target=null 49 333;ID=match$770_49_333;signature_desc=Borrelia lipoprotein;Name=PF00921;status=T;Dbxref="InterPro:IPR000680" |
| gene-bcCo53 001640 c | lp38 | pseudo | delta | Pfam | protein m | 5 | 281 | 5.40E-77 | + | . | Target=null 5 281;ID=match$978_5_281;signature_desc=Borrelia lipoprotein;Name=PF00921;status=T;Dbxref="InterPro:IPR000680" |
| gene-bcCo53 001641 c | lp38 | pseudo | gamma | Pfam | protein m | 21 | 219 | 2.90E-46 | + | . | Target=null 21 219;ID=match$1367_21_219;signature_desc=Borrelia lipoprotein;Name=PF00921;status=T;Dbxref="InterPro:IPR000680" |
| gene-bcCo53 001641 c | lp38 | | | Pfam | protein m | 3 | 89 | 5.00E-25 | + | . | Target=null 3 89;ID=match$2466_3_89;signature_desc=Borrelia lipoprotein;Name=PF00921;status=T;Dbxref="InterPro:IPR000680" |
| gene-bcCo53 001643 c | lp38 | gene | gamma | Pfam | protein m | 53 | 290 | 6.90E-50 | + | . | Target=null 53 290;ID=match$2750_53_290;signature_desc=Borrelia lipoprotein;Name=PF00921;status=T;Dbxref="InterPro:IPR000680" |
| gene-bcCo53 001645 c | lp38 | gene | alpha | Pfam | protein m | 50 | 346 | 4.40E-62 | + | . | Target=null 50 346;ID=match$2105_50_346;signature_desc=Borrelia lipoprotein;Name=PF00921;status=T;Dbxref="InterPro:IPR000680" |
| gene-bcCo53 001646 c | lp38 | gene | beta | Pfam | protein m | 50 | 333 | 9.30E-41 | + | . | Target=null 50 333;ID=match$1681_50_333;signature_desc=Borrelia lipoprotein;Name=PF00921;status=T;Dbxref="InterPro:IPR000680" |
| gene-bcCo53 001647 c | lp38 | pseudo | gamma | Pfam | protein m | 52 | 221 | 2.40E-35 | + | . | Target=null 52 221;ID=match$2192_52_221;signature_desc=Borrelia lipoprotein;Name=PF00921;status=T;Dbxref="InterPro:IPR000680" |
| gene-bcCo53 001647 c | lp38 | | | Pfam | protein m | 1 | 65 | 9.50E-16 | + | . | Target=null 1 65;ID=match$2378_1_65;signature_desc=Borrelia lipoprotein;Name=PF00921;status=T;Dbxref="InterPro:IPR000680" |
| gene-bcCo53 001647 c | lp38 | | | Pfam | protein m | 25 | 63 | 7.80E-11 | + | . | Target=null 25 63;ID=match$476_25_63;signature_desc=Borrelia lipoprotein;Name=PF00921;status=T;Dbxref="InterPro:IPR000680" |
| gene-bcCo53 001691 c | lp39 | gene | delta | Pfam | protein m | 49 | 329 | 5.20E-78 | + | . | Target=null 49 329;ID=match$3054_49_329;signature_desc=Borrelia lipoprotein;Name=PF00921;status=T;Dbxref="InterPro:IPR000680" |
| gene-bcCo53 001702 c | lp60 | gene | gamma | Pfam | protein m | 49 | 337 | 2.30E-67 | + | . | Target=null 49 337;ID=match$2226_49_337;signature_desc=Borrelia lipoprotein;Name=PF00921;status=T;Dbxref="InterPro:IPR000680" |
| gene-bcCo53 001703 c | lp60 | gene | gamma | Pfam | protein m | 49 | 351 | 1.30E-70 | + | . | Target=null 49 351;ID=match$1224_49_351;signature_desc=Borrelia lipoprotein;Name=PF00921;status=T;Dbxref="InterPro:IPR000680" |
| gene-bcCo53 001704 c | lp60 | gene | delta | Pfam | protein m | 49 | 336 | 6.30E-61 | + | . | Target=null 49 336;ID=match$1301_49_336;signature_desc=Borrelia lipoprotein;Name=PF00921;status=T;Dbxref="InterPro:IPR000680" |
| gene-bcCo53 001705 c | lp60 | gene | delta | Pfam | protein m | 49 | 327 | 1.10E-82 | + | . | Target=null 49 327;ID=match$296_49_327;signature_desc=Borrelia lipoprotein;Name=PF00921;status=T;Dbxref="InterPro:IPR000680" |
| gene-bcCo53 001706 c | lp60 | gene | delta | Pfam | protein m | 49 | 337 | 5.70E-58 | + | . | Target=null 49 337;ID=match$2458_49_337;signature_desc=Borrelia lipoprotein;Name=PF00921;status=T;Dbxref="InterPro:IPR000680" |
| gene-bcCo53 001707 c | lp60 | pseudo | gamma | Pfam | protein m | 7 | 45 | 3.70E-13 | + | . | Target=null 7 45;ID=match$3203_7_45;signature_desc=Borrelia lipoprotein;Name=PF00921;status=T;Dbxref="InterPro:IPR000680" |
| gene-bcCo53 001708 c | lp60 | gene | gamma | Pfam | protein m | 44 | 335 | 9.10E-82 | + | . | Target=null 44 335;ID=match$2653_44_335;signature_desc=Borrelia lipoprotein;Name=PF00921;status=T;Dbxref="InterPro:IPR000680" |
| gene-bcCo53 001709 c | lp60 | gene | beta | Pfam | protein m | 50 | 333 | 6.70E-47 | + | . | Target=null 50 333;ID=match$1374_50_333;signature_desc=Borrelia lipoprotein;Name=PF00921;status=T;Dbxref="InterPro:IPR000680" |
| gene-bcCo53 001710 c | lp60 | gene | gamma | Pfam | protein m | 52 | 339 | 6.40E-82 | + | . | Target=null 52 339;ID=match$3156_52_339;signature_desc=Borrelia lipoprotein;Name=PF00921;status=T;Dbxref="InterPro:IPR000680" |
| gene-bcCo53 001711 c | lp60 | pseudo | beta | Pfam | protein m | 5 | 276 | 6.90E-40 | + | . | Target=null 5 276;ID=match$3128_5_276;signature_desc=Borrelia lipoprotein;Name=PF00921;status=T;Dbxref="InterPro:IPR000680" |
| gene-bcCo53 001712 c | lp60 | pseudo | delta | Pfam | protein m | 6 | 169 | 2.20E-40 | + | . | Target=null 6 169;ID=match$1354_6_169;signature_desc=Borrelia lipoprotein;Name=PF00921;status=T;Dbxref="InterPro:IPR000680" |
| gene-bcCo53 001712 c | lp60 | | | Pfam | protein m | 16 | 75 | 7.00E-16 | + | . | Target=null 16 75;ID=match$1627_16_75;signature_desc=Borrelia lipoprotein;Name=PF00921;status=T;Dbxref="InterPro:IPR000680" |
| gene-bcCo53 001712 c | lp60 | | | Pfam | protein m | 20 | 55 | 1.20E-10 | + | . | Target=null 20 55;ID=match$1597_20_55;signature_desc=Borrelia lipoprotein;Name=PF00921;status=T;Dbxref="InterPro:IPR000680" |
| gene-bcCo53 001715 c | lp60 | gene | gamma | Pfam | protein m | 50 | 331 | 5.50E-81 | + | . | Target=null 50 331;ID=match$1584_50_331;signature_desc=Borrelia lipoprotein;Name=PF00921;status=T;Dbxref="InterPro:IPR000680" |
| gene-bcCo53 001724 c | lp60 | gene | beta | Pfam | protein m | 95 | 353 | 4.80E-23 | + | . | Target=null 95 353;ID=match$2981_95_353;signature_desc=Borrelia lipoprotein;Name=PF00921;status=T;Dbxref="InterPro:IPR000680" |
| gene-bcCo53 001733 c | lp60 | gene | alpha | Pfam | protein m | 50 | 343 | 1.50E-51 | + | . | Target=null 50 343;ID=match$653_50_343;signature_desc=Borrelia lipoprotein;Name=PF00921;status=T;Dbxref="InterPro:IPR000680" |
| gene-bcCo53 001734 c | lp60 | gene | alpha | Pfam | protein m | 56 | 350 | 3.40E-46 | + | . | Target=null 56 350;ID=match$1059_56_350;signature_desc=Borrelia lipoprotein;Name=PF00921;status=T;Dbxref="InterPro:IPR000680" |
| gene-bcCo53 001735 c | lp60 | gene | gamma | Pfam | protein m | 49 | 337 | 1.00E-83 | + | . | Target=null 49 337;ID=match$1813_49_337;signature_desc=Borrelia lipoprotein;Name=PF00921;status=T;Dbxref="InterPro:IPR000680" |
| gene-bcCo53 001736 c | lp60 | gene | beta | Pfam | protein m | 50 | 335 | 2.60E-47 | + | . | Target=null 50 335;ID=match$2518_50_335;signature_desc=Borrelia lipoprotein;Name=PF00921;status=T;Dbxref="InterPro:IPR000680" |
| gene-bcCo53 001737 c | lp60 | gene | delta | Pfam | protein m | 49 | 327 | 1.10E-77 | + | . | Target=null 49 327;ID=match$774_49_327;signature_desc=Borrelia lipoprotein;Name=PF00921;status=T;Dbxref="InterPro:IPR000680" |
| gene-bcCo53 001738 c | lp60 | pseudo | beta | Pfam | protein m | 12 | 51 | 1.20E-07 | + | . | Target=null 12 51;ID=match$143_12_51;signature_desc=Borrelia lipoprotein;Name=PF00921;status=T;Dbxref="InterPro:IPR000680" |
| gene-bcCo53 001739 c | lp60 | gene | alpha | Pfam | protein m | 50 | 348 | 2.20E-66 | + | . | Target=null 50 348;ID=match$2823_50_348;signature_desc=Borrelia lipoprotein;Name=PF00921;status=T;Dbxref="InterPro:IPR000680" |
| gene-bcCo53 001740 c | lp60 | pseudo | gamma | Pfam | protein m | 49 | 261 | 1.30E-50 | + | . | Target=null 49 261;ID=match$3231_49_261;signature_desc=Borrelia lipoprotein;Name=PF00921;status=T;Dbxref="InterPro:IPR000680" |
| gene-bcCo53 001740 c | lp60 | | | Pfam | protein m | 5 | 70 | 1.10E-22 | + | . | Target=null 5 70;ID=match$2541_5_70;signature_desc=Borrelia lipoprotein;Name=PF00921;status=T;Dbxref="InterPro:IPR000680" |
| gene-bcCo53 001741 c | lp60 | gene | delta | Pfam | protein m | 49 | 338 | 1.30E-71 | + | . | Target=null 49 338;ID=match$2188_49_338;signature_desc=Borrelia lipoprotein;Name=PF00921;status=T;Dbxref="InterPro:IPR000680" |
| gene-bcCo53 001742 c | lp60 | pseudo | beta | Pfam | protein m | 50 | 330 | 4.80E-43 | + | . | Target=null 50 330;ID=match$1954_50_330;signature_desc=Borrelia lipoprotein;Name=PF00921;status=T;Dbxref="InterPro:IPR000680" |
| gene-bcCo53 001743 c | lp60 | gene | beta | Pfam | protein m | 50 | 341 | 1.40E-47 | + | . | Target=null 50 341;ID=match$3047_50_341;signature_desc=Borrelia lipoprotein;Name=PF00921;status=T;Dbxref="InterPro:IPR000680" |
| gene-bcCo53 001744 c | lp60 | gene | beta | Pfam | protein m | 50 | 332 | 7.40E-41 | + | . | Target=null 50 332;ID=match$3002_50_332;signature_desc=Borrelia lipoprotein;Name=PF00921;status=T;Dbxref="InterPro:IPR000680" |
| gene-bcCo53 001745 c | lp60 | gene | gamma | Pfam | protein m | 49 | 330 | 1.50E-83 | + | . | Target=null 49 330;ID=match$2342_49_330;signature_desc=Borrelia lipoprotein;Name=PF00921;status=T;Dbxref="InterPro:IPR000680" |
| gene-bcCo53 001746 c | lp60 | pseudo | delta | Pfam | protein m | 18 | 101 | 2.20E-24 | + | . | Target=null 18 101;ID=match$442_18_101;signature_desc=Borrelia lipoprotein;Name=PF00921;status=T;Dbxref="InterPro:IPR000680" |
| gene-bcCo53 001746 c | lp60 | | | Pfam | protein m | 17 | 209 | 1.30E-53 | + | . | Target=null 17 209;ID=match$1959_17_209;signature_desc=Borrelia lipoprotein;Name=PF00921;status=T;Dbxref="InterPro:IPR000680" |

| Vsp | replicon | gene/pseudo | | | | | | | | |
|---|---|---|---|---|---|---|---|---|---|---|
| gene-bpuSUM 001588 | lp25 | gene | Pfam | protein matc | 39 | 204 | 1E-51 | + | . | Target=null 39 204;ID=match$1794_39_204;signature_desc=Lipoprotein;Name=PF01441;status=T;Dbxref="InterPro:IPR001800" |
| gene-bpuSUM 001664 | lp35 | gene | Pfam | protein matc | 40 | 209 | 1.9E-49 | + | . | Target=null 40 209;ID=match$2768_40_209;signature_desc=Lipoprotein;Name=PF01441;status=T;Dbxref="InterPro:IPR001800" |
| gene-bpuSUM 001668 | lp35 | gene | Pfam | protein matc | 39 | 213 | 6.6E-52 | + | . | Target=null 39 213;ID=match$333_39_213;signature_desc=Lipoprotein;Name=PF01441;status=T;Dbxref="InterPro:IPR001800" |
| gene-bpuSUM 001683 | lp35 | gene | Pfam | protein matc | 39 | 206 | 9.3E-50 | + | . | Target=null 39 206;ID=match$1302_39_206;signature_desc=Lipoprotein;Name=PF01441;status=T;Dbxref="InterPro:IPR001800" |
| gene-bpuSUM 001706 | lp39 | gene | Pfam | protein matc | 39 | 208 | 2.7E-55 | + | . | Target=null 39 208;ID=match$3336_39_208;signature_desc=Lipoprotein;Name=PF01441;status=T;Dbxref="InterPro:IPR001800" |
| gene-bpuSUM 001711 | lp39 | gene | Pfam | protein matc | 39 | 206 | 7.2E-40 | + | . | Target=null 39 206;ID=match$3437_39_206;signature_desc=Lipoprotein;Name=PF01441;status=T;Dbxref="InterPro:IPR001800" |
| gene-bpuSUM 001713 | lp39 | pseudo | Pfam | protein matc | 28 | 79 | 1.1E-18 | + | . | Target=null 28 79;ID=match$2180_28_79;signature_desc=Lipoprotein;Name=PF01441;status=T;Dbxref="InterPro:IPR001800" |
| gene-bpuSUM 001713 | lp39 | pseudo | Pfam | protein matc | 1 | 119 | 1.1E-27 | + | . | Target=null 1 119;ID=match$2197_1_119;signature_desc=Lipoprotein;Name=PF01441;status=T;Dbxref="InterPro:IPR001800" |
| gene-bpuSUM 001717 | lp39 | gene | Pfam | protein matc | 39 | 207 | 2.5E-60 | + | . | Target=null 39 207;ID=match$2090_39_207;signature_desc=Lipoprotein;Name=PF01441;status=T;Dbxref="InterPro:IPR001800" |
| gene-bpuSUM 001722 | lp39 | gene | Pfam | protein matc | 39 | 211 | 6.7E-47 | + | . | Target=null 39 211;ID=match$6_39_211;signature_desc=Lipoprotein;Name=PF01441;status=T;Dbxref="InterPro:IPR001800" |
| gene-bpuSUM 001750 | lp41 | gene | Pfam | protein matc | 39 | 201 | 7.5E-49 | + | . | Target=null 39 201;ID=match$110_39_201;signature_desc=Lipoprotein;Name=PF01441;status=T;Dbxref="InterPro:IPR001800" |
| gene-bpuSUM 001764 | lp41 | gene | Pfam | protein matc | 40 | 209 | 1.9E-49 | + | . | Target=null 40 209;ID=match$2770_40_209;signature_desc=Lipoprotein;Name=PF01441;status=T;Dbxref="InterPro:IPR001800" |
| gene-bpuSUM 001778 | lp46 | gene | Pfam | protein matc | 39 | 207 | 1.6E-58 | + | . | Target=null 39 207;ID=match$2332_39_207;signature_desc=Lipoprotein;Name=PF01441;status=T;Dbxref="InterPro:IPR001800" |
| gene-bpuSUM 001781 | lp46 | pseudo | Pfam | protein matc | 39 | 145 | 3.2E-31 | + | . | Target=null 39 145;ID=match$1777_39_145;signature_desc=Lipoprotein;Name=PF01441;status=T;Dbxref="InterPro:IPR001800" |
| gene-bpuSUM 001781 | lp46 | pseudo | Pfam | protein matc | 11 | 74 | 2.7E-16 | + | . | Target=null 11 74;ID=match$2312_11_74;signature_desc=Lipoprotein;Name=PF01441;status=T;Dbxref="InterPro:IPR001800" |
| gene-bpuSUM 001784 | lp46 | pseudo | Pfam | protein matc | 39 | 147 | 1.8E-36 | + | . | Target=null 39 147;ID=match$2721_39_147;signature_desc=Lipoprotein;Name=PF01441;status=T;Dbxref="InterPro:IPR001800" |
| gene-bpuSUM 001784 | lp46 | pseudo | Pfam | protein matc | 2 | 61 | 3.7E-13 | + | . | Target=null 2 61;ID=match$3611_2_61;signature_desc=Lipoprotein;Name=PF01441;status=T;Dbxref="InterPro:IPR001800" |
| gene-bpuSUM 001787 | lp46 | gene | Pfam | protein matc | 39 | 202 | 6E-42 | + | . | Target=null 39 202;ID=match$1316_39_202;signature_desc=Lipoprotein;Name=PF01441;status=T;Dbxref="InterPro:IPR001800" |
| gene-bpuSUM 001795 | lp46 | gene | Pfam | protein matc | 38 | 205 | 6E-65 | + | . | Target=null 38 205;ID=match$3743_38_205;signature_desc=Lipoprotein;Name=PF01441;status=T;Dbxref="InterPro:IPR001800" |
| gene-bpuSUM 001801 | lp46 | gene | Pfam | protein matc | 39 | 206 | 3.4E-65 | + | . | Target=null 39 206;ID=match$2638_39_206;signature_desc=Lipoprotein;Name=PF01441;status=T;Dbxref="InterPro:IPR001800" |
| gene-bpuSUM 001840 | lp47 | gene | Pfam | protein matc | 39 | 202 | 6E-42 | + | . | Target=null 39 202;ID=match$1314_39_202;signature_desc=Lipoprotein;Name=PF01441;status=T;Dbxref="InterPro:IPR001800" |
| gene-bpuSUM 001843 | lp47 | pseudo | Pfam | protein matc | 39 | 147 | 1.8E-36 | + | . | Target=null 39 147;ID=match$2719_39_147;signature_desc=Lipoprotein;Name=PF01441;status=T;Dbxref="InterPro:IPR001800" |
| gene-bpuSUM 001843 | lp47 | pseudo | Pfam | protein matc | 2 | 61 | 3.7E-13 | + | . | Target=null 2 61;ID=match$3607_2_61;signature_desc=Lipoprotein;Name=PF01441;status=T;Dbxref="InterPro:IPR001800" |
| gene-bpuSUM 001846 | lp47 | pseudo | Pfam | protein matc | 39 | 145 | 3.2E-31 | + | . | Target=null 39 145;ID=match$1775_39_145;signature_desc=Lipoprotein;Name=PF01441;status=T;Dbxref="InterPro:IPR001800" |
| gene-bpuSUM 001846 | lp47 | pseudo | Pfam | protein matc | 11 | 74 | 2.7E-16 | + | . | Target=null 11 74;ID=match$2314_11_74;signature_desc=Lipoprotein;Name=PF01441;status=T;Dbxref="InterPro:IPR001800" |
| gene-bpuSUM 001849 | lp47 | gene | Pfam | protein matc | 39 | 207 | 1.6E-58 | + | . | Target=null 39 207;ID=match$2330_39_207;signature_desc=Lipoprotein;Name=PF01441;status=T;Dbxref="InterPro:IPR001800" |
| gene-bpuSUM 001862 | lp57 | gene | Pfam | protein matc | 39 | 207 | 1.6E-58 | + | . | Target=null 39 207;ID=match$2334_39_207;signature_desc=Lipoprotein;Name=PF01441;status=T;Dbxref="InterPro:IPR001800" |
| gene-bpuSUM 001864 | lp57 | gene | Pfam | protein matc | 39 | 207 | 2.4E-65 | + | . | Target=null 39 207;ID=match$83_39_207;signature_desc=Lipoprotein;Name=PF01441;status=T;Dbxref="InterPro:IPR001800" |
| gene-bpuSUM 001873 | lp57 | gene | Pfam | protein matc | 39 | 211 | 6.1E-49 | + | . | Target=null 39 211;ID=match$3568_39_211;signature_desc=Lipoprotein;Name=PF01441;status=T;Dbxref="InterPro:IPR001800" |
| gene-bpuSUM 001894 | lp57 | gene | Pfam | protein matc | 39 | 206 | 2.3E-54 | + | . | Target=null 39 206;ID=match$1408_39_206;signature_desc=Lipoprotein;Name=PF01441;status=T;Dbxref="InterPro:IPR001800" |
| gene-bpuSUM 001896 | lp57 | gene | Pfam | protein matc | 28 | 196 | 2.7E-55 | + | . | Target=null 28 196;ID=match$3575_28_196;signature_desc=Lipoprotein;Name=PF01441;status=T;Dbxref="InterPro:IPR001800" |
| gene-bpuSUM 001899 | lp57 | gene | Pfam | protein matc | 38 | 206 | 1.6E-63 | + | . | Target=null 38 206;ID=match$214_38_206;signature_desc=Lipoprotein;Name=PF01441;status=T;Dbxref="InterPro:IPR001800" |
| gene-bpuSUM 001905 | lp57 | gene | Pfam | protein matc | 39 | 211 | 6.7E-47 | + | . | Target=null 39 211;ID=match$4_39_211;signature_desc=Lipoprotein;Name=PF01441;status=T;Dbxref="InterPro:IPR001800" |
| gene-bpuSUM 001918 | lp65 | gene | Pfam | protein matc | 28 | 196 | 3.8E-62 | + | . | Target=null 28 196;ID=match$2605_28_196;signature_desc=Lipoprotein;Name=PF01441;status=T;Dbxref="InterPro:IPR001800" |
| gene-bpuSUM 001930 | lp65 | gene | Pfam | protein matc | 39 | 206 | 8.7E-61 | + | . | Target=null 39 206;ID=match$1157_39_206;signature_desc=Lipoprotein;Name=PF01441;status=T;Dbxref="InterPro:IPR001800" |
| gene-bpuSUM 001935 | lp65 | gene | Pfam | protein matc | 40 | 206 | 1.9E-63 | + | . | Target=null 40 206;ID=match$2241_40_206;signature_desc=Lipoprotein;Name=PF01441;status=T;Dbxref="InterPro:IPR001800" |
| gene-bpuSUM 001937 | lp65 | gene | Pfam | protein matc | 39 | 206 | 1.2E-56 | + | . | Target=null 39 206;ID=match$923_39_206;signature_desc=Lipoprotein;Name=PF01441;status=T;Dbxref="InterPro:IPR001800" |
| gene-bpuSUM 001952 | lp65 | gene | Pfam | protein matc | 40 | 208 | 4.1E-63 | + | . | Target=null 40 208;ID=match$3700_40_208;signature_desc=Lipoprotein;Name=PF01441;status=T;Dbxref="InterPro:IPR001800" |
| gene-bpuSUM 001956 | lp65 | gene | Pfam | protein matc | 38 | 205 | 6E-65 | + | . | Target=null 38 205;ID=match$3745_38_205;signature_desc=Lipoprotein;Name=PF01441;status=T;Dbxref="InterPro:IPR001800" |
| gene-bpuSUM 001964 | lp65 | gene | Pfam | protein matc | 39 | 202 | 6E-42 | + | . | Target=null 39 202;ID=match$1318_39_202;signature_desc=Lipoprotein;Name=PF01441;status=T;Dbxref="InterPro:IPR001800" |
| gene-bpuSUM 001967 | lp65 | pseudo | Pfam | protein matc | 39 | 147 | 1.8E-36 | + | . | Target=null 39 147;ID=match$2723_39_147;signature_desc=Lipoprotein;Name=PF01441;status=T;Dbxref="InterPro:IPR001800" |
| gene-bpuSUM 001967 | lp65 | pseudo | Pfam | protein matc | 2 | 61 | 3.7E-13 | + | . | Target=null 2 61;ID=match$3609_2_61;signature_desc=Lipoprotein;Name=PF01441;status=T;Dbxref="InterPro:IPR001800" |
| gene-bpuSUM 001970 | lp65 | gene | Pfam | protein matc | 40 | 208 | 1.1E-49 | + | . | Target=null 40 208;ID=match$1149_40_208;signature_desc=Lipoprotein;Name=PF01441;status=T;Dbxref="InterPro:IPR001800" |
| gene-bpuSUM 001973 | lp65 | gene | Pfam | protein matc | 39 | 207 | 1.6E-58 | + | . | Target=null 39 207;ID=match$2336_39_207;signature_desc=Lipoprotein;Name=PF01441;status=T;Dbxref="InterPro:IPR001800" |

| Vlp | | | Subfamily | | | | | | | | |
|---|---|---|---|---|---|---|---|---|---|---|---|
| gene-bpuSUM 001573 | lp25 | pseudo | beta | Pfam | protein matc | 2 | 116 | 3.6E-11 | + | . | Target=null 2 116;ID=match$422_2_116;signature_desc=Borrelia lipoprotein;Name=PF00921;status=T;Dbxref="InterPro:IPR000680" |
| gene-bpuSUM 001573 | lp25 | pseudo | | Pfam | protein matc | 109 | 159 | 9.8E-11 | + | . | Target=null 109 159;ID=match$3635_109_159;signature_desc=Borrelia lipoprotein;Name=PF00921;status=T;Dbxref="InterPro:IPR000680" |
| gene-bpuSUM 001573 | lp25 | pseudo | | Pfam | protein matc | 33 | 110 | 0.000019 | + | . | Target=null 33 110;ID=match$3635_33_110;signature_desc=Borrelia lipoprotein;Name=PF00921;status=T;Dbxref="InterPro:IPR000680" |
| gene-bpuSUM 001574 | lp25 | gene | alpha | Pfam | protein matc | 58 | 347 | 1.4E-68 | + | . | Target=null 58 347;ID=match$2290_58_347;signature_desc=Borrelia lipoprotein;Name=PF00921;status=T;Dbxref="InterPro:IPR000680" |
| gene-bpuSUM 001586 | lp25 | gene | gamma | Pfam | protein matc | 1 | 270 | 2.6E-81 | + | . | Target=null 1 270;ID=match$2519_1_270;signature_desc=Borrelia lipoprotein;Name=PF00921;status=T;Dbxref="InterPro:IPR000680" |
| gene-bpuSUM 001589 | lp25 | gene | delta | Pfam | protein matc | 19 | 298 | 2.2E-85 | + | . | Target=null 19 298;ID=match$660_19_298;signature_desc=Borrelia lipoprotein;Name=PF00921;status=T;Dbxref="InterPro:IPR000680" |
| gene-bpuSUM 001590 | lp25 | gene | gamma | Pfam | protein matc | 213 | 269 | 2.7E-18 | + | . | Target=null 213 269;ID=match$1873_213_269;signature_desc=Borrelia lipoprotein;Name=PF00921;status=T;Dbxref="InterPro:IPR000680" |
| gene-bpuSUM 001590 | lp25 | gene | | Pfam | protein matc | 34 | 215 | 5.2E-44 | + | . | Target=null 34 215;ID=match$1873_34_215;signature_desc=Borrelia lipoprotein;Name=PF00921;status=T;Dbxref="InterPro:IPR000680" |
| gene-bpuSUM 001661 | lp35 | pseudo | gamma | Pfam | protein matc | 34 | 82 | 2.6E-16 | + | . | Target=null 34 82;ID=match$3347_34_82;signature_desc=Borrelia lipoprotein;Name=PF00921;status=T;Dbxref="InterPro:IPR000680" |
| gene-bpuSUM 001661 | lp35 | pseudo | | Pfam | protein matc | 3 | 46 | 0.00000052 | + | . | Target=null 3 46;ID=match$542_3_46;signature_desc=Borrelia lipoprotein;Name=PF00921;status=T;Dbxref="InterPro:IPR000680" |
| gene-bpuSUM 001661 | lp35 | pseudo | | Pfam | protein matc | 19 | 197 | 4.4E-45 | + | . | Target=null 19 197;ID=match$882_19_197;signature_desc=Borrelia lipoprotein;Name=PF00921;status=T;Dbxref="InterPro:IPR000680" |
| gene-bpuSUM 001662 | lp35 | pseudo | delta | Pfam | protein matc | 14 | 288 | 2.7E-83 | + | . | Target=null 14 288;ID=match$1031_14_288;signature_desc=Borrelia lipoprotein;Name=PF00921;status=T;Dbxref="InterPro:IPR000680" |
| gene-bpuSUM 001663 | lp35 | pseudo | alpha | Pfam | protein matc | 56 | 305 | 6E-55 | + | . | Target=null 56 305;ID=match$2669_56_305;signature_desc=Borrelia lipoprotein;Name=PF00921;status=T;Dbxref="InterPro:IPR000680" |
| gene-bpuSUM 001663 | lp35 | pseudo | | Pfam | protein matc | 2 | 34 | 0.00000061 | + | . | Target=null 2 34;ID=match$1458_2_34;signature_desc=Borrelia lipoprotein;Name=PF00921;status=T;Dbxref="InterPro:IPR000680" |
| gene-bpuSUM 001665 | lp35 | gene | gamma | Pfam | protein matc | 34 | 307 | 2.6E-83 | + | . | Target=null 34 307;ID=match$1755_34_307;signature_desc=Borrelia lipoprotein;Name=PF00921;status=T;Dbxref="InterPro:IPR000680" |
| gene-bpuSUM 001667 | lp35 | pseudo | beta | Pfam | protein matc | 19 | 60 | 0.000065 | + | . | Target=null 19 60;ID=match$1282_19_60;signature_desc=Borrelia lipoprotein;Name=PF00921;status=T;Dbxref="InterPro:IPR000680" |
| gene-bpuSUM 001667 | lp35 | pseudo | | Pfam | protein matc | 27 | 155 | 3.1E-25 | + | . | Target=null 27 155;ID=match$2921_27_155;signature_desc=Borrelia lipoprotein;Name=PF00921;status=T;Dbxref="InterPro:IPR000680" |
| gene-bpuSUM 001669 | lp35 | pseudo | delta | Pfam | protein matc | 44 | 120 | 4.5E-21 | + | . | Target=null 44 120;ID=match$2034_44_120;signature_desc=Borrelia lipoprotein;Name=PF00921;status=T;Dbxref="InterPro:IPR000680" |
| gene-bpuSUM 001669 | lp35 | pseudo | | Pfam | protein matc | 8 | 209 | 2.2E-56 | + | . | Target=null 8 209;ID=match$3039_8_209;signature_desc=Borrelia lipoprotein;Name=PF00921;status=T;Dbxref="InterPro:IPR000680" |
| gene-bpuSUM 001672 | lp35 | gene | delta | Pfam | protein matc | 44 | 324 | 1.5E-85 | + | . | Target=null 44 324;ID=match$2227_44_324;signature_desc=Borrelia lipoprotein;Name=PF00921;status=T;Dbxref="InterPro:IPR000680" |
| gene-bpuSUM 001673 | lp35 | gene | beta | Pfam | protein matc | 44 | 333 | 2E-46 | + | . | Target=null 44 333;ID=match$270_44_333;signature_desc=Borrelia lipoprotein;Name=PF00921;status=T;Dbxref="InterPro:IPR000680" |
| gene-bpuSUM 001682 | lp35 | pseudo | delta | Pfam | protein matc | 34 | 155 | 7.9E-37 | + | . | Target=null 34 155;ID=match$1056_34_155;signature_desc=Borrelia lipoprotein;Name=PF00921;status=T;Dbxref="InterPro:IPR000680" |
| gene-bpuSUM 001682 | lp35 | pseudo | | Pfam | protein matc | 2 | 147 | 4.7E-39 | + | . | Target=null 2 147;ID=match$1243_2_147;signature_desc=Borrelia lipoprotein;Name=PF00921;status=T;Dbxref="InterPro:IPR000680" |
| gene-bpuSUM 001684 | lp35 | gene | alpha | Pfam | protein matc | 7 | 303 | 3.3E-68 | + | . | Target=null 7 303;ID=match$3708_7_303;signature_desc=Borrelia lipoprotein;Name=PF00921;status=T;Dbxref="InterPro:IPR000680" |
| gene-bpuSUM 001685 | lp35 | pseudo | beta | Pfam | protein matc | 14 | 260 | 3.7E-35 | + | . | Target=null 14 260;ID=match$2337_14_260;signature_desc=Borrelia lipoprotein;Name=PF00921;status=T;Dbxref="InterPro:IPR000680" |
| gene-bpuSUM 001685 | lp35 | pseudo | | Pfam | protein matc | 2 | 38 | 2.7E-09 | + | . | Target=null 2 38;ID=match$3350_2_38;signature_desc=Borrelia lipoprotein;Name=PF00921;status=T;Dbxref="InterPro:IPR000680" |
| gene-bpuSUM 001686 | lp35 | gene | alpha | Pfam | protein matc | 49 | 341 | 6.6E-71 | + | . | Target=null 49 341;ID=match$1292_49_341;signature_desc=Borrelia lipoprotein;Name=PF00921;status=T;Dbxref="InterPro:IPR000680" |
| gene-bpuSUM 001687 | lp35 | pseudo | beta | Pfam | protein matc | 9 | 292 | 4E-47 | + | . | Target=null 9 292;ID=match$1190_9_292;signature_desc=Borrelia lipoprotein;Name=PF00921;status=T;Dbxref="InterPro:IPR000680" |
| gene-bpuSUM 001691 | lp39 | pseudo | delta | Pfam | protein matc | 34 | 60 | 0.000014 | + | . | Target=null 34 60;ID=match$639_34_60;signature_desc=Borrelia lipoprotein;Name=PF00921;status=T;Dbxref="InterPro:IPR000680" |
| gene-bpuSUM 001691 | lp39 | pseudo | | Pfam | protein matc | 1 | 268 | 1.2E-75 | + | . | Target=null 1 268;ID=match$1624_1_268;signature_desc=Borrelia lipoprotein;Name=PF00921;status=T;Dbxref="InterPro:IPR000680" |
| gene-bpuSUM 001704 | lp39 | pseudo | gamma | Pfam | protein matc | 2 | 51 | 3.1E-10 | + | . | Target=null 2 51;ID=match$1740_2_51;signature_desc=Borrelia lipoprotein;Name=PF00921;status=T;Dbxref="InterPro:IPR000680" |
| gene-bpuSUM 001704 | lp39 | pseudo | | Pfam | protein matc | 2 | 110 | 9.6E-19 | + | . | Target=null 2 110;ID=match$2501_2_110;signature_desc=Borrelia lipoprotein;Name=PF00921;status=T;Dbxref="InterPro:IPR000680" |
| gene-bpuSUM 001705 | lp39 | gene | delta | Pfam | protein matc | 1 | 270 | 6.8E-80 | + | . | Target=null 1 270;ID=match$2737_1_270;signature_desc=Borrelia lipoprotein;Name=PF00921;status=T;Dbxref="InterPro:IPR000680" |
| gene-bpuSUM 001707 | lp39 | gene | delta | Pfam | protein matc | 32 | 166 | 3.7E-36 | + | . | Target=null 32 166;ID=match$3007_32_166;signature_desc=Borrelia lipoprotein;Name=PF00921;status=T;Dbxref="InterPro:IPR000680" |
| gene-bpuSUM 001709 | lp39 | gene | alpha | Pfam | protein matc | 52 | 344 | 3.9E-70 | + | . | Target=null 52 344;ID=match$2879_52_344;signature_desc=Borrelia lipoprotein;Name=PF00921;status=T;Dbxref="InterPro:IPR000680" |
| gene-bpuSUM 001710 | lp39 | gene | beta | Pfam | protein matc | 23 | 304 | 3.1E-48 | + | . | Target=null 23 304;ID=match$2043_23_304;signature_desc=Borrelia lipoprotein;Name=PF00921;status=T;Dbxref="InterPro:IPR000680" |
| gene-bpuSUM 001712 | lp39 | pseudo | delta | Pfam | protein matc | 8 | 115 | 7.6E-26 | + | . | Target=null 8 115;ID=match$1102_8_115;signature_desc=Borrelia lipoprotein;Name=PF00921;status=T;Dbxref="InterPro:IPR000680" |
| gene-bpuSUM 001712 | lp39 | pseudo | | Pfam | protein matc | 4 | 164 | 1.5E-37 | + | . | Target=null 4 164;ID=match$505_4_164;signature_desc=Borrelia lipoprotein;Name=PF00921;status=T;Dbxref="InterPro:IPR000680" |
| gene-bpuSUM 001714 | lp39 | gene | gamma | Pfam | protein matc | 34 | 308 | 1.6E-83 | + | . | Target=null 34 308;ID=match$2038_34_308;signature_desc=Borrelia lipoprotein;Name=PF00921;status=T;Dbxref="InterPro:IPR000680" |
| gene-bpuSUM 001715 | lp39 | pseudo | beta | Pfam | protein matc | 34 | 92 | 0.00000079 | + | . | Target=null 34 92;ID=match$213_34_92;signature_desc=Borrelia lipoprotein;Name=PF00921;status=T;Dbxref="InterPro:IPR000680" |
| gene-bpuSUM 001715 | lp39 | pseudo | | Pfam | protein matc | 5 | 236 | 2.1E-29 | + | . | Target=null 5 236;ID=match$3767_5_236;signature_desc=Borrelia lipoprotein;Name=PF00921;status=T;Dbxref="InterPro:IPR000680" |
| gene-bpuSUM 001716 | lp39 | gene | gamma | Pfam | protein matc | 34 | 313 | 9E-89 | + | . | Target=null 34 313;ID=match$1387_34_313;signature_desc=Borrelia lipoprotein;Name=PF00921;status=T;Dbxref="InterPro:IPR000680" |
| gene-bpuSUM 001718 | lp39 | pseudo | delta | Pfam | protein matc | 19 | 288 | 8.5E-77 | + | . | Target=null 19 288;ID=match$586_19_288;signature_desc=Borrelia lipoprotein;Name=PF00921;status=T;Dbxref="InterPro:IPR000680" |
| gene-bpuSUM 001720 | lp39 | gene | delta | Pfam | protein matc | 44 | 334 | 2.8E-88 | + | . | Target=null 44 334;ID=match$510_44_334;signature_desc=Borrelia lipoprotein;Name=PF00921;status=T;Dbxref="InterPro:IPR000680" |
| gene-bpuSUM 001721 | lp39 | gene | gamma | Pfam | protein matc | 44 | 318 | 2.1E-81 | + | . | Target=null 44 318;ID=match$188_44_318;signature_desc=Borrelia lipoprotein;Name=PF00921;status=T;Dbxref="InterPro:IPR000680" |
| gene-bpuSUM 001723 | lp39 | gene | delta | Pfam | protein matc | 33 | 306 | 5E-85 | + | . | Target=null 33 306;ID=match$3192_33_306;signature_desc=Borrelia lipoprotein;Name=PF00921;status=T;Dbxref="InterPro:IPR000680" |
| gene-bpuSUM 001726 | lp39 | gene | gamma | Pfam | protein matc | 44 | 330 | 4.6E-91 | + | . | Target=null 44 330;ID=match$3180_44_330;signature_desc=Borrelia lipoprotein;Name=PF00921;status=T;Dbxref="InterPro:IPR000680" |
| gene-bpuSUM 001727 | lp39 | pseudo | alpha | Pfam | protein matc | 53 | 335 | 7E-69 | + | . | Target=null 53 335;ID=match$1219_53_335;signature_desc=Borrelia lipoprotein;Name=PF00921;status=T;Dbxref="InterPro:IPR000680" |
| gene-bpuSUM 001741 | lp41 | pseudo | gamma | Pfam | protein matc | 34 | 121 | 3.6E-26 | + | . | Target=null 34 121;ID=match$325_34_121;signature_desc=Borrelia lipoprotein;Name=PF00921;status=T;Dbxref="InterPro:IPR000680" |
| gene-bpuSUM 001741 | lp41 | pseudo | | Pfam | protein matc | 12 | 61 | 3.9E-09 | + | . | Target=null 12 61;ID=match$1681_12_61;signature_desc=Borrelia lipoprotein;Name=PF00921;status=T;Dbxref="InterPro:IPR000680" |
| gene-bpuSUM 001741 | lp41 | pseudo | | Pfam | protein matc | 22 | 150 | 2.4E-35 | + | . | Target=null 22 150;ID=match$857_22_150;signature_desc=Borrelia lipoprotein;Name=PF00921;status=T;Dbxref="InterPro:IPR000680" |
| gene-bpuSUM 001748 | lp41 | pseudo | delta | Pfam | protein matc | 34 | 235 | 1.1E-48 | + | . | Target=null 34 235;ID=match$267_34_235;signature_desc=Borrelia lipoprotein;Name=PF00921;status=T;Dbxref="InterPro:IPR000680" |
| gene-bpuSUM 001748 | lp41 | pseudo | | Pfam | protein matc | 3 | 45 | 2E-13 | + | . | Target=null 3 45;ID=match$3086_3_45;signature_desc=Borrelia lipoprotein;Name=PF00921;status=T;Dbxref="InterPro:IPR000680" |
| gene-bpuSUM 001749 | lp41 | gene | gamma | Pfam | protein matc | 44 | 326 | 4.8E-87 | + | . | Target=null 44 326;ID=match$965_44_326;signature_desc=Borrelia lipoprotein;Name=PF00921;status=T;Dbxref="InterPro:IPR000680" |
| gene-bpuSUM 001751 | lp41 | pseudo | alpha | Pfam | protein matc | 47 | 79 | 0.00000084 | + | . | Target=null 47 79;ID=match$2029_47_79;signature_desc=Borrelia lipoprotein;Name=PF00921;status=T;Dbxref="InterPro:IPR000680" |
| gene-bpuSUM 001751 | lp41 | pseudo | | Pfam | protein matc | 5 | 266 | 1.6E-52 | + | . | Target=null 5 266;ID=match$11_5_266;signature_desc=Borrelia lipoprotein;Name=PF00921;status=T;Dbxref="InterPro:IPR000680" |
| gene-bpuSUM 001752 | lp41 | gene | delta | Pfam | protein matc | 1 | 250 | 4E-69 | + | . | Target=null 1 250;ID=match$1918_1_250;signature_desc=Borrelia lipoprotein;Name=PF00921;status=T;Dbxref="InterPro:IPR000680" |
| gene-bpuSUM 001753 | lp41 | gene | gamma | Pfam | protein matc | 34 | 319 | 1E-83 | + | . | Target=null 34 319;ID=match$2298_34_319;signature_desc=Borrelia lipoprotein;Name=PF00921;status=T;Dbxref="InterPro:IPR000680" |
| gene-bpuSUM 001757 | lp41 | gene | beta | Pfam | protein matc | 44 | 331 | 1E-46 | + | . | Target=null 44 331;ID=match$1726_44_331;signature_desc=Borrelia lipoprotein;Name=PF00921;status=T;Dbxref="InterPro:IPR000680" |
| gene-bpuSUM 001758 | lp41 | gene | delta | Pfam | protein matc | 45 | 323 | 1.2E-88 | + | . | Target=null 45 323;ID=match$2596_45_323;signature_desc=Borrelia lipoprotein;Name=PF00921;status=T;Dbxref="InterPro:IPR000680" |
| gene-bpuSUM 001759 | lp41 | gene | alpha | Pfam | protein matc | 42 | 333 | 2.6E-70 | + | . | Target=null 42 333;ID=match$2110_42_333;signature_desc=Borrelia lipoprotein;Name=PF00921;status=T;Dbxref="InterPro:IPR000680" |
| gene-bpuSUM 001760 | lp41 | gene | beta | Pfam | protein matc | 13 | 301 | 4.2E-50 | + | . | Target=null 13 301;ID=match$1695_13_301;signature_desc=Borrelia lipoprotein;Name=PF00921;status=T;Dbxref="InterPro:IPR000680" |
| gene-bpuSUM 001761 | lp41 | gene | gamma | Pfam | protein matc | 44 | 318 | 1.9E-83 | + | . | Target=null 44 318;ID=match$2141_44_318;signature_desc=Borrelia lipoprotein;Name=PF00921;status=T;Dbxref="InterPro:IPR000680" |
| gene-bpuSUM 001762 | lp41 | gene | alpha | Pfam | protein matc | 52 | 344 | 2.2E-70 | + | . | Target=null 52 344;ID=match$3505_52_344;signature_desc=Borrelia lipoprotein;Name=PF00921;status=T;Dbxref="InterPro:IPR000680" |
| gene-bpuSUM 001763 | lp41 | pseudo | beta | Pfam | protein matc | 23 | 178 | 2.2E-15 | + | . | Target=null 23 178;ID=match$2199_23_178;signature_desc=Borrelia lipoprotein;Name=PF00921;status=T;Dbxref="InterPro:IPR000680" |
| gene-bpuSUM 001763 | lp41 | pseudo | | Pfam | protein matc | 3 | 124 | 1.8E-25 | + | . | Target=null 3 124;ID=match$1978_3_124;signature_desc=Borrelia lipoprotein;Name=PF00921;status=T;Dbxref="InterPro:IPR000680" |
| gene-bpuSUM 001765 | lp41 | pseudo | alpha | Pfam | protein matc | 56 | 305 | 6E-55 | + | . | Target=null 56 305;ID=match$2667_56_305;signature_desc=Borrelia lipoprotein;Name=PF00921;status=T;Dbxref="InterPro:IPR000680" |
| gene-bpuSUM 001765 | lp41 | pseudo | | Pfam | protein matc | 2 | 34 | 0.00000061 | + | . | Target=null 2 34;ID=match$1456_2_34;signature_desc=Borrelia lipoprotein;Name=PF00921;status=T;Dbxref="InterPro:IPR000680" |
| gene-bpuSUM 001766 | lp41 | pseudo | delta | Pfam | protein matc | 1 | 265 | 7.3E-77 | + | . | Target=null 1 265;ID=match$265_1_265;signature_desc=Borrelia lipoprotein;Name=PF00921;status=T;Dbxref="InterPro:IPR000680" |
| gene-bpuSUM 001767 | lp41 | pseudo | gamma | Pfam | protein matc | 34 | 82 | 2.6E-16 | + | . | Target=null 34 82;ID=match$3349_34_82;signature_desc=Borrelia lipoprotein;Name=PF00921;status=T;Dbxref="InterPro:IPR000680" |
| gene-bpuSUM 001767 | lp41 | pseudo | | Pfam | protein matc | 3 | 46 | 0.00000052 | + | . | Target=null 3 46;ID=match$541_3_46;signature_desc=Borrelia lipoprotein;Name=PF00921;status=T;Dbxref="InterPro:IPR000680" |
| gene-bpuSUM 001767 | lp41 | pseudo | | Pfam | protein matc | 19 | 197 | 4.4E-45 | + | . | Target=null 19 197;ID=match$884_19_197;signature_desc=Borrelia lipoprotein;Name=PF00921;status=T;Dbxref="InterPro:IPR000680" |
| gene-bpuSUM 001775 | lp46 | pseudo | alpha | Pfam | protein matc | 8 | 62 | 2.2E-13 | + | . | Target=null 8 62;ID=match$2316_8_62;signature_desc=Borrelia lipoprotein;Name=PF00921;status=T;Dbxref="InterPro:IPR000680" |
| gene-bpuSUM 001775 | lp46 | pseudo | | Pfam | protein matc | 11 | 245 | 5.1E-51 | + | . | Target=null 11 245;ID=match$2701_11_245;signature_desc=Borrelia lipoprotein;Name=PF00921;status=T;Dbxref="InterPro:IPR000680" |
| gene-bpuSUM 001776 | lp46 | pseudo | gamma | Pfam | protein matc | 19 | 150 | 3.4E-44 | + | . | Target=null 19 150;ID=match$1917_19_150;signature_desc=Borrelia lipoprotein;Name=PF00921;status=T;Dbxref="InterPro:IPR000680" |
| gene-bpuSUM 001776 | lp46 | pseudo | | Pfam | protein matc | 5 | 163 | 6.6E-42 | + | . | Target=null 5 163;ID=match$1951_5_163;signature_desc=Borrelia lipoprotein;Name=PF00921;status=T;Dbxref="InterPro:IPR000680" |
| gene-bpuSUM 001777 | lp46 | gene | delta | Pfam | protein matc | 30 | 315 | 7E-85 | + | . | Target=null 30 315;ID=match$1685_30_315;signature_desc=Borrelia lipoprotein;Name=PF00921;status=T;Dbxref="InterPro:IPR000680" |
| gene-bpuSUM 001779 | lp46 | gene | delta | Pfam | protein matc | 45 | 329 | 3.6E-85 | + | . | Target=null 45 329;ID=match$104_45_329;signature_desc=Borrelia lipoprotein;Name=PF00921;status=T;Dbxref="InterPro:IPR000680" |
| gene-bpuSUM 001780 | lp46 | gene | delta | Pfam | protein matc | 19 | 309 | 4.5E-88 | + | . | Target=null 19 309;ID=match$1596_19_309;signature_desc=Borrelia lipoprotein;Name=PF00921;status=T;Dbxref="InterPro:IPR000680" |
| gene-bpuSUM 001782 | lp46 | gene | delta | Pfam | protein matc | 45 | 323 | 2.4E-76 | + | . | Target=null 45 323;ID=match$3100_45_323;signature_desc=Borrelia lipoprotein;Name=PF00921;status=T;Dbxref="InterPro:IPR000680" |
| gene-bpuSUM 001783 | lp46 | pseudo | gamma | Pfam | protein matc | 34 | 191 | 6.4E-43 | + | . | Target=null 34 191;ID=match$1584_34_191;signature_desc=Borrelia lipoprotein;Name=PF00921;status=T;Dbxref="InterPro:IPR000680" |
| gene-bpuSUM 001783 | lp46 | pseudo | | Pfam | protein matc | 3 | 133 | 2.8E-38 | + | . | Target=null 3 133;ID=match$1439_3_133;signature_desc=Borrelia lipoprotein;Name=PF00921;status=T;Dbxref="InterPro:IPR000680" |
| gene-bpuSUM 001785 | lp46 | gene | beta | Pfam | protein matc | 20 | 309 | 1.9E-42 | + | . | Target=null 20 309;ID=match$2954_20_309;signature_desc=Borrelia lipoprotein;Name=PF00921;status=T;Dbxref="InterPro:IPR000680" |
| gene-bpuSUM 001786 | lp46 | gene | alpha | Pfam | protein matc | 7 | 302 | 9.9E-69 | + | . | Target=null 7 302;ID=match$1453_7_302;signature_desc=Borrelia lipoprotein;Name=PF00921;status=T;Dbxref="InterPro:IPR000680" |
| gene-bpuSUM 001790 | lp46 | gene | delta | Pfam | protein matc | 45 | 327 | 6.8E-86 | + | . | Target=null 45 327;ID=match$2127_45_327;signature_desc=Borrelia lipoprotein;Name=PF00921;status=T;Dbxref="InterPro:IPR000680" |
| gene-bpuSUM 001791 | lp46 | gene | beta | Pfam | protein matc | 34 | 321 | 2.1E-43 | + | . | Target=null 34 321;ID=match$1084_34_321;signature_desc=Borrelia lipoprotein;Name=PF00921;status=T;Dbxref="InterPro:IPR000680" |
| gene-bpuSUM 001792 | lp46 | gene | alpha | Pfam | protein matc | 49 | 345 | 9.2E-72 | + | . | Target=null 49 345;ID=match$1442_49_345;signature_desc=Borrelia lipoprotein;Name=PF00921;status=T;Dbxref="InterPro:IPR000680" |
| gene-bpuSUM 001793 | lp46 | gene | beta | Pfam | protein matc | 56 | 344 | 3.8E-42 | + | . | Target=null 56 344;ID=match$1277_56_344;signature_desc=Borrelia lipoprotein;Name=PF00921;status=T;Dbxref="InterPro:IPR000680" |
| gene-bpuSUM 001794 | lp46 | pseudo | beta | Pfam | protein matc | 19 | 61 | 0.0000031 | + | . | Target=null 19 61;ID=match$933_19_61;signature_desc=Borrelia lipoprotein;Name=PF00921;status=T;Dbxref="InterPro:IPR000680" |
| gene-bpuSUM 001794 | lp46 | pseudo | | Pfam | protein matc | 11 | 197 | 1.1E-31 | + | . | Target=null 11 197;ID=match$2499_11_197;signature_desc=Borrelia lipoprotein;Name=PF00921;status=T;Dbxref="InterPro:IPR000680" |
| gene-bpuSUM 001798 | lp46 | pseudo | gamma | Pfam | protein matc | 51 | 100 | 3.7E-08 | + | . | Target=null 51 100;ID=match$1796_51_100;signature_desc=Borrelia lipoprotein;Name=PF00921;status=T;Dbxref="InterPro:IPR000680" |
| gene-bpuSUM 001800 | lp46 | gene | alpha | Pfam | protein matc | 7 | 301 | 1.4E-71 | + | . | Target=null 7 301;ID=match$2463_7_301;signature_desc=Borrelia lipoprotein;Name=PF00921;status=T;Dbxref="InterPro:IPR000680" |
| gene-bpuSUM 001803 | lp46 | pseudo | gamma | Pfam | protein matc | 45 | 178 | 1E-43 | + | . | Target=null 45 178;ID=match$57_45_178;signature_desc=Borrelia lipoprotein;Name=PF00921;status=T;Dbxref="InterPro:IPR000680" |
| gene-bpuSUM 001803 | lp46 | pseudo | | Pfam | protein matc | 13 | 161 | 1.9E-35 | + | . | Target=null 13 161;ID=match$3110_13_161;signature_desc=Borrelia lipoprotein;Name=PF00921;status=T;Dbxref="InterPro:IPR000680" |
| gene-bpuSUM 001804 | lp46 | gene | beta | Pfam | protein matc | 45 | 331 | 2E-47 | + | . | Target=null 45 331;ID=match$2111_45_331;signature_desc=Borrelia lipoprotein;Name=PF00921;status=T;Dbxref="InterPro:IPR000680" |
| gene-bpuSUM 001806 | lp46 | gene | delta | Pfam | protein matc | 1 | 61 | 1.1E-17 | + | . | Target=null 1 61;ID=match$697_1_61;signature_desc=Borrelia lipoprotein;Name=PF00921;status=T;Dbxref="InterPro:IPR000680" |
| gene-bpuSUM 001812 | lp46 | gene | delta | Pfam | protein matc | 8 | 42 | 0.0000034 | + | . | Target=null 8 42;ID=match$3148_8_42;signature_desc=Borrelia lipoprotein;Name=PF00921;status=T;Dbxref="InterPro:IPR000680" |
| gene-bpuSUM 001841 | lp47 | gene | alpha | Pfam | protein matc | 7 | 302 | 9.9E-69 | + | . | Target=null 7 302;ID=match$1451_7_302;signature_desc=Borrelia lipoprotein;Name=PF00921;status=T;Dbxref="InterPro:IPR000680" |
| gene-bpuSUM 001842 | lp47 | gene | beta | Pfam | protein matc | 20 | 309 | 1.9E-42 | + | . | Target=null 20 309;ID=match$2956_20_309;signature_desc=Borrelia lipoprotein;Name=PF00921;status=T;Dbxref="InterPro:IPR000680" |
| gene-bpuSUM 001844 | lp47 | pseudo | gamma | Pfam | protein matc | 34 | 191 | 6.4E-43 | + | . | Target=null 34 191;ID=match$1586_34_191;signature_desc=Borrelia lipoprotein;Name=PF00921;status=T;Dbxref="InterPro:IPR000680" |
| gene-bpuSUM 001844 | lp47 | pseudo | | Pfam | protein matc | 3 | 133 | 2.8E-38 | + | . | Target=null 3 133;ID=match$1441_3_133;signature_desc=Borrelia lipoprotein;Name=PF00921;status=T;Dbxref="InterPro:IPR000680" |
| gene-bpuSUM 001845 | lp47 | gene | delta | Pfam | protein matc | 45 | 323 | 2.4E-76 | + | . | Target=null 45 323;ID=match$3098_45_323;signature_desc=Borrelia lipoprotein;Name=PF00921;status=T;Dbxref="InterPro:IPR000680" |
| gene-bpuSUM 001847 | lp47 | gene | delta | Pfam | protein matc | 19 | 309 | 4.5E-88 | + | . | Target=null 19 309;ID=match$1598_19_309;signature_desc=Borrelia lipoprotein;Name=PF00921;status=T;Dbxref="InterPro:IPR000680" |
| gene-bpuSUM 001848 | lp47 | gene | delta | Pfam | protein matc | 45 | 329 | 3.6E-85 | + | . | Target=null 45 329;ID=match$106_45_329;signature_desc=Borrelia lipoprotein;Name=PF00921;status=T;Dbxref="InterPro:IPR000680" |
| gene-bpuSUM 001850 | lp47 | gene | delta | Pfam | protein matc | 30 | 315 | 7E-85 | + | . | Target=null 30 315;ID=match$1691_30_315;signature_desc=Borrelia lipoprotein;Name=PF00921;status=T;Dbxref="InterPro:IPR000680" |
| gene-bpuSUM 001851 | lp47 | gene | gamma | Pfam | protein matc | 44 | 330 | 4.6E-91 | + | . | Target=null 44 330;ID=match$3184_44_330;signature_desc=Borrelia lipoprotein;Name=PF00921;status=T;Dbxref="InterPro:IPR000680" |
| gene-bpuSUM 001852 | lp47 | pseudo | alpha | Pfam | protein matc | 53 | 335 | 7E-69 | + | . | Target=null 53 335;ID=match$1215_53_335;signature_desc=Borrelia lipoprotein;Name=PF00921;status=T;Dbxref="InterPro:IPR000680" |
| gene-bpuSUM 001859 | lp57 | pseudo | alpha | Pfam | protein matc | 53 | 335 | 7E-69 | + | . | Target=null 53 335;ID=match$1217_53_335;signature_desc=Borrelia lipoprotein;Name=PF00921;status=T;Dbxref="InterPro:IPR000680" |
| gene-bpuSUM 001860 | lp57 | gene | gamma | Pfam | protein matc | 44 | 330 | 4.6E-91 | + | . | Target=null 44 330;ID=match$3182_44_330;signature_desc=Borrelia lipoprotein;Name=PF00921;status=T;Dbxref="InterPro:IPR000680" |
| gene-bpuSUM 001861 | lp57 | gene | delta | Pfam | protein matc | 30 | 315 | 7E-85 | + | . | Target=null 30 315;ID=match$1689_30_315;signature_desc=Borrelia lipoprotein;Name=PF00921;status=T;Dbxref="InterPro:IPR000680" |
| gene-bpuSUM 001863 | lp57 | gene | delta | Pfam | protein matc | 19 | 303 | 1.4E-87 | + | . | Target=null 19 303;ID=match$2287_19_303;signature_desc=Borrelia lipoprotein;Name=PF00921;status=T;Dbxref="InterPro:IPR000680" |
| gene-bpuSUM 001865 | lp57 | gene | beta | Pfam | protein matc | 61 | 349 | 2E-45 | + | . | Target=null 61 349;ID=match$635_61_349;signature_desc=Borrelia lipoprotein;Name=PF00921;status=T;Dbxref="InterPro:IPR000680" |
| gene-bpuSUM 001866 | lp57 | gene | beta | Pfam | protein matc | 5 | 294 | 3.8E-41 | + | . | Target=null 5 294;ID=match$1968_5_294;signature_desc=Borrelia lipoprotein;Name=PF00921;status=T;Dbxref="InterPro:IPR000680" |
| gene-bpuSUM 001867 | lp57 | gene | alpha | Pfam | protein matc | 22 | 311 | 1E-74 | + | . | Target=null 22 311;ID=match$331_22_311;signature_desc=Borrelia lipoprotein;Name=PF00921;status=T;Dbxref="InterPro:IPR000680" |
| gene-bpuSUM 001868 | lp57 | gene | gamma | Pfam | protein matc | 44 | 326 | 3.5E-91 | + | . | Target=null 44 326;ID=match$1104_44_326;signature_desc=Borrelia lipoprotein;Name=PF00921;status=T;Dbxref="InterPro:IPR000680" |
| gene-bpuSUM 001871 | lp57 | gene | gamma | Pfam | protein matc | 44 | 323 | 6.8E-87 | + | . | Target=null 44 323;ID=match$2188_44_323;signature_desc=Borrelia lipoprotein;Name=PF00921;status=T;Dbxref="InterPro:IPR000680" |
| gene-bpuSUM 001872 | lp57 | pseudo | delta | Pfam | protein matc | 19 | 167 | 1.5E-38 | + | . | Target=null 19 167;ID=match$2238_19_167;signature_desc=Borrelia lipoprotein;Name=PF00921;status=T;Dbxref="InterPro:IPR000680" |
| gene-bpuSUM 001872 | lp57 | pseudo | | Pfam | protein matc | 2 | 114 | 1.4E-30 | + | . | Target=null 2 114;ID=match$2893_2_114;signature_desc=Borrelia lipoprotein;Name=PF00921;status=T;Dbxref="InterPro:IPR000680" |
| gene-bpuSUM 001874 | lp57 | gene | delta | Pfam | protein matc | 45 | 324 | 8.7E-87 | + | . | Target=null 45 324;ID=match$1230_45_324;signature_desc=Borrelia lipoprotein;Name=PF00921;status=T;Dbxref="InterPro:IPR000680" |
| gene-bpuSUM 001876 | lp57 | gene | delta | Pfam | protein matc | 34 | 315 | 1.5E-84 | + | . | Target=null 34 315;ID=match$1654_34_315;signature_desc=Borrelia lipoprotein;Name=PF00921;status=T;Dbxref="InterPro:IPR000680" |
| gene-bpuSUM 001890 | lp57 | gene | gamma | Pfam | protein matc | 19 | 295 | 3.2E-83 | + | . | Target=null 19 295;ID=match$2896_19_295;signature_desc=Borrelia lipoprotein;Name=PF00921;status=T;Dbxref="InterPro:IPR000680" |
| gene-bpuSUM 001891 | lp57 | gene | gamma | Pfam | protein matc | 38 | 328 | 3.6E-92 | + | . | Target=null 38 328;ID=match$3420_38_328;signature_desc=Borrelia lipoprotein;Name=PF00921;status=T;Dbxref="InterPro:IPR000680" |
| gene-bpuSUM 001892 | lp57 | gene | delta | Pfam | protein matc | 19 | 292 | 3.5E-80 | + | . | Target=null 19 292;ID=match$3472_19_292;signature_desc=Borrelia lipoprotein;Name=PF00921;status=T;Dbxref="InterPro:IPR000680" |
| gene-bpuSUM 001893 | lp57 | gene | beta | Pfam | protein matc | 45 | 333 | 3E-44 | + | . | Target=null 45 333;ID=match$508_45_333;signature_desc=Borrelia lipoprotein;Name=PF00921;status=T;Dbxref="InterPro:IPR000680" |
| gene-bpuSUM 001895 | lp57 | pseudo | gamma | Pfam | protein matc | 1 | 268 | 3.9E-78 | + | . | Target=null 1 268;ID=match$2469_1_268;signature_desc=Borrelia lipoprotein;Name=PF00921;status=T;Dbxref="InterPro:IPR000680" |
| gene-bpuSUM 001897 | lp57 | gene | alpha | Pfam | protein matc | 7 | 300 | 1.7E-70 | + | . | Target=null 7 300;ID=match$1488_7_300;signature_desc=Borrelia lipoprotein;Name=PF00921;status=T;Dbxref="InterPro:IPR000680" |
| gene-bpuSUM 001898 | lp57 | gene | delta | Pfam | protein matc | 34 | 317 | 4.7E-85 | + | . | Target=null 34 317;ID=match$2446_34_317;signature_desc=Borrelia lipoprotein;Name=PF00921;status=T;Dbxref="InterPro:IPR000680" |
| gene-bpuSUM 001900 | lp57 | gene | alpha | Pfam | protein matc | 7 | 301 | 5E-67 | + | . | Target=null 7 301;ID=match$385_7_301;signature_desc=Borrelia lipoprotein;Name=PF00921;status=T;Dbxref="InterPro:IPR000680" |
| gene-bpuSUM 001901 | lp57 | gene | beta | Pfam | protein matc | 35 | 321 | 4.5E-49 | + | . | Target=null 35 321;ID=match$3657_35_321;signature_desc=Borrelia lipoprotein;Name=PF00921;status=T;Dbxref="InterPro:IPR000680" |
| gene-bpuSUM 001902 | lp57 | gene | alpha | Pfam | protein matc | 42 | 337 | 2.2E-65 | + | . | Target=null 42 337;ID=match$1328_42_337;signature_desc=Borrelia lipoprotein;Name=PF00921;status=T;Dbxref="InterPro:IPR000680" |
| gene-bpuSUM 001903 | lp57 | gene | beta | Pfam | protein matc | 20 | 307 | 1.2E-51 | + | . | Target=null 20 307;ID=match$3222_20_307;signature_desc=Borrelia lipoprotein;Name=PF00921;status=T;Dbxref="InterPro:IPR000680" |
| gene-bpuSUM 001904 | lp57 | gene | gamma | Pfam | protein matc | 34 | 308 | 1.9E-81 | + | . | Target=null 34 308;ID=match$2448_34_308;signature_desc=Borrelia lipoprotein;Name=PF00921;status=T;Dbxref="InterPro:IPR000680" |
| gene-bpuSUM 001906 | lp57 | gene | delta | Pfam | protein matc | 33 | 306 | 5E-85 | + | . | Target=null 33 306;ID=match$3190_33_306;signature_desc=Borrelia lipoprotein;Name=PF00921;status=T;Dbxref="InterPro:IPR000680" |
| gene-bpuSUM 001909 | lp57 | gene | gamma | Pfam | protein matc | 44 | 330 | 4.6E-91 | + | . | Target=null 44 330;ID=match$3176_44_330;signature_desc=Borrelia lipoprotein;Name=PF00921;status=T;Dbxref="InterPro:IPR000680" |
| gene-bpuSUM 001910 | lp57 | pseudo | alpha | Pfam | protein matc | 53 | 107 | 9.8E-13 | + | . | Target=null 53 107;ID=match$21_53_107;signature_desc=Borrelia lipoprotein;Name=PF00921;status=T;Dbxref="InterPro:IPR000680" |
| gene-bpuSUM 001910 | lp57 | pseudo | | Pfam | protein matc | 11 | 245 | 5.1E-51 | + | . | Target=null 11 245;ID=match$2699_11_245;signature_desc=Borrelia lipoprotein;Name=PF00921;status=T;Dbxref="InterPro:IPR000680" |
| gene-bpuSUM 001917 | lp65 | pseudo | gamma | Pfam | protein matc | 35 | 223 | 1.3E-54 | + | . | Target=null 35 223;ID=match$2864_35_223;signature_desc=Borrelia lipoprotein;Name=PF00921;status=T;Dbxref="InterPro:IPR000680" |
| gene-bpuSUM 001917 | lp65 | pseudo | | Pfam | protein matc | 8 | 89 | 2.3E-27 | + | . | Target=null 8 89;ID=match$253_8_89;signature_desc=Borrelia lipoprotein;Name=PF00921;status=T;Dbxref="InterPro:IPR000680" |
| gene-bpuSUM 001919 | lp65 | gene | delta | Pfam | protein matc | 45 | 326 | 1.4E-88 | + | . | Target=null 45 326;ID=match$3067_45_326;signature_desc=Borrelia lipoprotein;Name=PF00921;status=T;Dbxref="InterPro:IPR000680" |
| gene-bpuSUM 001922 | lp65 | gene | delta | Pfam | protein matc | 45 | 325 | 1.1E-82 | + | . | Target=null 45 325;ID=match$2992_45_325;signature_desc=Borrelia lipoprotein;Name=PF00921;status=T;Dbxref="InterPro:IPR000680" |
| gene-bpuSUM 001923 | lp65 | gene | alpha | Pfam | protein matc | 39 | 328 | 5.5E-68 | + | . | Target=null 39 328;ID=match$849_39_328;signature_desc=Borrelia lipoprotein;Name=PF00921;status=T;Dbxref="InterPro:IPR000680" |
| gene-bpuSUM 001924 | lp65 | gene | delta | Pfam | protein matc | 44 | 333 | 1.6E-86 | + | . | Target=null 44 333;ID=match$3143_44_333;signature_desc=Borrelia lipoprotein;Name=PF00921;status=T;Dbxref="InterPro:IPR000680" |
| gene-bpuSUM 001926 | lp65 | pseudo | delta | Pfam | protein matc | 45 | 129 | 2.7E-26 | + | . | Target=null 45 129;ID=match$2847_45_129;signature_desc=Borrelia lipoprotein;Name=PF00921;status=T;Dbxref="InterPro:IPR000680" |
| gene-bpuSUM 001926 | lp65 | pseudo | | Pfam | protein matc | 16 | 209 | 1.3E-55 | + | . | Target=null 16 209;ID=match$517_16_209;signature_desc=Borrelia lipoprotein;Name=PF00921;status=T;Dbxref="InterPro:IPR000680" |
| gene-bpuSUM 001927 | lp65 | gene | beta | Pfam | protein matc | 45 | 328 | 5.1E-48 | + | . | Target=null 45 328;ID=match$3378_45_328;signature_desc=Borrelia lipoprotein;Name=PF00921;status=T;Dbxref="InterPro:IPR000680" |
| gene-bpuSUM 001928 | lp65 | pseudo | beta | Pfam | protein matc | 20 | 107 | 1.5E-09 | + | . | Target=null 20 107;ID=match$514_20_107;signature_desc=Borrelia lipoprotein;Name=PF00921;status=T;Dbxref="InterPro:IPR000680" |
| gene-bpuSUM 001929 | lp65 | pseudo | | Pfam | protein matc | 7 | 85 | 8.1E-16 | + | . | Target=null 7 85;ID=match$274_7_85;signature_desc=Borrelia lipoprotein;Name=PF00921;status=T;Dbxref="InterPro:IPR000680" |
| gene-bpuSUM 001929 | lp65 | pseudo | alpha | Pfam | protein matc | 6 | 206 | 4.5E-47 | + | . | Target=null 6 206;ID=match$2407_6_206;signature_desc=Borrelia lipoprotein;Name=PF00921;status=T;Dbxref="InterPro:IPR000680" |
| gene-bpuSUM 001931 | lp65 | pseudo | beta | Pfam | protein matc | 16 | 150 | 3.4E-19 | + | . | Target=null 16 150;ID=match$646_16_150;signature_desc=Borrelia lipoprotein;Name=PF00921;status=T;Dbxref="InterPro:IPR000680" |
| gene-bpuSUM 001932 | lp65 | gene | alpha | Pfam | protein matc | 49 | 345 | 6.6E-71 | + | . | Target=null 49 345;ID=match$2495_49_345;signature_desc=Borrelia lipoprotein;Name=PF00921;status=T;Dbxref="InterPro:IPR000680" |
| gene-bpuSUM 001934 | lp65 | gene | delta | Pfam | protein matc | 30 | 316 | 1.4E-89 | + | . | Target=null 30 316;ID=match$2135_30_316;signature_desc=Borrelia lipoprotein;Name=PF00921;status=T;Dbxref="InterPro:IPR000680" |
| gene-bpuSUM 001936 | lp65 | pseudo | gamma | Pfam | protein matc | 21 | 233 | 4.9E-59 | + | . | Target=null 21 233;ID=match$1790_21_233;signature_desc=Borrelia lipoprotein;Name=PF00921;status=T;Dbxref="InterPro:IPR000680" |
| gene-bpuSUM 001936 | lp65 | pseudo | | Pfam | protein matc | 24 | 87 | 2E-24 | + | . | Target=null 24 87;ID=match$2168_24_87;signature_desc=Borrelia lipoprotein;Name=PF00921;status=T;Dbxref="InterPro:IPR000680" |
| gene-bpuSUM 001938 | lp65 | gene | delta | Pfam | protein matc | 8 | 298 | 1.6E-83 | + | . | Target=null 8 298;ID=match$1261_8_298;signature_desc=Borrelia lipoprotein;Name=PF00921;status=T;Dbxref="InterPro:IPR000680" |
| gene-bpuSUM 001953 | lp65 | gene | delta | Pfam | protein matc | 42 | 326 | 1.6E-83 | + | . | Target=null 42 326;ID=match$2827_42_326;signature_desc=Borrelia lipoprotein;Name=PF00921;status=T;Dbxref="InterPro:IPR000680" |
| gene-bpuSUM 001957 | lp65 | pseudo | beta | Pfam | protein matc | 19 | 61 | 0.0000031 | + | . | Target=null 19 61;ID=match$931_19_61;signature_desc=Borrelia lipoprotein;Name=PF00921;status=T;Dbxref="InterPro:IPR000680" |
| gene-bpuSUM 001957 | lp65 | pseudo | | Pfam | protein matc | 11 | 197 | 1.1E-31 | + | . | Target=null 11 197;ID=match$2497_11_197;signature_desc=Borrelia lipoprotein;Name=PF00921;status=T;Dbxref="InterPro:IPR000680" |
| gene-bpuSUM 001958 | lp65 | gene | beta | Pfam | protein matc | 56 | 344 | 3.8E-42 | + | . | Target=null 56 344;ID=match$1279_56_344;signature_desc=Borrelia lipoprotein;Name=PF00921;status=T;Dbxref="InterPro:IPR000680" |
| gene-bpuSUM 001959 | lp65 | gene | alpha | Pfam | protein matc | 49 | 345 | 9.2E-72 | + | . | Target=null 49 345;ID=match$1444_49_345;signature_desc=Borrelia lipoprotein;Name=PF00921;status=T;Dbxref="InterPro:IPR000680" |
| gene-bpuSUM 001960 | lp65 | gene | beta | Pfam | protein matc | 37 | 324 | 2.2E-43 | + | . | Target=null 37 324;ID=match$3311_37_324;signature_desc=Borrelia lipoprotein;Name=PF00921;status=T;Dbxref="InterPro:IPR000680" |
| gene-bpuSUM 001961 | lp65 | gene | delta | Pfam | protein matc | 45 | 327 | 6.8E-86 | + | . | Target=null 45 327;ID=match$2129_45_327;signature_desc=Borrelia lipoprotein;Name=PF00921;status=T;Dbxref="InterPro:IPR000680" |
| gene-bpuSUM 001965 | lp65 | gene | alpha | Pfam | protein matc | 7 | 302 | 9.9E-69 | + | . | Target=null 7 302;ID=match$1449_7_302;signature_desc=Borrelia lipoprotein;Name=PF00921;status=T;Dbxref="InterPro:IPR000680" |
| gene-bpuSUM 001966 | lp65 | gene | beta | Pfam | protein matc | 20 | 309 | 1.9E-42 | + | . | Target=null 20 309;ID=match$2958_20_309;signature_desc=Borrelia lipoprotein;Name=PF00921;status=T;Dbxref="InterPro:IPR000680" |
| gene-bpuSUM 001968 | lp65 | pseudo | gamma | Pfam | protein matc | 34 | 191 | 6.4E-43 | + | . | Target=null 34 191;ID=match$1588_34_191;signature_desc=Borrelia lipoprotein;Name=PF00921;status=T;Dbxref="InterPro:IPR000680" |
| gene-bpuSUM 001968 | lp65 | pseudo | | Pfam | protein matc | 3 | 133 | 2.8E-38 | + | . | Target=null 3 133;ID=match$1437_3_133;signature_desc=Borrelia lipoprotein;Name=PF00921;status=T;Dbxref="InterPro:IPR000680" |
| gene-bpuSUM 001969 | lp65 | gene | delta | Pfam | protein matc | 45 | 323 | 2.4E-76 | + | . | Target=null 45 323;ID=match$3102_45_323;signature_desc=Borrelia lipoprotein;Name=PF00921;status=T;Dbxref="InterPro:IPR000680" |
| gene-bpuSUM 001971 | lp65 | gene | delta | Pfam | protein matc | 19 | 309 | 4.5E-88 | + | . | Target=null 19 309;ID=match$1594_19_309;signature_desc=Borrelia lipoprotein;Name=PF00921;status=T;Dbxref="InterPro:IPR000680" |
| gene-bpuSUM 001972 | lp65 | gene | delta | Pfam | protein matc | 45 | 329 | 3.6E-85 | + | . | Target=null 45 329;ID=match$108_45_329;signature_desc=Borrelia lipoprotein;Name=PF00921;status=T;Dbxref="InterPro:IPR000680" |
| gene-bpuSUM 001974 | lp65 | gene | delta | Pfam | protein matc | 30 | 315 | 7E-85 | + | . | Target=null 30 315;ID=match$1687_30_315;signature_desc=Borrelia lipoprotein;Name=PF00921;status=T;Dbxref="InterPro:IPR000680" |
| gene-bpuSUM 001975 | lp65 | gene | gamma | Pfam | protein matc | 44 | 330 | 4.6E-91 | + | . | Target=null 44 330;ID=match$3178_44_330;signature_desc=Borrelia lipoprotein;Name=PF00921;status=T;Dbxref="InterPro:IPR000680" |
| gene-bpuSUM 001976 | lp65 | pseudo | alpha | Pfam | protein matc | 53 | 335 | 7E-69 | + | . | Target=null 53 335;ID=match$1213_53_335;signature_desc=Borrelia lipoprotein;Name=PF00921;status=T;Dbxref="InterPro:IPR000680" |

| Vsp | | replicon | gene/pseudo | | | | | | | | | | |
|---|---|---|---|---|---|---|---|---|---|---|---|---|---|
| gene-bpSLO | 001244 | lp28 | gene | Pfam | | protein | matc | 39 | 206 | 1.7E-67 | + | . | Target=null 39 206;ID=match$2082_39_206;signature_desc=Lipoprotein;Name=PF01441;status=T;Dbxref="InterPro:IPR001800" |
| gene-bpSLO | 001281 | Lp30-1 | gene | Pfam | | protein | matc | 29 | 195 | 7.8E-56 | + | . | Target=null 29 195;ID=match$1518_29_195;signature_desc=Lipoprotein;Name=PF01441;status=T;Dbxref="InterPro:IPR001800" |
| gene-bpSLO | 001340 | lp32 | gene | Pfam | | protein | matc | 40 | 205 | 3E-63 | + | . | Target=null 40 205;ID=match$1120_40_205;signature_desc=Lipoprotein;Name=PF01441;status=T;Dbxref="InterPro:IPR001800" |
| gene-bpSLO | 001360 | Lp34-1 | gene | Pfam | | protein | matc | 40 | 209 | 1.4E-58 | + | . | Target=null 40 209;ID=match$2441_40_209;signature_desc=Lipoprotein;Name=PF01441;status=T;Dbxref="InterPro:IPR001800" |
| gene-bpSLO | 001374 | Lp34-1 | gene | Pfam | | protein | matc | 39 | 205 | 1.6E-58 | + | . | Target=null 39 205;ID=match$1090_39_205;signature_desc=Lipoprotein;Name=PF01441;status=T;Dbxref="InterPro:IPR001800" |
| gene-bpSLO | 001378 | Lp34-1 | gene | Pfam | | protein | matc | 40 | 208 | 3.7E-56 | + | . | Target=null 40 208;ID=match$1034_40_208;signature_desc=Lipoprotein;Name=PF01441;status=T;Dbxref="InterPro:IPR001800" |
| gene-bpSLO | 001412 | Lp34-2 | gene | Pfam | | protein | matc | 40 | 209 | 1.4E-58 | + | . | Target=null 40 209;ID=match$2445_40_209;signature_desc=Lipoprotein;Name=PF01441;status=T;Dbxref="InterPro:IPR001800" |
| gene-bpSLO | 001429 | lp42 | gene | Pfam | | protein | matc | 39 | 205 | 9.9E-58 | + | . | Target=null 39 205;ID=match$1133_39_205;signature_desc=Lipoprotein;Name=PF01441;status=T;Dbxref="InterPro:IPR001800" |
| gene-bpSLO | 001434 | lp42 | gene | Pfam | | protein | matc | 40 | 207 | 2.3E-69 | + | . | Target=null 40 207;ID=match$1707_40_207;signature_desc=Lipoprotein;Name=PF01441;status=T;Dbxref="InterPro:IPR001800" |
| gene-bpSLO | 001438 | lp42 | gene | Pfam | | protein | matc | 40 | 208 | 1.3E-54 | + | . | Target=null 40 208;ID=match$1082_40_208;signature_desc=Lipoprotein;Name=PF01441;status=T;Dbxref="InterPro:IPR001800" |
| gene-bpSLO | 001475 | lp58 | gene | Pfam | | protein | matc | 39 | 207 | 2.8E-64 | + | . | Target=null 39 207;ID=match$1030_39_207;signature_desc=Lipoprotein;Name=PF01441;status=T;Dbxref="InterPro:IPR001800" |
| gene-bpSLO | 001486 | lp58 | gene | Pfam | | protein | matc | 39 | 206 | 7.7E-65 | + | . | Target=null 39 206;ID=match$2188_39_206;signature_desc=Lipoprotein;Name=PF01441;status=T;Dbxref="InterPro:IPR001800" |
| gene-bpSLO | 001512 | lp58 | gene | Pfam | | protein | matc | 40 | 204 | 5.9E-51 | + | . | Target=null 40 204;ID=match$803_40_204;signature_desc=Lipoprotein;Name=PF01441;status=T;Dbxref="InterPro:IPR001800" |
| gene-bpSLO | 001530 | lp60 | gene | Pfam | | protein | matc | 40 | 209 | 1.4E-58 | + | . | Target=null 40 209;ID=match$2443_40_209;signature_desc=Lipoprotein;Name=PF01441;status=T;Dbxref="InterPro:IPR001800" |
| gene-bpSLO | 001533 | lp60 | gene | Pfam | | protein | matc | 39 | 205 | 3.9E-49 | + | . | Target=null 39 205;ID=match$2012_39_205;signature_desc=Lipoprotein;Name=PF01441;status=T;Dbxref="InterPro:IPR001800" |
| gene-bpSLO | 001538 | lp60 | gene | Pfam | | protein | matc | 40 | 208 | 7.8E-62 | + | . | Target=null 40 208;ID=match$2837_40_208;signature_desc=Lipoprotein;Name=PF01441;status=T;Dbxref="InterPro:IPR001800" |
| gene-bpSLO | 001541 | lp60 | gene | Pfam | | protein | matc | 40 | 205 | 4.4E-58 | + | . | Target=null 40 205;ID=match$1305_40_205;signature_desc=Lipoprotein;Name=PF01441;status=T;Dbxref="InterPro:IPR001800" |
| gene-bpSLO | 001546 | lp60 | gene | Pfam | | protein | matc | 38 | 206 | 3.9E-64 | + | . | Target=null 38 206;ID=match$1640_38_206;signature_desc=Lipoprotein;Name=PF01441;status=T;Dbxref="InterPro:IPR001800" |
| gene-bpSLO | 001567 | lp60 | gene | Pfam | | protein | matc | 23 | 192 | 6.1E-52 | + | . | Target=null 23 192;ID=match$383_23_192;signature_desc=Lipoprotein;Name=PF01441;status=T;Dbxref="InterPro:IPR001800" |
| gene-bpSLO | 001569 | lp60 | gene | Pfam | | protein | matc | 39 | 206 | 2E-64 | + | . | Target=null 39 206;ID=match$725_39_206;signature_desc=Lipoprotein;Name=PF01441;status=T;Dbxref="InterPro:IPR001800" |
| gene-bpSLO | 001579 | lp60 | gene | Pfam | | protein | matc | 39 | 211 | 1.6E-46 | + | . | Target=null 39 211;ID=match$2817_39_211;signature_desc=Lipoprotein;Name=PF01441;status=T;Dbxref="InterPro:IPR001800" |
| gene-bpSLO | 001585 | lp60 | gene | Pfam | | protein | matc | 39 | 207 | 1.8E-54 | + | . | Target=null 39 207;ID=match$1058_39_207;signature_desc=Lipoprotein;Name=PF01441;status=T;Dbxref="InterPro:IPR001800" |
| Vlp | | | | | Subfamily | | | | | | | | |
| gene-bpSLO | 001190 | lp18 | pseudo | Pfam | beta | protein | matc | 7 | 292 | 2.8E-46 | + | . | Target=null 7 292;ID=match$1742_7_292;signature_desc=Borrelia lipoprotein;Name=PF00921;status=T;Dbxref="InterPro:IPR000680" |
| gene-bpSLO | 001191 | lp18 | gene | Pfam | beta | protein | matc | 48 | 331 | 3.3E-45 | + | . | Target=null 48 331;ID=match$1797_48_331;signature_desc=Borrelia lipoprotein;Name=PF00921;status=T;Dbxref="InterPro:IPR000680" |
| gene-bpSLO | 001204 | lp18 | gene | Pfam | gamma | protein | matc | 1 | 278 | 4.2E-83 | + | . | Target=null 1 278;ID=match$1146_1_278;signature_desc=Borrelia lipoprotein;Name=PF00921;status=T;Dbxref="InterPro:IPR000680" |
| gene-bpSLO | 001209 | lp24 | gene | Pfam | alpha | protein | matc | 51 | 334 | 1.3E-64 | + | . | Target=null 51 334;ID=match$2455_51_334;signature_desc=Borrelia lipoprotein;Name=PF00921;status=T;Dbxref="InterPro:IPR000680" |
| gene-bpSLO | 001211 | lp24 | gene | Pfam | beta | protein | matc | 48 | 338 | 2.5E-48 | + | . | Target=null 48 338;ID=match$2797_48_338;signature_desc=Borrelia lipoprotein;Name=PF00921;status=T;Dbxref="InterPro:IPR000680" |
| gene-bpSLO | 001213 | lp24 | gene | Pfam | gamma | protein | matc | 34 | 318 | 2.6E-86 | + | . | Target=null 34 318;ID=match$2297_34_318;signature_desc=Borrelia lipoprotein;Name=PF00921;status=T;Dbxref="InterPro:IPR000680" |
| gene-bpSLO | 001214 | lp24 | gene | Pfam | delta | protein | matc | 45 | 325 | 1.9E-83 | + | . | Target=null 45 325;ID=match$2027_45_325;signature_desc=Borrelia lipoprotein;Name=PF00921;status=T;Dbxref="InterPro:IPR000680" |
| gene-bpSLO | 001215 | lp24 | gene | Pfam | delta | protein | matc | 34 | 317 | 1.9E-85 | + | . | Target=null 34 317;ID=match$1323_34_317;signature_desc=Borrelia lipoprotein;Name=PF00921;status=T;Dbxref="InterPro:IPR000680" |
| gene-bpSLO | 001216 | lp24 | gene | Pfam | delta | protein | matc | 44 | 332 | 1.5E-81 | + | . | Target=null 44 332;ID=match$2626_44_332;signature_desc=Borrelia lipoprotein;Name=PF00921;status=T;Dbxref="InterPro:IPR000680" |
| gene-bpSLO | 001217 | lp24 | gene | Pfam | delta | protein | matc | 44 | 329 | 1.6E-78 | + | . | Target=null 44 329;ID=match$2972_44_329;signature_desc=Borrelia lipoprotein;Name=PF00921;status=T;Dbxref="InterPro:IPR000680" |
| gene-bpSLO | 001218 | lp24 | gene | Pfam | gamma | protein | matc | 34 | 330 | 2.8E-86 | + | . | Target=null 34 330;ID=match$1174_34_330;signature_desc=Borrelia lipoprotein;Name=PF00921;status=T;Dbxref="InterPro:IPR000680" |
| gene-bpSLO | 001219 | lp24 | gene | Pfam | delta | protein | matc | 45 | 326 | 5E-83 | + | . | Target=null 45 326;ID=match$995_45_326;signature_desc=Borrelia lipoprotein;Name=PF00921;status=T;Dbxref="InterPro:IPR000680" |
| gene-bpSLO | 001220 | lp24 | gene | Pfam | beta | protein | matc | 20 | 309 | 1E-47 | + | . | Target=null 20 309;ID=match$1177_20_309;signature_desc=Borrelia lipoprotein;Name=PF00921;status=T;Dbxref="InterPro:IPR000680" |
| gene-bpSLO | 001221 | lp24 | gene | Pfam | alpha | protein | matc | 52 | 347 | 2.1E-71 | + | . | Target=null 52 347;ID=match$1467_52_347;signature_desc=Borrelia lipoprotein;Name=PF00921;status=T;Dbxref="InterPro:IPR000680" |
| gene-bpSLO | 001222 | lp24 | gene | Pfam | gamma | protein | matc | 1 | 55 | 2.9E-14 | + | . | Target=null 1 55;ID=match$677_1_55;signature_desc=Borrelia lipoprotein;Name=PF00921;status=T;Dbxref="InterPro:IPR000680" |
| gene-bpSLO | 001228 | lp24 | gene | Pfam | delta | protein | matc | 6 | 39 | 7.6E-08 | + | . | Target=null 6 39;ID=match$575_6_39;signature_desc=Borrelia lipoprotein;Name=PF00921;status=T;Dbxref="InterPro:IPR000680" |
| gene-bpSLO | 001233 | lp28 | gene | Pfam | alpha | protein | matc | 51 | 334 | 1.3E-64 | + | . | Target=null 51 334;ID=match$2453_51_334;signature_desc=Borrelia lipoprotein;Name=PF00921;status=T;Dbxref="InterPro:IPR000680" |
| gene-bpSLO | 001235 | lp28 | gene | Pfam | beta | protein | matc | 49 | 338 | 2E-46 | + | . | Target=null 49 338;ID=match$2177_49_338;signature_desc=Borrelia lipoprotein;Name=PF00921;status=T;Dbxref="InterPro:IPR000680" |
| gene-bpSLO | 001236 | lp28 | gene | Pfam | delta | protein | matc | 45 | 325 | 1.9E-84 | + | . | Target=null 45 325;ID=match$1067_45_325;signature_desc=Borrelia lipoprotein;Name=PF00921;status=T;Dbxref="InterPro:IPR000680" |
| gene-bpSLO | 001237 | lp28 | gene | Pfam | beta | protein | matc | 20 | 302 | 8.4E-45 | + | . | Target=null 20 302;ID=match$1420_20_302;signature_desc=Borrelia lipoprotein;Name=PF00921;status=T;Dbxref="InterPro:IPR000680" |
| gene-bpSLO | 001238 | lp28 | gene | Pfam | alpha | protein | matc | 54 | 349 | 3.1E-69 | + | . | Target=null 54 349;ID=match$1422_54_349;signature_desc=Borrelia lipoprotein;Name=PF00921;status=T;Dbxref="InterPro:IPR000680" |
| gene-bpSLO | 001239 | lp28 | gene | Pfam | beta | protein | matc | 49 | 337 | 2.2E-44 | + | . | Target=null 49 337;ID=match$2187_49_337;signature_desc=Borrelia lipoprotein;Name=PF00921;status=T;Dbxref="InterPro:IPR000680" |
| gene-bpSLO | 001240 | lp28 | gene | Pfam | delta | protein | matc | 44 | 333 | 1E-82 | + | . | Target=null 44 333;ID=match$1953_44_333;signature_desc=Borrelia lipoprotein;Name=PF00921;status=T;Dbxref="InterPro:IPR000680" |
| gene-bpSLO | 001243 | lp28 | gene | Pfam | delta | protein | matc | 38 | 322 | 4.1E-85 | + | . | Target=null 38 322;ID=match$2128_38_322;signature_desc=Borrelia lipoprotein;Name=PF00921;status=T;Dbxref="InterPro:IPR000680" |
| gene-bpSLO | 001246 | lp28 | gene | Pfam | gamma | protein | matc | 54 | 122 | 6.1E-26 | + | . | Target=null 54 122;ID=match$2466_54_122;signature_desc=Borrelia lipoprotein;Name=PF00921;status=T;Dbxref="InterPro:IPR000680" |
| gene-bpSLO | 001253 | lp28 | gene | Pfam | gamma | protein | matc | 1 | 28 | 1.1E-07 | + | . | Target=null 1 28;ID=match$2517_1_28;signature_desc=Borrelia lipoprotein;Name=PF00921;status=T;Dbxref="InterPro:IPR000680" |
| gene-bpSLO | 001254 | lp28 | gene | Pfam | gamma | protein | matc | 7 | 31 | 1.2E-07 | + | . | Target=null 7 31;ID=match$359_7_31;signature_desc=Borrelia lipoprotein;Name=PF00921;status=T;Dbxref="InterPro:IPR000680" |
| gene-bpSLO | 001257 | lp30-1 | gene | Pfam | alpha | protein | matc | 51 | 334 | 1.3E-64 | + | . | Target=null 51 334;ID=match$2451_51_334;signature_desc=Borrelia lipoprotein;Name=PF00921;status=T;Dbxref="InterPro:IPR000680" |
| gene-bpSLO | 001258 | lp30-1 | gene | Pfam | delta | protein | matc | 45 | 329 | 4.9E-86 | + | . | Target=null 45 329;ID=match$2340_45_329;signature_desc=Borrelia lipoprotein;Name=PF00921;status=T;Dbxref="InterPro:IPR000680" |
| gene-bpSLO | 001260 | lp30-1 | gene | Pfam | delta | protein | matc | 34 | 319 | 6.5E-86 | + | . | Target=null 34 319;ID=match$674_34_319;signature_desc=Borrelia lipoprotein;Name=PF00921;status=T;Dbxref="InterPro:IPR000680" |
| gene-bpSLO | 001261 | lp30-1 | gene | Pfam | gamma | protein | matc | 34 | 334 | 1.7E-79 | + | . | Target=null 34 334;ID=match$694_34_334;signature_desc=Borrelia lipoprotein;Name=PF00921;status=T;Dbxref="InterPro:IPR000680" |
| gene-bpSLO | 001282 | lp30-1 | gene | Pfam | beta | protein | matc | 20 | 303 | 5E-48 | + | . | Target=null 20 303;ID=match$798_20_303;signature_desc=Borrelia lipoprotein;Name=PF00921;status=T;Dbxref="InterPro:IPR000680" |
| gene-bpSLO | 001283 | lp30-1 | gene | Pfam | delta | protein | matc | 45 | 334 | 9E-82 | + | . | Target=null 45 334;ID=match$2483_45_334;signature_desc=Borrelia lipoprotein;Name=PF00921;status=T;Dbxref="InterPro:IPR000680" |
| gene-bpSLO | 001287 | lp30-1 | gene | Pfam | delta | protein | matc | 45 | 312 | 1.4E-79 | + | . | Target=null 45 312;ID=match$2970_45_312;signature_desc=Borrelia lipoprotein;Name=PF00921;status=T;Dbxref="InterPro:IPR000680" |
| gene-bpSLO | 001288 | lp30-1 | gene | Pfam | beta | protein | matc | 14 | 305 | 1.7E-50 | + | . | Target=null 14 305;ID=match$2376_14_305;signature_desc=Borrelia lipoprotein;Name=PF00921;status=T;Dbxref="InterPro:IPR000680" |
| gene-bpSLO | 001325 | lp32 | gene | Pfam | beta | protein | matc | 49 | 332 | 1.9E-45 | + | . | Target=null 49 332;ID=match$2854_49_332;signature_desc=Borrelia lipoprotein;Name=PF00921;status=T;Dbxref="InterPro:IPR000680" |
| gene-bpSLO | 001326 | lp32 | pseudo | Pfam | beta | protein | matc | 17 | 305 | 2.5E-50 | + | . | Target=null 17 305;ID=match$2795_17_305;signature_desc=Borrelia lipoprotein;Name=PF00921;status=T;Dbxref="InterPro:IPR000680" |
| gene-bpSLO | 001339 | lp32 | gene | Pfam | delta | protein | matc | 34 | 314 | 5.9E-87 | + | . | Target=null 34 314;ID=match$2772_34_314;signature_desc=Borrelia lipoprotein;Name=PF00921;status=T;Dbxref="InterPro:IPR000680" |
| gene-bpSLO | 001341 | lp32 | gene | Pfam | alpha | protein | matc | 43 | 338 | 8.5E-69 | + | . | Target=null 43 338;ID=match$1136_43_338;signature_desc=Borrelia lipoprotein;Name=PF00921;status=T;Dbxref="InterPro:IPR000680" |
| gene-bpSLO | 001342 | lp32 | gene | Pfam | beta | protein | matc | 20 | 306 | 9.9E-48 | + | . | Target=null 20 306;ID=match$2370_20_306;signature_desc=Borrelia lipoprotein;Name=PF00921;status=T;Dbxref="InterPro:IPR000680" |
| gene-bpSLO | 001343 | lp32 | gene | Pfam | gamma | protein | matc | 34 | 316 | 2.1E-87 | + | . | Target=null 34 316;ID=match$3038_34_316;signature_desc=Borrelia lipoprotein;Name=PF00921;status=T;Dbxref="InterPro:IPR000680" |
| gene-bpSLO | 001344 | lp32 | gene | Pfam | delta | protein | matc | 44 | 333 | 2.7E-80 | + | . | Target=null 44 333;ID=match$1038_44_333;signature_desc=Borrelia lipoprotein;Name=PF00921;status=T;Dbxref="InterPro:IPR000680" |
| gene-bpSLO | 001345 | lp32 | gene | Pfam | delta | protein | matc | 44 | 328 | 7.2E-87 | + | . | Target=null 44 328;ID=match$1934_44_328;signature_desc=Borrelia lipoprotein;Name=PF00921;status=T;Dbxref="InterPro:IPR000680" |
| gene-bpSLO | 001346 | lp32 | gene | Pfam | beta | protein | matc | 49 | 338 | 2.4E-46 | + | . | Target=null 49 338;ID=match$439_49_338;signature_desc=Borrelia lipoprotein;Name=PF00921;status=T;Dbxref="InterPro:IPR000680" |
| gene-bpSLO | 001347 | lp32 | gene | Pfam | delta | protein | matc | 45 | 326 | 7.9E-86 | + | . | Target=null 45 326;ID=match$2405_45_326;signature_desc=Borrelia lipoprotein;Name=PF00921;status=T;Dbxref="InterPro:IPR000680" |
| gene-bpSLO | 001348 | lp32 | gene | Pfam | gamma | protein | matc | 34 | 314 | 6.1E-92 | + | . | Target=null 34 314;ID=match$369_34_314;signature_desc=Borrelia lipoprotein;Name=PF00921;status=T;Dbxref="InterPro:IPR000680" |
| gene-bpSLO | 001351 | lp32 | gene | Pfam | gamma | protein | matc | 34 | 319 | 1.8E-84 | + | . | Target=null 34 319;ID=match$301_34_319;signature_desc=Borrelia lipoprotein;Name=PF00921;status=T;Dbxref="InterPro:IPR000680" |
| gene-bpSLO | 001352 | lp32 | gene | Pfam | alpha | protein | matc | 51 | 336 | 6E-65 | + | . | Target=null 51 336;ID=match$2424_51_336;signature_desc=Borrelia lipoprotein;Name=PF00921;status=T;Dbxref="InterPro:IPR000680" |
| gene-bpSLO | 001357 | lp34-1 | gene | Pfam | gamma | protein | matc | 14 | 301 | 6.2E-48 | + | . | Target=null 14 301;ID=match$1018_14_301;signature_desc=Borrelia lipoprotein;Name=PF00921;status=T;Dbxref="InterPro:IPR000680" |
| gene-bpSLO | 001358 | lp34-1 | gene | Pfam | alpha | protein | matc | 43 | 340 | 2.1E-59 | + | . | Target=null 43 340;ID=match$2890_43_340;signature_desc=Borrelia lipoprotein;Name=PF00921;status=T;Dbxref="InterPro:IPR000680" |
| gene-bpSLO | 001359 | lp34-1 | gene | Pfam | gamma | protein | matc | 178 | 268 | 2.1E-27 | + | . | Target=null 178 268;ID=match$678_178_268;signature_desc=Borrelia lipoprotein;Name=PF00921;status=T;Dbxref="InterPro:IPR000680" |
| gene-bpSLO | 001359 | lp34-1 | gene | Pfam | | protein | matc | 28 | 198 | 7E-47 | + | . | Target=null 28 198;ID=match$678_28_198;signature_desc=Borrelia lipoprotein;Name=PF00921;status=T;Dbxref="InterPro:IPR000680" |
| gene-bpSLO | 001372 | lp34-1 | gene | Pfam | gamma | protein | matc | 1 | 273 | 2.3E-79 | + | . | Target=null 1 273;ID=match$152_1_273;signature_desc=Borrelia lipoprotein;Name=PF00921;status=T;Dbxref="InterPro:IPR000680" |
| gene-bpSLO | 001373 | lp34-1 | gene | Pfam | delta | protein | matc | 44 | 321 | 3.1E-86 | + | . | Target=null 44 321;ID=match$1334_44_321;signature_desc=Borrelia lipoprotein;Name=PF00921;status=T;Dbxref="InterPro:IPR000680" |
| gene-bpSLO | 001375 | lp34-1 | gene | Pfam | gamma | protein | matc | 7 | 287 | 2.6E-79 | + | . | Target=null 7 287;ID=match$137_7_287;signature_desc=Borrelia lipoprotein;Name=PF00921;status=T;Dbxref="InterPro:IPR000680" |
| gene-bpSLO | 001376 | lp34-1 | gene | Pfam | alpha | protein | matc | 55 | 354 | 5.6E-70 | + | . | Target=null 55 354;ID=match$1529_55_354;signature_desc=Borrelia lipoprotein;Name=PF00921;status=T;Dbxref="InterPro:IPR000680" |
| gene-bpSLO | 001377 | lp34-1 | gene | Pfam | beta | protein | matc | 14 | 301 | 1.2E-44 | + | . | Target=null 14 301;ID=match$17_14_301;signature_desc=Borrelia lipoprotein;Name=PF00921;status=T;Dbxref="InterPro:IPR000680" |
| gene-bpSLO | 001379 | lp34-1 | gene | Pfam | gamma | protein | matc | 1 | 278 | 5.6E-82 | + | . | Target=null 1 278;ID=match$176_1_278;signature_desc=Borrelia lipoprotein;Name=PF00921;status=T;Dbxref="InterPro:IPR000680" |
| gene-bpSLO | 001380 | lp34-1 | gene | Pfam | alpha | protein | matc | 45 | 341 | 1.8E-73 | + | . | Target=null 45 341;ID=match$242_45_341;signature_desc=Borrelia lipoprotein;Name=PF00921;status=T;Dbxref="InterPro:IPR000680" |
| gene-bpSLO | 001381 | lp34-1 | gene | Pfam | beta | protein | matc | 20 | 305 | 5.7E-46 | + | . | Target=null 20 305;ID=match$2734_20_305;signature_desc=Borrelia lipoprotein;Name=PF00921;status=T;Dbxref="InterPro:IPR000680" |
| gene-bpSLO | 001382 | lp34-1 | gene | Pfam | delta | protein | matc | 45 | 335 | 1.6E-84 | + | . | Target=null 45 335;ID=match$1711_45_335;signature_desc=Borrelia lipoprotein;Name=PF00921;status=T;Dbxref="InterPro:IPR000680" |
| gene-bpSLO | 001384 | lp34-1 | gene | Pfam | gamma | protein | matc | 34 | 319 | 4.8E-93 | + | . | Target=null 34 319;ID=match$1340_34_319;signature_desc=Borrelia lipoprotein;Name=PF00921;status=T;Dbxref="InterPro:IPR000680" |
| gene-bpSLO | 001385 | lp34-1 | gene | Pfam | delta | protein | matc | 45 | 333 | 1.6E-86 | + | . | Target=null 45 333;ID=match$1559_45_333;signature_desc=Borrelia lipoprotein;Name=PF00921;status=T;Dbxref="InterPro:IPR000680" |
| gene-bpSLO | 001386 | lp34-1 | gene | Pfam | alpha | protein | matc | 55 | 338 | 4.9E-63 | + | . | Target=null 55 338;ID=match$2802_55_338;signature_desc=Borrelia lipoprotein;Name=PF00921;status=T;Dbxref="InterPro:IPR000680" |
| gene-bpSLO | 001389 | lp34-2 | gene | Pfam | alpha | protein | matc | 5 | 191 | 8.7E-43 | + | . | Target=null 5 191;ID=match$1254_5_191;signature_desc=Borrelia lipoprotein;Name=PF00921;status=T;Dbxref="InterPro:IPR000680" |
| gene-bpSLO | 001413 | lp34-2 | gene | Pfam | gamma | protein | matc | 178 | 268 | 2.1E-27 | + | . | Target=null 178 268;ID=match$680_178_268;signature_desc=Borrelia lipoprotein;Name=PF00921;status=T;Dbxref="InterPro:IPR000680" |
| gene-bpSLO | 001413 | lp34-2 | gene | Pfam | | protein | matc | 28 | 198 | 7E-47 | + | . | Target=null 28 198;ID=match$680_28_198;signature_desc=Borrelia lipoprotein;Name=PF00921;status=T;Dbxref="InterPro:IPR000680" |
| gene-bpSLO | 001414 | lp34-2 | gene | Pfam | alpha | protein | matc | 43 | 340 | 2.1E-59 | + | . | Target=null 43 340;ID=match$2886_43_340;signature_desc=Borrelia lipoprotein;Name=PF00921;status=T;Dbxref="InterPro:IPR000680" |
| gene-bpSLO | 001415 | lp34-2 | gene | Pfam | beta | protein | matc | 14 | 301 | 6.2E-48 | + | . | Target=null 14 301;ID=match$1020_14_301;signature_desc=Borrelia lipoprotein;Name=PF00921;status=T;Dbxref="InterPro:IPR000680" |
| gene-bpSLO | 001420 | lp42 | gene | Pfam | alpha | protein | matc | 51 | 336 | 6E-65 | + | . | Target=null 51 336;ID=match$2422_51_336;signature_desc=Borrelia lipoprotein;Name=PF00921;status=T;Dbxref="InterPro:IPR000680" |
| gene-bpSLO | 001421 | lp42 | gene | Pfam | gamma | protein | matc | 34 | 319 | 1.8E-84 | + | . | Target=null 34 319;ID=match$303_34_319;signature_desc=Borrelia lipoprotein;Name=PF00921;status=T;Dbxref="InterPro:IPR000680" |
| gene-bpSLO | 001424 | lp42 | gene | Pfam | gamma | protein | matc | 34 | 314 | 6.1E-92 | + | . | Target=null 34 314;ID=match$365_34_314;signature_desc=Borrelia lipoprotein;Name=PF00921;status=T;Dbxref="InterPro:IPR000680" |
| gene-bpSLO | 001425 | lp42 | gene | Pfam | delta | protein | matc | 45 | 326 | 7.9E-86 | + | . | Target=null 45 326;ID=match$2407_45_326;signature_desc=Borrelia lipoprotein;Name=PF00921;status=T;Dbxref="InterPro:IPR000680" |
| gene-bpSLO | 001426 | lp42 | gene | Pfam | beta | protein | matc | 49 | 338 | 2.4E-46 | + | . | Target=null 49 338;ID=match$441_49_338;signature_desc=Borrelia lipoprotein;Name=PF00921;status=T;Dbxref="InterPro:IPR000680" |
| gene-bpSLO | 001427 | lp42 | gene | Pfam | delta | protein | matc | 44 | 323 | 1.4E-87 | + | . | Target=null 44 323;ID=match$2512_44_323;signature_desc=Borrelia lipoprotein;Name=PF00921;status=T;Dbxref="InterPro:IPR000680" |
| gene-bpSLO | 001428 | lp42 | gene | Pfam | delta | protein | matc | 8 | 291 | 2.1E-83 | + | . | Target=null 8 291;ID=match$2224_8_291;signature_desc=Borrelia lipoprotein;Name=PF00921;status=T;Dbxref="InterPro:IPR000680" |
| gene-bpSLO | 001430 | lp42 | gene | Pfam | delta | protein | matc | 34 | 324 | 7E-85 | + | . | Target=null 34 324;ID=match$1519_34_324;signature_desc=Borrelia lipoprotein;Name=PF00921;status=T;Dbxref="InterPro:IPR000680" |
| gene-bpSLO | 001431 | lp42 | gene | Pfam | beta | protein | matc | 20 | 310 | 1.8E-49 | + | . | Target=null 20 310;ID=match$2500_20_310;signature_desc=Borrelia lipoprotein;Name=PF00921;status=T;Dbxref="InterPro:IPR000680" |
| gene-bpSLO | 001432 | lp42 | gene | Pfam | alpha | protein | matc | 7 | 304 | 1.2E-72 | + | . | Target=null 7 304;ID=match$380_7_304;signature_desc=Borrelia lipoprotein;Name=PF00921;status=T;Dbxref="InterPro:IPR000680" |
| gene-bpSLO | 001433 | lp42 | pseudo | Pfam | delta | protein | matc | 19 | 62 | 1.2E-13 | + | . | Target=null 19 62;ID=match$2218_19_62;signature_desc=Borrelia lipoprotein;Name=PF00921;status=T;Dbxref="InterPro:IPR000680" |
| gene-bpSLO | 001435 | lp42 | gene | Pfam | delta | protein | matc | 34 | 319 | 3.3E-89 | + | . | Target=null 34 319;ID=match$200_34_319;signature_desc=Borrelia lipoprotein;Name=PF00921;status=T;Dbxref="InterPro:IPR000680" |
| gene-bpSLO | 001437 | lp42 | gene | Pfam | delta | protein | matc | 8 | 144 | 1.7E-37 | + | . | Target=null 8 144;ID=match$2793_8_144;signature_desc=Borrelia lipoprotein;Name=PF00921;status=T;Dbxref="InterPro:IPR000680" |
| gene-bpSLO | 001439 | lp42 | gene | Pfam | gamma | protein | matc | 11 | 50 | 1.5E-10 | + | . | Target=null 11 50;ID=match$1894_11_50;signature_desc=Borrelia lipoprotein;Name=PF00921;status=T;Dbxref="InterPro:IPR000680" |
| gene-bpSLO | 001445 | lp42 | gene | Pfam | beta | protein | matc | 49 | 335 | 5.8E-51 | + | . | Target=null 49 335;ID=match$2640_49_335;signature_desc=Borrelia lipoprotein;Name=PF00921;status=T;Dbxref="InterPro:IPR000680" |
| gene-bpSLO | 001448 | lp42 | pseudo | Pfam | beta | protein | matc | 48 | 187 | 1.5E-22 | + | . | Target=null 48 187;ID=match$2645_48_187;signature_desc=Borrelia lipoprotein;Name=PF00921;status=T;Dbxref="InterPro:IPR000680" |
| gene-bpSLO | 001448 | lp42 | pseudo | Pfam | | protein | matc | 22 | 137 | 1.4E-19 | + | . | Target=null 22 137;ID=match$1819_22_137;signature_desc=Borrelia lipoprotein;Name=PF00921;status=T;Dbxref="InterPro:IPR000680" |
| gene-bpSLO | 001450 | lp42 | pseudo | Pfam | gamma | protein | matc | 124 | 197 | 1.4E-17 | + | . | Target=null 124 197;ID=match$1052_124_197;signature_desc=Borrelia lipoprotein;Name=PF00921;status=T;Dbxref="InterPro:IPR000680" |
| gene-bpSLO | 001450 | lp42 | pseudo | Pfam | | protein | matc | 34 | 132 | 2.7E-30 | + | . | Target=null 34 132;ID=match$1052_34_132;signature_desc=Borrelia lipoprotein;Name=PF00921;status=T;Dbxref="InterPro:IPR000680" |
| gene-bpSLO | 001462 | lp58 | gene | Pfam | alpha | protein | matc | 51 | 336 | 6E-65 | + | . | Target=null 51 336;ID=match$2420_51_336;signature_desc=Borrelia lipoprotein;Name=PF00921;status=T;Dbxref="InterPro:IPR000680" |
| gene-bpSLO | 001463 | lp58 | gene | Pfam | gamma | protein | matc | 34 | 319 | 1.8E-84 | + | . | Target=null 34 319;ID=match$299_34_319;signature_desc=Borrelia lipoprotein;Name=PF00921;status=T;Dbxref="InterPro:IPR000680" |
| gene-bpSLO | 001466 | lp58 | gene | Pfam | gamma | protein | matc | 34 | 314 | 6.1E-92 | + | . | Target=null 34 314;ID=match$367_34_314;signature_desc=Borrelia lipoprotein;Name=PF00921;status=T;Dbxref="InterPro:IPR000680" |
| gene-bpSLO | 001467 | lp58 | gene | Pfam | delta | protein | matc | 45 | 326 | 7.9E-86 | + | . | Target=null 45 326;ID=match$2409_45_326;signature_desc=Borrelia lipoprotein;Name=PF00921;status=T;Dbxref="InterPro:IPR000680" |
| gene-bpSLO | 001468 | lp58 | gene | Pfam | beta | protein | matc | 49 | 338 | 2.4E-46 | + | . | Target=null 49 338;ID=match$443_49_338;signature_desc=Borrelia lipoprotein;Name=PF00921;status=T;Dbxref="InterPro:IPR000680" |
| gene-bpSLO | 001469 | lp58 | gene | Pfam | delta | protein | matc | 44 | 323 | 1.4E-87 | + | . | Target=null 44 323;ID=match$2514_44_323;signature_desc=Borrelia lipoprotein;Name=PF00921;status=T;Dbxref="InterPro:IPR000680" |
| gene-bpSLO | 001470 | lp58 | gene | Pfam | delta | protein | matc | 45 | 331 | 2.1E-83 | + | . | Target=null 45 331;ID=match$822_45_331;signature_desc=Borrelia lipoprotein;Name=PF00921;status=T;Dbxref="InterPro:IPR000680" |
| gene-bpSLO | 001474 | lp58 | gene | Pfam | delta | protein | matc | 38 | 159 | 1.3E-34 | + | . | Target=null 38 159;ID=match$1845_38_159;signature_desc=Borrelia lipoprotein;Name=PF00921;status=T;Dbxref="InterPro:IPR000680" |
| gene-bpSLO | 001476 | lp58 | gene | Pfam | beta | protein | matc | 29 | 318 | 4.9E-46 | + | . | Target=null 29 318;ID=match$943_29_318;signature_desc=Borrelia lipoprotein;Name=PF00921;status=T;Dbxref="InterPro:IPR000680" |
| gene-bpSLO | 001477 | lp58 | gene | Pfam | alpha | protein | matc | 7 | 303 | 3.6E-71 | + | . | Target=null 7 303;ID=match$522_7_303;signature_desc=Borrelia lipoprotein;Name=PF00921;status=T;Dbxref="InterPro:IPR000680" |
| gene-bpSLO | 001478 | lp58 | gene | Pfam | gamma | protein | matc | 34 | 325 | 4.6E-87 | + | . | Target=null 34 325;ID=match$834_34_325;signature_desc=Borrelia lipoprotein;Name=PF00921;status=T;Dbxref="InterPro:IPR000680" |
| gene-bpSLO | 001479 | lp58 | gene | Pfam | delta | protein | matc | 44 | 334 | 8.7E-86 | + | . | Target=null 44 334;ID=match$2022_44_334;signature_desc=Borrelia lipoprotein;Name=PF00921;status=T;Dbxref="InterPro:IPR000680" |
| gene-bpSLO | 001482 | lp58 | gene | Pfam | delta | protein | matc | 34 | 315 | 6.2E-88 | + | . | Target=null 34 315;ID=match$3003_34_315;signature_desc=Borrelia lipoprotein;Name=PF00921;status=T;Dbxref="InterPro:IPR000680" |
| gene-bpSLO | 001484 | lp58 | pseudo | Pfam | beta | protein | matc | 3 | 171 | 2.2E-15 | + | . | Target=null 3 171;ID=match$88_3_171;signature_desc=Borrelia lipoprotein;Name=PF00921;status=T;Dbxref="InterPro:IPR000680" |
| gene-bpSLO | 001484 | lp58 | pseudo | Pfam | | protein | matc | 165 | 212 | 7.7E-09 | + | . | Target=null 165 212;ID=match$88_165_212;signature_desc=Borrelia lipoprotein;Name=PF00921;status=T;Dbxref="InterPro:IPR000680" |
| gene-bpSLO | 001485 | lp58 | gene | Pfam | gamma | protein | matc | 117 | 246 | 3.4E-35 | + | . | Target=null 117 246;ID=match$2893_117_246;signature_desc=Borrelia lipoprotein;Name=PF00921;status=T;Dbxref="InterPro:IPR000680" |
| gene-bpSLO | 001485 | lp58 | gene | Pfam | | protein | matc | 28 | 119 | 1.8E-33 | + | . | Target=null 28 119;ID=match$2893_28_119;signature_desc=Borrelia lipoprotein;Name=PF00921;status=T;Dbxref="InterPro:IPR000680" |
| gene-bpSLO | 001489 | lp58 | gene | Pfam | gamma | protein | matc | 7 | 173 | 1.8E-44 | + | . | Target=null 7 173;ID=match$2150_7_173;signature_desc=Borrelia lipoprotein;Name=PF00921;status=T;Dbxref="InterPro:IPR000680" |
| gene-bpSLO | 001507 | lp58 | gene | Pfam | delta | protein | matc | 45 | 244 | 5.4E-51 | + | . | Target=null 45 244;ID=match$390_45_244;signature_desc=Borrelia lipoprotein;Name=PF00921;status=T;Dbxref="InterPro:IPR000680" |
| gene-bpSLO | 001507 | lp58 | gene | Pfam | | protein | matc | 239 | 280 | 4.3E-10 | + | . | Target=null 239 280;ID=match$390_239_280;signature_desc=Borrelia lipoprotein;Name=PF00921;status=T;Dbxref="InterPro:IPR000680" |
| gene-bpSLO | 001510 | lp58 | gene | Pfam | delta | protein | matc | 45 | 329 | 2.8E-85 | + | . | Target=null 45 329;ID=match$1092_45_329;signature_desc=Borrelia lipoprotein;Name=PF00921;status=T;Dbxref="InterPro:IPR000680" |
| gene-bpSLO | 001513 | lp58 | pseudo | Pfam | gamma | protein | matc | 43 | 165 | 6.7E-39 | + | . | Target=null 43 165;ID=match$2098_43_165;signature_desc=Borrelia lipoprotein;Name=PF00921;status=T;Dbxref="InterPro:IPR000680" |
| gene-bpSLO | 001513 | lp58 | pseudo | Pfam | | protein | matc | 4 | 166 | 6.7E-47 | + | . | Target=null 4 166;ID=match$1537_4_166;signature_desc=Borrelia lipoprotein;Name=PF00921;status=T;Dbxref="InterPro:IPR000680" |
| gene-bpSLO | 001516 | lp58 | gene | Pfam | alpha | protein | matc | 57 | 357 | 3.1E-69 | + | . | Target=null 57 357;ID=match$3196_57_357;signature_desc=Borrelia lipoprotein;Name=PF00921;status=T;Dbxref="InterPro:IPR000680" |
| gene-bpSLO | 001517 | lp58 | pseudo | Pfam | beta | protein | matc | 14 | 154 | 1.1E-18 | + | . | Target=null 14 154;ID=match$2502_14_154;signature_desc=Borrelia lipoprotein;Name=PF00921;status=T;Dbxref="InterPro:IPR000680" |
| gene-bpSLO | 001518 | lp58 | gene | Pfam | delta | protein | matc | 34 | 320 | 1.4E-88 | + | . | Target=null 34 320;ID=match$2786_34_320;signature_desc=Borrelia lipoprotein;Name=PF00921;status=T;Dbxref="InterPro:IPR000680" |
| gene-bpSLO | 001520 | lp58 | gene | Pfam | gamma | protein | matc | 34 | 319 | 4.8E-93 | + | . | Target=null 34 319;ID=match$1342_34_319;signature_desc=Borrelia lipoprotein;Name=PF00921;status=T;Dbxref="InterPro:IPR000680" |
| gene-bpSLO | 001521 | lp58 | gene | Pfam | delta | protein | matc | 44 | 332 | 1.8E-86 | + | . | Target=null 44 332;ID=match$655_44_332;signature_desc=Borrelia lipoprotein;Name=PF00921;status=T;Dbxref="InterPro:IPR000680" |
| gene-bpSLO | 001522 | lp58 | gene | Pfam | alpha | protein | matc | 55 | 338 | 4.9E-63 | + | . | Target=null 55 338;ID=match$2800_55_338;signature_desc=Borrelia lipoprotein;Name=PF00921;status=T;Dbxref="InterPro:IPR000680" |
| gene-bpSLO | 001527 | lp60 | gene | Pfam | beta | protein | matc | 14 | 301 | 6.2E-48 | + | . | Target=null 14 301;ID=match$1016_14_301;signature_desc=Borrelia lipoprotein;Name=PF00921;status=T;Dbxref="InterPro:IPR000680" |
| gene-bpSLO | 001528 | lp60 | gene | Pfam | alpha | protein | matc | 43 | 340 | 2.1E-59 | + | . | Target=null 43 340;ID=match$2888_43_340;signature_desc=Borrelia lipoprotein;Name=PF00921;status=T;Dbxref="InterPro:IPR000680" |
| gene-bpSLO | 001529 | lp60 | gene | Pfam | gamma | protein | matc | 178 | 268 | 2.1E-27 | + | . | Target=null 178 268;ID=match$682_178_268;signature_desc=Borrelia lipoprotein;Name=PF00921;status=T;Dbxref="InterPro:IPR000680" |
| gene-bpSLO | 001529 | lp60 | gene | Pfam | | protein | matc | 28 | 198 | 7E-47 | + | . | Target=null 28 198;ID=match$682_28_198;signature_desc=Borrelia lipoprotein;Name=PF00921;status=T;Dbxref="InterPro:IPR000680" |
| gene-bpSLO | 001531 | lp60 | gene | Pfam | gamma | protein | matc | 44 | 319 | 2.6E-86 | + | . | Target=null 44 319;ID=match$107_44_319;signature_desc=Borrelia lipoprotein;Name=PF00921;status=T;Dbxref="InterPro:IPR000680" |
| gene-bpSLO | 001532 | lp60 | gene | Pfam | delta | protein | matc | 74 | 358 | 5E-86 | + | . | Target=null 74 358;ID=match$2869_74_358;signature_desc=Borrelia lipoprotein;Name=PF00921;status=T;Dbxref="InterPro:IPR000680" |
| gene-bpSLO | 001534 | lp60 | gene | Pfam | beta | protein | matc | 115 | 244 | 4.9E-24 | + | . | Target=null 115 244;ID=match$3055_115_244;signature_desc=Borrelia lipoprotein;Name=PF00921;status=T;Dbxref="InterPro:IPR000680" |
| gene-bpSLO | 001534 | lp60 | gene | Pfam | | protein | matc | 14 | 110 | 2.3E-10 | + | . | Target=null 14 110;ID=match$3055_14_110;signature_desc=Borrelia lipoprotein;Name=PF00921;status=T;Dbxref="InterPro:IPR000680" |
| gene-bpSLO | 001535 | lp60 | gene | Pfam | alpha | protein | matc | 55 | 354 | 1.3E-71 | + | . | Target=null 55 354;ID=match$2268_55_354;signature_desc=Borrelia lipoprotein;Name=PF00921;status=T;Dbxref="InterPro:IPR000680" |
| gene-bpSLO | 001537 | lp60 | gene | Pfam | gamma | protein | matc | 1 | 284 | 4.7E-85 | + | . | Target=null 1 284;ID=match$191_1_284;signature_desc=Borrelia lipoprotein;Name=PF00921;status=T;Dbxref="InterPro:IPR000680" |
| gene-bpSLO | 001540 | lp60 | gene | Pfam | gamma | protein | matc | 1 | 279 | 2E-83 | + | . | Target=null 1 279;ID=match$1281_1_279;signature_desc=Borrelia lipoprotein;Name=PF00921;status=T;Dbxref="InterPro:IPR000680" |
| gene-bpSLO | 001544 | lp60 | gene | Pfam | delta | protein | matc | 45 | 324 | 1.1E-84 | + | . | Target=null 45 324;ID=match$2619_45_324;signature_desc=Borrelia lipoprotein;Name=PF00921;status=T;Dbxref="InterPro:IPR000680" |
| gene-bpSLO | 001545 | lp60 | pseudo | Pfam | delta | protein | matc | 13 | 88 | 1.6E-22 | + | . | Target=null 13 88;ID=match$1850_13_88;signature_desc=Borrelia lipoprotein;Name=PF00921;status=T;Dbxref="InterPro:IPR000680" |
| gene-bpSLO | 001545 | lp60 | pseudo | Pfam | | protein | matc | 12 | 72 | 1.1E-08 | + | . | Target=null 12 72;ID=match$2815_12_72;signature_desc=Borrelia lipoprotein;Name=PF00921;status=T;Dbxref="InterPro:IPR000680" |
| gene-bpSLO | 001545 | lp60 | pseudo | Pfam | | protein | matc | 74 | 141 | 1.4E-28 | + | . | Target=null 74 141;ID=match$2815_74_141;signature_desc=Borrelia lipoprotein;Name=PF00921;status=T;Dbxref="InterPro:IPR000680" |
| gene-bpSLO | 001568 | lp60 | gene | Pfam | delta | protein | matc | 29 | 313 | 6.3E-80 | + | . | Target=null 29 313;ID=match$1862_29_313;signature_desc=Borrelia lipoprotein;Name=PF00921;status=T;Dbxref="InterPro:IPR000680" |
| gene-bpSLO | 001570 | lp60 | gene | Pfam | alpha | protein | matc | 54 | 350 | 6.4E-68 | + | . | Target=null 54 350;ID=match$760_54_350;signature_desc=Borrelia lipoprotein;Name=PF00921;status=T;Dbxref="InterPro:IPR000680" |
| gene-bpSLO | 001571 | lp60 | pseudo | Pfam | beta | protein | matc | 20 | 270 | 6.3E-31 | + | . | Target=null 20 270;ID=match$2951_20_270;signature_desc=Borrelia lipoprotein;Name=PF00921;status=T;Dbxref="InterPro:IPR000680" |
| gene-bpSLO | 001571 | lp60 | pseudo | Pfam | | protein | matc | 12 | 50 | 4.4E-11 | + | . | Target=null 12 50;ID=match$1553_12_50;signature_desc=Borrelia lipoprotein;Name=PF00921;status=T;Dbxref="InterPro:IPR000680" |
| gene-bpSLO | 001572 | lp60 | gene | Pfam | gamma | protein | matc | 35 | 313 | 1.7E-86 | + | . | Target=null 35 313;ID=match$695_35_313;signature_desc=Borrelia lipoprotein;Name=PF00921;status=T;Dbxref="InterPro:IPR000680" |
| gene-bpSLO | 001573 | lp60 | pseudo | Pfam | delta | protein | matc | 44 | 250 | 6.9E-53 | + | . | Target=null 44 250;ID=match$1054_44_250;signature_desc=Borrelia lipoprotein;Name=PF00921;status=T;Dbxref="InterPro:IPR000680" |
| gene-bpSLO | 001573 | lp60 | pseudo | Pfam | | protein | matc | 5 | 77 | 5.5E-28 | + | . | Target=null 5 77;ID=match$2068_5_77;signature_desc=Borrelia lipoprotein;Name=PF00921;status=T;Dbxref="InterPro:IPR000680" |
| gene-bpSLO | 001574 | lp60 | gene | Pfam | delta | protein | matc | 45 | 324 | 7.8E-82 | + | . | Target=null 45 324;ID=match$612_45_324;signature_desc=Borrelia lipoprotein;Name=PF00921;status=T;Dbxref="InterPro:IPR000680" |
| gene-bpSLO | 001575 | lp60 | gene | Pfam | gamma | protein | matc | 34 | 319 | 1.2E-84 | + | . | Target=null 34 319;ID=match$2417_34_319;signature_desc=Borrelia lipoprotein;Name=PF00921;status=T;Dbxref="InterPro:IPR000680" |
| gene-bpSLO | 001578 | lp60 | gene | Pfam | delta | protein | matc | 44 | 330 | 2.3E-87 | + | . | Target=null 44 330;ID=match$567_44_330;signature_desc=Borrelia lipoprotein;Name=PF00921;status=T;Dbxref="InterPro:IPR000680" |
| gene-bpSLO | 001580 | lp60 | gene | Pfam | gamma | protein | matc | 43 | 335 | 1.9E-91 | + | . | Target=null 43 335;ID=match$2282_43_335;signature_desc=Borrelia lipoprotein;Name=PF00921;status=T;Dbxref="InterPro:IPR000680" |
| gene-bpSLO | 001584 | lp60 | gene | Pfam | delta | protein | matc | 44 | 330 | 1.9E-88 | + | . | Target=null 44 330;ID=match$1111_44_330;signature_desc=Borrelia lipoprotein;Name=PF00921;status=T;Dbxref="InterPro:IPR000680" |
| gene-bpSLO | 001586 | lp60 | pseudo | Pfam | gamma | protein | matc | 36 | 73 | 3.8E-13 | + | . | Target=null 36 73;ID=match$437_36_73;signature_desc=Borrelia lipoprotein;Name=PF00921;status=T;Dbxref="InterPro:IPR000680" |
| gene-bpSLO | 001586 | lp60 | pseudo | Pfam | | protein | matc | 2 | 245 | 7.9E-67 | + | . | Target=null 2 245;ID=match$820_2_245;signature_desc=Borrelia lipoprotein;Name=PF00921;status=T;Dbxref="InterPro:IPR000680" |
| gene-bpSLO | 001587 | lp60 | gene | Pfam | delta | protein | matc | 44 | 323 | 1.9E-85 | + | . | Target=null 44 323;ID=match$1431_44_323;signature_desc=Borrelia lipoprotein;Name=PF00921;status=T;Dbxref="InterPro:IPR000680" |
| gene-bpSLO | 001590 | lp60 | gene | Pfam | beta | protein | matc | 48 | 334 | 2.7E-51 | + | . | Target=null 48 334;ID=match$762_48_334;signature_desc=Borrelia lipoprotein;Name=PF00921;status=T;Dbxref="InterPro:IPR000680" |

| Vsp | replicon | gene/pseudo | | | | | | | | | |
|---|---|---|---|---|---|---|---|---|---|---|---|
| gene-bvRMA01_001093 | lp25 | gene | | Pfam | protein_matc | 39 | 210 | 1.2E-46 | + | . | Target=null 39 210;ID=match$653_39_210;signature_desc=Lipoprotein;Name=PF01441;status=T;Dbxref="InterPro:IPR001800" |
| gene-bvRMA01_001109 | lp30 | gene | | Pfam | protein_matc | 39 | 209 | 2E-58 | + | . | Target=null 39 209;ID=match$1736_39_209;signature_desc=Lipoprotein;Name=PF01441;status=T;Dbxref="InterPro:IPR001800" |
| gene-bvRMA01_001121 | lp30 | gene | | Pfam | protein_matc | 38 | 204 | 1E-62 | + | . | Target=null 38 204;ID=match$1966_38_204;signature_desc=Lipoprotein;Name=PF01441;status=T;Dbxref="InterPro:IPR001800" |
| gene-bvRMA01_001128 | lp30 | gene | | Pfam | protein_matc | 39 | 208 | 1.7E-65 | + | . | Target=null 39 208;ID=match$2240_39_208;signature_desc=Lipoprotein;Name=PF01441;status=T;Dbxref="InterPro:IPR001800" |
| gene-bvRMA01_001142 | lp31 | gene | | Pfam | protein_matc | 39 | 206 | 8.2E-60 | + | . | Target=null 39 206;ID=match$114_39_206;signature_desc=Lipoprotein;Name=PF01441;status=T;Dbxref="InterPro:IPR001800" |
| gene-bvRMA01_001144 | lp31 | gene | | Pfam | protein_matc | 39 | 207 | 3.1E-53 | + | . | Target=null 39 207;ID=match$218_39_207;signature_desc=Lipoprotein;Name=PF01441;status=T;Dbxref="InterPro:IPR001800" |
| gene-bvRMA01_001160 | lp31 | gene | | Pfam | protein_matc | 39 | 206 | 7.7E-54 | + | . | Target=null 39 206;ID=match$2686_39_206;signature_desc=Lipoprotein;Name=PF01441;status=T;Dbxref="InterPro:IPR001800" |
| gene-bvRMA01_001196 | lp35 | gene | | Pfam | protein_matc | 39 | 209 | 2E-58 | + | . | Target=null 39 209;ID=match$1734_39_209;signature_desc=Lipoprotein;Name=PF01441;status=T;Dbxref="InterPro:IPR001800" |
| gene-bvRMA01_001228 | lp37 | pseudo | | Pfam | protein_matc | 38 | 117 | 5.1E-35 | + | . | Target=null 38 117;ID=match$613_38_117;signature_desc=Lipoprotein;Name=PF01441;status=T;Dbxref="InterPro:IPR001800" |
| gene-bvRMA01_001228 | lp37 | pseudo | | Pfam | protein_matc | 3 | 94 | 1.3E-27 | + | . | Target=null 3 94;ID=match$1272_3_94;signature_desc=Lipoprotein;Name=PF01441;status=T;Dbxref="InterPro:IPR001800" |

| Vlp | | | Subfamily | | | | | | | | |
|---|---|---|---|---|---|---|---|---|---|---|---|
| gene-bvRMA01_001081 | lp17-2 | gene | delta | Pfam | protein_matc | 34 | 316 | 1E-86 | + | . | Target=null 34 316;ID=match$2672_34_316;signature_desc=Borrelia lipoprotein;Name=PF00921;status=T;Dbxref="InterPro:IPR000680" |
| gene-bvRMA01_001087 | lp25 | gene | alpha | Pfam | protein_matc | 52 | 343 | 2E-70 | + | . | Target=null 52 343;ID=match$999_52_343;signature_desc=Borrelia lipoprotein;Name=PF00921;status=T;Dbxref="InterPro:IPR000680" |
| gene-bvRMA01_001088 | lp25 | gene | gamma | Pfam | protein_matc | 34 | 309 | 2E-86 | + | . | Target=null 34 309;ID=match$335_34_309;signature_desc=Borrelia lipoprotein;Name=PF00921;status=T;Dbxref="InterPro:IPR000680" |
| gene-bvRMA01_001089 | lp25 | gene | delta | Pfam | protein_matc | 39 | 163 | 1.1E-40 | + | . | Target=null 39 163;ID=match$2251_39_163;signature_desc=Borrelia lipoprotein;Name=PF00921;status=T;Dbxref="InterPro:IPR000680" |
| gene-bvRMA01_001090 | lp25 | gene | alpha | Pfam | protein_matc | 52 | 344 | 5.4E-69 | + | . | Target=null 52 344;ID=match$295_52_344;signature_desc=Borrelia lipoprotein;Name=PF00921;status=T;Dbxref="InterPro:IPR000680" |
| gene-bvRMA01_001092 | lp25 | gene | delta | Pfam | protein_matc | 48 | 325 | 2.9E-89 | + | . | Target=null 48 325;ID=match$1979_48_325;signature_desc=Borrelia lipoprotein;Name=PF00921;status=T;Dbxref="InterPro:IPR000680" |
| gene-bvRMA01_001096 | lp25 | gene | delta | Pfam | protein_matc | 9 | 61 | 2.8E-18 | + | . | Target=null 9 61;ID=match$2624_9_61;signature_desc=Borrelia lipoprotein;Name=PF00921;status=T;Dbxref="InterPro:IPR000680" |
| gene-bvRMA01_001102 | lp25 | gene | gamma | Pfam | protein_matc | 6 | 42 | 0.0000015 | + | . | Target=null 6 42;ID=match$367_6_42;signature_desc=Borrelia lipoprotein;Name=PF00921;status=T;Dbxref="InterPro:IPR000680" |
| gene-bvRMA01_001107 | lp30 | gene | beta | Pfam | protein_matc | 45 | 326 | 7.5E-45 | + | . | Target=null 45 326;ID=match$2712_45_326;signature_desc=Borrelia lipoprotein;Name=PF00921;status=T;Dbxref="InterPro:IPR000680" |
| gene-bvRMA01_001108 | lp30 | gene | gamma | Pfam | protein_matc | 44 | 319 | 1.8E-85 | + | . | Target=null 44 319;ID=match$8_44_319;signature_desc=Borrelia lipoprotein;Name=PF00921;status=T;Dbxref="InterPro:IPR000680" |
| gene-bvRMA01_001110 | lp30 | gene | gamma | Pfam | protein_matc | 34 | 312 | 4.9E-90 | + | . | Target=null 34 312;ID=match$1078_34_312;signature_desc=Borrelia lipoprotein;Name=PF00921;status=T;Dbxref="InterPro:IPR000680" |
| gene-bvRMA01_001120 | lp30 | gene | delta | Pfam | protein_matc | 45 | 330 | 5.5E-88 | + | . | Target=null 45 330;ID=match$2261_45_330;signature_desc=Borrelia lipoprotein;Name=PF00921;status=T;Dbxref="InterPro:IPR000680" |
| gene-bvRMA01_001122 | lp30 | pseudo | alpha | Pfam | protein_matc | 7 | 185 | 1.4E-32 | + | . | Target=null 7 185;ID=match$1297_7_185;signature_desc=Borrelia lipoprotein;Name=PF00921;status=T;Dbxref="InterPro:IPR000680" |
| gene-bvRMA01_001122 | lp30 | pseudo | | Pfam | protein_matc | 1 | 117 | 3E-33 | + | . | Target=null 1 117;ID=match$2150_1_117;signature_desc=Borrelia lipoprotein;Name=PF00921;status=T;Dbxref="InterPro:IPR000680" |
| gene-bvRMA01_001125 | lp30 | pseudo | beta | Pfam | protein_matc | 51 | 100 | 2.3E-08 | + | . | Target=null 51 100;ID=match$2674_51_100;signature_desc=Borrelia lipoprotein;Name=PF00921;status=T;Dbxref="InterPro:IPR000680" |
| gene-bvRMA01_001129 | lp30 | gene | alpha | Pfam | protein_matc | 7 | 299 | 3.3E-69 | + | . | Target=null 7 299;ID=match$1634_7_299;signature_desc=Borrelia lipoprotein;Name=PF00921;status=T;Dbxref="InterPro:IPR000680" |
| gene-bvRMA01_001130 | lp30 | gene | delta | Pfam | protein_matc | 39 | 163 | 1.1E-40 | + | . | Target=null 39 163;ID=match$2249_39_163;signature_desc=Borrelia lipoprotein;Name=PF00921;status=T;Dbxref="InterPro:IPR000680" |
| gene-bvRMA01_001131 | lp30 | gene | gamma | Pfam | protein_matc | 34 | 309 | 2E-86 | + | . | Target=null 34 309;ID=match$333_34_309;signature_desc=Borrelia lipoprotein;Name=PF00921;status=T;Dbxref="InterPro:IPR000680" |
| gene-bvRMA01_001132 | lp30 | gene | alpha | Pfam | protein_matc | 52 | 343 | 2E-70 | + | . | Target=null 52 343;ID=match$997_52_343;signature_desc=Borrelia lipoprotein;Name=PF00921;status=T;Dbxref="InterPro:IPR000680" |
| gene-bvRMA01_001138 | lp31 | gene | beta | Pfam | protein_matc | 45 | 326 | 7.5E-45 | + | . | Target=null 45 326;ID=match$2714_45_326;signature_desc=Borrelia lipoprotein;Name=PF00921;status=T;Dbxref="InterPro:IPR000680" |
| gene-bvRMA01_001139 | lp31 | gene | gamma | Pfam | protein_matc | 34 | 313 | 1.7E-87 | + | . | Target=null 34 313;ID=match$1195_34_313;signature_desc=Borrelia lipoprotein;Name=PF00921;status=T;Dbxref="InterPro:IPR000680" |
| gene-bvRMA01_001140 | lp31 | gene | delta | Pfam | protein_matc | 45 | 306 | 5.5E-78 | + | . | Target=null 45 306;ID=match$1081_45_306;signature_desc=Borrelia lipoprotein;Name=PF00921;status=T;Dbxref="InterPro:IPR000680" |
| gene-bvRMA01_001141 | lp31 | gene | beta | Pfam | protein_matc | 34 | 319 | 6.1E-46 | + | . | Target=null 34 319;ID=match$2746_34_319;signature_desc=Borrelia lipoprotein;Name=PF00921;status=T;Dbxref="InterPro:IPR000680" |
| gene-bvRMA01_001143 | lp31 | gene | gamma | Pfam | protein_matc | 31 | 310 | 1.8E-78 | + | . | Target=null 31 310;ID=match$2416_31_310;signature_desc=Borrelia lipoprotein;Name=PF00921;status=T;Dbxref="InterPro:IPR000680" |
| gene-bvRMA01_001159 | lp31 | pseudo | gamma | Pfam | protein_matc | 1 | 162 | 1E-34 | + | . | Target=null 1 162;ID=match$1208_1_162;signature_desc=Borrelia lipoprotein;Name=PF00921;status=T;Dbxref="InterPro:IPR000680" |
| gene-bvRMA01_001159 | lp31 | pseudo | | Pfam | protein_matc | 160 | 232 | 1.2E-25 | + | . | Target=null 160 232;ID=match$1208_160_232;signature_desc=Borrelia lipoprotein;Name=PF00921;status=T;Dbxref="InterPro:IPR000680" |
| gene-bvRMA01_001161 | lp31 | pseudo | alpha | Pfam | protein_matc | 3 | 75 | 3.5E-10 | + | . | Target=null 3 75;ID=match$42_3_75;signature_desc=Borrelia lipoprotein;Name=PF00921;status=T;Dbxref="InterPro:IPR000680" |
| gene-bvRMA01_001161 | lp31 | pseudo | | Pfam | protein_matc | 113 | 168 | 2.8E-10 | + | . | Target=null 113 168;ID=match$964_113_168;signature_desc=Borrelia lipoprotein;Name=PF00921;status=T;Dbxref="InterPro:IPR000680" |
| gene-bvRMA01_001161 | lp31 | pseudo | | Pfam | protein_matc | 7 | 125 | 5.4E-15 | + | . | Target=null 7 125;ID=match$964_7_125;signature_desc=Borrelia lipoprotein;Name=PF00921;status=T;Dbxref="InterPro:IPR000680" |
| gene-bvRMA01_001162 | lp31 | gene | beta | Pfam | protein_matc | 14 | 197 | 3E-14 | + | . | Target=null 14 197;ID=match$2345_14_197;signature_desc=Borrelia lipoprotein;Name=PF00921;status=T;Dbxref="InterPro:IPR000680" |
| gene-bvRMA01_001162 | lp31 | gene | | Pfam | protein_matc | 198 | 383 | 2.2E-29 | + | . | Target=null 198 383;ID=match$2345_198_383;signature_desc=Borrelia lipoprotein;Name=PF00921;status=T;Dbxref="InterPro:IPR000680" |
| gene-bvRMA01_001165 | lp31 | gene | gamma | Pfam | protein_matc | 45 | 326 | 2.2E-90 | + | . | Target=null 45 326;ID=match$545_45_326;signature_desc=Borrelia lipoprotein;Name=PF00921;status=T;Dbxref="InterPro:IPR000680" |
| gene-bvRMA01_001166 | lp31 | pseudo | alpha | Pfam | protein_matc | 58 | 272 | 3.8E-39 | + | . | Target=null 58 272;ID=match$2557_58_272;signature_desc=Borrelia lipoprotein;Name=PF00921;status=T;Dbxref="InterPro:IPR000680" |
| gene-bvRMA01_001166 | lp31 | pseudo | | Pfam | protein_matc | 13 | 95 | 7.6E-30 | + | . | Target=null 13 95;ID=match$1449_13_95;signature_desc=Borrelia lipoprotein;Name=PF00921;status=T;Dbxref="InterPro:IPR000680" |
| gene-bvRMA01_001167 | lp31 | gene | beta | Pfam | protein_matc | 55 | 342 | 8.2E-48 | + | . | Target=null 55 342;ID=match$657_55_342;signature_desc=Borrelia lipoprotein;Name=PF00921;status=T;Dbxref="InterPro:IPR000680" |
| gene-bvRMA01_001197 | lp35 | gene | gamma | Pfam | protein_matc | 44 | 319 | 1.8E-85 | + | . | Target=null 44 319;ID=match$6_44_319;signature_desc=Borrelia lipoprotein;Name=PF00921;status=T;Dbxref="InterPro:IPR000680" |
| gene-bvRMA01_001198 | lp35 | gene | beta | Pfam | protein_matc | 45 | 326 | 7.5E-45 | + | . | Target=null 45 326;ID=match$2710_45_326;signature_desc=Borrelia lipoprotein;Name=PF00921;status=T;Dbxref="InterPro:IPR000680" |
| gene-bvRMA01_001229 | lp37 | gene | alpha | Pfam | protein_matc | 7 | 299 | 3.3E-69 | + | . | Target=null 7 299;ID=match$1632_7_299;signature_desc=Borrelia lipoprotein;Name=PF00921;status=T;Dbxref="InterPro:IPR000680" |
| gene-bvRMA01_001230 | lp37 | gene | delta | Pfam | protein_matc | 39 | 163 | 1.1E-40 | + | . | Target=null 39 163;ID=match$2253_39_163;signature_desc=Borrelia lipoprotein;Name=PF00921;status=T;Dbxref="InterPro:IPR000680" |
| gene-bvRMA01_001231 | lp37 | gene | gamma | Pfam | protein_matc | 34 | 309 | 2E-86 | + | . | Target=null 34 309;ID=match$337_34_309;signature_desc=Borrelia lipoprotein;Name=PF00921;status=T;Dbxref="InterPro:IPR000680" |
| gene-bvRMA01_001232 | lp37 | gene | alpha | Pfam | protein_matc | 52 | 343 | 2E-70 | + | . | Target=null 52 343;ID=match$1001_52_343;signature_desc=Borrelia lipoprotein;Name=PF00921;status=T;Dbxref="InterPro:IPR000680" |

| Vsp | replicon | gene/pseudo | | | | | | | | | |
|---|---|---|---|---|---|---|---|---|---|---|---|
| gene-bt91E135_001351 | lp38 | gene | | Pfam | protein_matc | 39 | 211 | 4.9E-48 | + | . | Target=null 39 211;ID=match$1758_39_211;signature_desc=Lipoprotein;Name=PF01441;status=T;Dbxref="InterPro:IPR001800" |
| gene-bt91E135_001353 | lp38 | gene | | Pfam | protein_matc | 29 | 194 | 1.2E-57 | + | . | Target=null 29 194;ID=match$2858_29_194;signature_desc=Lipoprotein;Name=PF01441;status=T;Dbxref="InterPro:IPR001800" |
| gene-bt91E135_001361 | lp38 | gene | | Pfam | protein_matc | 30 | 196 | 2.4E-39 | + | . | Target=null 30 196;ID=match$3081_30_196;signature_desc=Lipoprotein;Name=PF01441;status=T;Dbxref="InterPro:IPR001800" |
| gene-bt91E135_001369 | lp38 | gene | | Pfam | protein_matc | 2 | 170 | 2.4E-52 | + | . | Target=null 2 170;ID=match$2165_2_170;signature_desc=Lipoprotein;Name=PF01441;status=T;Dbxref="InterPro:IPR001800" |
| gene-bt91E135_001372 | lp38 | gene | | Pfam | protein_matc | 38 | 206 | 9.4E-61 | + | . | Target=null 38 206;ID=match$456_38_206;signature_desc=Lipoprotein;Name=PF01441;status=T;Dbxref="InterPro:IPR001800" |
| gene-bt91E135_001407 | lp40 | gene | | Pfam | protein_matc | 39 | 211 | 4.9E-48 | + | . | Target=null 39 211;ID=match$1756_39_211;signature_desc=Lipoprotein;Name=PF01441;status=T;Dbxref="InterPro:IPR001800" |
| gene-bt91E135_001429 | lp43 | pseudo | | Pfam | protein_matc | 23 | 159 | 5.6E-51 | + | . | Target=null 23 159;ID=match$1807_23_159;signature_desc=Lipoprotein;Name=PF01441;status=T;Dbxref="InterPro:IPR001800" |
| gene-bt91E135_001442 | lp43 | gene | | Pfam | protein_matc | 38 | 208 | 9.3E-50 | + | . | Target=null 38 208;ID=match$2569_38_208;signature_desc=Lipoprotein;Name=PF01441;status=T;Dbxref="InterPro:IPR001800" |
| gene-bt91E135_001447 | lp43 | gene | | Pfam | protein_matc | 38 | 204 | 1.2E-56 | + | . | Target=null 38 204;ID=match$568_38_204;signature_desc=Lipoprotein;Name=PF01441;status=T;Dbxref="InterPro:IPR001800" |
| gene-bt91E135_001469 | lp48 | gene | | Pfam | protein_matc | 38 | 205 | 2.2E-55 | + | . | Target=null 38 205;ID=match$452_38_205;signature_desc=Lipoprotein;Name=PF01441;status=T;Dbxref="InterPro:IPR001800" |
| gene-bt91E135_001473 | lp48 | gene | | Pfam | protein_matc | 37 | 206 | 2.8E-54 | + | . | Target=null 37 206;ID=match$2827_37_206;signature_desc=Lipoprotein;Name=PF01441;status=T;Dbxref="InterPro:IPR001800" |
| gene-bt91E135_001475 | lp48 | gene | | Pfam | protein_matc | 38 | 207 | 5.4E-53 | + | . | Target=null 38 207;ID=match$453_38_207;signature_desc=Lipoprotein;Name=PF01441;status=T;Dbxref="InterPro:IPR001800" |
| gene-bt91E135_001480 | lp48 | gene | | Pfam | protein_matc | 38 | 206 | 1.1E-59 | + | . | Target=null 38 206;ID=match$465_38_206;signature_desc=Lipoprotein;Name=PF01441;status=T;Dbxref="InterPro:IPR001800" |
| gene-bt91E135_001483 | lp48 | gene | | Pfam | protein_matc | 38 | 212 | 1.8E-47 | + | . | Target=null 38 212;ID=match$1315_38_212;signature_desc=Lipoprotein;Name=PF01441;status=T;Dbxref="InterPro:IPR001800" |
| gene-bt91E135_001525 | lp53 | gene | | Pfam | protein_matc | 39 | 205 | 1.2E-56 | + | . | Target=null 39 205;ID=match$508_39_205;signature_desc=Lipoprotein;Name=PF01441;status=T;Dbxref="InterPro:IPR001800" |
| gene-bt91E135_001536 | lp53 | pseudo | | Pfam | protein_matc | 29 | 106 | 4.4E-31 | + | . | Target=null 29 106;ID=match$1622_29_106;signature_desc=Lipoprotein;Name=PF01441;status=T;Dbxref="InterPro:IPR001800" |
| gene-bt91E135_001536 | lp53 | pseudo | | Pfam | protein_matc | 3 | 92 | 1.6E-25 | + | . | Target=null 3 92;ID=match$314_3_92;signature_desc=Lipoprotein;Name=PF01441;status=T;Dbxref="InterPro:IPR001800" |
| gene-bt91E135_001538 | lp53 | gene | | Pfam | protein_matc | 38 | 208 | 1.5E-47 | + | . | Target=null 38 208;ID=match$1775_38_208;signature_desc=Lipoprotein;Name=PF01441;status=T;Dbxref="InterPro:IPR001800" |
| gene-bt91E135_001542 | lp53 | gene | | Pfam | protein_matc | 38 | 206 | 2.5E-58 | + | . | Target=null 38 206;ID=match$1095_38_206;signature_desc=Lipoprotein;Name=PF01441;status=T;Dbxref="InterPro:IPR001800" |

| Vlp | | | Subfamily | | | | | | | | |
|---|---|---|---|---|---|---|---|---|---|---|---|
| gene-bt91E135_001241 | lp19 | gene | gamma | Pfam | protein_matc | 51 | 331 | 2.5E-88 | + | . | Target=null 51 331;ID=match$2504_51_331;signature_desc=Borrelia lipoprotein;Name=PF00921;status=T;Dbxref="InterPro:IPR000680" |
| gene-bt91E135_001242 | lp19 | gene | alpha | Pfam | protein_matc | 41 | 341 | 4.9E-68 | + | . | Target=null 41 341;ID=match$2697_41_341;signature_desc=Borrelia lipoprotein;Name=PF00921;status=T;Dbxref="InterPro:IPR000680" |
| gene-bt91E135_001243_ | lp19 | gene | beta | Pfam | protein_matc | 20 | 304 | 2.1E-49 | + | . | Target=null 20 304;ID=match$762_20_304;signature_desc=Borrelia lipoprotein;Name=PF00921;status=T;Dbxref="InterPro:IPR000680" |
| gene-bt91E135_001308 | lp36 | pseudo | beta | Pfam | protein_matc | 16 | 304 | 3.2E-47 | + | . | Target=null 16 304;ID=match$1629_16_304;signature_desc=Borrelia lipoprotein;Name=PF00921;status=T;Dbxref="InterPro:IPR000680" |
| gene-bt91E135_001309 | lp36 | gene | alpha | Pfam | protein_matc | 58 | 355 | 2.2E-71 | + | . | Target=null 58 355;ID=match$2642_58_355;signature_desc=Borrelia lipoprotein;Name=PF00921;status=T;Dbxref="InterPro:IPR000680" |
| gene-bt91E135_001310 | lp36 | gene | beta | Pfam | protein_matc | 20 | 309 | 2.4E-49 | + | . | Target=null 20 309;ID=match$2371_20_309;signature_desc=Borrelia lipoprotein;Name=PF00921;status=T;Dbxref="InterPro:IPR000680" |
| gene-bt91E135_001311 | lp36 | gene | alpha | Pfam | protein_matc | 57 | 345 | 1E-67 | + | . | Target=null 57 345;ID=match$1838_57_345;signature_desc=Borrelia lipoprotein;Name=PF00921;status=T;Dbxref="InterPro:IPR000680" |
| gene-bt91E135_001312 | lp36 | gene | alpha | Pfam | protein_matc | 57 | 352 | 9.3E-71 | + | . | Target=null 57 352;ID=match$2543_57_352;signature_desc=Borrelia lipoprotein;Name=PF00921;status=T;Dbxref="InterPro:IPR000680" |
| gene-bt91E135_001314 | lp36 | gene | beta | Pfam | protein_matc | 55 | 338 | 7.4E-47 | + | . | Target=null 55 338;ID=match$1554_55_338;signature_desc=Borrelia lipoprotein;Name=PF00921;status=T;Dbxref="InterPro:IPR000680" |
| gene-bt91E135_001315 | lp36 | pseudo | beta | Pfam | protein_matc | 23 | 314 | 7.4E-47 | + | . | Target=null 23 314;ID=match$713_23_314;signature_desc=Borrelia lipoprotein;Name=PF00921;status=T;Dbxref="InterPro:IPR000680" |
| gene-bt91E135_001316 | lp36 | gene | alpha | Pfam | protein_matc | 58 | 355 | 1.4E-69 | + | . | Target=null 58 355;ID=match$2656_58_355;signature_desc=Borrelia lipoprotein;Name=PF00921;status=T;Dbxref="InterPro:IPR000680" |
| gene-bt91E135_001319 | lp36 | gene | beta | Pfam | protein_matc | 20 | 302 | 2.1E-44 | + | . | Target=null 20 302;ID=match$630_20_302;signature_desc=Borrelia lipoprotein;Name=PF00921;status=T;Dbxref="InterPro:IPR000680" |
| gene-bt91E135_001320 | lp36 | gene | alpha | Pfam | protein_matc | 57 | 353 | 2E-67 | + | . | Target=null 57 353;ID=match$2430_57_353;signature_desc=Borrelia lipoprotein;Name=PF00921;status=T;Dbxref="InterPro:IPR000680" |
| gene-bt91E135_001323 | lp36 | gene | gamma | Pfam | protein_matc | 34 | 317 | 1.1E-87 | + | . | Target=null 34 317;ID=match$2998_34_317;signature_desc=Borrelia lipoprotein;Name=PF00921;status=T;Dbxref="InterPro:IPR000680" |
| gene-bt91E135_001326 | lp36 | gene | delta | Pfam | protein_matc | 44 | 328 | 5.4E-85 | + | . | Target=null 44 328;ID=match$359_44_328;signature_desc=Borrelia lipoprotein;Name=PF00921;status=T;Dbxref="InterPro:IPR000680" |
| gene-bt91E135_001331 | lp36 | gene | gamma | Pfam | protein_matc | 9 | 50 | 1.5E-10 | + | . | Target=null 9 50;ID=match$1170_9_50;signature_desc=Borrelia lipoprotein;Name=PF00921;status=T;Dbxref="InterPro:IPR000680" |
| gene-bt91E135_001336 | lp36 | pseudo | alpha | Pfam | protein_matc | 45 | 121 | 7.1E-14 | + | . | Target=null 45 121;ID=match$1194_45_121;signature_desc=Borrelia lipoprotein;Name=PF00921;status=T;Dbxref="InterPro:IPR000680" |
| gene-bt91E135_001336 | lp36 | pseudo | | Pfam | protein_matc | 11 | 81 | 5.4E-07 | + | . | Target=null 11 81;ID=match$2369_11_81;signature_desc=Borrelia lipoprotein;Name=PF00921;status=T;Dbxref="InterPro:IPR000680" |
| gene-bt91E135_001336 | lp36 | pseudo | | Pfam | protein_matc | 6 | 129 | 7.3E-32 | + | . | Target=null 6 129;ID=match$2971_6_129;signature_desc=Borrelia lipoprotein;Name=PF00921;status=T;Dbxref="InterPro:IPR000680" |
| gene-bt91E135_001338 | lp36 | pseudo | alpha | Pfam | protein_matc | 8 | 72 | 3.8E-13 | + | . | Target=null 8 72;ID=match$2502_8_72;signature_desc=Borrelia lipoprotein;Name=PF00921;status=T;Dbxref="InterPro:IPR000680" |
| gene-bt91E135_001338 | lp36 | pseudo | | Pfam | protein_matc | 7 | 107 | 2.9E-16 | + | . | Target=null 7 107;ID=match$1214_7_107;signature_desc=Borrelia lipoprotein;Name=PF00921;status=T;Dbxref="InterPro:IPR000680" |
| gene-bt91E135_001339 | lp36 | pseudo | beta | Pfam | protein_matc | 2 | 40 | 0.000038 | + | . | Target=null 2 40;ID=match$1906_2_40;signature_desc=Borrelia lipoprotein;Name=PF00921;status=T;Dbxref="InterPro:IPR000680" |
| gene-bt91E135_001341 | lp36 | gene | beta | Pfam | protein_matc | 192 | 445 | 1.7E-29 | + | . | Target=null 192 445;ID=match$2469_192_445;signature_desc=Borrelia lipoprotein;Name=PF00921;status=T;Dbxref="InterPro:IPR000680" |
| gene-bt91E135_001345 | lp38 | gene | beta | Pfam | protein_matc | 136 | 257 | 3E-26 | + | . | Target=null 136 257;ID=match$205_136_257;signature_desc=Borrelia lipoprotein;Name=PF00921;status=T;Dbxref="InterPro:IPR000680" |
| gene-bt91E135_001345 | lp38 | gene | | Pfam | protein_matc | 11 | 136 | 3.8E-30 | + | . | Target=null 11 136;ID=match$205_11_136;signature_desc=Borrelia lipoprotein;Name=PF00921;status=T;Dbxref="InterPro:IPR000680" |
| gene-bt91E135_001346 | lp38 | gene | delta | Pfam | protein_matc | 1 | 45 | 1.7E-08 | + | . | Target=null 1 45;ID=match$652_1_45;signature_desc=Borrelia lipoprotein;Name=PF00921;status=T;Dbxref="InterPro:IPR000680" |
| gene-bt91E135_001347 | lp38 | pseudo | alpha | Pfam | protein_matc | 57 | 253 | 1.8E-39 | + | . | Target=null 57 253;ID=match$3074_57_253;signature_desc=Borrelia lipoprotein;Name=PF00921;status=T;Dbxref="InterPro:IPR000680" |
| gene-bt91E135_001347_ | lp38 | pseudo | | Pfam | protein_matc | 7 | 106 | 3.2E-28 | + | . | Target=null 7 106;ID=match$649_7_106;signature_desc=Borrelia lipoprotein;Name=PF00921;status=T;Dbxref="InterPro:IPR000680" |
| gene-bt91E135_001349 | lp38 | gene | beta | Pfam | protein_matc | 14 | 248 | 4.9E-26 | + | . | Target=null 14 248;ID=match$7_14_248;signature_desc=Borrelia lipoprotein;Name=PF00921;status=T;Dbxref="InterPro:IPR000680" |
| gene-bt91E135_001349 | lp38 | gene | | Pfam | protein_matc | 236 | 277 | 0.0000018 | + | . | Target=null 236 277;ID=match$7_236_277;signature_desc=Borrelia lipoprotein;Name=PF00921;status=T;Dbxref="InterPro:IPR000680" |
| gene-bt91E135_001350 | lp38 | gene | alpha | Pfam | protein_matc | 7 | 306 | 2.2E-70 | + | . | Target=null 7 306;ID=match$2643_7_306;signature_desc=Borrelia lipoprotein;Name=PF00921;status=T;Dbxref="InterPro:IPR000680" |
| gene-bt91E135_001352 | lp38 | gene | delta | Pfam | protein_matc | 19 | 298 | 3.2E-85 | + | . | Target=null 19 298;ID=match$1713_19_298;signature_desc=Borrelia lipoprotein;Name=PF00921;status=T;Dbxref="InterPro:IPR000680" |
| gene-bt91E135_001354 | lp38 | gene | gamma | Pfam | protein_matc | 34 | 310 | 1.8E-78 | + | . | Target=null 34 310;ID=match$1859_34_310;signature_desc=Borrelia lipoprotein;Name=PF00921;status=T;Dbxref="InterPro:IPR000680" |
| gene-bt91E135_001356 | lp38 | gene | delta | Pfam | protein_matc | 14 | 81 | 1.4E-22 | + | . | Target=null 14 81;ID=match$2259_14_81;signature_desc=Borrelia lipoprotein;Name=PF00921;status=T;Dbxref="InterPro:IPR000680" |
| gene-bt91E135_001358 | lp38 | gene | beta | Pfam | protein_matc | 59 | 341 | 6.8E-46 | + | . | Target=null 59 341;ID=match$795_59_341;signature_desc=Borrelia lipoprotein;Name=PF00921;status=T;Dbxref="InterPro:IPR000680" |
| gene-bt91E135_001360 | lp38 | gene | delta | Pfam | protein_matc | 57 | 107 | 3.4E-16 | + | . | Target=null 57 107;ID=match$2388_57_107;signature_desc=Borrelia lipoprotein;Name=PF00921;status=T;Dbxref="InterPro:IPR000680" |
| gene-bt91E135_001360 | lp38 | gene | | Pfam | protein_matc | 8 | 58 | 3.6E-14 | + | . | Target=null 8 58;ID=match$2388_8_58;signature_desc=Borrelia lipoprotein;Name=PF00921;status=T;Dbxref="InterPro:IPR000680" |
| gene-bt91E135_001362 | lp38 | gene | delta | Pfam | protein_matc | 44 | 325 | 5.2E-86 | + | . | Target=null 44 325;ID=match$603_44_325;signature_desc=Borrelia lipoprotein;Name=PF00921;status=T;Dbxref="InterPro:IPR000680" |
| gene-bt91E135_001365 | lp38 | gene | beta | Pfam | protein_matc | 20 | 307 | 2.4E-47 | + | . | Target=null 20 307;ID=match$1334_20_307;signature_desc=Borrelia lipoprotein;Name=PF00921;status=T;Dbxref="InterPro:IPR000680" |
| gene-bt91E135_001366 | lp38 | gene | alpha | Pfam | protein_matc | 14 | 55 | 1.9E-09 | + | . | Target=null 14 55;ID=match$1656_14_55;signature_desc=Borrelia lipoprotein;Name=PF00921;status=T;Dbxref="InterPro:IPR000680" |
| gene-bt91E135_001367 | lp38 | gene | delta | Pfam | protein_matc | 34 | 317 | 1.6E-82 | + | . | Target=null 34 317;ID=match$2537_34_317;signature_desc=Borrelia lipoprotein;Name=PF00921;status=T;Dbxref="InterPro:IPR000680" |
| gene-bt91E135_001368 | lp38 | gene | delta | Pfam | protein_matc | 8 | 292 | 7.3E-87 | + | . | Target=null 8 292;ID=match$375_8_292;signature_desc=Borrelia lipoprotein;Name=PF00921;status=T;Dbxref="InterPro:IPR000680" |
| gene-bt91E135_001370 | lp38 | gene | beta | Pfam | protein_matc | 14 | 297 | 2.1E-47 | + | . | Target=null 14 297;ID=match$641_14_297;signature_desc=Borrelia lipoprotein;Name=PF00921;status=T;Dbxref="InterPro:IPR000680" |
| gene-bt91E135_001371 | lp38 | gene | alpha | Pfam | protein_matc | 50 | 272 | 1.1E-36 | + | . | Target=null 50 272;ID=match$1141_50_272;signature_desc=Borrelia lipoprotein;Name=PF00921;status=T;Dbxref="InterPro:IPR000680" |
| gene-bt91E135_001371 | lp38 | gene | | Pfam | protein_matc | 260 | 315 | 4.1E-09 | + | . | Target=null 260 315;ID=match$1141_260_315;signature_desc=Borrelia lipoprotein;Name=PF00921;status=T;Dbxref="InterPro:IPR000680" |
| gene-bt91E135_001373 | lp38 | pseudo | gamma | Pfam | protein_matc | 6 | 47 | 3E-11 | + | . | Target=null 6 47;ID=match$1316_6_47;signature_desc=Borrelia lipoprotein;Name=PF00921;status=T;Dbxref="InterPro:IPR000680" |
| gene-bt91E135_001378 | lp38 | gene | delta | Pfam | protein_matc | 42 | 76 | 5.7E-09 | + | . | Target=null 42 76;ID=match$2975_42_76;signature_desc=Borrelia lipoprotein;Name=PF00921;status=T;Dbxref="InterPro:IPR000680" |
| gene-bt91E135_001408 | lp40 | gene | alpha | Pfam | protein_matc | 7 | 306 | 2.2E-70 | + | . | Target=null 7 306;ID=match$2645_7_306;signature_desc=Borrelia lipoprotein;Name=PF00921;status=T;Dbxref="InterPro:IPR000680" |
| gene-bt91E135_001409 | lp40 | gene | beta | Pfam | protein_matc | 14 | 248 | 4.9E-26 | + | . | Target=null 14 248;ID=match$5_14_248;signature_desc=Borrelia lipoprotein;Name=PF00921;status=T;Dbxref="InterPro:IPR000680" |
| gene-bt91E135_001409 | lp40 | gene | | Pfam | protein_matc | 236 | 277 | 0.0000018 | + | . | Target=null 236 277;ID=match$5_236_277;signature_desc=Borrelia lipoprotein;Name=PF00921;status=T;Dbxref="InterPro:IPR000680" |
| gene-bt91E135_001412 | lp40 | gene | alpha | Pfam | protein_matc | 57 | 355 | 6.2E-72 | + | . | Target=null 57 355;ID=match$1957_57_355;signature_desc=Borrelia lipoprotein;Name=PF00921;status=T;Dbxref="InterPro:IPR000680" |
| gene-bt91E135_001413 | lp40 | gene | delta | Pfam | protein_matc | 1 | 45 | 1.7E-08 | + | . | Target=null 1 45;ID=match$650_1_45;signature_desc=Borrelia lipoprotein;Name=PF00921;status=T;Dbxref="InterPro:IPR000680" |
| gene-bt91E135_001414_ | lp40 | gene | beta | Pfam | protein_matc | 136 | 257 | 3E-26 | + | . | Target=null 136 257;ID=match$208_136_257;signature_desc=Borrelia lipoprotein;Name=PF00921;status=T;Dbxref="InterPro:IPR000680" |
| gene-bt91E135_001414 | lp40 | gene | | Pfam | protein_matc | 11 | 136 | 3.8E-30 | + | . | Target=null 11 136;ID=match$208_11_136;signature_desc=Borrelia lipoprotein;Name=PF00921;status=T;Dbxref="InterPro:IPR000680" |
| gene-bt91E135_001422 | lp43 | gene | beta | Pfam | protein_matc | 14 | 299 | 2.1E-47 | + | . | Target=null 14 299;ID=match$834_14_299;signature_desc=Borrelia lipoprotein;Name=PF00921;status=T;Dbxref="InterPro:IPR000680" |
| gene-bt91E135_001423 | lp43 | gene | alpha | Pfam | protein_matc | 49 | 217 | 1.2E-24 | + | . | Target=null 49 217;ID=match$1220_49_217;signature_desc=Borrelia lipoprotein;Name=PF00921;status=T;Dbxref="InterPro:IPR000680" |
| gene-bt91E135_001423 | lp43 | gene | | Pfam | protein_matc | 216 | 250 | 0.0000065 | + | . | Target=null 216 250;ID=match$1220_216_250;signature_desc=Borrelia lipoprotein;Name=PF00921;status=T;Dbxref="InterPro:IPR000680" |
| gene-bt91E135_001424 | lp43 | gene | delta | Pfam | protein_matc | 44 | 328 | 3.6E-85 | + | . | Target=null 44 328;ID=match$1191_44_328;signature_desc=Borrelia lipoprotein;Name=PF00921;status=T;Dbxref="InterPro:IPR000680" |
| gene-bt91E135_001425 | lp43 | gene | gamma | Pfam | protein_matc | 34 | 322 | 3.4E-93 | + | . | Target=null 34 322;ID=match$969_34_322;signature_desc=Borrelia lipoprotein;Name=PF00921;status=T;Dbxref="InterPro:IPR000680" |
| gene-bt91E135_001428 | lp43 | gene | delta | Pfam | protein_matc | 34 | 316 | 5.6E-90 | + | . | Target=null 34 316;ID=match$1399_34_316;signature_desc=Borrelia lipoprotein;Name=PF00921;status=T;Dbxref="InterPro:IPR000680" |
| gene-bt91E135_001429 | lp43 | pseudo | gamma | Pfam | protein_matc | 10 | 86 | 6E-25 | + | . | Target=null 10 86;ID=match$42_10_86;signature_desc=Borrelia lipoprotein;Name=PF00921;status=T;Dbxref="InterPro:IPR000680" |
| gene-bt91E135_001440 | lp43 | gene | alpha | Pfam | protein_matc | 51 | 344 | 4.5E-67 | + | . | Target=null 51 344;ID=match$1591_51_344;signature_desc=Borrelia lipoprotein;Name=PF00921;status=T;Dbxref="InterPro:IPR000680" |
| gene-bt91E135_001441 | lp43 | gene | beta | Pfam | protein_matc | 20 | 306 | 2.4E-45 | + | . | Target=null 20 306;ID=match$2769_20_306;signature_desc=Borrelia lipoprotein;Name=PF00921;status=T;Dbxref="InterPro:IPR000680" |
| gene-bt91E135_001443 | lp43 | gene | delta | Pfam | protein_matc | 8 | 292 | 1.9E-87 | + | . | Target=null 8 292;ID=match$2837_8_292;signature_desc=Borrelia lipoprotein;Name=PF00921;status=T;Dbxref="InterPro:IPR000680" |
| gene-bt91E135_001444 | lp43 | gene | gamma | Pfam | protein_matc | 34 | 319 | 6.2E-85 | + | . | Target=null 34 319;ID=match$1512_34_319;signature_desc=Borrelia lipoprotein;Name=PF00921;status=T;Dbxref="InterPro:IPR000680" |
| gene-bt91E135_001446 | lp43 | gene | delta | Pfam | protein_matc | 34 | 318 | 1.7E-86 | + | . | Target=null 34 318;ID=match$2852_34_318;signature_desc=Borrelia lipoprotein;Name=PF00921;status=T;Dbxref="InterPro:IPR000680" |
| gene-bt91E135_001448 | lp43 | gene | delta | Pfam | protein_matc | 48 | 327 | 2.9E-86 | + | . | Target=null 48 327;ID=match$116_48_327;signature_desc=Borrelia lipoprotein;Name=PF00921;status=T;Dbxref="InterPro:IPR000680" |
| gene-bt91E135_001449 | lp43 | gene | delta | Pfam | protein_matc | 34 | 315 | 7.4E-84 | + | . | Target=null 34 315;ID=match$2460_34_315;signature_desc=Borrelia lipoprotein;Name=PF00921;status=T;Dbxref="InterPro:IPR000680" |
| gene-bt91E135_001452 | lp43 | gene | alpha | Pfam | protein_matc | 57 | 342 | 8.8E-64 | + | . | Target=null 57 342;ID=match$1379_57_342;signature_desc=Borrelia lipoprotein;Name=PF00921;status=T;Dbxref="InterPro:IPR000680" |
| gene-bt91E135_001457 | lp43 | pseudo | delta | Pfam | protein_matc | 34 | 244 | 2.7E-53 | + | . | Target=null 34 244;ID=match$1983_34_244;signature_desc=Borrelia lipoprotein;Name=PF00921;status=T;Dbxref="InterPro:IPR000680" |
| gene-bt91E135_001457 | lp43 | pseudo | | Pfam | protein_matc | 1 | 67 | 2.2E-27 | + | . | Target=null 1 67;ID=match$2260_1_67;signature_desc=Borrelia lipoprotein;Name=PF00921;status=T;Dbxref="InterPro:IPR000680" |
| gene-bt91E135_001458 | lp43 | gene | alpha | Pfam | protein_matc | 3 | 41 | 1.2E-11 | + | . | Target=null 3 41;ID=match$1729_3_41;signature_desc=Borrelia lipoprotein;Name=PF00921;status=T;Dbxref="InterPro:IPR000680" |
| gene-bt91E135_001459 | lp43 | pseudo | beta | Pfam | protein_matc | 14 | 246 | 2.3E-29 | + | . | Target=null 14 246;ID=match$2320_14_246;signature_desc=Borrelia lipoprotein;Name=PF00921;status=T;Dbxref="InterPro:IPR000680" |
| gene-bt91E135_001459 | lp43 | pseudo | | Pfam | protein_matc | 4 | 60 | 9E-16 | + | . | Target=null 4 60;ID=match$3032_4_60;signature_desc=Borrelia lipoprotein;Name=PF00921;status=T;Dbxref="InterPro:IPR000680" |
| gene-bt91E135_001468 | lp48 | pseudo | delta | Pfam | protein_matc | 4 | 52 | 0.000064 | + | . | Target=null 4 52;ID=match$1624_4_52;signature_desc=Borrelia lipoprotein;Name=PF00921;status=T;Dbxref="InterPro:IPR000680" |
| gene-bt91E135_001468 | lp48 | pseudo | | Pfam | protein_matc | 8 | 100 | 1.4E-27 | + | . | Target=null 8 100;ID=match$113_8_100;signature_desc=Borrelia lipoprotein;Name=PF00921;status=T;Dbxref="InterPro:IPR000680" |
| gene-bt91E135_001472 | lp48 | gene | gamma | Pfam | protein_matc | 1 | 282 | 2.5E-85 | + | . | Target=null 1 282;ID=match$2909_1_282;signature_desc=Borrelia lipoprotein;Name=PF00921;status=T;Dbxref="InterPro:IPR000680" |
| gene-bt91E135_001474_ | lp48 | gene | beta | Pfam | protein_matc | 20 | 305 | 2.3E-44 | + | . | Target=null 20 305;ID=match$808_20_305;signature_desc=Borrelia lipoprotein;Name=PF00921;status=T;Dbxref="InterPro:IPR000680" |
| gene-bt91E135_001476 | lp48 | gene | delta | Pfam | protein_matc | 34 | 310 | 7.6E-87 | + | . | Target=null 34 310;ID=match$2589_34_310;signature_desc=Borrelia lipoprotein;Name=PF00921;status=T;Dbxref="InterPro:IPR000680" |
| gene-bt91E135_001478 | lp48 | gene | delta | Pfam | protein_matc | 34 | 314 | 1.2E-85 | + | . | Target=null 34 314;ID=match$2897_34_314;signature_desc=Borrelia lipoprotein;Name=PF00921;status=T;Dbxref="InterPro:IPR000680" |
| gene-bt91E135_001479 | lp48 | gene | gamma | Pfam | protein_matc | 1 | 276 | 6.5E-79 | + | . | Target=null 1 276;ID=match$1911_1_276;signature_desc=Borrelia lipoprotein;Name=PF00921;status=T;Dbxref="InterPro:IPR000680" |
| gene-bt91E135_001482 | lp48 | gene | delta | Pfam | protein_matc | 147 | 189 | 5E-12 | + | . | Target=null 147 189;ID=match$867_147_189;signature_desc=Borrelia lipoprotein;Name=PF00921;status=T;Dbxref="InterPro:IPR000680" |
| gene-bt91E135_001482 | lp48 | gene | | Pfam | protein_matc | 22 | 151 | 7.5E-32 | + | . | Target=null 22 151;ID=match$867_22_151;signature_desc=Borrelia lipoprotein;Name=PF00921;status=T;Dbxref="InterPro:IPR000680" |
| gene-bt91E135_001484 | lp48 | gene | beta | Pfam | protein_matc | 20 | 309 | 3.1E-47 | + | . | Target=null 20 309;ID=match$1921_20_309;signature_desc=Borrelia lipoprotein;Name=PF00921;status=T;Dbxref="InterPro:IPR000680" |
| gene-bt91E135_001485 | lp48 | gene | alpha | Pfam | protein_matc | 57 | 345 | 1.6E-67 | + | . | Target=null 57 345;ID=match$2886_57_345;signature_desc=Borrelia lipoprotein;Name=PF00921;status=T;Dbxref="InterPro:IPR000680" |
| gene-bt91E135_001486 | lp48 | gene | delta | Pfam | protein_matc | 44 | 327 | 1.2E-76 | + | . | Target=null 44 327;ID=match$2648_44_327;signature_desc=Borrelia lipoprotein;Name=PF00921;status=T;Dbxref="InterPro:IPR000680" |
| gene-bt91E135_001487 | lp48 | gene | gamma | Pfam | protein_matc | 44 | 330 | 1.1E-90 | + | . | Target=null 44 330;ID=match$1504_44_330;signature_desc=Borrelia lipoprotein;Name=PF00921;status=T;Dbxref="InterPro:IPR000680" |
| gene-bt91E135_001506 | lp48 | gene | alpha | Pfam | protein_matc | 58 | 354 | 9.4E-70 | + | . | Target=null 58 354;ID=match$1350_58_354;signature_desc=Borrelia lipoprotein;Name=PF00921;status=T;Dbxref="InterPro:IPR000680" |
| gene-bt91E135_001507 | lp48 | pseudo | beta | Pfam | protein_matc | 55 | 136 | 1.3E-08 | + | . | Target=null 55 136;ID=match$2468_55_136;signature_desc=Borrelia lipoprotein;Name=PF00921;status=T;Dbxref="InterPro:IPR000680" |
| gene-bt91E135_001507 | lp48 | pseudo | | Pfam | protein_matc | 29 | 129 | 9.1E-24 | + | . | Target=null 29 129;ID=match$256_29_129;signature_desc=Borrelia lipoprotein;Name=PF00921;status=T;Dbxref="InterPro:IPR000680" |
| gene-bt91E135_001513 | lp53 | pseudo | beta | Pfam | protein_matc | 14 | 246 | 2.3E-29 | + | . | Target=null 14 246;ID=match$2318_14_246;signature_desc=Borrelia lipoprotein;Name=PF00921;status=T;Dbxref="InterPro:IPR000680" |
| gene-bt91E135_001513 | lp53 | pseudo | | Pfam | protein_matc | 4 | 60 | 9E-16 | + | . | Target=null 4 60;ID=match$3034_4_60;signature_desc=Borrelia lipoprotein;Name=PF00921;status=T;Dbxref="InterPro:IPR000680" |
| gene-bt91E135_001514 | lp53 | gene | alpha | Pfam | protein_matc | 3 | 41 | 1.2E-11 | + | . | Target=null 3 41;ID=match$1731_3_41;signature_desc=Borrelia lipoprotein;Name=PF00921;status=T;Dbxref="InterPro:IPR000680" |
| gene-bt91E135_001515 | lp53 | pseudo | delta | Pfam | protein_matc | 34 | 244 | 2.7E-53 | + | . | Target=null 34 244;ID=match$1981_34_244;signature_desc=Borrelia lipoprotein;Name=PF00921;status=T;Dbxref="InterPro:IPR000680" |
| gene-bt91E135_001515 | lp53 | pseudo | | Pfam | protein_matc | 5 | 53 | 8.1E-16 | + | . | Target=null 5 53;ID=match$606_5_53;signature_desc=Borrelia lipoprotein;Name=PF00921;status=T;Dbxref="InterPro:IPR000680" |
| gene-bt91E135_001520 | lp53 | gene | alpha | Pfam | protein_matc | 57 | 342 | 8.8E-64 | + | . | Target=null 57 342;ID=match$1381_57_342;signature_desc=Borrelia lipoprotein;Name=PF00921;status=T;Dbxref="InterPro:IPR000680" |
| gene-bt91E135_001523 | lp53 | gene | delta | Pfam | protein_matc | 34 | 315 | 7.4E-84 | + | . | Target=null 34 315;ID=match$2462_34_315;signature_desc=Borrelia lipoprotein;Name=PF00921;status=T;Dbxref="InterPro:IPR000680" |
| gene-bt91E135_001524 | lp53 | gene | delta | Pfam | protein_matc | 48 | 327 | 2.9E-86 | + | . | Target=null 48 327;ID=match$118_48_327;signature_desc=Borrelia lipoprotein;Name=PF00921;status=T;Dbxref="InterPro:IPR000680" |
| gene-bt91E135_001526 | lp53 | pseudo | delta | Pfam | protein_matc | 6 | 47 | 4.6E-10 | + | . | Target=null 6 47;ID=match$79_6_47;signature_desc=Borrelia lipoprotein;Name=PF00921;status=T;Dbxref="InterPro:IPR000680" |
| gene-bt91E135_001527 | lp53 | gene | delta | Pfam | protein_matc | 4 | 31 | 0.0000033 | + | . | Target=null 4 31;ID=match$371_4_31;signature_desc=Borrelia lipoprotein;Name=PF00921;status=T;Dbxref="InterPro:IPR000680" |
| gene-bt91E135_001528 | lp53 | gene | alpha | Pfam | protein_matc | 58 | 353 | 9.1E-68 | + | . | Target=null 58 353;ID=match$3091_58_353;signature_desc=Borrelia lipoprotein;Name=PF00921;status=T;Dbxref="InterPro:IPR000680" |
| gene-bt91E135_001531 | lp53 | gene | beta | Pfam | protein_matc | 48 | 330 | 2.5E-45 | + | . | Target=null 48 330;ID=match$335_48_330;signature_desc=Borrelia lipoprotein;Name=PF00921;status=T;Dbxref="InterPro:IPR000680" |
| gene-bt91E135_001534_ | lp53 | gene | beta | Pfam | protein_matc | 48 | 336 | 9.3E-47 | + | . | Target=null 48 336;ID=match$1550_48_336;signature_desc=Borrelia lipoprotein;Name=PF00921;status=T;Dbxref="InterPro:IPR000680" |
| gene-bt91E135_001535 | lp53 | gene | delta | Pfam | protein_matc | 19 | 298 | 4.8E-88 | + | . | Target=null 19 298;ID=match$866_19_298;signature_desc=Borrelia lipoprotein;Name=PF00921;status=T;Dbxref="InterPro:IPR000680" |
| gene-bt91E135_001537 | lp53 | gene | gamma | Pfam | protein_matc | 1 | 267 | 1.9E-72 | + | . | Target=null 1 267;ID=match$1486_1_267;signature_desc=Borrelia lipoprotein;Name=PF00921;status=T;Dbxref="InterPro:IPR000680" |
| gene-bt91E135_001539 | lp53 | gene | delta | Pfam | protein_matc | 44 | 335 | 1.8E-86 | + | . | Target=null 44 335;ID=match$1069_44_335;signature_desc=Borrelia lipoprotein;Name=PF00921;status=T;Dbxref="InterPro:IPR000680" |
| gene-bt91E135_001541 | lp53 | gene | delta | Pfam | protein_matc | 31 | 317 | 6.5E-88 | + | . | Target=null 31 317;ID=match$622_31_317;signature_desc=Borrelia lipoprotein;Name=PF00921;status=T;Dbxref="InterPro:IPR000680" |
| gene-bt91E135_001545 | lp53 | gene | delta | Pfam | protein_matc | 44 | 329 | 6.6E-88 | + | . | Target=null 44 329;ID=match$1681_44_329;signature_desc=Borrelia lipoprotein;Name=PF00921;status=T;Dbxref="InterPro:IPR000680" |
| gene-bt91E135_001556 | lp53 | gene | gamma | Pfam | protein_matc | 1 | 281 | 1.2E-77 | + | . | Target=null 1 281;ID=match$2289_1_281;signature_desc=Borrelia lipoprotein;Name=PF00921;status=T;Dbxref="InterPro:IPR000680" |
| gene-bt91E135_001560 | lp53 | gene | delta | Pfam | protein_matc | 34 | 313 | 5.5E-85 | + | . | Target=null 34 313;ID=match$2488_34_313;signature_desc=Borrelia lipoprotein;Name=PF00921;status=T;Dbxref="InterPro:IPR000680" |
| gene-bt91E135_001562 | lp53 | gene | alpha | Pfam | protein_matc | 58 | 351 | 4.7E-66 | + | . | Target=null 58 351;ID=match$1583_58_351;signature_desc=Borrelia lipoprotein;Name=PF00921;status=T;Dbxref="InterPro:IPR000680" |
| gene-bt91E135_001563 | lp53 | gene | beta | Pfam | protein_matc | 20 | 205 | 1.5E-22 | + | . | Target=null 20 205;ID=match$15_20_205;signature_desc=Borrelia lipoprotein;Name=PF00921;status=T;Dbxref="InterPro:IPR000680" |

| Vsp | replicon | gene/pseudo | | | | | | | | | |
|---|---|---|---|---|---|---|---|---|---|---|---|
| gene-btBTE5EL_001273 | lp28 | gene | | Pfam | protein_matc | 38 | 205 | 1.9E-58 | + | . | Target=null 38 205;ID=match$1317_38_205;signature_desc=Lipoprotein;Name=PF01441;status=T;Dbxref="InterPro:IPR001800" |
| gene-btBTE5EL_001281 | lp28 | gene | | Pfam | protein_matc | 38 | 206 | 4.4E-64 | + | . | Target=null 38 206;ID=match$3074_38_206;signature_desc=Lipoprotein;Name=PF01441;status=T;Dbxref="InterPro:IPR001800" |
| gene-btBTE5EL_001283 | lp28 | gene | | Pfam | protein_matc | 23 | 190 | 2.3E-58 | + | . | Target=null 23 190;ID=match$23_23_190;signature_desc=Lipoprotein;Name=PF01441;status=T;Dbxref="InterPro:IPR001800" |
| gene-btBTE5EL_001333 | lp32 | gene | | Pfam | protein_matc | 38 | 206 | 4E-66 | + | . | Target=null 38 206;ID=match$3062_38_206;signature_desc=Lipoprotein;Name=PF01441;status=T;Dbxref="InterPro:IPR001800" |
| gene-btBTE5EL_001336 | lp32 | gene | | Pfam | protein_matc | 38 | 206 | 2.7E-53 | + | . | Target=null 38 206;ID=match$585_38_206;signature_desc=Lipoprotein;Name=PF01441;status=T;Dbxref="InterPro:IPR001800" |
| gene-btBTE5EL_001338 | lp32 | gene | | Pfam | protein_matc | 22 | 190 | 4.8E-62 | + | . | Target=null 22 190;ID=match$1879_22_190;signature_desc=Lipoprotein;Name=PF01441;status=T;Dbxref="InterPro:IPR001800" |
| gene-btBTE5EL_001374 | lp34 | pseudo | | Pfam | protein_matc | 27 | 78 | 1E-21 | + | . | Target=null 27 78;ID=match$1346_27_78;signature_desc=Lipoprotein;Name=PF01441;status=T;Dbxref="InterPro:IPR001800" |
| gene-btBTE5EL_001376 | lp34 | gene | | Pfam | protein_matc | 39 | 210 | 3.3E-50 | + | . | Target=null 39 210;ID=match$1524_39_210;signature_desc=Lipoprotein;Name=PF01441;status=T;Dbxref="InterPro:IPR001800" |
| gene-btBTE5EL_001396 | lp34 | gene | | Pfam | protein_matc | 1 | 167 | 4.1E-60 | + | . | Target=null 1 167;ID=match$458_1_167;signature_desc=Lipoprotein;Name=PF01441;status=T;Dbxref="InterPro:IPR001800" |
| gene-btBTE5EL_001410 | lp37 | gene | | Pfam | protein_matc | 28 | 195 | 1.1E-61 | + | . | Target=null 28 195;ID=match$1079_28_195;signature_desc=Lipoprotein;Name=PF01441;status=T;Dbxref="InterPro:IPR001800" |
| gene-btBTE5EL_001430 | lp37 | gene | | Pfam | protein_matc | 28 | 195 | 1.1E-61 | + | . | Target=null 28 195;ID=match$1081_28_195;signature_desc=Lipoprotein;Name=PF01441;status=T;Dbxref="InterPro:IPR001800" |
| gene-btBTE5EL_001446 | lp42 | gene | | Pfam | protein_matc | 38 | 207 | 3.8E-53 | + | . | Target=null 38 207;ID=match$1866_38_207;signature_desc=Lipoprotein;Name=PF01441;status=T;Dbxref="InterPro:IPR001800" |
| gene-btBTE5EL_001448 | lp42 | gene | | Pfam | protein_matc | 38 | 207 | 1.7E-59 | + | . | Target=null 38 207;ID=match$370_38_207;signature_desc=Lipoprotein;Name=PF01441;status=T;Dbxref="InterPro:IPR001800" |
| gene-btBTE5EL_001451 | lp42 | gene | | Pfam | protein_matc | 23 | 188 | 2.2E-56 | + | . | Target=null 23 188;ID=match$2853_23_188;signature_desc=Lipoprotein;Name=PF01441;status=T;Dbxref="InterPro:IPR001800" |
| gene-btBTE5EL_001458 | lp42 | pseudo | | Pfam | protein_matc | 39 | 151 | 2E-39 | + | . | Target=null 39 151;ID=match$2909_39_151;signature_desc=Lipoprotein;Name=PF01441;status=T;Dbxref="InterPro:IPR001800" |
| gene-btBTE5EL_001458 | lp42 | pseudo | | Pfam | protein_matc | 2 | 39 | 0.00000018 | + | . | Target=null 2 39;ID=match$1850_2_39;signature_desc=Lipoprotein;Name=PF01441;status=T;Dbxref="InterPro:IPR001800" |
| gene-btBTE5EL_001467 | lp42 | gene | | Pfam | protein_matc | 38 | 206 | 1.6E-60 | + | . | Target=null 38 206;ID=match$2402_38_206;signature_desc=Lipoprotein;Name=PF01441;status=T;Dbxref="InterPro:IPR001800" |
| gene-btBTE5EL_001508 | lp45 | gene | | Pfam | protein_matc | 38 | 206 | 4.4E-64 | + | . | Target=null 38 206;ID=match$3076_38_206;signature_desc=Lipoprotein;Name=PF01441;status=T;Dbxref="InterPro:IPR001800" |
| gene-btBTE5EL_001510 | lp45 | gene | | Pfam | protein_matc | 23 | 190 | 2.3E-58 | + | . | Target=null 23 190;ID=match$25_23_190;signature_desc=Lipoprotein;Name=PF01441;status=T;Dbxref="InterPro:IPR001800" |
| gene-btBTE5EL_001530 | lp46 | gene | | Pfam | protein_matc | 38 | 201 | 2.7E-53 | + | . | Target=null 38 201;ID=match$2589_38_201;signature_desc=Lipoprotein;Name=PF01441;status=T;Dbxref="InterPro:IPR001800" |
| gene-btBTE5EL_001580 | lp49 | gene | | Pfam | protein_matc | 38 | 205 | 4E-64 | + | . | Target=null 38 205;ID=match$742_38_205;signature_desc=Lipoprotein;Name=PF01441;status=T;Dbxref="InterPro:IPR001800" |
| gene-btBTE5EL_001599 | lp49 | gene | | Pfam | protein_matc | 39 | 212 | 2.4E-47 | + | . | Target=null 39 212;ID=match$1537_39_212;signature_desc=Lipoprotein;Name=PF01441;status=T;Dbxref="InterPro:IPR001800" |
| gene-btBTE5EL_001615 | lp49 | gene | | Pfam | protein_matc | 37 | 206 | 2.2E-53 | + | . | Target=null 37 206;ID=match$1062_37_206;signature_desc=Lipoprotein;Name=PF01441;status=T;Dbxref="InterPro:IPR001800" |
| Vlp | | | Subfamily | | | | | | | | |
| gene-btBTE5EL_001272 | lp28 | pseudo | alpha | Pfam | protein_matc | 40 | 182 | 2.3E-24 | + | . | Target=null 40 182;ID=match$1502_40_182;signature_desc=Borrelia lipoprotein;Name=PF00921;status=T;Dbxref="InterPro:IPR000680" |
| gene-btBTE5EL_001272 | lp28 | pseudo | | Pfam | protein_matc | 4 | 143 | 1.4E-33 | + | . | Target=null 4 143;ID=match$496_4_143;signature_desc=Borrelia lipoprotein;Name=PF00921;status=T;Dbxref="InterPro:IPR000680" |
| gene-btBTE5EL_001275 | lp28 | pseudo | gamma | Pfam | protein_matc | 3 | 42 | 0.000065 | + | . | Target=null 3 42;ID=match$1708_3_42;signature_desc=Borrelia lipoprotein;Name=PF00921;status=T;Dbxref="InterPro:IPR000680" |
| gene-btBTE5EL_001275 | lp28 | pseudo | | Pfam | protein_matc | 50 | 161 | 2.6E-33 | + | . | Target=null 50 161;ID=match$330_50_161;signature_desc=Borrelia lipoprotein;Name=PF00921;status=T;Dbxref="InterPro:IPR000680" |
| gene-btBTE5EL_001282 | lp28 | gene | gamma | Pfam | protein_matc | 48 | 333 | 1.3E-88 | + | . | Target=null 48 333;ID=match$1927_48_333;signature_desc=Borrelia lipoprotein;Name=PF00921;status=T;Dbxref="InterPro:IPR000680" |
| gene-btBTE5EL_001284 | lp28 | pseudo | delta | Pfam | protein_matc | 32 | 317 | 3E-83 | + | . | Target=null 32 317;ID=match$1794_32_317;signature_desc=Borrelia lipoprotein;Name=PF00921;status=T;Dbxref="InterPro:IPR000680" |
| gene-btBTE5EL_001286 | lp28 | gene | delta | Pfam | protein_matc | 44 | 335 | 5E-87 | + | . | Target=null 44 335;ID=match$1160_44_335;signature_desc=Borrelia lipoprotein;Name=PF00921;status=T;Dbxref="InterPro:IPR000680" |
| gene-btBTE5EL_001287 | lp28 | gene | beta | Pfam | protein_matc | 55 | 343 | 2.7E-47 | + | . | Target=null 55 343;ID=match$2806_55_343;signature_desc=Borrelia lipoprotein;Name=PF00921;status=T;Dbxref="InterPro:IPR000680" |
| gene-btBTE5EL_001289 | lp28 | gene | alpha | Pfam | protein_matc | 59 | 343 | 1.3E-63 | + | . | Target=null 59 343;ID=match$1714_59_343;signature_desc=Borrelia lipoprotein;Name=PF00921;status=T;Dbxref="InterPro:IPR000680" |
| gene-btBTE5EL_001290 | lp28 | gene | beta | Pfam | protein_matc | 14 | 297 | 2.9E-43 | + | . | Target=null 14 297;ID=match$388_14_297;signature_desc=Borrelia lipoprotein;Name=PF00921;status=T;Dbxref="InterPro:IPR000680" |
| gene-btBTE5EL_001291 | lp28 | pseudo | gamma | Pfam | protein_matc | 34 | 115 | 6.4E-29 | + | . | Target=null 34 115;ID=match$965_34_115;signature_desc=Borrelia lipoprotein;Name=PF00921;status=T;Dbxref="InterPro:IPR000680" |
| gene-btBTE5EL_001291 | lp28 | pseudo | | Pfam | protein_matc | 4 | 203 | 2.8E-49 | + | . | Target=null 4 203;ID=match$3079_4_203;signature_desc=Borrelia lipoprotein;Name=PF00921;status=T;Dbxref="InterPro:IPR000680" |
| gene-btBTE5EL_001298 | lp30 | gene | gamma | Pfam | protein_matc | 150 | 278 | 2.1E-23 | + | . | Target=null 150 278;ID=match$3052_150_278;signature_desc=Borrelia lipoprotein;Name=PF00921;status=T;Dbxref="InterPro:IPR000680" |
| gene-btBTE5EL_001298 | lp30 | gene | | Pfam | protein_matc | 6 | 138 | 5.1E-30 | + | . | Target=null 6 138;ID=match$3052_6_138;signature_desc=Borrelia lipoprotein;Name=PF00921;status=T;Dbxref="InterPro:IPR000680" |
| gene-btBTE5EL_001301 | lp30 | gene | gamma | Pfam | protein_matc | 34 | 308 | 3.6E-77 | + | . | Target=null 34 308;ID=match$3099_34_308;signature_desc=Borrelia lipoprotein;Name=PF00921;status=T;Dbxref="InterPro:IPR000680" |
| gene-btBTE5EL_001319 | lp30 | gene | gamma | Pfam | protein_matc | 1 | 276 | 3.1E-77 | + | . | Target=null 1 276;ID=match$979_1_276;signature_desc=Borrelia lipoprotein;Name=PF00921;status=T;Dbxref="InterPro:IPR000680" |
| gene-btBTE5EL_001320 | lp30 | pseudo | beta | Pfam | protein_matc | 48 | 208 | 6.2E-16 | + | . | Target=null 48 208;ID=match$596_48_208;signature_desc=Borrelia lipoprotein;Name=PF00921;status=T;Dbxref="InterPro:IPR000680" |
| gene-btBTE5EL_001320 | lp30 | pseudo | | Pfam | protein_matc | 20 | 58 | 2.2E-09 | + | . | Target=null 20 58;ID=match$1511_20_58;signature_desc=Borrelia lipoprotein;Name=PF00921;status=T;Dbxref="InterPro:IPR000680" |
| gene-btBTE5EL_001322 | lp30 | gene | delta | Pfam | protein_matc | 44 | 329 | 1.2E-87 | + | . | Target=null 44 329;ID=match$808_44_329;signature_desc=Borrelia lipoprotein;Name=PF00921;status=T;Dbxref="InterPro:IPR000680" |
| gene-btBTE5EL_001323 | lp30 | gene | gamma | Pfam | protein_matc | 34 | 312 | 1.9E-84 | + | . | Target=null 34 312;ID=match$326_34_312;signature_desc=Borrelia lipoprotein;Name=PF00921;status=T;Dbxref="InterPro:IPR000680" |
| gene-btBTE5EL_001325 | lp30 | gene | alpha | Pfam | protein_matc | 58 | 350 | 1.8E-74 | + | . | Target=null 58 350;ID=match$2586_58_350;signature_desc=Borrelia lipoprotein;Name=PF00921;status=T;Dbxref="InterPro:IPR000680" |
| gene-btBTE5EL_001326 | lp30 | gene | beta | Pfam | protein_matc | 20 | 308 | 1.1E-50 | + | . | Target=null 20 308;ID=match$2406_20_308;signature_desc=Borrelia lipoprotein;Name=PF00921;status=T;Dbxref="InterPro:IPR000680" |
| gene-btBTE5EL_001332 | lp32 | gene | beta | Pfam | protein_matc | 20 | 309 | 5.2E-48 | + | . | Target=null 20 309;ID=match$1125_20_309;signature_desc=Borrelia lipoprotein;Name=PF00921;status=T;Dbxref="InterPro:IPR000680" |
| gene-btBTE5EL_001335 | lp32 | gene | delta | Pfam | protein_matc | 19 | 295 | 1.6E-84 | + | . | Target=null 19 295;ID=match$1289_19_295;signature_desc=Borrelia lipoprotein;Name=PF00921;status=T;Dbxref="InterPro:IPR000680" |
| gene-btBTE5EL_001337 | lp32 | pseudo | delta | Pfam | protein_matc | 4 | 282 | 2.3E-82 | + | . | Target=null 4 282;ID=match$2221_4_282;signature_desc=Borrelia lipoprotein;Name=PF00921;status=T;Dbxref="InterPro:IPR000680" |
| gene-btBTE5EL_001356 | lp32 | gene | gamma | Pfam | protein_matc | 1 | 278 | 1.2E-72 | + | . | Target=null 1 278;ID=match$2044_1_278;signature_desc=Borrelia lipoprotein;Name=PF00921;status=T;Dbxref="InterPro:IPR000680" |
| gene-btBTE5EL_001358 | lp32 | gene | alpha | Pfam | protein_matc | 58 | 354 | 7.4E-65 | + | . | Target=null 58 354;ID=match$2937_58_354;signature_desc=Borrelia lipoprotein;Name=PF00921;status=T;Dbxref="InterPro:IPR000680" |
| gene-btBTE5EL_001359 | lp32 | pseudo | beta | Pfam | protein_matc | 23 | 160 | 2.9E-16 | + | . | Target=null 23 160;ID=match$1921_23_160;signature_desc=Borrelia lipoprotein;Name=PF00921;status=T;Dbxref="InterPro:IPR000680" |
| gene-btBTE5EL_001369 | lp34 | gene | delta | Pfam | protein_matc | 44 | 202 | 9.5E-37 | + | . | Target=null 44 202;ID=match$2535_44_202;signature_desc=Borrelia lipoprotein;Name=PF00921;status=T;Dbxref="InterPro:IPR000680" |
| gene-btBTE5EL_001369 | lp34 | gene | | Pfam | protein_matc | 210 | 290 | 3.4E-28 | + | . | Target=null 210 290;ID=match$2535_210_290;signature_desc=Borrelia lipoprotein;Name=PF00921;status=T;Dbxref="InterPro:IPR000680" |
| gene-btBTE5EL_001370 | lp34 | gene | gamma | Pfam | protein_matc | 34 | 319 | 4.2E-87 | + | . | Target=null 34 319;ID=match$1446_34_319;signature_desc=Borrelia lipoprotein;Name=PF00921;status=T;Dbxref="InterPro:IPR000680" |
| gene-btBTE5EL_001371 | lp34 | gene | delta | Pfam | protein_matc | 44 | 323 | 2.4E-82 | + | . | Target=null 44 323;ID=match$468_44_323;signature_desc=Borrelia lipoprotein;Name=PF00921;status=T;Dbxref="InterPro:IPR000680" |
| gene-btBTE5EL_001373 | lp34 | gene | delta | Pfam | protein_matc | 30 | 121 | 1.5E-29 | + | . | Target=null 30 121;ID=match$2620_30_121;signature_desc=Borrelia lipoprotein;Name=PF00921;status=T;Dbxref="InterPro:IPR000680" |
| gene-btBTE5EL_001375 | lp34 | gene | beta | Pfam | protein_matc | 20 | 307 | 1.7E-49 | + | . | Target=null 20 307;ID=match$982_20_307;signature_desc=Borrelia lipoprotein;Name=PF00921;status=T;Dbxref="InterPro:IPR000680" |
| gene-btBTE5EL_001389 | lp34 | pseudo | gamma | Pfam | protein_matc | 1 | 90 | 2.1E-22 | + | . | Target=null 1 90;ID=match$48_1_90;signature_desc=Borrelia lipoprotein;Name=PF00921;status=T;Dbxref="InterPro:IPR000680" |
| gene-btBTE5EL_001389 | lp34 | pseudo | | Pfam | protein_matc | 26 | 208 | 9.2E-48 | + | . | Target=null 26 208;ID=match$2828_26_208;signature_desc=Borrelia lipoprotein;Name=PF00921;status=T;Dbxref="InterPro:IPR000680" |
| gene-btBTE5EL_001390 | lp34 | gene | beta | Pfam | protein_matc | 38 | 322 | 4.3E-44 | + | . | Target=null 38 322;ID=match$512_38_322;signature_desc=Borrelia lipoprotein;Name=PF00921;status=T;Dbxref="InterPro:IPR000680" |
| gene-btBTE5EL_001393 | lp34 | gene | alpha | Pfam | protein_matc | 53 | 348 | 2.3E-67 | + | . | Target=null 53 348;ID=match$2532_53_348;signature_desc=Borrelia lipoprotein;Name=PF00921;status=T;Dbxref="InterPro:IPR000680" |
| gene-btBTE5EL_001394 | lp34 | pseudo | beta | Pfam | protein_matc | 14 | 261 | 2.1E-34 | + | . | Target=null 14 261;ID=match$2171_14_261;signature_desc=Borrelia lipoprotein;Name=PF00921;status=T;Dbxref="InterPro:IPR000680" |
| gene-btBTE5EL_001394 | lp34 | pseudo | | Pfam | protein_matc | 2 | 39 | 2.2E-10 | + | . | Target=null 2 39;ID=match$2182_2_39;signature_desc=Borrelia lipoprotein;Name=PF00921;status=T;Dbxref="InterPro:IPR000680" |
| gene-btBTE5EL_001395 | lp34 | gene | delta | Pfam | protein_matc | 34 | 315 | 3.9E-88 | + | . | Target=null 34 315;ID=match$2099_34_315;signature_desc=Borrelia lipoprotein;Name=PF00921;status=T;Dbxref="InterPro:IPR000680" |
| gene-btBTE5EL_001397 | lp34 | pseudo | delta | Pfam | protein_matc | 8 | 166 | 2.8E-38 | + | . | Target=null 8 166;ID=match$184_8_166;signature_desc=Borrelia lipoprotein;Name=PF00921;status=T;Dbxref="InterPro:IPR000680" |
| gene-btBTE5EL_001397 | lp34 | pseudo | | Pfam | protein_matc | 11 | 84 | 2E-28 | + | . | Target=null 11 84;ID=match$247_11_84;signature_desc=Borrelia lipoprotein;Name=PF00921;status=T;Dbxref="InterPro:IPR000680" |
| gene-btBTE5EL_001402 | lp37 | gene | beta | Pfam | protein_matc | 171 | 423 | 1.2E-26 | + | . | Target=null 171 423;ID=match$240_171_423;signature_desc=Borrelia lipoprotein;Name=PF00921;status=T;Dbxref="InterPro:IPR000680" |
| gene-btBTE5EL_001404 | lp37 | gene | delta | Pfam | protein_matc | 44 | 327 | 1.4E-84 | + | . | Target=null 44 327;ID=match$1256_44_327;signature_desc=Borrelia lipoprotein;Name=PF00921;status=T;Dbxref="InterPro:IPR000680" |
| gene-btBTE5EL_001407 | lp37 | pseudo | beta | Pfam | protein_matc | 3 | 248 | 6.1E-31 | + | . | Target=null 3 248;ID=match$798_3_248;signature_desc=Borrelia lipoprotein;Name=PF00921;status=T;Dbxref="InterPro:IPR000680" |
| gene-btBTE5EL_001408 | lp37 | gene | alpha | Pfam | protein_matc | 58 | 354 | 1.9E-70 | + | . | Target=null 58 354;ID=match$3174_58_354;signature_desc=Borrelia lipoprotein;Name=PF00921;status=T;Dbxref="InterPro:IPR000680" |
| gene-btBTE5EL_001409 | lp37 | gene | delta | Pfam | protein_matc | 19 | 302 | 6.4E-88 | + | . | Target=null 19 302;ID=match$2000_19_302;signature_desc=Borrelia lipoprotein;Name=PF00921;status=T;Dbxref="InterPro:IPR000680" |
| gene-btBTE5EL_001411 | lp37 | gene | beta | Pfam | protein_matc | 20 | 303 | 6.5E-43 | + | . | Target=null 20 303;ID=match$268_20_303;signature_desc=Borrelia lipoprotein;Name=PF00921;status=T;Dbxref="InterPro:IPR000680" |
| gene-btBTE5EL_001412 | lp37 | gene | alpha | Pfam | protein_matc | 62 | 344 | 1.2E-64 | + | . | Target=null 62 344;ID=match$1265_62_344;signature_desc=Borrelia lipoprotein;Name=PF00921;status=T;Dbxref="InterPro:IPR000680" |
| gene-btBTE5EL_001413 | lp37 | gene | gamma | Pfam | protein_matc | 1 | 274 | 2.8E-79 | + | . | Target=null 1 274;ID=match$739_1_274;signature_desc=Borrelia lipoprotein;Name=PF00921;status=T;Dbxref="InterPro:IPR000680" |
| gene-btBTE5EL_001427 | lp37 | gene | gamma | Pfam | protein_matc | 1 | 274 | 2.8E-79 | + | . | Target=null 1 274;ID=match$737_1_274;signature_desc=Borrelia lipoprotein;Name=PF00921;status=T;Dbxref="InterPro:IPR000680" |
| gene-btBTE5EL_001428 | lp37 | pseudo | alpha | Pfam | protein_matc | 48 | 330 | 1E-64 | + | . | Target=null 48 330;ID=match$2292_48_330;signature_desc=Borrelia lipoprotein;Name=PF00921;status=T;Dbxref="InterPro:IPR000680" |
| gene-btBTE5EL_001429 | lp37 | gene | beta | Pfam | protein_matc | 20 | 303 | 6.5E-43 | + | . | Target=null 20 303;ID=match$270_20_303;signature_desc=Borrelia lipoprotein;Name=PF00921;status=T;Dbxref="InterPro:IPR000680" |
| gene-btBTE5EL_001431 | lp37 | gene | delta | Pfam | protein_matc | 19 | 302 | 6.4E-88 | + | . | Target=null 19 302;ID=match$1998_19_302;signature_desc=Borrelia lipoprotein;Name=PF00921;status=T;Dbxref="InterPro:IPR000680" |
| gene-btBTE5EL_001432 | lp37 | gene | alpha | Pfam | protein_matc | 58 | 354 | 1.9E-70 | + | . | Target=null 58 354;ID=match$3176_58_354;signature_desc=Borrelia lipoprotein;Name=PF00921;status=T;Dbxref="InterPro:IPR000680" |
| gene-btBTE5EL_001433 | lp37 | pseudo | beta | Pfam | protein_matc | 3 | 248 | 6.1E-31 | + | . | Target=null 3 248;ID=match$796_3_248;signature_desc=Borrelia lipoprotein;Name=PF00921;status=T;Dbxref="InterPro:IPR000680" |
| gene-btBTE5EL_001436 | lp37 | gene | delta | Pfam | protein_matc | 44 | 327 | 1.4E-84 | + | . | Target=null 44 327;ID=match$1254_44_327;signature_desc=Borrelia lipoprotein;Name=PF00921;status=T;Dbxref="InterPro:IPR000680" |
| gene-btBTE5EL_001438 | lp37 | gene | beta | Pfam | protein_matc | 171 | 423 | 1.2E-26 | + | . | Target=null 171 423;ID=match$242_171_423;signature_desc=Borrelia lipoprotein;Name=PF00921;status=T;Dbxref="InterPro:IPR000680" |
| gene-btBTE5EL_001443 | lp42 | pseudo | beta | Pfam | protein_matc | 16 | 304 | 3.2E-47 | + | . | Target=null 16 304;ID=match$1358_16_304;signature_desc=Borrelia lipoprotein;Name=PF00921;status=T;Dbxref="InterPro:IPR000680" |
| gene-btBTE5EL_001444 | lp42 | gene | alpha | Pfam | protein_matc | 53 | 350 | 6.2E-72 | + | . | Target=null 53 350;ID=match$432_53_350;signature_desc=Borrelia lipoprotein;Name=PF00921;status=T;Dbxref="InterPro:IPR000680" |
| gene-btBTE5EL_001445 | lp42 | gene | delta | Pfam | protein_matc | 31 | 315 | 1.8E-86 | + | . | Target=null 31 315;ID=match$488_31_315;signature_desc=Borrelia lipoprotein;Name=PF00921;status=T;Dbxref="InterPro:IPR000680" |
| gene-btBTE5EL_001447 | lp42 | gene | gamma | Pfam | protein_matc | 1 | 277 | 2E-76 | + | . | Target=null 1 277;ID=match$3206_1_277;signature_desc=Borrelia lipoprotein;Name=PF00921;status=T;Dbxref="InterPro:IPR000680" |
| gene-btBTE5EL_001449 | lp42 | gene | beta | Pfam | protein_matc | 20 | 306 | 3.9E-50 | + | . | Target=null 20 306;ID=match$2781_20_306;signature_desc=Borrelia lipoprotein;Name=PF00921;status=T;Dbxref="InterPro:IPR000680" |
| gene-btBTE5EL_001450 | lp42 | gene | alpha | Pfam | protein_matc | 8 | 304 | 2.2E-57 | + | . | Target=null 8 304;ID=match$2249_8_304;signature_desc=Borrelia lipoprotein;Name=PF00921;status=T;Dbxref="InterPro:IPR000680" |
| gene-btBTE5EL_001452 | lp42 | gene | delta | Pfam | protein_matc | 44 | 323 | 1E-83 | + | . | Target=null 44 323;ID=match$3047_44_323;signature_desc=Borrelia lipoprotein;Name=PF00921;status=T;Dbxref="InterPro:IPR000680" |
| gene-btBTE5EL_001457 | lp42 | gene | alpha | Pfam | protein_matc | 7 | 134 | 3.7E-26 | + | . | Target=null 7 134;ID=match$2251_7_134;signature_desc=Borrelia lipoprotein;Name=PF00921;status=T;Dbxref="InterPro:IPR000680" |
| gene-btBTE5EL_001459 | lp42 | gene | gamma | Pfam | protein_matc | 41 | 320 | 2.3E-85 | + | . | Target=null 41 320;ID=match$2490_41_320;signature_desc=Borrelia lipoprotein;Name=PF00921;status=T;Dbxref="InterPro:IPR000680" |
| gene-btBTE5EL_001464 | lp42 | gene | gamma | Pfam | protein_matc | 1 | 279 | 5E-84 | + | . | Target=null 1 279;ID=match$2890_1_279;signature_desc=Borrelia lipoprotein;Name=PF00921;status=T;Dbxref="InterPro:IPR000680" |
| gene-btBTE5EL_001465 | lp42 | gene | delta | Pfam | protein_matc | 44 | 329 | 7.1E-90 | + | . | Target=null 44 329;ID=match$1164_44_329;signature_desc=Borrelia lipoprotein;Name=PF00921;status=T;Dbxref="InterPro:IPR000680" |
| gene-btBTE5EL_001468 | lp42 | gene | alpha | Pfam | protein_matc | 7 | 301 | 5.8E-65 | + | . | Target=null 7 301;ID=match$1916_7_301;signature_desc=Borrelia lipoprotein;Name=PF00921;status=T;Dbxref="InterPro:IPR000680" |
| gene-btBTE5EL_001469 | lp42 | pseudo | beta | Pfam | protein_matc | 16 | 295 | 3.4E-44 | + | . | Target=null 16 295;ID=match$3082_16_295;signature_desc=Borrelia lipoprotein;Name=PF00921;status=T;Dbxref="InterPro:IPR000680" |
| gene-btBTE5EL_001472 | lp42 | gene | alpha | Pfam | protein_matc | 62 | 346 | 1.3E-63 | + | . | Target=null 62 346;ID=match$1057_62_346;signature_desc=Borrelia lipoprotein;Name=PF00921;status=T;Dbxref="InterPro:IPR000680" |
| gene-btBTE5EL_001473 | lp42 | gene | beta | Pfam | protein_matc | 14 | 297 | 4.9E-44 | + | . | Target=null 14 297;ID=match$1936_14_297;signature_desc=Borrelia lipoprotein;Name=PF00921;status=T;Dbxref="InterPro:IPR000680" |
| gene-btBTE5EL_001474 | lp42 | gene | gamma | Pfam | protein_matc | 44 | 329 | 7.3E-85 | + | . | Target=null 44 329;ID=match$548_44_329;signature_desc=Borrelia lipoprotein;Name=PF00921;status=T;Dbxref="InterPro:IPR000680" |
| gene-btBTE5EL_001509 | lp45 | gene | gamma | Pfam | protein_matc | 48 | 333 | 1.3E-88 | + | . | Target=null 48 333;ID=match$1925_48_333;signature_desc=Borrelia lipoprotein;Name=PF00921;status=T;Dbxref="InterPro:IPR000680" |
| gene-btBTE5EL_001511 | lp45 | pseudo | delta | Pfam | protein_matc | 32 | 317 | 3E-83 | + | . | Target=null 32 317;ID=match$1796_32_317;signature_desc=Borrelia lipoprotein;Name=PF00921;status=T;Dbxref="InterPro:IPR000680" |
| gene-btBTE5EL_001513 | lp45 | gene | delta | Pfam | protein_matc | 44 | 335 | 5E-87 | + | . | Target=null 44 335;ID=match$1158_44_335;signature_desc=Borrelia lipoprotein;Name=PF00921;status=T;Dbxref="InterPro:IPR000680" |
| gene-btBTE5EL_001514 | lp45 | gene | beta | Pfam | protein_matc | 55 | 343 | 2.7E-47 | + | . | Target=null 55 343;ID=match$2808_55_343;signature_desc=Borrelia lipoprotein;Name=PF00921;status=T;Dbxref="InterPro:IPR000680" |
| gene-btBTE5EL_001516 | lp45 | gene | alpha | Pfam | protein_matc | 59 | 343 | 1.3E-63 | + | . | Target=null 59 343;ID=match$1716_59_343;signature_desc=Borrelia lipoprotein;Name=PF00921;status=T;Dbxref="InterPro:IPR000680" |
| gene-btBTE5EL_001517 | lp45 | gene | beta | Pfam | protein_matc | 14 | 297 | 2.9E-43 | + | . | Target=null 14 297;ID=match$390_14_297;signature_desc=Borrelia lipoprotein;Name=PF00921;status=T;Dbxref="InterPro:IPR000680" |
| gene-btBTE5EL_001518 | lp45 | gene | gamma | Pfam | protein_matc | 34 | 313 | 2.5E-85 | + | . | Target=null 34 313;ID=match$888_34_313;signature_desc=Borrelia lipoprotein;Name=PF00921;status=T;Dbxref="InterPro:IPR000680" |
| gene-btBTE5EL_001525 | lp46 | pseudo | delta | Pfam | protein_matc | 44 | 202 | 5.1E-38 | + | . | Target=null 44 202;ID=match$2247_44_202;signature_desc=Borrelia lipoprotein;Name=PF00921;status=T;Dbxref="InterPro:IPR000680" |
| gene-btBTE5EL_001525 | lp46 | pseudo | | Pfam | protein_matc | 11 | 84 | 2E-28 | + | . | Target=null 11 84;ID=match$249_11_84;signature_desc=Borrelia lipoprotein;Name=PF00921;status=T;Dbxref="InterPro:IPR000680" |
| gene-btBTE5EL_001526 | lp46 | gene | gamma | Pfam | protein_matc | 34 | 319 | 4.2E-87 | + | . | Target=null 34 319;ID=match$1448_34_319;signature_desc=Borrelia lipoprotein;Name=PF00921;status=T;Dbxref="InterPro:IPR000680" |
| gene-btBTE5EL_001527 | lp46 | pseudo | delta | Pfam | protein_matc | 44 | 184 | 7.7E-39 | + | . | Target=null 44 184;ID=match$2495_44_184;signature_desc=Borrelia lipoprotein;Name=PF00921;status=T;Dbxref="InterPro:IPR000680" |
| gene-btBTE5EL_001527 | lp46 | pseudo | | Pfam | protein_matc | 24 | 159 | 5.3E-40 | + | . | Target=null 24 159;ID=match$1286_24_159;signature_desc=Borrelia lipoprotein;Name=PF00921;status=T;Dbxref="InterPro:IPR000680" |
| gene-btBTE5EL_001529 | lp46 | gene | delta | Pfam | protein_matc | 19 | 297 | 5E-87 | + | . | Target=null 19 297;ID=match$1532_19_297;signature_desc=Borrelia lipoprotein;Name=PF00921;status=T;Dbxref="InterPro:IPR000680" |
| gene-btBTE5EL_001551 | lp46 | pseudo | alpha | Pfam | protein_matc | 62 | 343 | 2.3E-65 | + | . | Target=null 62 343;ID=match$1333_62_343;signature_desc=Borrelia lipoprotein;Name=PF00921;status=T;Dbxref="InterPro:IPR000680" |
| gene-btBTE5EL_001560 | lp46 | pseudo | gamma | Pfam | protein_matc | 34 | 61 | 2.7E-10 | + | . | Target=null 34 61;ID=match$306_34_61;signature_desc=Borrelia lipoprotein;Name=PF00921;status=T;Dbxref="InterPro:IPR000680" |
| gene-btBTE5EL_001560 | lp46 | pseudo | | Pfam | protein_matc | 1 | 114 | 1.6E-23 | + | . | Target=null 1 114;ID=match$1770_1_114;signature_desc=Borrelia lipoprotein;Name=PF00921;status=T;Dbxref="InterPro:IPR000680" |
| gene-btBTE5EL_001560 | lp46 | pseudo | | Pfam | protein_matc | 117 | 192 | 2E-23 | + | . | Target=null 117 192;ID=match$1770_117_192;signature_desc=Borrelia lipoprotein;Name=PF00921;status=T;Dbxref="InterPro:IPR000680" |
| gene-btBTE5EL_001573 | lp49 | pseudo | beta | Pfam | protein_matc | 36 | 164 | 5E-24 | + | . | Target=null 36 164;ID=match$328_36_164;signature_desc=Borrelia lipoprotein;Name=PF00921;status=T;Dbxref="InterPro:IPR000680" |
| gene-btBTE5EL_001574 | lp49 | pseudo | delta | Pfam | protein_matc | 5 | 119 | 7.3E-26 | + | . | Target=null 5 119;ID=match$2115_5_119;signature_desc=Borrelia lipoprotein;Name=PF00921;status=T;Dbxref="InterPro:IPR000680" |
| gene-btBTE5EL_001577 | lp49 | gene | beta | Pfam | protein_matc | 48 | 331 | 7.7E-44 | + | . | Target=null 48 331;ID=match$3173_48_331;signature_desc=Borrelia lipoprotein;Name=PF00921;status=T;Dbxref="InterPro:IPR000680" |
| gene-btBTE5EL_001579 | lp49 | pseudo | gamma | Pfam | protein_matc | 1 | 103 | 3.1E-29 | + | . | Target=null 1 103;ID=match$461_1_103;signature_desc=Borrelia lipoprotein;Name=PF00921;status=T;Dbxref="InterPro:IPR000680" |
| gene-btBTE5EL_001579 | lp49 | pseudo | | Pfam | protein_matc | 33 | 205 | 7.8E-49 | + | . | Target=null 33 205;ID=match$2277_33_205;signature_desc=Borrelia lipoprotein;Name=PF00921;status=T;Dbxref="InterPro:IPR000680" |
| gene-btBTE5EL_001591 | lp49 | gene | delta | Pfam | protein_matc | 44 | 323 | 1.9E-79 | + | . | Target=null 44 323;ID=match$3081_44_323;signature_desc=Borrelia lipoprotein;Name=PF00921;status=T;Dbxref="InterPro:IPR000680" |
| gene-btBTE5EL_001593 | lp49 | gene | delta | Pfam | protein_matc | 34 | 315 | 2E-87 | + | . | Target=null 34 315;ID=match$2416_34_315;signature_desc=Borrelia lipoprotein;Name=PF00921;status=T;Dbxref="InterPro:IPR000680" |
| gene-btBTE5EL_001594 | lp49 | gene | delta | Pfam | protein_matc | 44 | 323 | 2.9E-84 | + | . | Target=null 44 323;ID=match$2876_44_323;signature_desc=Borrelia lipoprotein;Name=PF00921;status=T;Dbxref="InterPro:IPR000680" |
| gene-btBTE5EL_001597 | lp49 | gene | delta | Pfam | protein_matc | 44 | 332 | 1.4E-80 | + | . | Target=null 44 332;ID=match$355_44_332;signature_desc=Borrelia lipoprotein;Name=PF00921;status=T;Dbxref="InterPro:IPR000680" |
| gene-btBTE5EL_001600 | lp49 | gene | delta | Pfam | protein_matc | 19 | 294 | 1.2E-83 | + | . | Target=null 19 294;ID=match$899_19_294;signature_desc=Borrelia lipoprotein;Name=PF00921;status=T;Dbxref="InterPro:IPR000680" |
| gene-btBTE5EL_001603 | lp49 | gene | gamma | Pfam | protein_matc | 34 | 322 | 8.3E-94 | + | . | Target=null 34 322;ID=match$680_34_322;signature_desc=Borrelia lipoprotein;Name=PF00921;status=T;Dbxref="InterPro:IPR000680" |
| gene-btBTE5EL_001604 | lp49 | gene | delta | Pfam | protein_matc | 44 | 311 | 1.1E-78 | + | . | Target=null 44 311;ID=match$2920_44_311;signature_desc=Borrelia lipoprotein;Name=PF00921;status=T;Dbxref="InterPro:IPR000680" |
| gene-btBTE5EL_001607 | lp49 | gene | gamma | Pfam | protein_matc | 34 | 312 | 2.3E-89 | + | . | Target=null 34 312;ID=match$2_34_312;signature_desc=Borrelia lipoprotein;Name=PF00921;status=T;Dbxref="InterPro:IPR000680" |
| gene-btBTE5EL_001608 | lp49 | gene | delta | Pfam | protein_matc | 44 | 325 | 6.3E-85 | + | . | Target=null 44 325;ID=match$1934_44_325;signature_desc=Borrelia lipoprotein;Name=PF00921;status=T;Dbxref="InterPro:IPR000680" |
| gene-btBTE5EL_001611 | lp49 | gene | gamma | Pfam | protein_matc | 44 | 335 | 7.1E-85 | + | . | Target=null 44 335;ID=match$119_44_335;signature_desc=Borrelia lipoprotein;Name=PF00921;status=T;Dbxref="InterPro:IPR000680" |
| gene-btBTE5EL_001612 | lp49 | gene | delta | Pfam | protein_matc | 44 | 334 | 2.8E-83 | + | . | Target=null 44 334;ID=match$2350_44_334;signature_desc=Borrelia lipoprotein;Name=PF00921;status=T;Dbxref="InterPro:IPR000680" |
| gene-btBTE5EL_001613 | lp49 | pseudo | alpha | Pfam | protein_matc | 47 | 341 | 2.4E-64 | + | . | Target=null 47 341;ID=match$366_47_341;signature_desc=Borrelia lipoprotein;Name=PF00921;status=T;Dbxref="InterPro:IPR000680" |
| gene-btBTE5EL_001614 | lp49 | pseudo | beta | Pfam | protein_matc | 16 | 296 | 1.3E-46 | + | . | Target=null 16 296;ID=match$1095_16_296;signature_desc=Borrelia lipoprotein;Name=PF00921;status=T;Dbxref="InterPro:IPR000680" |
| gene-btBTE5EL_001616 | lp49 | gene | delta | Pfam | protein_matc | 8 | 292 | 2.8E-86 | + | . | Target=null 8 292;ID=match$2580_8_292;signature_desc=Borrelia lipoprotein;Name=PF00921;status=T;Dbxref="InterPro:IPR000680" |
| gene-btBTE5EL_001617 | lp49 | gene | delta | Pfam | protein_matc | 44 | 325 | 4E-85 | + | . | Target=null 44 325;ID=match$34_44_325;signature_desc=Borrelia lipoprotein;Name=PF00921;status=T;Dbxref="InterPro:IPR000680" |
| gene-btBTE5EL_001619 | lp49 | pseudo | alpha | Pfam | protein_matc | 41 | 183 | 2.3E-24 | + | . | Target=null 41 183;ID=match$774_41_183;signature_desc=Borrelia lipoprotein;Name=PF00921;status=T;Dbxref="InterPro:IPR000680" |
| gene-btBTE5EL_001619 | lp49 | pseudo | | Pfam | protein_matc | 4 | 143 | 1.4E-33 | + | . | Target=null 4 143;ID=match$494_4_143;signature_desc=Borrelia lipoprotein;Name=PF00921;status=T;Dbxref="InterPro:IPR000680" |
